# Evaluation of the stereochemical quality of predicted RNA 3D models in the RNA-Puzzles submissions

**DOI:** 10.1101/2021.08.22.457258

**Authors:** Francisco Carrascoza, Maciej Antczak, Zhichao Miao, Eric Westhof, Marta Szachniuk

**Affiliations:** Institute of Computing Science and European Centre for Bioinformatics and Genomics, Poznan University of Technology, 60-965 Poznan, Poland; Institute of Bioorganic Chemistry, Polish Academy of Sciences, 61-704 Poznan, Poland; European Molecular Biology Laboratory, European Bioinformatics Institute (EMBL-EBI), Wellcome Genome Campus, CB10 1SD, UK; Translational Research Institute of Brain and Brain-Like Intelligence and Department of Anesthesiology, Shanghai Fourth People’s Hospital Affiliated to Tongji University School of Medicine, Shanghai 200081, China; Université de Strasbourg, Institut de biologie moléculaire et cellulaire CNRS, Architecture et Réactivité de l’ARN, 12 allée Konrad Roentgen, 67084 Strasbourg, France

**Keywords:** stereochemistry, quality validation, RNA structure, 3D structure prediction, RNA-Puzzles

## Abstract

*In silico* prediction is a well-established approach to derive a general shape of an RNA molecule based on its sequence or secondary structure. This paper reports an analysis of the stereochemical quality of the RNA three-dimensional models predicted using dedicated computer programs. The stereochemistry of 1,052 RNA 3D structures, including 1,030 models predicted by fully automated and human-guided approaches within 22 RNA-Puzzles challenges and reference structures, is analysed. The evaluation is based on standards of RNA stereochemistry that the Protein Data Bank requires from deposited experimental structures. Deviations from standard bond lengths and angles, planarity or chirality are quantified. A reduction in the number of such deviations should help in the improvement of RNA 3D structure modelling approaches.

## INTRODUCTION

Knowledge of the RNA atomic structure is crucial to address biological problems, therefore computational tools for the prediction of RNA three-dimensional models from sequence, have been developed to help or bypass some hurdles of laboratory procedures (Miao and Westhof 2017; Lukasiak et al. 2015; Magnus et al. 2020; Gumna et al. 2020; Li et al. 2020).

The first decade of the 21st century resulted in several computer programs and protocols, which paved the way for automated modelling of RNA 3D structures: S2S (Jossinet and Westhof 2005), FARFAR (Das and Baker 2007), iFoldRNA (Ding et al. 2008), MC-Fold/MC-Sym (Parisien and Major 2008), NAST (Jonikas et al. 2009). Some of them developed into highly specialized programs, which are used for either fully automatic or human-guided prediction. In the following years, this collection grew to include other tools such as ModeRNA (Rother et al. 2011), RNAComposer (Popenda et al. 2012), 3dRNA (Zhao et al. 2012), Vfold (Xu et al. 2014), and SimRNA (Boniecki et al. 2016).

To stimulate the improvement of quality in RNA prediction, RNA-Puzzles was organized ten years ago (Cruz et al. 2012). RNA-Puzzles is a community-wide assessment of RNA 3D structure prediction that aims to understand the bottlenecks in current RNA 3D structure prediction to promote the improvement of prediction methods. Before the publication of an experimentally determined RNA structure, the sequence is disseminated among the community and prediction results are submitted within 3-4 weeks. Assessment against the experimental structure is performed after the release of the structure. There are two categories of challenges depending on the protocols used to obtain the models: they can originate from fully automated webservices or human experts running various prediction programs. The starting point for each challenge is a novel experimentally determined RNA 3D structure, the conformation of which is unknown to the predictors. The webservers have 48 hours and human experts 3-4 weeks for submitting their models. After the deadline, the predictions are evaluated, and the results are published with the ranking of the submitted models. Presently, 28 crystallographic structures have been part of the contest. Eighteen of them have been the basis of four scientific papers published by the RNA-Puzzles community (Cruz et al. 2012; Miao et al. 2015; Miao et al. 2017; Miao et al. 2020). As of October 2020, 22 challenges have been concluded with assessment results available on the RNA-Puzzles website (rnapuzzles.org). It provides accuracy assessments determined by comparison to the reference structure and calculation of several global similarity and distance measures (Magnus et al. 2020): root-meansquare-deviation (RMSD) (Kabsch 1978), Deformation Index (DI) that normalizes RMSD with the sequence length (Parisien et al. 2009), Interaction Network Fidelity (INF) including Watson-Crick, non-canonical, and stacking interactions (Parisien et al. 2009), and – more recently – Mean of Circular Quantities operating in torsion angle space (Zok et al. 2014; Wiedemann et al. 2017). RMSD serves as the main criterion to rank the predicted models, although it is only capable of assessing the minimum average distance between two 3D structures represented as two sets of atomic coordinates. The remaining metrics allow a focus on base-pairs and torsion angles. Additionally, RNA-Puzzles use the Clashscore – as defined in the MolProbity software package (Williams et al. 2018) – for assessing the accuracy in a non-comparative procedure by finding overlapping or too close atoms in the models and used as an overall evaluation of the stereochemistry.

Nevertheless, current biological problems are setting new thresholds of what acceptable geometry qualities should be. Catalytic features, for instance, highlight that not only the model’s geometry is important but that its stereochemistry is an important factor as well. One example is the torsion-angle based dependence between active and non-active conformation of base pairs in some ribozyme active site (White et al. 2018). Moreover, self-cleaving ribozymes can provide another example, in which the correct description of phosphate backbone stereochemistry is critical to assess correctly the reaction pathways of these mechanisms. (Teplova et al. 2020). Yet another recent case is drug development that targets RNA (e.g., against viruses (Aftab et al. 2020)). Therefore, there is a clear need to advance technology to provide useful and trustable tools capable to address these challenges.

Proper stereochemistry is at the core of biomolecular structure modelling. The geometries and stereochemistry of the nucleic acid building blocks are very well known and with high precision (Clowney et al. 1996; Gelbin et al. 1996; Schneider et al. 1996). Inaccuracies in molecular geometry can result from geometry optimizations that fall into local minima that may lead to a metastable conformer, different form the native one or another biologically irrelevant conformation. Inappropriate geometry may mask incorrect choice in torsion angle space (for example, a base incorrectly in the *syn* conformer can lead to geometrical distortions in the sugar-phosphate backbone). Biomolecular structures are extremely well fine-tuned and the whole variety of physicochemical interactions is exploited in the folded native structure. Neglect of some type of interactions, or an inappropriate calibration, can lead to wrong conformations that can produce molecular distortions under insufficiently controlled structural refinement (Popenda et al. 2021).

Here, we revisit the evaluation of the stereochemistry of predicted models beyond interatomic non-covalent distances. We follow a routine recommended to experimenters who deposit their structure data in the Protein Data Bank (Berman et al. 2000) and the Biological Magnetic Resonance Data Bank (Ulrich et al. 2008) – both contributing to the wwPDB partnership (Berman et al. 2003). wwPDB stresses the importance of careful examination of structures by providing tools that set the standards for 3D structure submission. In 2017, it introduced OneDep – a unified system applying the deposition, biocuration, and validation pipelines for structural data (Gore et al. 2017; Young et al. 2017). OneDep is an extensive suite of programs operating on different metrics to assess the accuracy of structures. It implements stereochemistry analysis through MAXIT (Feng et al. 1998; Berman et al. 2000).

To evaluate the stereochemistry of RNA tertiary structure predictions, we analysed the results of all RNA-Puzzles challenges with the standardized data available as of November 2020 – *i.e.,* Puzzles 1-15, 17-21, and 24 (Puzzle 14 in two versions, bound 14a and free 14b) – and 22 corresponding reference structures. We downloaded 1,030 predicted RNA models from the standardized dataset belonging to RNA-Puzzles resources (located at https://github.com/RNA-Puzzles; Magnus et al. 2020). Among those, 797 models were in the human category and 233 models in the webserver category. From these data, we created 23 clusters by participants – each containing models submitted by a single human group or a webserver (Table 1). An additional 24th cluster included the reference structures (Table 2). Next, we processed the structures in all of these subsets using MAXIT software (Feng et al. 1998; Berman et al. 2000) and compare results with the MolProbity software (Williams et al. 2018). Next, we used Barnaba (Bottaro et al. 2019) and X3DNA-DSSR (Lu and Olson 2003) to verify base-paring geometries and handedness of helices, respectively. Finally, we conducted a simple statistical analysis by computing the average value, standard deviation, and median for every subset including more than one model (see Materials and Methods section).

**Table 1.** Clusters of RNA 3D models predicted within RNA-Puzzles organized by participants.

| Category | Cluster | Name of the group/ server | Number of challenges | Total number of predicted models | Puzzle (challenge) numbers |
| --- | --- | --- | --- | --- | --- |
| human groups | H1 | Adamiak | 14 | 64 | 4,5,7,8,11-15,17,19-21,24 |
|  | H2 | Bujnicki | 20 | 131 | 1-14,17,19-21,24 |
|  | H3 | Chen | 20 | 145 | 1-19,21 |
|  | H4 | Das | 21 | 188 | 1-14,17-21,24 |
|  | H5 | Ding | 12 | 124 | 7-9,11-14,17-19,24 |
|  | H6 | Dokholyan | 14 | 52 | 1-10,13,17-19 |
|  | H7 | Kollmann | 1 | 10 | 24 |
|  | H8 | Lee | 1 | 5 | 18 |
|  | H9 | Major | 7 | 32 | 1-4,6-7,17 |
|  | H10 | Mikolajczak | 1 | 1 | 4 |
|  | H11 | Sanbonmatsu | 1 | 4 | 21 |
|  | H12 | Santalucia | 3 | 3 | 1,2,4 |
|  | H13 | Weeks | 1 | 3 | 12 |
|  | H14 | Wildauer | 1 | 1 | 2 |
|  | H15 | Xiao | 6 | 33 | 5,11-13,17,20 |
|  | H16 | Yagoubali | 1 | 1 | 18 |
| webservers | W1 | 3dRNA | 5 | 30 | 15,18-19,21,24 |
|  | W2 | FARFAR | 6 | 70 | 15,18-21,24 |
|  | W3 | iFoldRNA | 1 | 5 | 24 |
|  | W4 | LeeServer | 2 | 10 | 18-19 |
|  | W5 | RNAComposer | 7 | 49 | 15,17-21,24 |
|  | W6 | SimRNA | 7 | 59 | 15,17-21,24 |
|  | W7 | Vfold | 1 | 10 | 24 |

**Table 2.** Cluster with the reference structures and their prediction-related data.

| No | Puzzle id | PDB id | Number of predicted models |  |  | Number of participants |  |  |
| --- | --- | --- | --- | --- | --- | --- | --- | --- |
|  |  |  | Humans | Webservers | Total | Humans | Webservers | Total |
| 1 | PZ01 | 3MEI | 14 | 0 | 14 | 6 | 0 | 6 |
| 2 | PZ02 | 3P59 | 13 | 0 | 13 | 7 | 0 | 7 |
| 3 | PZ03 | 3OWZ | 12 | 0 | 12 | 5 | 0 | 5 |
| 4 | PZ04 | 3V7E | 30 | 0 | 30 | 8 | 0 | 8 |
| 5 | PZ05 | 4P9R | 25 | 0 | 25 | 7 | 0 | 7 |
| 6 | PZ06 | 4GXY | 34 | 0 | 34 | 5 | 0 | 5 |
| 7 | PZ07 | 4R4V | 52 | 0 | 52 | 7 | 0 | 7 |
| 8 | PZ08 | 4L81 | 42 | 0 | 42 | 6 | 0 | 6 |
| 9 | PZ09 | 5KPY | 34 | 0 | 34 | 5 | 0 | 5 |
| 10 | PZ10 | 4LCK | 26 | 0 | 26 | 4 | 0 | 4 |
| 11 | PZ11 | 5IEM | 53 | 0 | 53 | 6 | 0 | 6 |
| 12 | PZ12 | 4QLM | 51 | 0 | 51 | 7 | 0 | 7 |
| 13 | PZ13 | 4XW7 | 55 | 0 | 55 | 7 | 0 | 7 |
| 14 | PZ14a | 5DDO | 61 | 0 | 61 | 5 | 0 | 5 |
| 15 | PZ14b | 5DDO | 52 | 0 | 52 | 5 | 0 | 4 |
| 16 | PZ15 | 5DI4 | 20 | 54 | 74 | 2 | 4 | 6 |
| 17 | PZ17 | 5K7C | 69 | 38 | 107 | 8 | 2 | 10 |
| 18 | PZ18 | 5TPY | 24 | 28 | 52 | 6 | 5 | 11 |
| 19 | PZ19 | 5T5A | 26 | 28 | 54 | 6 | 5 | 11 |
| 20 | PZ20 | 5Y85 | 20 | 20 | 40 | 4 | 3 | 7 |
| 21 | PZ21 | 5NZ6 | 34 | 25 | 59 | 5 | 4 | 9 |
| 22 | PZ24 | 6OL3 | 50 | 40 | 90 | 5 | 6 | 11 |

## RESULTS

For each model, MAXIT returned a report of abnormal stereochemical parameters (falling into six categories: close contacts, bond length deviations, bond angle deviations, deviation from planarity, chirality issues, and phosphate bond linkages (Supplementary Materials include tables with the error numbers in every model). Using MAXIT we examined them first for the subset of 22 reference structures (Figure 1 and Supplementary Table S1). Most of them contained some types of geometrical deviations from standard dictionaries. We found the highest incidence of errors in the bond angles (183 errors in 17 structures), followed by close contacts (54 errors in 7 structures) and bond lengths (32 errors in 5 structures). Among the worst cases (PZ07, PZ01, and PZ21), two are for structures at a resolution worse than 2.5Å *(c.f.* Figure S1 in the Supplementary Materials). The software X3DNA-DSSR (Lu and Olson 2003) does not reveal any left-handed helix/dinucleotide step in RNA-Puzzles submissions nor experimentally determined RNA 3D structures (c.f. Tables S46-S69 in the Supplementary Materials). In Figure 1, one can also observe that there are no chirality issues, while deviations from planarity occur only in 2 instances (9 errors in total). For polymer linkage (i.e. deviations in P-O bond lengths), we found 7 structures with a total of 9 reported inaccuracies, making an average of less than one error per structure, the same as for errors in planarity.

**Figure 1.**
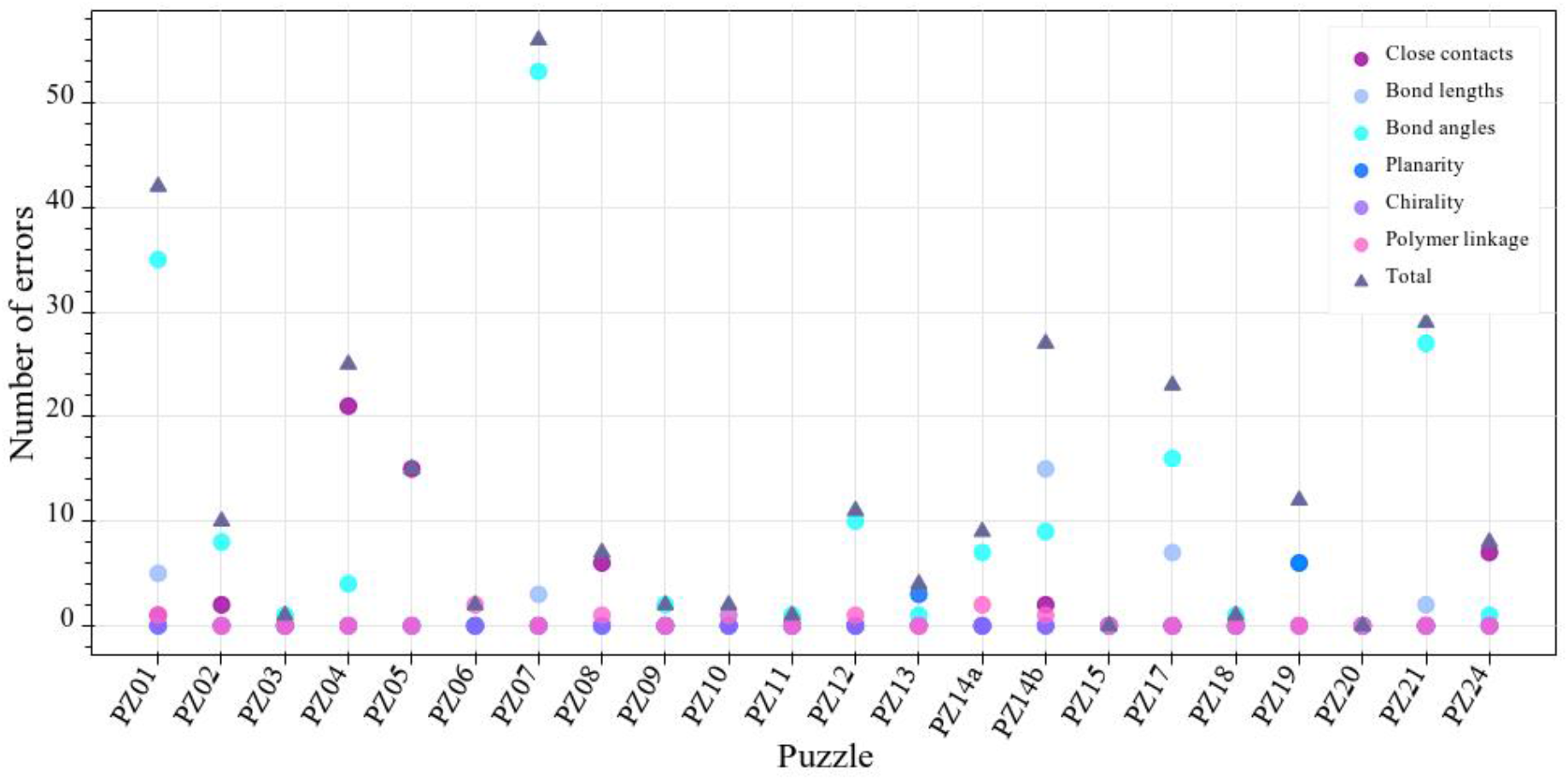
Stereochemical errors reported by MAXIT for the reference structures.

We have analysed separately clusters with models predicted by human experts and webservers. Each of these 23 collections contains the predictions submitted by one participant within all the considered challenges available in the standardized dataset of RNA-Puzzles resource *(c.f.* Table 1). Their cardinalities range from 1 to 188. Within each of these clusters, except for those including only one model *(i.e.,* H10, H14, and H16), we determined the total number of errors of each type (Tables S2-S24 in the Supplementary Materials), the average number over all the errors and the standard deviation (Figure 2) and confirmed these results using MolProbity software version 4.5.1 (Williams et al. 2018) (Figures S2-S4 and Tables S71-S92 in the Supplementary Material). We did the same for each of the six types of stereochemical properties – we further computed the average value of each error and the standard deviation per cluster (Figure 3). One can observe that some of the applied prediction methods have an advantage over others in terms of the total number of errors. However, most submissions have stereochemical issues to address. Interestingly, there is no visible difference between the quality of human versus webserver predictions as far as the average number of all the inaccuracies is concerned. In both categories, we can observe both good and bad scores. The average number of errors per model in the human category equals 106, while in the webserver category it is 103.

**Figure 2.**
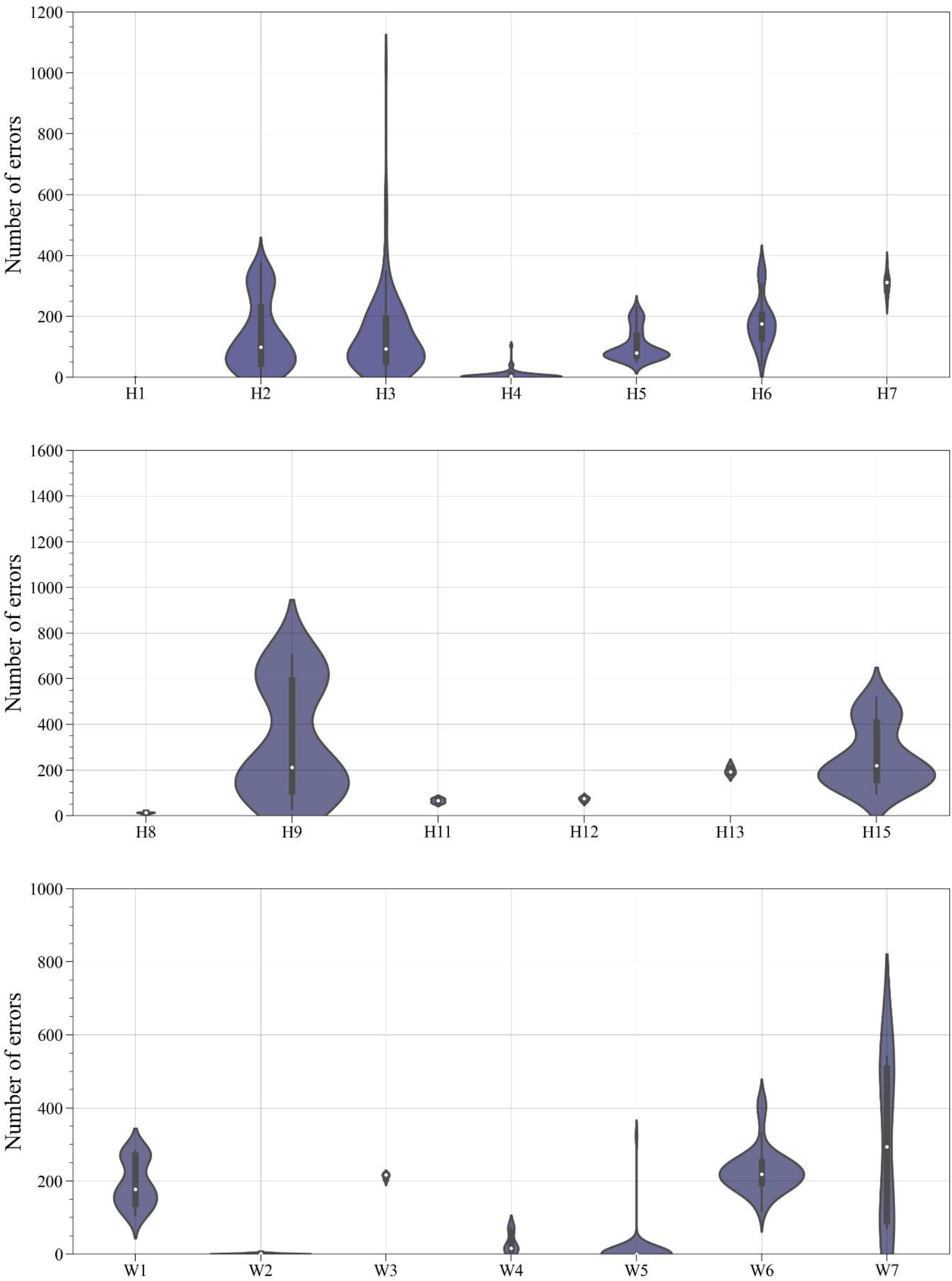
The number of stereochemical errors identified in all the considered models by participants. The white dot at the centre of every violin plot represents a median. The black bar corresponds to the interquartile range. The first and the third quartile are represented as wicks up and down from the interquartile range. The violin shape shows error distribution.

**Figure 3.**
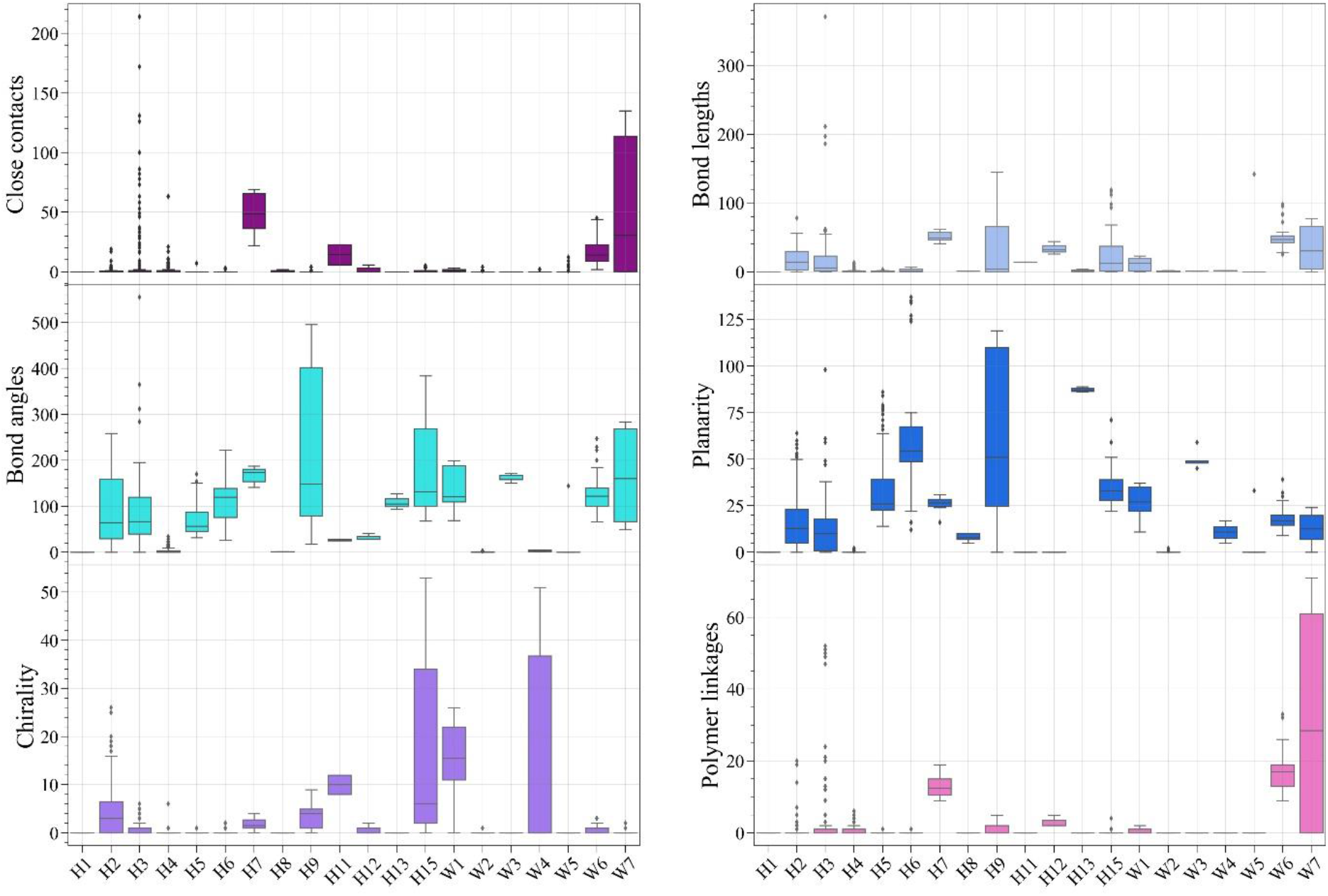
The number of identified stereochemical errors per error type and participant. The box plot of each participant shows the interquartile range. The black middle line in every box depicts a median. The first and third quartile are represented as wicks up and down from the interquartile range. Separated dots outside boxes correspond to outliers.

Due to the significant difference between cluster cardinalities, there is no statistical consistency between them, but there is statistical consistency within each cluster – the results of a single participant. For instance, the H9 set, for which the total score is significant in Figure 2, has only 32 items – we should remember that in a small set, one highly defective object significantly affects the average value and standard deviation. The most numerous clusters (over 100 models) are H2, H3, H4, and H5. The sets labelled as H1, H6, W2, and W6 include 50-100 items. The remaining ones have less than 50 models each. By clustering and comparing the predictions submitted, one can observe that H1 (Average Total Number of errors, ATN=0), H4 (ATN=7.69), and H8 (ATN=11.60) groups apply methods performing the best in the category of human experts; W2 (ATN=1.30), W5 (ATN=7.61), and W4 (ATN=31.20) are most successful among the webservers – their average number of all errors per model is less than 50 (Figure 2). For clusters H2, H3, H5, H6, H11, H12, H13, and W1, the average total number of inaccuracies is in the range 50-200. However, the significant standard deviations indicate a large spread in stereochemical issues for the prediction methods used to obtain the models collected in these clusters. In the other clusters, the average total number of errors falls in the range 200-350 (if we do not consider single-model clusters).

A comparison between Figures 1 and 2 shows the gap between reference structures and predicted models. The most notable conformational errors in predicted models occur in bond lengths and angles. On average, MAXIT has identified over a hundred of such inaccuracies per the predicted RNA 3D model and less than 10 per the reference structure (on average). Chirality (or incorrect sugar substituent) is correct in the experimentally determined RNAs, while 28% of the predicted models have problems with it. Quite many deviations from the average planes of aromatic rings are observed. Polymer linkage assesses bond lengths between the adjacent nucleotides by measuring the P-O bond distances. This parameter has the lowest error rate in computationally generated structures – errors of this type occurred in 22% of all analysed RNAs.

Figure 3 presents MAXIT results separately for each error type and allows to take a closer look into the weaknesses of the protocols embedded within various prediction programs. The plots reveal the highest number of inaccuracies especially in bond angles (70,184). Virtually every prediction method generates numbers of errors in covalent geometries, and the exceptional models with no such issue are not necessarily the most similar to the reference structures(s) in terms of overall RMSD. In the human category, models collected in H1, H4, and H8 clusters have little or no geometric issues (although, at the same time, four H4 models in Puzzle 19 have the largest bond length error with 05’–C5’ length >100Å), while predictions in H7, H9, and H15 are among those with the highest number of inaccuracies. In the webserver category, W2, W4, and W5 perform the best as far as bond lengths and angles are concerned, while W3, W6, and W7 are at the end of the ranking. If we consider deviations from ring planarity (Figure 3), of which the total number is 17,594, their average per model for every cluster is below 90 errors. Models in H6, H9, and H13 have a significant number of these issues. The largest identified deviation from planarity equals 0.791 Å and occurred in H15 model 1 predicted for Puzzle 12. An example error of this type is depicted in Figure S5.

The average number of chirality errors is below 20 for all clusters. For some clusters, MAXIT reported zero or one issue of this type in total: H1, H8, and H13 within the human category, W2, W3, and W5 within the webservers. Let us add that for H4 – having the longest track of submissions – the number of errors in this category is also negligible. Some approaches (H2, H15, and W4) scored higher as far as the average number of chirality inaccuracies is concerned.

In total, 2,130 abnormalities classified by MAXIT as chirality errors occurred in 291 predicted models. The most common form of chiral error is the interchange of the hydroxyl group and hydrogen atom on the same carbon atom at the ribose moiety (Figure 4). Such an interchange does not lead to a chiral error, it produces another sugar type (for example arabinose or xylose instead of ribose). Such improper sugar construction represents 94.9% of all chiral errors identified by MAXIT. The remaining 5.1% are planar inaccuracies in the sugar ring, and they occur when the improper torsion angle at a sp^3^ carbon atom is close to zero instead of being around −122 or +122 degrees. Such a situation occurs in the furanose ring with distorted or flat sugar rings.

**Figure 4.**
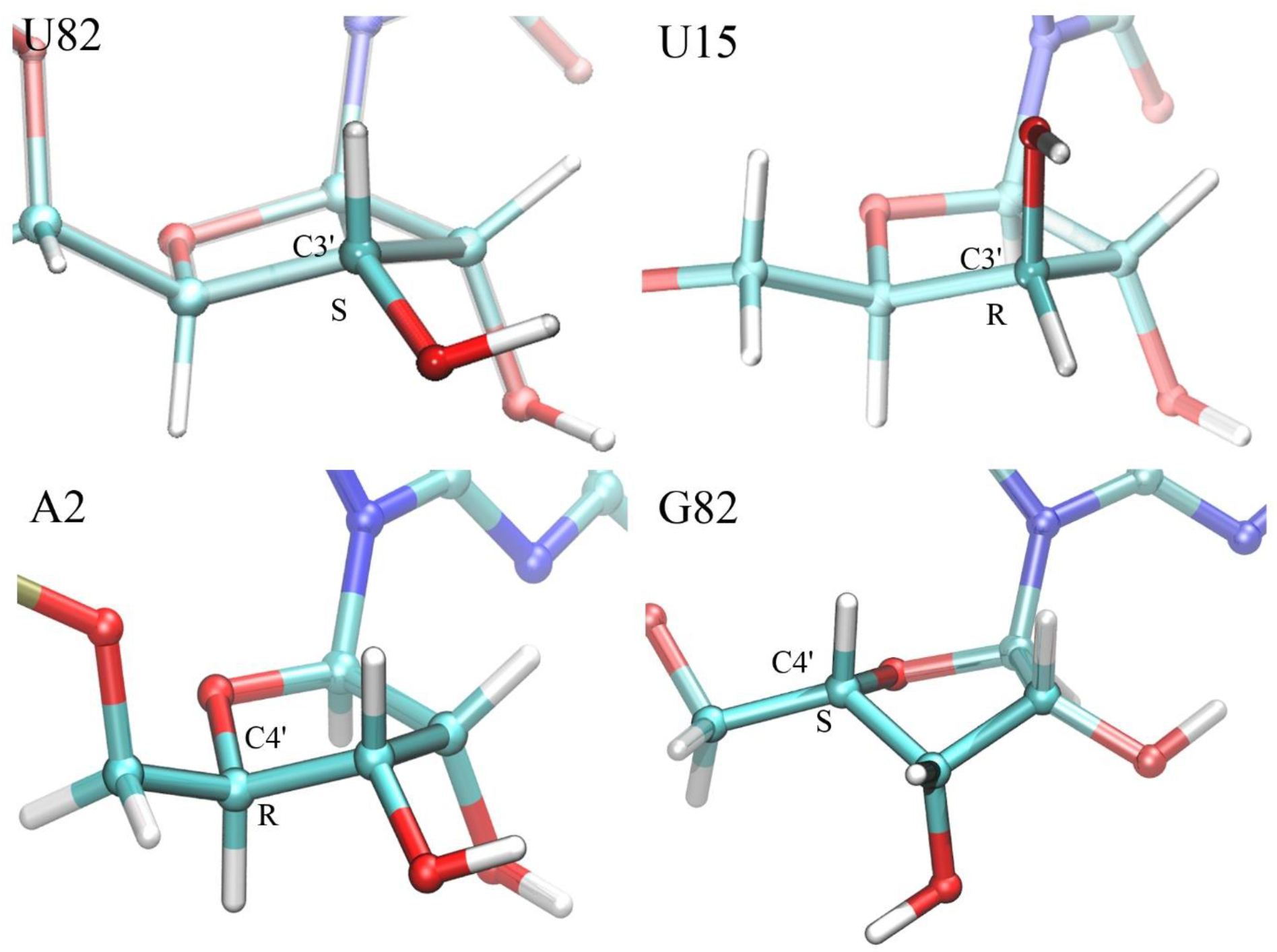
Example chiral errors in H3 model from PZ07 (top) and H4 model from PZ06 (bottom). Top left: C3’ atom in U82 with correct chiral centre; top right: U15 with incorrect chiral inversion at carbon atom C3’, actually changing a ribose to a xylose moiety. Bottom left: A2 with correct chirality at C4’; bottom right: G82 with incorrect chirality at C4’.

A distribution of chiral errors among nucleotides is shown in Table 3. We can see a high frequency in Guanine (692 inaccuracies, which make 32.5% of all chiral errors) and a lower one for Uracil (410 inaccuracies, which makes 19.2% of all chiral errors). This relationship is visible for both sugar construction inversions and planar errors. However, the frequencies are affected by the nucleotide content in the analysed RNA structures. Thus, in Table 3, we also present the total number of Adenines, Cytosines, Guanines, and Uracils in the analysed dataset, and the percentage of these nucleotides having erroneous chirality. Let us add that among all nucleotides with chiral errors 91% are *anti*-while 9% are *syn* nucleotides. A similar distribution is observed for each of the individual nucleotide types. *Syn/anti* conformation characterizes a relative orientation of base and sugar and is determined based on the chi torsion angle (defined by O4’-C1’-N1-C2 for pyrimidines and O4’-C1’-N9-C4 for purines). Usually, chi falls into the ranges [+90,+180] or [-180,-90] corresponding to the *anti* conformation. Occasionally, we observe its value in [-90,+90], which refers to the *syn* conformation. Some chiral errors (11%) appear when the conformation of a nucleotide in the predicted model differs from that in the reference structure. However, in most cases (89%), these errors cannot result from the conformation change (Table 4). Regarding the distribution of errors among chiral atomic centres, we can observe that 43% of inaccuracies occur at C4’ *(c.f.* Figure 4), around 25% at C2’ and C3’ *(c.f.* Figure 5), and only 5.9% at C1’ atom (Table 5).

**Table 3.** Chiral or planar sp3 atom errors by nucleotide type. The percentage of erroneous nucleotides of a given type is calculated about all nucleotides in the set of analysed models.

| Nucleotide | Nucleotide<br>content in<br>the dataset | Number of erroneous nts |  |  | Percentage of erroneous nts [%] |  |  |
| --- | --- | --- | --- | --- | --- | --- | --- |
|  |  | Total | Reversed<br>geometry | Planar<br>sugar | Total | Reversed<br>geometry | Planar<br>sugar |
| A | 1,184,711 | 585 | 554 | 31 | 0.049 | 0.047 | 0.002 |
| C | 1,198,575 | 443 | 419 | 24 | 0.037 | 0.035 | 0.002 |
| G | 1,679,526 | 692 | 655 | 37 | 0.041 | 0.039 | 0.002 |
| U | 872,633 | 410 | 393 | 17 | 0.047 | 0.045 | 0.002 |
| Total | 4,935,445 | 2,130 | 2,021 | 109 | 0.043 | 0.041 | 0.002 |

**Table 4.** Chiral or planar sp^3^ atom errors by nucleotide type and conformation (*anti* or *syn*) depending on whether it is different or the same as in the reference structure. The percentage of erroneous nucleotides of a given type and conformation is calculated about all (2,130) erroneous nucleotides in the set.

| Nucleotide | Number of erroneous nts |  |  | Percentage of erroneous nts [%] |  |  |
| --- | --- | --- | --- | --- | --- | --- |
|  | Total | Same conformation | Changed conformation | Total | Same conformation | Changed conformation |
| A | 585 | 524 | 61 | 27.46 | 24.60 | 2.86 |
| C | 443 | 405 | 38 | 20.80 | 19.01 | 1.79 |
| G | 692 | 623 | 69 | 32.49 | 29.25 | 3.24 |
| U | 410 | 339 | 71 | 19.25 | 15.92 | 3.33 |
| Total | 2,130 | 1,891 | 239 | 100 | 88.78 | 11.22 |

**Table 5.** Chiral or planar sp3 atom errors. An inversion at C1’ would replace a beta-ribose nucleotide into an alpha-ribose nucleotide, an inversion at C2’ leads to an arabinose instead of a ribose, an inversion at C3’ leads to a xylose instead of a ribose, an inversion at C4’ would lead to a real chiral inversion from a D-ribose to a L-ribose.

| Atom | C1' | C2' | C3' | C4' | Total |
| --- | --- | --- | --- | --- | --- |
|  | Beta/Alpha | R/Arabino | R/Xylose | D/L |  |
| Number of errors | 163 | 521 | 589 | 857 | 2,130 |
| Percentage of errors [%] | 7.65 | 24.46 | 27.65 | 40.23 | 100 |

Polymer linkage errors are rare for most predicted models (Figure S6). We have found that MAXIT may report false-positives in this category – whenever it comes to the truncation of the sequence in the model, MAXIT fails to recognize different chains properly. This kind of artifact is clear for the reference structures *(e.g.,* PZ12 and PZ21). However, when it comes to RNA 3D models predicted by webservers or human experts, we have not found such false-positives since different chains are labelled correctly by the prediction methods (about 10% of submissions contain double-chain models). Thus, polymer linkage inaccuracies depicted in Figure S6 are true-positive errors, and they come from incorrect linkage bond length between oxygen and phosphate group of two neighbouring nucleotides in the polymer chain (Figure S6). Such errors were suspected to occur during the assembly building process of RNA fragments since generally nucleotides start at 5’-P and ends at O3’, however, our analysis revealed that models predicted by assembly-based methods did not show errors of this type.

The analysis of the RNA-Puzzles data set containing 1,052 RNAs reveals 2,431 polymer linkage errors, including 2,422 errors in 230 predicted models and 9 errors in 7 reference structures. H2 and H7 clusters have the highest average number of these errors among human expert predictions. For the webservers, MAXIT has found the highest number of this type of inaccuracy in W6 and W7. The remaining prediction methods do not tend to generate errors in this category. By default, MAXIT reports such error whenever the distance between oxygen atom O3’ and phosphorus atom P of the next nucleotide in the polymer chain is longer than a typical length of a covalent bond between these atoms (Schneider et al. 1996). Some of these errors are small deviations, but major ones occur as well. For example, in H15 model 3 predicted within Puzzle 13, MAXIT identified a bond of length 82.52Å between A70 and A71 – it is the highest inaccuracy of this type identified within the dataset. The distribution of errors by the nucleotide type and *syn/anti* conformation is presented in Table 6. One can observe that linkage errors involving Adenine (19%) are least frequent, and those with Guanine (32%) occur most often. However, as a function of the relative contents of the four nucleotides in the analysed RNA molecules, Cytosine has the most linkage errors and Adenine the least.

**Table 6.** Polymer linkage errors by the types and conformation (*anti* or *syn*) of nucleotides depending on whether they have same or different conformation as in the reference structure. The percentage of erroneous linkages of a given type and conformation is calculated about all (2,422) erroneous linkages in the set.

| Linkage | Number of erroneous linkages |  |  |  | Percentage of erroneous linkages [%] |  |  |  |
| --- | --- | --- | --- | --- | --- | --- | --- | --- |
|  | Total | Conformation* |  |  | Total | Conformation* |  |  |
|  |  | Same | 1 changed | 2 changed |  | Same | 1 changed | 2 changed |
| A(O3')–P | 461 | 378 | 72 | 11 | 19.03 | 15.61 | 2.97 | 0.45 |
| C(O3')–P | 692 | 652 | 38 | 2 | 28.57 | 26.92 | 1.58 | 0.07 |
| G(O3')–P | 783 | 714 | 60 | 9 | 32.33 | 29.48 | 2.48 | 0.37 |
| U(O3')–P | 486 | 428 | 41 | 17 | 20.07 | 17.67 | 1.69 | 0.71 |
| Total | 2,422 | 2,172 | 211 | 39 | 100 | 89.68 | 8.71 | 1.61 |

\* Conformation “same” – both residues in the linkage have the same conformations in the predicted model and the reference structure; “1 changed” – one of the predicted residues is in different conformation than in the reference structure; “2 changed” – both predicted residues are in different conformation than in the reference structure.

Then we analysed the dataset with the Barnaba software (Bottaro et al. 2019), we computed the Backbone Root Mean Square Deviation (herein called BBRMSD) and Basepairing Interactions Root Mean Square Deviations (eRMSD) for nucleobases. BBRMSD can be interpreted as the quality of the backbone structures expressed in RMSD values. eRMSD was taken to measure the quality of base pairs in the structures. Both values were computed using the reference structure given by the RNA-Puzzles organizers.

eRMSD values below 8Å are considered as low. For values less than 5Å, both the reference structure and target structure are very similar (Bottaro et al. 2019).

Our results (see Figure S7A) show that across all the puzzles, there is no sensitive trend in the average measurement of both BBRSMD and eRMSD. In the average RMSD values across participants (see Figure S7B), groups H1 to H4 and H12, performed better than the automated protocols, with exception of W7 which showed the lowest average in both, backbone and base-pairs quality. Some groups, from H6 to H9, have lower eRMSD than BBRMSD values, suggesting these groups focus their attention into the base pairs rather than the backbone structure. Other groups, like H11 and W5, perform better at deducing the volumetric backbone shape, but they have relatively worst base-pairs performance (Figure S7B).

## DISCUSSION

Stereochemical errors in the predicted RNA 3D models are primarily generated by computer programs used in both human-guided and webserver prediction. In many cases, these may be the result of relatively small rounding errors appearing at one of the calculation steps and propagating in subsequent iterations. Knowing the general approach used by the prediction method, it is possible to indicate the most sensitive stages at which the errors arise. Computational complexity is the main problem in *de novo* simulation of RNA folding. Therefore, different techniques are used to reduce the time cost of *de novo* prediction methods.

One of them is to evaluate the fold using simplified potentials, which do not take stereochemical parameters into account. The simulation *(e.g.,* Monte Carlo) converges towards an optimum defined in terms of the overall 3D shape of the molecule and gives a fold that is stereochemically oversimplified and far from ideal. The large number of calculations that are performed during the random sampling of the solution space also affects the generation of errors, as the errors that occur in one step are propagated further and often deteriorate the final solution. Another related problem concerns coarse-grained simulations. The transformation of a coarse-grained model to a full-atom model is a highly erroneous procedure.

In template-based approaches (homology modeling, fragment assembly methods), a crucial moment is the choice of the right template/fragment. A model based on a stereochemically erroneous template or of low-resolution may incorporate the incorrect stereochemical parameters. Stereochemical errors may also arise during nucleobase exchange, structural blocks insertion, or their assembly into a larger whole. In the latter case, the choice of the structural blocks that are rotated and translated is critical, since maneuvering a larger element is more erroneous.

Errors that arise during the structure modeling process can be avoided when applying a function that validates partial solutions based on their stereochemical parameters. However, it is very time-consuming and – in the case of *de novo* methods – completely unprofitable since it may cause the method not to return the solution in a reasonable time. Therefore, the best solution to the problem is to improve stereochemistry post-factum by minimizing the geometry or the energy after building the model, but bad local geometries are not easy to relieve when embedded in a large fold. At the same time, re-refining the templates used with standard dictionaries may alleviate the propagation of errors, leading to tight conformers with stereochemical errors difficult to energy minimize. In some tools, like FARFAR (Das and Baker 2007; Das et al. 2010) and RNAComposer (Popenda et al. 2012; Purzycka et al. 2014; Antczak et al. 2016), such a procedure has been implemented and successfully fulfills its role.

Erroneous bond lengths, bond angles, and planarity deviations are the most frequent errors in RNA 3D structure prediction, while incorrect sugar constructions or chirality and polymer linkage errors occurring less frequently (~10 issues per structure on average). Falsepositive errors, which are caused by improper identification of structural chains in multi-chain RNA structures, are found in the polymer linkage category of the MAXIT results. Most errors can be compensated by running energy minimization protocols – *e.g.,* CYANA (Güntert and Buchner 2015), NAMD (Philips et al. 2020), XPLOR-NIH (Schwieters et al. 2003) – for the preliminary models or ensuring a proper stereochemistry from the early stages of prediction. One can also process the predicted RNA structures using tools – for example, RNAfitme (Antczak et al. 2018; Zok et al. 2015) or QRNAS (Stasiewicz et al. 2019) – having the potential to refine the nucleic acid structure.

## CONCLUSIONS

We found that most RNA 3D structure prediction methods evaluated within RNA-Puzzles – either in human or webserver category – generate models with some incorrect stereochemical parameters. Even the best models, according to the RMSD-based rankings, are not free of such errors. One could argue that one can generate easily a very precise model that it is inaccurate and that precision in geometric and stereochemical parameters are of lesser importance. These geometric and stereochemical parameters are very well established and need to be implemented to be helpful in the future for modelling structures with catalytic or fine recognition properties. Thus, a similarity/distance measure assessing a model against a reference structure cannot be the only reliable indicator of the model quality and that all the predictors should ensure the stereochemical accuracy of their models before submission. We suggest that a detailed stereochemical analysis should enter regular evaluation processes for improving the accuracy of RNA-Puzzles submissions and promoting high-quality RNA 3D structure prediction.

## MATERIALS AND METHODS

In this research, we used MAXIT version 10 downloaded from RCSB PDB (https://sw-tools.rcsb.org/apps/MAXIT), Barnaba 0.1.7 obtained from https://github.com/srnas/barnaba (Bottaro et al. 2019), MolProbity 4.5.1 taken from https://github.com/rlabduke/MolProbity (Williams et al. 2018), and X3DNA-DSSR version 2.4 (Lu and Olson 2003). Structures were divided into 24 subsets, one subset with the reference structures and 23 subsets with predicted models (one for each participant), and the average values and standard deviations were computed for them. Three clusters, H10, H14, and H16, including predictions by human groups, were excluded from the statistical analysis since, in all the challenges, these groups submitted only one model each. However, their MAXIT reports are also available in the Supplementary Materials.

MAXIT reports the following stereochemical issues: (i) close contacts, (ii) bond length deviations, (iii) bond angle deviations, (iv) deviations from planarity, (v) chirality errors, and (vi) polymer linkage errors (the P-O bond lengths). For (i)-(iii) and (vi), the program identifies abnormality if the parameter exceeds the expected value six times the standard sigma value. The expected values and the attached sigmas are based on (Clowney et al. 1996; Gelbin et al. 1996; Parkinson et al. 1996; Schneider et al. 1996) and have been attached in the Supplementary Materials. The current source of these reference terms is the Cambridge Structural Database (Bruno and Groom 2014; Tickle 2012; Urzhumtseva et al. 2009). Atomic clashes are signalled whenever any intermolecular atom pair is closer than the sum of their respective van der Waals radii. In general, a clash is defined when the distance between two atoms is less than 2.2Å (if no H atom is involved) or 1.6Å (if one H atom is involved) *(c.f.* Table D1 in the Supplementary Materials).

Departures from the average best-fit plane centre yield the RMS deviations for all atoms from the plane. It is reported when higher than 6×0.02Å or when at least one atom has a deviation higher than 0.02Å. MAXIT also determines improper torsion angles in the furanose ring and reports deviations from puckering in the ring (imposed by the sp^3^ carbon atoms).

In the case of chirality assessment, MAXIT lists the residues that contain unexpected configuration of chiral centres (C1’, C2’, C3’, C4’). Improper dihedrals (Gelbin et al. 1996) are a measure of the chirality/planarity of the structure at a specific atom.Polymer linkage between the adjacent nucleotides is measured based on the distances computed for O3’-P and O5’-P atom pairs. By default, the O3’-P distance is evaluated. However, if it exceeds 2.5Å, MAXIT takes the minimum value out of these two for the consideration.

Figures 4–6 in the manuscript were prepared using Symmetry Tool Plug-in 1.3 implemented in VMD software version 1.94 (Humphrey et al. 1996).

## ACKNOWLEDGEMENTS

This work was supported by the Polish National Science Centre (grants 2016/23/B/ST6/03931 and 2019/35/B/ST6/03074 to MS), grant POIR.04.02.00-30-A004/16, as well as the statutory funds of Poznan University of Technology (SBAD), Poland, and the Institute of Bioorganic Chemistry, Polish Academy of Sciences.

## AUTHOR CONTRIBUTIONS

FC, MA and MS conceived the project. MS supervised the analysis made by FC and MA. ZM and EW analysed and assessed the results. All authors contributed to the manuscript preparation.

## REFERENCES

Aftab SO, Ghouri MZ, Masood MU, Haider Z, Khan Z, Ahmad A, Munawar N. 2020. Analysis of SARS-CoV-2 RNA-dependent RNA polymerase as a potential therapeutic drug target using a computational approach. Journal of Translational Medicine 18:1–15. doi:10.1186/s12967-020-02439-0

Antczak M, Popenda M, Zok T, Sarzynska J, Ratajczak T, Tomczyk K, Adamiak RW, Szachniuk M. 2016. New functionality of RNAComposer: an application to shape the axis of miR160 precursor structure. Acta Biochim Pol 63:737–744. doi:10.18388/abp.2016_1329

Antczak M, Zok T, Osowiecki M, Popenda M, Adamiak RW, Szachniuk M. 2018. RNAfitme: a webserver for modeling nucleobase and nucleoside residue conformation in fixed-backbone RNA structures. BMC Bioinformatics 19:304. doi:10.1186/s12859-018-2317-9

Berman HM, Westbrook J, Feng Z, Gilliland G, Bhat TN, Weissig H, Shindyalov IN, Bourne PE. 2000. The Protein Data Bank. Nucleic Acids Res 28:235–242. doi:10.1093/nar/28.1.235

Berman H, Henrick K, Nakamura H. 2003. Announcing the worldwide protein data bank. Nat Struct Biol 10:980. doi: 10.1038/nsb1203-980

Bruno IJ, Groom CR. 2014. A crystallographic perspective on sharing data and knowledge. J Comput Aided Mol Des 28:1015–1022. doi:10.1007/s10822-014-9780-9

Boniecki MJ, Lach G, Dawson WK, Tomala K, Lukasz P, Soltysinski T, Rother KM, Bujnicki JM. 2016. SimRNA: a coarse-grained method for RNA folding simulations and 3D structure prediction. Nucleic Acids Res 44:e63. doi:10.1093/nar/gkv1479

Bottaro S, Bussi G, Pinamonti G, Reißer S, Boomsma W, Lindorff-Larsen K. 2019. Barnaba: software for analysis of nucleic acid structures and trajectories. RNA 25:219–231. doi:10.1261/rna.067678.118

Clowney L, Jain SC, Srinivasan AR, Westbrook J, Olson WK, Berman HM. 1996. Geometric parameters in nucleic acids: nitrogenous bases. J Am Chem Soc 118:509–518. doi:10.1021/ja952883d

Cruz JA, Blanchet MF, Boniecki M, Bujnicki JM, Chen SJ, Cao S, Das R, Ding F, Dokholyan NV, Flores SC, et al. 2012. RNA-Puzzles: A CASP-like evaluation of RNA three-dimensional structure prediction. RNA 18:610–625. doi:10.1261/rna.031054.111

Das R, Baker D. 2007. Automated de novo prediction of native-like RNA tertiary structures. PNAS 104:14664–14669. doi:10.1073/pnas.0703836104

Das R, Karanicolas J, Baker D. 2010. Atomic accuracy in predicting and designing noncanonical RNA structure. Nature Methods 7:291–294. doi:10.1038/nmeth.1433

Ding F, Sharma S, Chalasani P, Demidov VV, Broude NE, Dokholyan NV. 2008. Ab initio RNA folding by discrete molecular dynamics: From structure prediction to folding mechanisms. RNA 14:1164–1173. doi:10.1261/rna.894608

Feng Z, Hsieh S, Gelbin A, Westbrook J. 1998. MAXIT: Macromolecular Exchange and Input Tool. Rutgers University, New Brunswick.

Gelbin A, Schneider B, Clowney L, Hsieh S-H, Olson WK, Berman HM. 1996. Geometric Parameters in Nucleic Acids: Sugar and Phosphate Constituents. J Am Chem Soc 118:519–529. doi:10.1021/ja9528846

Gore S, Sanz García E, Hendrickx PMS, Gutmanas A, Westbrook JD, Yang H, Feng Z, Baskaran K, Berrisford JM, Hudson BP, et al. 2017. Validation of Structures in the Protein Data Bank. Structure 25:1916–1927. doi:10.1016/j.str.2017.10.009

Gumna J, Zok T, Figurski K, Pachulska-Wieczorek K, Szachniuk M. 2020. RNAthor - fast, accurate normalization, visualization and statistical analysis of RNA probing data resolved by capillary electrophoresis. PLOS ONE 15:e0239287. doi:10.1371/journal.pone.0239287

Güntert P, Buchner L. 2015. Combined automated NOE assignment and structure calculation with CYANA. J Biomol NMR 62:453–471. doi:10.1007/s10858-015-9924-9

Humphrey W, Dalke A, Schulten K. 1996. VMD: Visual molecular dynamics. J Mol Graph 14:33–38. doi:10.1016/0263-7855(96)00018-5

Jonikas MA, Radmer RJ, Laederach A, Das R, Pearlman S, Herschlag D, Altman RB. 2009. Coarse-grained modeling of large RNA molecules with knowledge-based potentials and structural filters. RNA 15:189–199. doi:10.1261/rna.1270809

Jossinet F, Westhof E. 2005. Sequence to Structure (S2S): display, manipulate and interconnect RNA data from sequence to structure. Bioinformatics 21:3320–3332. doi:10.1093/bioinformatics/bti504

Kabsch W. 1978. A discussion of the solution for the best rotation to relate two sets of vectors. Acta Crystallogr A 34:827–828. doi:10.1107/S0567739478001680

Li B, Cao Y, Westhof E, Miao Z. 2020. Advances in RNA 3D structure modeling using experimental data. Frontiers in Genetics 11:1147. doi:10.3389/fgene.2020.574485

Lu XJ, Olson, WK. 2003. 3DNA: a software package for the analysis, rebuilding and visualization of three-dimensional nucleic acid structures. Nucleic Acids Res 31:5108–5121. doi:10.1093/nar/gkg680

Lukasiak P, Antczak M, Ratajczak T, Szachniuk M, Popenda M, Adamiak RW, Blazewicz J. 2015. RNAssess - a webserver for quality assessment of RNA 3D structures. Nucleic Acids Res 43:W502–W506. doi:10.1093/nar/gkv557

Magnus M, Antczak M, Zok T, Wiedemann J, Lukasiak P, Cao Y, Bujnicki JM, Westhof E, Szachniuk M, Miao Z. 2020. RNA-Puzzles toolkit: A computational resource of RNA 3D structure benchmark datasets, structure manipulation, and evaluation tools. Nucleic Acids Res 48:576–588. doi:10.1093/nar/gkz1108

Major F, Turcotte M, Gautheret D, Lapalme G, Fillion E, Cedergren R. 1991. The combination of symbolic and numerical computation for three-dimensional modeling of RNA. Science 253:1255–1260. doi:10.1126/science.1716375

Massire C, Westhof E. 1998. MANIP: An interactive tool for modelling RNA. J Mol Graph Model 16:197–205. doi:10.1016/s1093-3263(98)80004-1

Miao Z, Westhof E. 2017. RNA Structure: Advances and Assessment of 3D Structure Prediction. Annu Rev Biophys 46:483–503. doi:10.1146/annurev-biophys-070816-034125

Miao Z, Adamiak RW, Antczak M, Batey RT, Becka AJ, Biesiada M, Boniecki MJ, Bujnicki, JM, Chen SJ, Cheng CY, et al. 2017. RNA-Puzzles Round III: 3D RNA structure prediction of five riboswitches and one ribozyme. RNA 23:655–672. doi:10.1261/rna.060368.116

Miao Z, Adamiak RW, Antczak M, Boniecki MJ, Bujnicki JM, Chen S-J, Cheng CY, Cheng Y, Chou F-C, Das R, et al. 2020. RNA-Puzzles Round IV: 3D structure predictions of four ribozymes and two aptamers. RNA 26:982–995. doi:10.1261/rna.075341.120

Miao Z, Adamiak RW, Blanchet M-F, Boniecki M, Bujnicki JM, Chen S-J, Cheng C, Chojnowski G, Chou F-C, Cordero P, et al. 2015. RNA-Puzzles Round II: assessment of RNA structure prediction programs applied to three large RNA structures. RNA 21:1066–1084. doi:10.1261/rna.049502.114

Michel F, Westhof E. 1990. Modelling of the three-dimensional architecture of group I catalytic introns based on comparative sequence analysis. J Mol Biol 216:585–610. doi:10.1016/0022-2836(90)90386-z

Parisien M, Major F. 2008. The MC-Fold and MC-Sym pipeline infers RNA structure from sequence data. Nature 452:51–55. doi:10.1038/nature06684

Parisien M, Cruz JA, Westhof E, Major F. 2009. New metrics for comparing and assessing discrepancies between RNA 3D structures and models. RNA 15:1875–1885. doi:10.1261/rna.1700409

Parkinson G, Vojtechovsky J, Clowney L, Brünger AT, Berman HM. 1996. New parameters for the refinement of nucleic acid-containing structures. Acta Crystallographica Section D: Biological Crystallography 52:57–64. doi:10.1107/S0907444995011115

Phillips JC, Hardy DJ, Maia JDC, Stone JE, Ribeiro JV, Bernardi JC, Buch R, Fiorin G, Henin J, Jiang W, et al. 2020. Scalable molecular dynamics on CPU and GPU architectures with NAMD. J Chem Phys 153:044130. doi:10.1063/5.0014475

Popenda M, Szachniuk M, Antczak M, Purzycka KJ, Lukasiak P, Bartol N, Blazewicz J, Adamiak RW. 2012. Automated 3D structure composition for large RNAs. Nucleic Acids Res 40:e112. doi:10.1093/nar/gks339

Popenda M, Zok T, Sarzynska J, Korpeta A, Adamiak RW, Antczak M, Szachniuk M. 2021. Entanglements of structure elements revealed in RNA 3D models. Nucleic Acids Res 49:9625–9632. doi:10.1093/nar/gkab716

Purzycka KJ, Popenda M, Szachniuk M, Antczak M, Lukasiak P, Blazewicz J, Adamiak RW. 2014. Automated 3D RNA structure prediction using the RNAComposer method for riboswitches. Meth Enzymol 553:3–34. doi: 10.1016/bs.mie.2014.10.050

Rother M, Rother K, Puton T, Bujnicki JM. 2011. ModeRNA: a tool for comparative modeling of RNA 3D structure. Nucleic Acids Res 39:4007–4022. doi:10.1093/nar/gkq1320

Schneider B, Kabelac M, Hobza P. 1996. Geometry of the Phosphate Group and Its Interactions with Metal Cations in Crystals and ab Initio Calculations. J Am Chem Soc 118:12207–12217.

Schwieters CD, Kuszewski JJ, Tjandra N, Clore GM. 2003. The Xplor-NIH NMR Molecular Structure Determination Package. J Magn Res 160:66–74. doi:10.1016/s1090-7807(02)00014-9

Stasiewicz J, Mukherjee S, Nithin C, Bujnicki JM. 2019. QRNAS: software tool for refinement of nucleic acid structures. BMC Struct Biol 19:5. doi: 10.1186/s12900-019-0103-1

Teplova M, Falschlunger C, Krasheninina O, Egger M, Ren A, Patel DJ, Micura R. 2020. Crucial roles of two hydrated Mg2+ ions in reaction catalysis of the pistol ribozyme. Angewandte Chemie, 132:2859–2865. doi: 10.1002/ange.201912522

Tickle IJ. 2012. Statistical quality indicators for electron-density maps. Acta Crystallogr D 68:454–467. doi:10.1107/S0907444911035918

Ulrich EL, Akutsu H, Doreleijers JF, Harano Y, Ioannidis YE, Lin J, Livny M, Mading S, Maziuk D, Miller Z, et al. 2008. BioMagResBank. Nucleic Acids Res 36:D402–D408. doi:10.1093/nar/gkm957

Urzhumtseva L, Afonine PV, Adams PD, Urzhumtsev A. 2009. Crystallographic model quality at a glance. Acta Crystallogr D 65:297–300. doi:10.1107/S0907444908044296

Westhof E, Romby P, Romaniuk PJ, Ebel JP, Ehresmann C, Ehresmann B. 1989. Computer modeling from solution data of spinach chloroplast and of Xenopus laevis somatic and oocyte 5 S rRNAs. J Mol Biol 207:417–431. doi:10.1016/0022-2836(89)90264-7

White NA, Sumita M, Marquez VE, Hoogstraten CG. 2018. Coupling between conformational dynamics and catalytic function at the active site of the lead-dependent ribozyme. RNA 24:1542–1554. doi:10.1261/rna.067579.118

Wiedemann J, Zok T, Milostan M, Szachniuk M. 2017. LCS-TA to identify similar fragments in RNA 3D structures, BMC Bioinformatics 18:456. doi:10.1186/s12859-017-1867-6

Williams CJ, Headd JJ, Moriarty NW, Prisant MG, Videau LL, Deis LN, Verma V, Keedy DA, Hintze BJ, Chen VB, et al. 2018. MolProbity: More and better reference data for improved all-atom structure validation. Protein Sci 27:293–315. doi:10.1002/pro.3330

Xu X, Zhao P, Chen S-J. 2014. Vfold: A Web Server for RNA Structure and Folding Thermodynamics Prediction. PLOS ONE 9:e107504. doi:10.1371/journal.pone.0107504

Young JY, Westbrook JD, Feng Z, Sala R, Peisach E, Oldfield TJ, Sen S, Gutmanas A, Armstrong DR, Berrisford JM, et al. 2017. OneDep: unified wwPDB system for deposition, biocuration, and validation of macromolecular structures in the PDB archive. Structure 25:536–545. doi:10.1016/j.str.2017.01.004

Zhao Y, Huang Y, Gong Z, Wang Y, Man J, Xiao Y. 2012. Automated and fast building of three-dimensional RNA structures. Sci Rep 2:734. doi:10.1038/srep00734

Zok T, Popenda M, Szachniuk M. 2014. MCQ4Structures to compute similarity of molecule structures. Cent Eur J Oper Res 22:457–473. doi:10.1007/s10100-013-0296-5

Zok T, Antczak M, Riedel M, Nebel D, Villmann T, Lukasiak P, Blazewicz J, Szachniuk M. 2015. Building the library of RNA 3D nucleotide conformations using clustering approach. Int J Appl Math Comput Sci 25:689–700. doi:10.1515/amcs-2015-0050

