## Supplementary Material for "Evaluation of the stereochemical quality of predicted RNA 3D models in the RNA-Puzzles submissions"

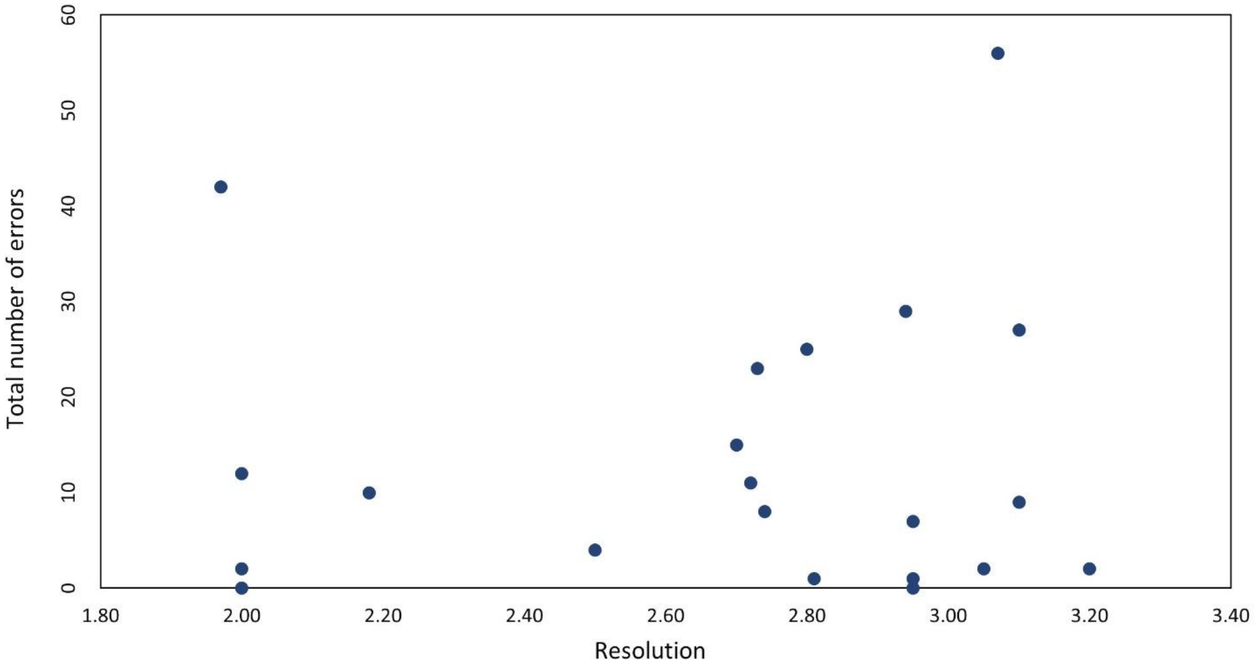

**Figure S1**. Resolution of reference structures versus number of errors reported by MAXIT.

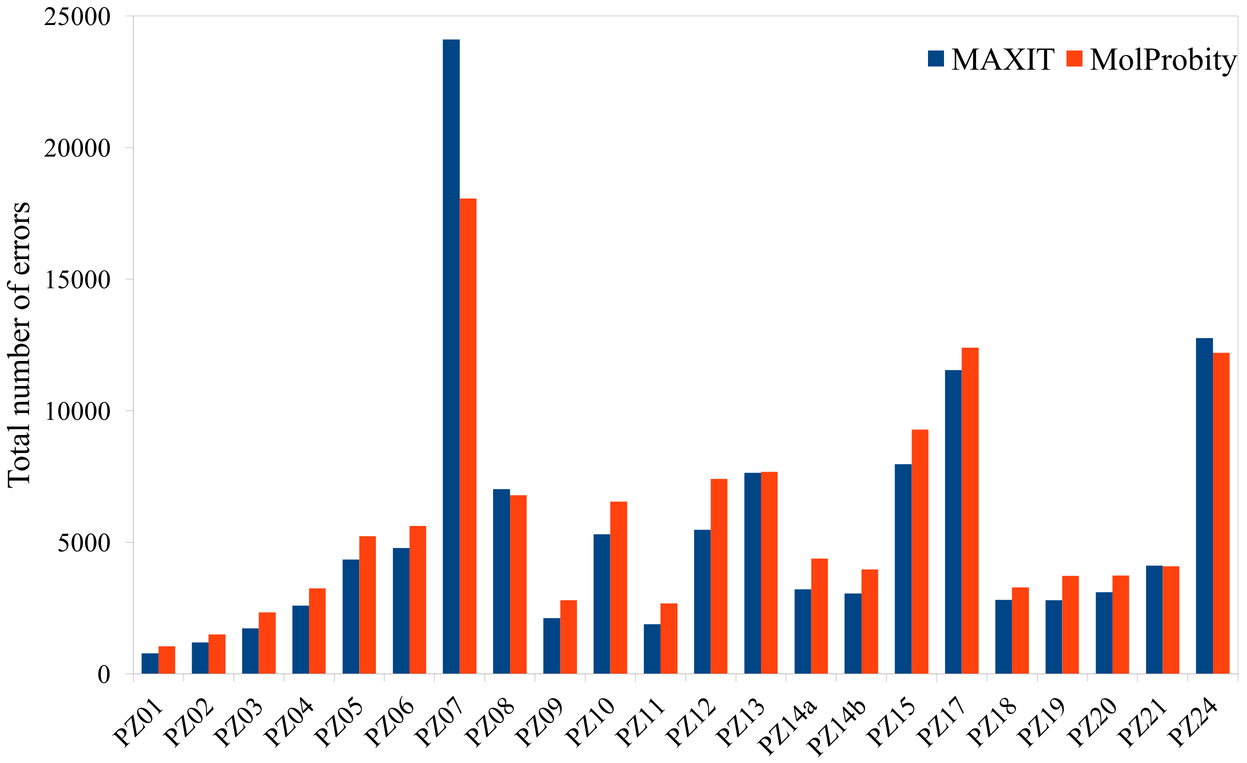

**Figure S2**. The total number of errors identified by MAXIT and MolProbity in all predicted models, divided by RNA-Puzzles challenges.

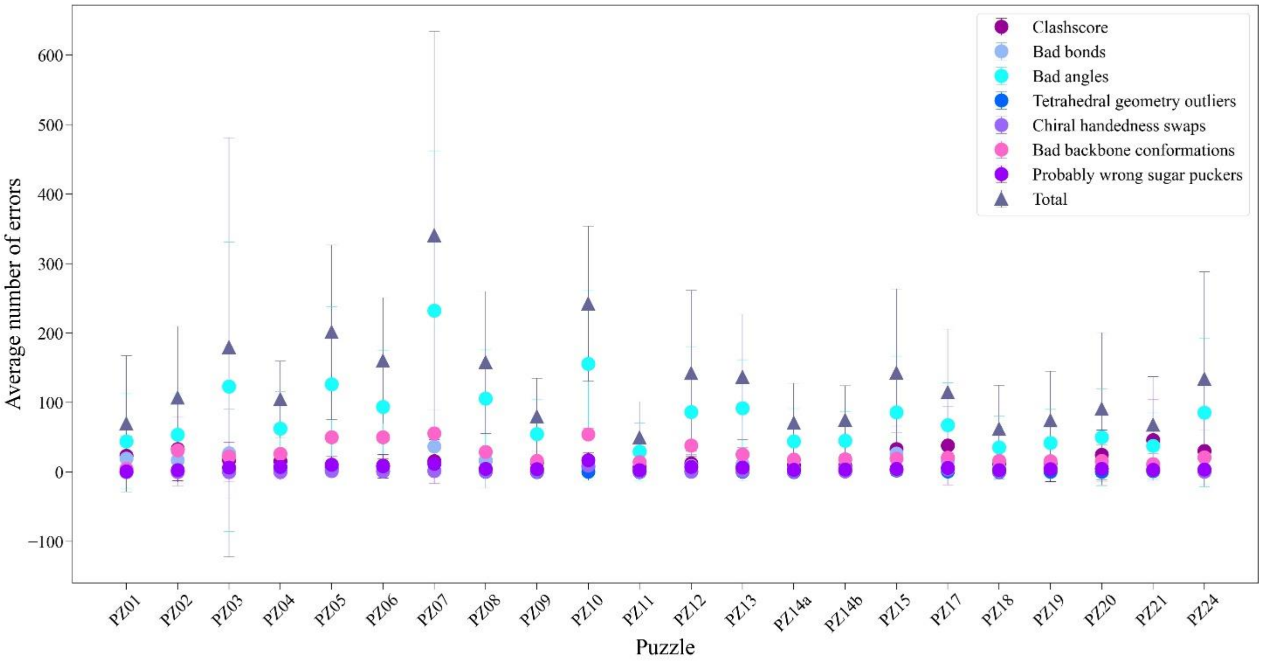

**Figure S3**. The average number of errors identified for each puzzle by MolProbity software.

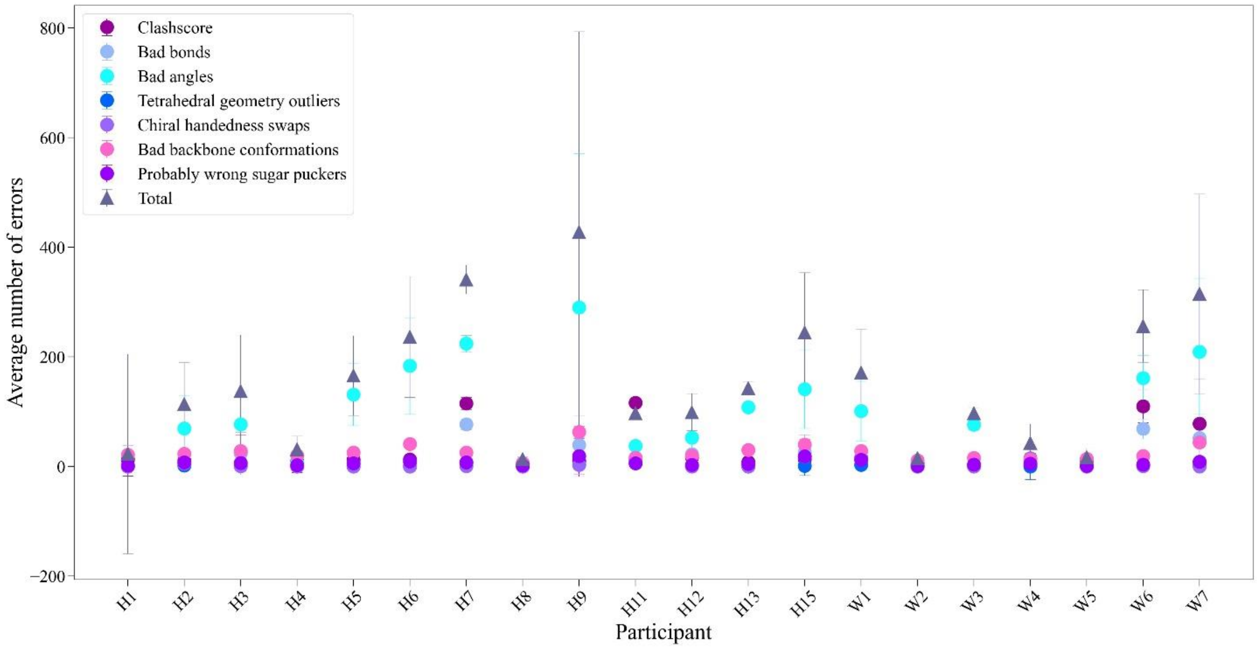

**Figure S4**. The average number of errors identified for each participant by MolProbity software.

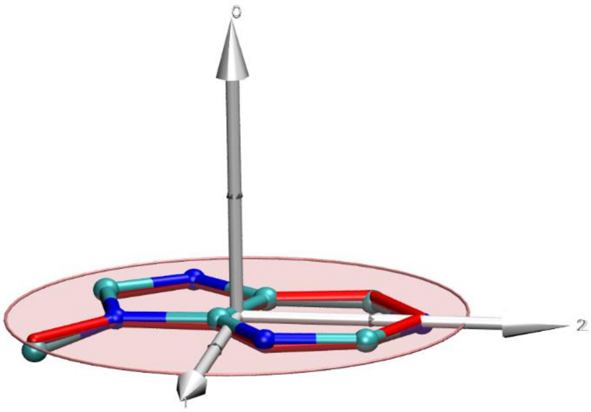

**Figure S5**. Planarity deviation for residue A28 in the PZ04 reference structure. Adenine from the target structure is depicted in ball-and-sticks representation with carbons in green and nitrogen in blue. Idealized structure is highlighted in red sticks. The reference plane appears in a red-transparent circle in its best fit from all atoms but hydrogen. Principle axes of inertia are shown with white arrows.The plane deviation presented in the figure equals 0.074Å. Although this value is not high, MAXIT indicates it as an error. MAXIT reports an error in this category whenever the deviations computed over all the atoms for the plane is higher than 6x0.02Å or at least one atom distance exceeds 0.02Å. In the presented example, the deviation of a single atom exceeds the threshold.

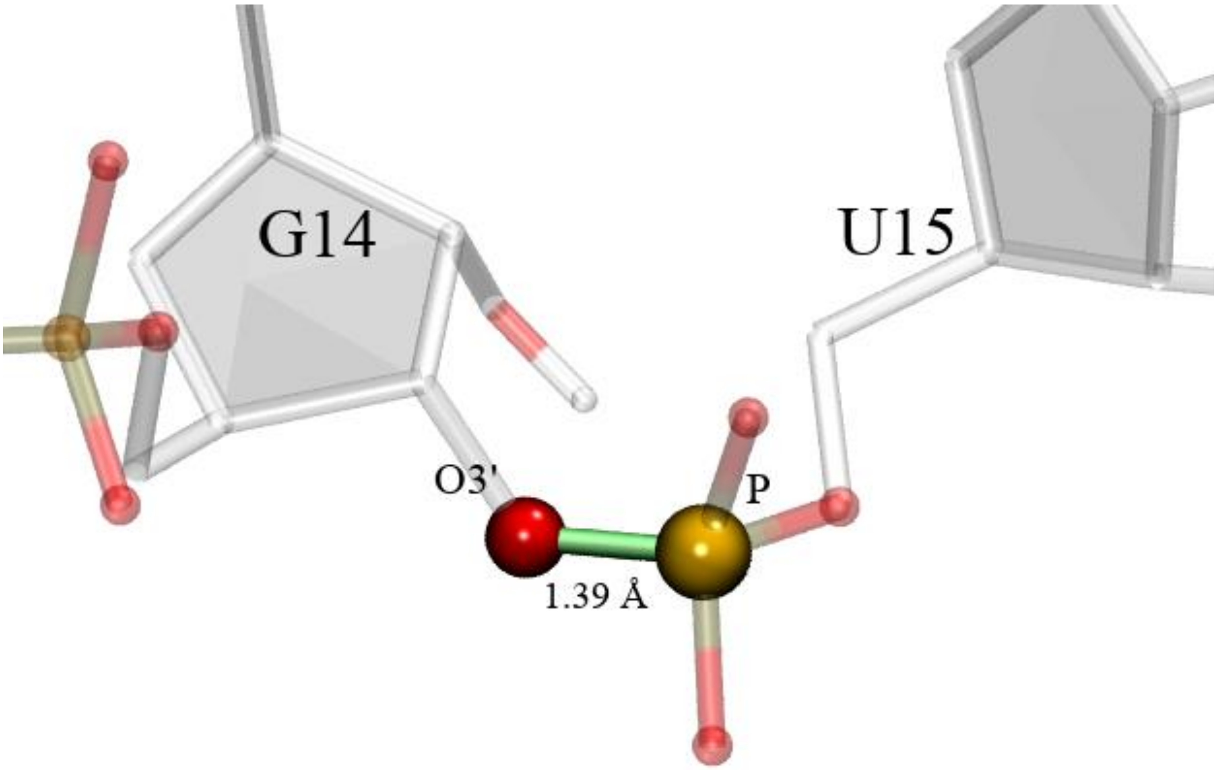

**Figure S6**. Polymer linkage abnormality found between Guanidine 14 and Uracil 15 (in the H7 model submitted to PZ24). The distance 1.39Å was measured by MAXIT (when it should be around 1.6Å with a sigma around 0.01).

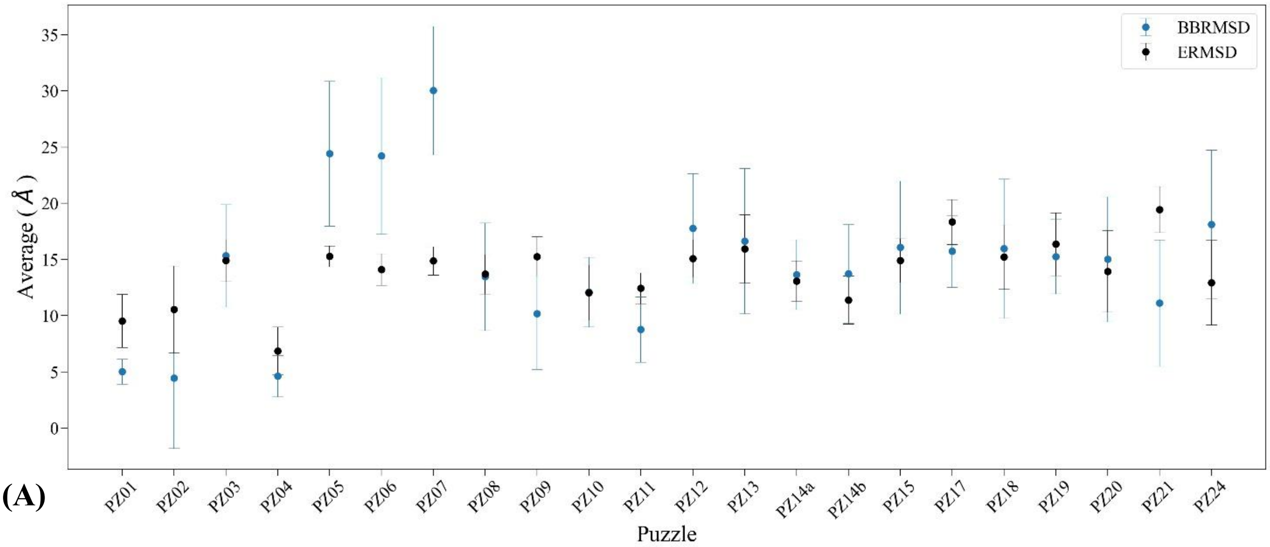

**(A)**

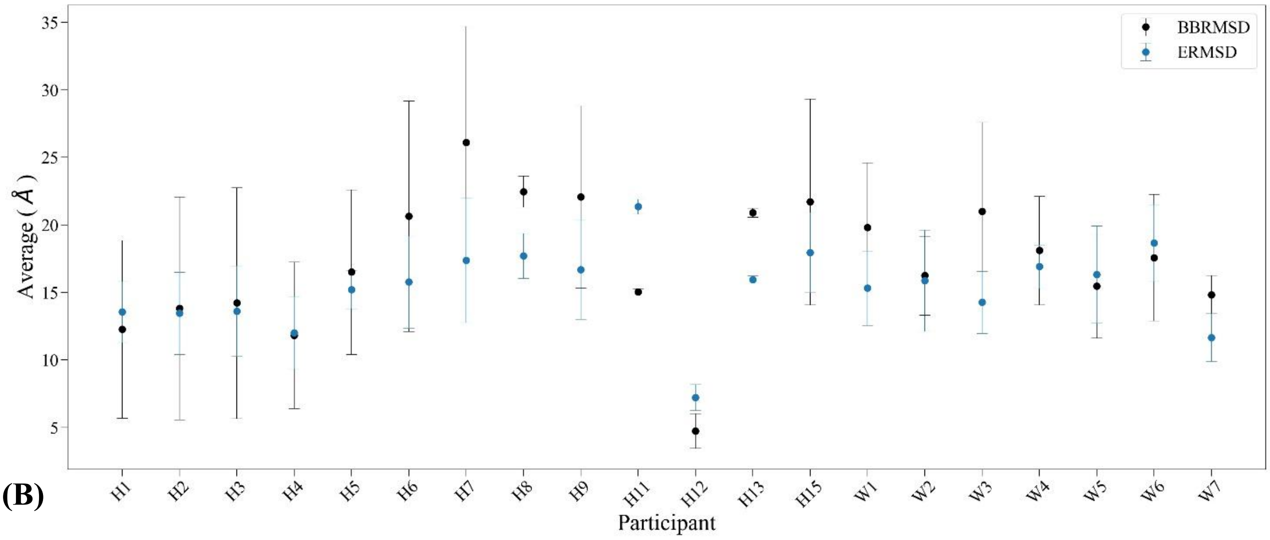

**(B)**

**Figure S7**. The average RMSD computed for the backbone (BBRMSD) and base pairs (eRMSD) over all models per puzzle (A), and per participant (B). Bars represent the standard deviation for every mean value.

RCSB MAXIT reports on the stereochemistry of experimentally determined 3D structures includes:

- the number of identified too-close contacts between symmetry-related molecules,
- bond lengths and bond angles,
- the number of planarity deviations,
- the number of chirality errors,
- the number of identified polymer linkage artefacts.

**Table S1**. MAXIT report on the reference structures.

| **No.** | **Puzzle id** | **PDB id** | **Reso-lution** | **Close con-tacts** | **Bond lengths** | **Bond angles** | **Plana-rity** | **Chira-lity** | **Poly-mer linkage** | **To-tal** |
|---|---|---|---|---|---|---|---|---|---|---|
| 1 | PZ01 | 3MEI | 1.97 | 1 | 5 | 35 | 0 | 0 | 1 | 42 |
| 2 | PZ02 | 3P59 | 2.18 | 2 | 0 | 8 | 0 | 0 | 0 | 10 |
| 3 | PZ03 | 3OWZ | 2.95 | 0 | 0 | 1 | 0 | 0 | 0 | 1 |
| 4 | PZ04 | 3V7E | 2.80 | 21 | 0 | 4 | 0 | 0 | 0 | 25 |
| 5 | PZ05 | 4P9R | 2.70 | 15 | 0 | 0 | 0 | 0 | 0 | 15 |
| 6 | PZ06 | 4GXY | 3.05 | 0 | 0 | 0 | 0 | 0 | 2 | 2 |
| 7 | PZ07 | 4R4V | 3.07 | 0 | 3 | 53 | 0 | 0 | 0 | 56 |
| 8 | PZ08 | 4L81 | 2.95 | 6 | 0 | 0 | 0 | 0 | 1 | 7 |
| 9 | PZ09 | 5KPY | 2.00 | 0 | 0 | 2 | 0 | 0 | 0 | 2 |
| 10 | PZ10 | 4LCK | 3.20 | 0 | 0 | 1 | 0 | 0 | 1 | 2 |
| 11 | PZ11 | 5IEM | - | 0 | 0 | 1 | 0 | 0 | 0 | 1 |
| 12 | PZ12 | 4QLM | 2.72 | 0 | 0 | 10 | 0 | 0 | 1 | 11 |
| 13 | PZ13 | 4XW7 | 2.50 | 0 | 0 | 1 | 3 | 0 | 0 | 4 |
| 14 | PZ14a | 5DDO | 3.10 | 0 | 0 | 7 | 0 | 0 | 2 | 9 |
| 15 | PZ14b | 5DDO | 3.10 | 2 | 15 | 9 | 0 | 0 | 1 | 27 |
| 16 | PZ15 | 5DI4 | 2.95 | 0 | 0 | 0 | 0 | 0 | 0 | 0 |
| 17 | PZ17 | 5K7C | 2.73 | 0 | 7 | 16 | 0 | 0 | 0 | 23 |
| 18 | PZ18 | 5TPY | 2.81 | 0 | 0 | 1 | 0 | 0 | 0 | 1 |
| 19 | PZ19 | 5T5A | 2.00 | 0 | 0 | 6 | 6 | 0 | 0 | 12 |
| 20 | PZ20 | 5Y85 | 2.00 | 0 | 0 | 0 | 0 | 0 | 0 | 0 |
| 21 | PZ21 | 5NZ6 | 2.94 | 0 | 2 | 27 | 0 | 0 | 0 | 29 |
| 22 | PZ24 | 6OL3 | 2.74 | 7 | 0 | 1 | 0 | 0 | 0 | 8 |
| Average | | | | 2.45 | 1.45 | 8.32 | 0.41 | 0.00 | 0.41 | 13.05 |
| Standard deviation | | | | 5.47 | 3.56 | 13.54 | 1.40 | 0.00 | 0.67 | 14.95 |
| Median | | | | 0 | 0 | 1.5 | 0 | 0 | 0 | 8.5 |

**Table S2**. MAXIT report on structures in Puzzle 01.

| No. | 3D RNA model | Close con-tacts | Bond lengths | Bond angles | Plana-rity | Chira-lity | Poly-mer lin-kage | Total |
|---|---|---|---|---|---|---|---|---|
| 1 | PZ1_Das_1 | 0 | 0 | 0 | 0 | 0 | 0 | 0 |
| 2 | PZ1_Das_3 | 0 | 0 | 0 | 0 | 0 | 0 | 0 |
| 3 | PZ01_Das_4 | 0 | 0 | 0 | 0 | 0 | 0 | 0 |
| 4 | PZ01_Das_2 | 0 | 0 | 1 | 0 | 0 | 0 | 1 |
| 5 | PZ01_Das_5 | 0 | 0 | 3 | 0 | 0 | 0 | 3 |
| 6 | PZ01_Bujnicki_1 | 0 | 0 | 15 | 9 | 1 | 0 | 25 |
| 7 | PZ01_refstructure | 0 | 2 | 27 | 0 | 0 | 0 | 29 |
| 8 | PZ01_Dokholyan_1 | 0 | 0 | 27 | 12 | 0 | 1 | 40 |
| 9 | PZ01_Chen_1 | 0 | 1 | 30 | 9 | 0 | 0 | 40 |
| 10 | PZ01_Santalucia_1 | 0 | 44 | 29 | 0 | 0 | 2 | 75 |
| 11 | PZ01_Bujnicki_3 | 3 | 23 | 33 | 1 | 0 | 19 | 79 |
| 12 | PZ01_Bujnicki_4 | 17 | 25 | 40 | 1 | 0 | 7 | 90 |
| 13 | PZ01_Bujnicki_5 | 19 | 20 | 46 | 1 | 0 | 14 | 100 |
| 14 | PZ01_Bujnicki_2 | 4 | 30 | 55 | 1 | 0 | 20 | 110 |
| 15 | PZ01_Major_1 | 1 | 85 | 209 | 29 | 0 | 0 | 324 |
| Average | | 2.93 | 15.33 | 34.33 | 4.20 | 0.07 | 4.20 | 61.07 |
| Standard deviation | | 6.25 | 24.04 | 51.59 | 7.96 | 0.26 | 7.30 | 82.71 |
| Median | | 0 | 1 | 27 | 1 | 0 | 0 | 40 |

**Table S3**. MAXIT report on structures in Puzzle 02.

| No. | 3D RNA model | Close con-tacts | Bond lengths | Bond angles | Plana-rity | Chira-lity | Poly-mer lin-kage | Total |
|---|---|---|---|---|---|---|---|---|
| 1 | PZ02_Das_3 | 0 | 0 | 9 | 0 | 0 | 0 | 9 |
| 2 | PZ02_Das_1 | 1 | 0 | 9 | 0 | 0 | 0 | 10 |
| 3 | PZ02_Das_2 | 1 | 0 | 9 | 0 | 0 | 0 | 10 |
| 4 | PZ02_Das_4 | 1 | 1 | 9 | 0 | 0 | 0 | 11 |
| 5 | PZ02_Das_5 | 2 | 1 | 9 | 0 | 0 | 0 | 12 |
| 6 | PZ02_Major_1 | 0 | 12 | 18 | 0 | 0 | 0 | 30 |
| 7 | PZ02_refstructure | 1 | 5 | 35 | 0 | 0 | 1 | 42 |
| 8 | PZ02_Bujnicki_3 | 0 | 0 | 35 | 29 | 0 | 0 | 64 |
| 9 | PZ02_Bujnicki_2 | 0 | 3 | 36 | 30 | 0 | 0 | 69 |
| 10 | PZ02_Dokholyan_1 | 0 | 0 | 52 | 22 | 0 | 0 | 74 |
| 11 | PZ02_Santalucia_1 | 6 | 32 | 41 | 0 | 0 | 2 | 81 |
| 12 | PZ02_Bujnicki_1 | 0 | 4 | 70 | 20 | 1 | 0 | 95 |
| 13 | PZ02_Chen_1 | 5 | 21 | 196 | 30 | 2 | 0 | 254 |
| 14 | PZ02_Wildauer_1 | 271 | 39 | 210 | 0 | 0 | 0 | 520 |
| Average | | 20.57 | 8.43 | 52.71 | 9.36 | 0.21 | 0.21 | 91.50 |
| Standard deviation | | 72.10 | 12.97 | 66.43 | 13.30 | 0.58 | 0.58 | 138.91 |
| Median | | 1 | 2 | 35 | 0 | 0 | 0 | 53 |

**Table S4**. MAXIT report on structures in Puzzle 03.

| No. | 3D RNA model | Close con-tacts | Bond lengths | Bond angles | Plana-rity | Chira-lity | Poly-mer lin-kage | Total |
|---|---|---|---|---|---|---|---|---|
| 1 | PZ03_Das_4 | 0 | 0 | 0 | 0 | 0 | 1 | 1 |
| 2 | PZ03_refstructure | 0 | 0 | 1 | 0 | 0 | 0 | 1 |
| 3 | PZ03_Das_1 | 0 | 0 | 0 | 0 | 0 | 1 | 1 |
| 4 | PZ03_Das_2 | 0 | 0 | 0 | 0 | 0 | 1 | 1 |
| 5 | PZ03_Das_3 | 0 | 0 | 2 | 0 | 0 | 0 | 2 |
| 6 | PZ03_Das_5 | 0 | 1 | 0 | 0 | 0 | 1 | 2 |
| 7 | PZ03_Bujnicki_2 | 0 | 0 | 35 | 29 | 0 | 0 | 64 |
| 8 | PZ03_Bujnicki_1 | 0 | 0 | 44 | 37 | 0 | 0 | 81 |
| 9 | PZ03_Dokholyan_2 | 0 | 0 | 59 | 30 | 0 | 0 | 89 |
| 10 | PZ03_Dokholyan_1 | 0 | 0 | 68 | 29 | 0 | 0 | 97 |
| 11 | PZ03_Chen_1 | 0 | 1 | 114 | 25 | 0 | 0 | 140 |
| 12 | PZ03_Major_2 | 0 | 145 | 398 | 48 | 1 | 0 | 592 |
| 13 | PZ03_Major_1 | 0 | 143 | 415 | 36 | 1 | 0 | 595 |
| Average | | 0.00 | 22.31 | 87.38 | 18.00 | 0.15 | 0.31 | 128.15 |
| Standard deviation | | 0.00 | 54.01 | 145.95 | 18.18 | 0.38 | 0.48 | 211.85 |
| Median | | 0 | 0 | 35 | 25 | 0 | 0 | 64 |

**Table S5**. MAXIT report on structures in Puzzle 04.

| No. | 3D RNA model | Close con-tacts | Bond lengths | Bond angles | Plana-rity | Chira-lity | Poly-mer lin-kage | Total |
|---|---|---|---|---|---|---|---|---|
| 1 | PZ04_Adamiak_1 | 0 | 0 | 0 | 0 | 0 | 0 | 0 |
| 2 | PZ04_Adamiak_2 | 0 | 0 | 0 | 0 | 0 | 0 | 0 |
| 3 | PZ04_Adamiak_3 | 0 | 0 | 0 | 0 | 0 | 0 | 0 |
| 4 | PZ04_Adamiak_4 | 0 | 0 | 0 | 0 | 0 | 0 | 0 |
| 5 | PZ04_Adamiak_5 | 0 | 0 | 0 | 0 | 0 | 0 | 0 |
| 6 | PZ04_Das_2 | 0 | 0 | 1 | 0 | 0 | 0 | 1 |
| 7 | PZ04_Das_3 | 0 | 0 | 1 | 0 | 0 | 0 | 1 |
| 8 | PZ04_Das_4 | 0 | 0 | 1 | 0 | 0 | 0 | 1 |
| 9 | PZ04_Das_5 | 0 | 0 | 1 | 0 | 0 | 0 | 1 |
| 10 | PZ04_Das_1 | 0 | 0 | 1 | 0 | 0 | 0 | 1 |
| 11 | PZ04_refstructure | 0 | 0 | 6 | 6 | 0 | 0 | 12 |
| 12 | PZ04_Bujnicki_4 | 1 | 18 | 27 | 2 | 0 | 2 | 50 |
| 13 | PZ04_Bujnicki_2 | 1 | 19 | 27 | 2 | 0 | 2 | 51 |
| 14 | PZ04_Dokholyan_1 | 0 | 0 | 37 | 16 | 2 | 0 | 55 |
| 15 | PZ04_Dokholyan_2 | 0 | 0 | 37 | 16 | 2 | 0 | 55 |
| 16 | PZ04_Bujnicki_5 | 1 | 19 | 30 | 6 | 0 | 2 | 58 |
| 17 | PZ04_Bujnicki_3 | 1 | 18 | 31 | 8 | 0 | 2 | 60 |
| 18 | PZ04_Santalucia_1 | 0 | 26 | 29 | 0 | 2 | 5 | 62 |
| 19 | PZ04_Bujnicki_1 | 1 | 20 | 34 | 7 | 0 | 2 | 64 |
| 20 | PZ04_Mikolajczak_1 | 26 | 15 | 24 | 6 | 0 | 1 | 72 |
| 21 | PZ04_Chen_1 | 0 | 1 | 122 | 16 | 0 | 0 | 139 |

| | | | | | | | | |
|---|---|---|---|---|---|---|---|---|
| 22 | PZ04_Chen_4 | 0 | 3 | 135 | 22 | 0 | 0 | 160 |
| 23 | PZ04_Chen_2 | 0 | 2 | 149 | 19 | 0 | 0 | 170 |
| 24 | PZ04_Major_1 | 0 | 0 | 150 | 32 | 1 | 0 | 183 |
| 25 | PZ04_Chen_8 | 0 | 5 | 160 | 21 | 0 | 0 | 186 |
| 26 | PZ04_Chen_6 | 0 | 2 | 163 | 24 | 0 | 0 | 189 |
| 27 | PZ04_Chen_10 | 0 | 4 | 171 | 19 | 0 | 0 | 194 |
| 28 | PZ04_Chen_7 | 0 | 3 | 169 | 23 | 0 | 0 | 195 |
| 29 | PZ04_Chen_3 | 0 | 7 | 173 | 20 | 0 | 0 | 200 |
| 30 | PZ04_Chen_5 | 0 | 14 | 177 | 18 | 0 | 0 | 209 |
| 31 | PZ04_Chen_9 | 0 | 18 | 186 | 20 | 0 | 0 | 224 |
| Average | | 1.00 | 6.26 | 65.87 | 9.77 | 0.23 | 0.52 | 83.65 |
| Standard deviation | | 4.65 | 8.35 | 72.53 | 9.89 | 0.62 | 1.12 | 81.98 |
| Median | | 0 | 2 | 30 | 6 | 0 | 0 | 58 |

**Table S6**. MAXIT report on structures in Puzzle 05.

| **No.** | **3D RNA model** | **Close con-tacts** | **Bond lengths** | **Bond angles** | **Plana-rity** | **Chira-lity** | **Poly-mer lin-kage** | **Total** |
|---|---|---|---|---|---|---|---|---|
| 1 | PZ05_Adamiak_1 | 0 | 0 | 0 | 0 | 0 | 0 | 0 |
| 2 | PZ05_refstructure | 2 | 0 | 8 | 0 | 0 | 0 | 10 |
| 3 | PZ05_Bujnicki_4 | 0 | 0 | 22 | 0 | 0 | 0 | 22 |
| 4 | PZ05_Bujnicki_1 | 0 | 0 | 25 | 0 | 0 | 0 | 25 |
| 5 | PZ05_Bujnicki_5 | 0 | 0 | 29 | 0 | 0 | 0 | 29 |
| 6 | PZ05_Bujnicki_3 | 2 | 0 | 31 | 0 | 0 | 0 | 33 |
| 7 | PZ05_Bujnicki_2 | 1 | 0 | 34 | 0 | 0 | 0 | 35 |
| 8 | PZ05_Das_1 | 0 | 10 | 29 | 0 | 0 | 1 | 40 |
| 9 | PZ05_Das_2 | 0 | 7 | 34 | 0 | 0 | 2 | 43 |
| 10 | PZ05_Chen_7 | 0 | 0 | 65 | 3 | 5 | 0 | 73 |
| 11 | PZ05_Chen_6 | 0 | 0 | 67 | 5 | 5 | 0 | 77 |
| 12 | PZ05_Chen_1 | 1 | 10 | 94 | 9 | 5 | 0 | 119 |
| 13 | PZ05_Chen_4 | 0 | 17 | 93 | 14 | 5 | 0 | 129 |
| 14 | PZ05_Chen_2 | 0 | 20 | 97 | 16 | 5 | 0 | 138 |
| 15 | PZ05_Chen_3 | 0 | 22 | 115 | 10 | 4 | 0 | 151 |
| 16 | PZ05_Chen_5 | 1 | 26 | 108 | 7 | 6 | 3 | 151 |
| 17 | PZ05_Dokholyan_6 | 0 | 0 | 145 | 59 | 0 | 0 | 204 |
| 18 | PZ05_Dokholyan_7 | 0 | 0 | 149 | 59 | 0 | 0 | 208 |
| 19 | PZ05_Dokholyan_8 | 0 | 1 | 162 | 60 | 0 | 0 | 223 |
| 20 | PZ05_Dokholyan_3 | 0 | 4 | 178 | 125 | 0 | 0 | 307 |
| 21 | PZ05_Dokholyan_5 | 0 | 3 | 191 | 134 | 0 | 0 | 328 |
| 22 | PZ05_Dokholyan_4 | 0 | 5 | 217 | 135 | 0 | 0 | 357 |
| 23 | PZ05_Dokholyan_2 | 0 | 5 | 217 | 135 | 0 | 0 | 357 |
| 24 | PZ05_Dokholyan_1 | 0 | 3 | 222 | 135 | 0 | 0 | 360 |
| 25 | PZ05_Xiao_1 | 0 | 14 | 331 | 71 | 3 | 0 | 419 |
| 26 | PZ05_Xiao_2 | 0 | 45 | 368 | 71 | 5 | 4 | 493 |
| Average | | 0.27 | 7.38 | 116.58 | 40.31 | 1.65 | 0.38 | 166.58 |

| | | | | | | | |
|---|---|---|---|---|---|---|---|
| Standard deviation | 0.60 | 10.93 | 97.85 | 51.91 | 2.37 | 1.02 | 145.96 |
| Median | 0 | 3 | 96.5 | 9.5 | 0 | 0 | 133.5 |

**Table S7**. MAXIT report on structures in Puzzle 06.

| No. | 3D RNA model | Close con-tacts | Bond lengths | Bond angles | Plana-rity | Chira-lity | Poly-mer lin-kage | Total |
|---|---|---|---|---|---|---|---|---|
| 1 | PZ06_refstructure | 0 | 0 | 0 | 0 | 0 | 2 | 2 |
| 2 | PZ06_Bujnicki_3 | 1 | 0 | 13 | 0 | 0 | 0 | 14 |
| 3 | PZ06_Bujnicki_1 | 0 | 0 | 16 | 0 | 0 | 0 | 16 |
| 4 | PZ06_Bujnicki_4 | 3 | 0 | 16 | 0 | 0 | 0 | 19 |
| 5 | PZ06_Bujnicki_2 | 2 | 0 | 22 | 0 | 0 | 0 | 24 |
| 6 | PZ06_Das_7 | 5 | 8 | 12 | 1 | 0 | 3 | 29 |
| 7 | PZ06_Das_3 | 6 | 8 | 12 | 1 | 0 | 4 | 31 |
| 8 | PZ06_Das_9 | 6 | 11 | 18 | 0 | 0 | 3 | 38 |
| 9 | PZ06_Das_5 | 17 | 6 | 15 | 0 | 0 | 4 | 42 |
| 10 | PZ06_Das_8 | 17 | 6 | 16 | 0 | 0 | 3 | 42 |
| 11 | PZ06_Das_2 | 8 | 12 | 22 | 0 | 0 | 2 | 44 |
| 12 | PZ06_Das_10 | 21 | 13 | 17 | 1 | 0 | 4 | 56 |
| 13 | PZ06_Das_6 | 63 | 11 | 23 | 1 | 0 | 6 | 104 |
| 14 | PZ06_Das_1 | 63 | 12 | 23 | 1 | 0 | 5 | 104 |
| 15 | PZ06_Das_4 | 63 | 12 | 23 | 1 | 0 | 6 | 105 |
| 16 | PZ06_Chen_1 | 0 | 0 | 97 | 23 | 0 | 0 | 120 |
| 17 | PZ06_Chen_2 | 0 | 2 | 113 | 21 | 0 | 0 | 136 |
| 18 | PZ06_Chen_4 | 0 | 0 | 115 | 29 | 0 | 0 | 144 |
| 19 | PZ06_Chen_3 | 0 | 2 | 121 | 32 | 0 | 0 | 155 |
| 20 | PZ06_Chen_6 | 0 | 5 | 115 | 37 | 0 | 0 | 157 |
| 21 | PZ06_Dokholyan_1 | 0 | 0 | 113 | 50 | 0 | 0 | 163 |
| 22 | PZ06_Dokholyan_5 | 0 | 0 | 112 | 53 | 0 | 0 | 165 |
| 23 | PZ06_Dokholyan_4 | 0 | 0 | 122 | 46 | 0 | 0 | 168 |
| 24 | PZ06_Chen_7 | 2 | 20 | 124 | 28 | 0 | 1 | 175 |
| 25 | PZ06_Dokholyan_2 | 0 | 0 | 118 | 57 | 0 | 0 | 175 |
| 26 | PZ06_Dokholyan_3 | 0 | 0 | 122 | 55 | 0 | 0 | 177 |
| 27 | PZ06_Major_6 | 1 | 0 | 147 | 42 | 3 | 0 | 193 |
| 28 | PZ06_Major_4 | 0 | 0 | 142 | 56 | 4 | 0 | 202 |
| 29 | PZ06_Major_7 | 0 | 0 | 147 | 57 | 4 | 0 | 208 |
| 30 | PZ06_Major_1 | 0 | 0 | 143 | 63 | 4 | 0 | 210 |
| 31 | PZ06_Major_3 | 1 | 0 | 147 | 61 | 5 | 0 | 214 |
| 32 | PZ06_Major_2 | 0 | 0 | 160 | 54 | 7 | 0 | 221 |
| 33 | PZ06_Major_5 | 0 | 0 | 170 | 55 | 5 | 0 | 230 |
| 34 | PZ06_Chen_5 | 3 | 29 | 185 | 59 | 1 | 1 | 278 |
| 35 | PZ06_Dokholyan_6 | 0 | 2 | 199 | 124 | 0 | 0 | 325 |
| Average | | 8.06 | 4.54 | 84.57 | 28.80 | 0.94 | 1.26 | 128.17 |
| Standard deviation | | 17.86 | 6.86 | 63.18 | 29.85 | 1.92 | 1.93 | 84.66 |
| Median | | 0 | 0 | 113 | 28 | 0 | 0 | 144 |

**Table S8**. MAXIT report on structures in Puzzle 07.

| No. | 3D RNA model | Close con-tacts | Bond lengths | Bond angles | Plana-rity | Chira-lity | Poly-mer lin-kage | Total |
|---|---|---|---|---|---|---|---|---|
| 1 | PZ07_Adamiak_1 | 0 | 0 | 0 | 0 | 0 | 0 | 0 |
| 2 | PZ07_Adamiak_2 | 0 | 0 | 0 | 0 | 0 | 0 | 0 |
| 3 | PZ07_Adamiak_3 | 0 | 0 | 0 | 0 | 0 | 0 | 0 |
| 4 | PZ07_Adamiak_4 | 0 | 0 | 0 | 0 | 0 | 0 | 0 |
| 5 | PZ07_Adamiak_5 | 0 | 0 | 0 | 0 | 0 | 0 | 0 |
| 6 | PZ07_Das_5 | 0 | 1 | 2 | 0 | 0 | 0 | 3 |
| 7 | PZ07_Das_8 | 0 | 3 | 5 | 0 | 0 | 0 | 8 |
| 8 | PZ07_Das_3 | 0 | 3 | 5 | 0 | 0 | 1 | 9 |
| 9 | PZ07_Das_7 | 0 | 0 | 0 | 0 | 0 | 1 | 9 |
| 10 | PZ07_Das_6 | 1 | 3 | 6 | 0 | 0 | 0 | 10 |
| 11 | PZ07_Das_2 | 0 | 3 | 6 | 0 | 0 | 3 | 12 |
| 12 | PZ07_Das_4 | 0 | 4 | 6 | 0 | 0 | 2 | 12 |
| 13 | PZ07_Das_1 | 0 | 3 | 6 | 0 | 0 | 2 | 11 |
| 14 | PZ07_refstructure | 0 | 3 | 53 | 0 | 0 | 0 | 56 |
| 15 | PZ07_Ding_8 | 0 | 0 | 127 | 60 | 0 | 0 | 187 |
| 16 | PZ07_Ding_10 | 0 | 1 | 138 | 58 | 0 | 0 | 197 |
| 17 | PZ07_Ding_4 | 0 | 0 | 133 | 64 | 0 | 0 | 197 |
| 18 | PZ07_Ding_7 | 0 | 0 | 149 | 58 | 0 | 0 | 207 |
| 19 | PZ07_Ding_5 | 0 | 1 | 142 | 64 | 0 | 0 | 207 |
| 20 | PZ07_Ding_9 | 0 | 1 | 142 | 66 | 0 | 0 | 209 |
| 21 | PZ07_Ding_6 | 0 | 0 | 151 | 68 | 0 | 0 | 219 |
| 22 | PZ07_Ding_1 | 0 | 0 | 154 | 68 | 0 | 0 | 222 |
| 23 | PZ07_Ding_3 | 0 | 0 | 150 | 74 | 0 | 0 | 224 |
| 24 | PZ07_Ding_2 | 0 | 0 | 170 | 59 | 0 | 0 | 229 |
| 25 | PZ07_Chen_10 | 53 | 54 | 103 | 0 | 0 | 51 | 261 |
| 26 | PZ07_Chen_8 | 86 | 53 | 98 | 0 | 0 | 51 | 288 |
| 27 | PZ07_Chen_3 | 63 | 61 | 118 | 0 | 0 | 51 | 293 |
| 28 | PZ07_Chen_7 | 78 | 55 | 114 | 0 | 0 | 47 | 294 |
| 29 | PZ07_Chen_6 | 73 | 60 | 118 | 0 | 0 | 49 | 300 |
| 30 | PZ07_Bujnicki_5 | 0 | 37 | 212 | 58 | 7 | 0 | 314 |
| 31 | PZ07_Bujnicki_4 | 0 | 38 | 216 | 58 | 7 | 0 | 319 |
| 32 | PZ07_Bujnicki_6 | 0 | 41 | 230 | 50 | 10 | 0 | 331 |
| 33 | PZ07_Dokholyan_1 | 0 | 7 | 199 | 127 | 1 | 0 | 334 |
| 34 | PZ07_Bujnicki_3 | 0 | 44 | 229 | 53 | 9 | 0 | 335 |
| 35 | PZ07_Chen_2 | 131 | 54 | 106 | 0 | 0 | 50 | 341 |
| 36 | PZ07_Dokholyan_2 | 0 | 7 | 199 | 137 | 0 | 0 | 343 |
| 37 | PZ07_Bujnicki_7 | 0 | 33 | 242 | 64 | 5 | 0 | 344 |
| 38 | PZ07_Chen_5 | 126 | 59 | 113 | 0 | 0 | 49 | 347 |
| 39 | PZ07_Bujnicki_2 | 0 | 47 | 252 | 52 | 9 | 0 | 360 |
| 40 | PZ07_Bujnicki_1 | 0 | 44 | 259 | 50 | 10 | 0 | 363 |
| 41 | PZ07_Chen_4 | 172 | 62 | 125 | 0 | 0 | 50 | 409 |
| 42 | PZ07_Chen_9 | 214 | 53 | 96 | 0 | 0 | 52 | 415 |

| | | | | | | | | |
|---|---|---|---|---|---|---|---|---|
| 43 | PZ07_Major_4 | 0 | 59 | 379 | 113 | 6 | 2 | 559 |
| 44 | PZ07_Major_7 | 0 | 61 | 390 | 110 | 4 | 3 | 568 |
| 45 | PZ07_Major_5 | 0 | 57 | 394 | 112 | 4 | 2 | 569 |
| 46 | PZ07_Major_8 | 0 | 62 | 428 | 110 | 0 | 0 | 600 |
| 47 | PZ07_Major_1 | 0 | 57 | 434 | 112 | 4 | 2 | 609 |
| 48 | PZ07_Major_6 | 0 | 76 | 431 | 113 | 3 | 3 | 626 |
| 49 | PZ07_Major_3 | 0 | 80 | 471 | 119 | 6 | 5 | 681 |
| 50 | PZ07_Major_9 | 0 | 88 | 483 | 112 | 4 | 3 | 690 |
| 51 | PZ07_Major_2 | 0 | 89 | 491 | 109 | 6 | 4 | 699 |
| 52 | PZ07_Major_10 | 0 | 82 | 496 | 116 | 5 | 4 | 703 |
| 53 | PZ07_Chen_1 | 0 | 371 | 555 | 98 | 3 | 0 | 1027 |
| Average | | 18.81 | 36.17 | 179.74 | 47.40 | 1.94 | 9.19 | 293.40 |
| Standard deviation | | 47.08 | 55.71 | 160.40 | 46.97 | 3.10 | 18.69 | 238.71 |
| Median | | 0 | 33 | 142 | 53 | 0 | 0 | 293 |

**Table S9**. MAXIT report on structures in Puzzle 08.

| **No.** | **3D RNA model** | **Close contacts** | **Bond lengths** | **Bond angles** | **Planarity** | **Chirality** | **Polymer linkage** | **Total** |
|---|---|---|---|---|---|---|---|---|
| 1 | PZ08_Adamiak_1 | 0 | 0 | 0 | 0 | 0 | 0 | 0 |
| 2 | PZ08_Adamiak_2 | 0 | 0 | 0 | 0 | 0 | 0 | 0 |
| 3 | PZ08_Das_4 | 0 | 0 | 0 | 0 | 0 | 0 | 0 |
| 4 | PZ08_Das_6 | 0 | 0 | 0 | 0 | 0 | 0 | 0 |
| 5 | PZ08_Das_5 | 0 | 0 | 1 | 0 | 0 | 0 | 1 |
| 6 | PZ08_Das_2 | 2 | 0 | 0 | 0 | 0 | 0 | 2 |
| 7 | PZ08_Das_1 | 2 | 0 | 1 | 0 | 0 | 0 | 3 |
| 8 | PZ08_refstructure | 0 | 0 | 1 | 3 | 0 | 0 | 4 |
| 9 | PZ08_Das_3 | 2 | 1 | 2 | 0 | 0 | 1 | 6 |
| 10 | PZ08_Chen_2 | 0 | 0 | 52 | 19 | 0 | 0 | 71 |
| 11 | PZ08_Chen_5 | 0 | 2 | 59 | 16 | 0 | 0 | 77 |
| 12 | PZ08_Chen_3 | 0 | 5 | 68 | 18 | 0 | 0 | 91 |
| 13 | PZ08_Chen_4 | 0 | 0 | 75 | 22 | 0 | 0 | 97 |
| 14 | PZ08_Chen_10 | 0 | 5 | 70 | 21 | 2 | 0 | 98 |
| 15 | PZ08_Ding_7 | 0 | 0 | 65 | 38 | 0 | 0 | 103 |
| 16 | PZ08_Ding_8 | 0 | 0 | 68 | 35 | 0 | 0 | 103 |
| 17 | PZ08_Ding_4 | 0 | 1 | 72 | 34 | 0 | 0 | 107 |
| 18 | PZ08_Ding_5 | 0 | 0 | 79 | 30 | 0 | 0 | 109 |
| 19 | PZ08_Ding_6 | 0 | 0 | 76 | 37 | 0 | 0 | 113 |
| 20 | PZ08_Ding_1 | 0 | 0 | 78 | 36 | 0 | 0 | 114 |
| 21 | PZ08_Ding_10 | 0 | 0 | 83 | 35 | 0 | 0 | 118 |
| 22 | PZ08_Ding_9 | 0 | 0 | 82 | 37 | 0 | 0 | 119 |
| 23 | PZ08_Ding_2 | 0 | 0 | 84 | 38 | 0 | 0 | 122 |
| 24 | PZ08_Ding_3 | 0 | 0 | 96 | 35 | 0 | 0 | 131 |
| 25 | PZ08_Chen_1 | 0 | 19 | 91 | 22 | 1 | 0 | 133 |
| 26 | PZ08_Bujnicki_7 | 0 | 20 | 105 | 15 | 8 | 0 | 148 |

| | | | | | | | | |
|---|---|---|---|---|---|---|---|---|
| 27 | PZ08_Dokholyan_1 | 0 | 3 | 117 | 72 | 0 | 0 | 192 |
| 28 | PZ08_Dokholyan_3 | 0 | 1 | 126 | 73 | 0 | 0 | 200 |
| 29 | PZ08_Dokholyan_4 | 0 | 1 | 126 | 73 | 0 | 0 | 200 |
| 30 | PZ08_Chen_7 | 0 | 33 | 141 | 28 | 1 | 0 | 203 |
| 31 | PZ08_Bujnicki_9 | 0 | 29 | 141 | 33 | 2 | 0 | 205 |
| 32 | PZ08_Dokholyan_2 | 0 | 3 | 131 | 75 | 0 | 0 | 209 |
| 33 | PZ08_Bujnicki_8 | 0 | 29 | 163 | 37 | 8 | 0 | 237 |
| 34 | PZ08_Bujnicki_10 | 0 | 28 | 173 | 29 | 8 | 3 | 241 |
| 35 | PZ08_Bujnicki_6 | 0 | 28 | 203 | 26 | 4 | 0 | 261 |
| 36 | PZ08_Bujnicki_1 | 0 | 30 | 204 | 25 | 3 | 1 | 263 |
| 37 | PZ08_Bujnicki_4 | 0 | 32 | 235 | 22 | 1 | 1 | 291 |
| 38 | PZ08_Bujnicki_5 | 0 | 44 | 225 | 23 | 3 | 0 | 295 |
| 39 | PZ08_Bujnicki_2 | 0 | 30 | 254 | 26 | 4 | 0 | 314 |
| 40 | PZ08_Bujnicki_3 | 0 | 46 | 248 | 23 | 3 | 0 | 320 |
| 41 | PZ08_Chen_8 | 0 | 197 | 284 | 49 | 1 | 0 | 531 |
| 42 | PZ08_Chen_6 | 0 | 186 | 312 | 47 | 0 | 0 | 545 |
| 43 | PZ08_Chen_9 | 0 | 211 | 365 | 61 | 1 | 0 | 638 |
| Average | | 0.14 | 22.88 | 110.60 | 28.21 | 1.16 | 0.14 | 163.14 |
| Standard deviation | | 0.52 | 50.58 | 92.76 | 20.94 | 2.21 | 0.52 | 147.84 |
| Median | | 0 | 1 | 83 | 26 | 0 | 0 | 119 |

**Table S10**. MAXIT report on structures in Puzzle 09.

| **No.** | **3D RNA model** | **Close con-tacts** | **Bond lengths** | **Bond angles** | **Plana-rity** | **Chira-lity** | **Poly-mer lin-kage** | **Total** |
|---|---|---|---|---|---|---|---|---|
| 1 | PZ09_Das_6 | 0 | 0 | 0 | 1 | 0 | 0 | 1 |
| 2 | PZ09_Das_7 | 0 | 0 | 0 | 1 | 0 | 0 | 1 |
| 3 | PZ09_refstructure | 0 | 0 | 2 | 0 | 0 | 0 | 2 |
| 4 | PZ09_Das_8 | 0 | 0 | 1 | 1 | 0 | 0 | 2 |
| 5 | PZ09_Das_9 | 0 | 0 | 1 | 1 | 0 | 0 | 2 |
| 6 | PZ09_Das_1 | 0 | 1 | 3 | 1 | 0 | 0 | 5 |
| 7 | PZ09_Das_3 | 0 | 1 | 3 | 1 | 0 | 0 | 5 |
| 8 | PZ09_Das_4 | 0 | 1 | 3 | 1 | 0 | 0 | 5 |
| 9 | PZ09_Das_2 | 0 | 1 | 4 | 1 | 0 | 0 | 6 |
| 10 | PZ09_Das_5 | 0 | 1 | 4 | 1 | 0 | 0 | 6 |
| 11 | PZ09_Chen_4 | 17 | 0 | 2 | 0 | 0 | 2 | 21 |
| 12 | PZ09_Chen_3 | 16 | 1 | 5 | 0 | 0 | 2 | 24 |
| 13 | PZ09_Chen_2 | 20 | 1 | 4 | 0 | 0 | 1 | 26 |
| 14 | PZ09_Chen_1 | 7 | 3 | 21 | 0 | 0 | 2 | 33 |
| 15 | PZ09_Chen_5 | 22 | 2 | 8 | 0 | 0 | 1 | 33 |
| 16 | PZ09_Chen_7 | 24 | 3 | 17 | 0 | 0 | 1 | 45 |
| 17 | PZ09_Ding_5 | 0 | 0 | 42 | 24 | 0 | 0 | 66 |
| 18 | PZ09_Ding_7 | 0 | 0 | 57 | 14 | 0 | 0 | 71 |
| 19 | PZ09_Ding_6 | 0 | 0 | 52 | 21 | 0 | 0 | 73 |
| 20 | PZ09_Chen_8 | 37 | 5 | 26 | 0 | 0 | 12 | 80 |

| | | | | | | | | |
|---|---|---|---|---|---|---|---|---|
| 21 | PZ09_Ding_8 | 0 | 1 | 62 | 20 | 0 | 0 | 83 |
| 22 | PZ09_Ding_4 | 0 | 0 | 59 | 26 | 0 | 0 | 85 |
| 23 | PZ09_Ding_9 | 0 | 1 | 64 | 20 | 0 | 0 | 85 |
| 24 | PZ09_Ding_10 | 0 | 0 | 60 | 27 | 0 | 0 | 87 |
| 25 | PZ09_Ding_3 | 0 | 0 | 59 | 29 | 0 | 0 | 88 |
| 26 | PZ09_Chen_6 | 46 | 6 | 28 | 0 | 0 | 13 | 93 |
| 27 | PZ09_Ding_1 | 0 | 1 | 69 | 25 | 0 | 0 | 95 |
| 28 | PZ09_Ding_2 | 0 | 0 | 78 | 24 | 0 | 0 | 102 |
| 29 | PZ09_Dokholyan_3 | 0 | 2 | 68 | 49 | 0 | 0 | 119 |
| 30 | PZ09_Dokholyan_1 | 0 | 1 | 76 | 48 | 0 | 0 | 125 |
| 31 | PZ09_Bujnicki_3 | 0 | 14 | 86 | 15 | 12 | 0 | 127 |
| 32 | PZ09_Bujnicki_4 | 0 | 14 | 86 | 15 | 12 | 0 | 127 |
| 33 | PZ09_Bujnicki_5 | 0 | 14 | 86 | 15 | 12 | 0 | 127 |
| 34 | PZ09_Dokholyan_4 | 0 | 4 | 76 | 54 | 0 | 0 | 134 |
| 35 | PZ09_Dokholyan_2 | 0 | 4 | 81 | 53 | 0 | 0 | 138 |
| Average | | 5.40 | 2.34 | 36.94 | 13.94 | 1.03 | 0.97 | 60.63 |
| Standard deviation | | 11.53 | 3.93 | 32.80 | 16.95 | 3.41 | 2.95 | 48.03 |
| Median | | 0 | 1 | 28 | 1 | 0 | 0 | 71 |

**Table S11**. MAXIT report on structures in Puzzle 10.

| No. | 3D RNA model | Close contacts | Bond lengths | Bond angles | Planarity | Chirality | Polymer linkage | Total |
|---|---|---|---|---|---|---|---|---|
| 1 | PZ10_Das_5 | 1 | 3 | 7 | 0 | 0 | 2 | 13 |
| 2 | PZ10_refstructure | 15 | 0 | 0 | 0 | 0 | 0 | 15 |
| 3 | PZ10_Das_4 | 1 | 4 | 11 | 0 | 0 | 2 | 18 |
| 4 | PZ10_Das_1 | 1 | 4 | 11 | 0 | 0 | 3 | 19 |
| 5 | PZ10_Das_2 | 1 | 5 | 11 | 0 | 0 | 2 | 19 |
| 6 | PZ10_Das_3 | 1 | 5 | 13 | 0 | 0 | 3 | 22 |
| 7 | PZ10_Dokholyan_7 | 0 | 1 | 126 | 51 | 0 | 0 | 178 |
| 8 | PZ10_Dokholyan_5 | 0 | 0 | 125 | 61 | 0 | 0 | 186 |
| 9 | PZ10_Dokholyan_4 | 0 | 1 | 129 | 59 | 0 | 0 | 189 |
| 10 | PZ10_Dokholyan_3 | 0 | 1 | 129 | 62 | 0 | 0 | 192 |
| 11 | PZ10_Dokholyan_6 | 0 | 0 | 134 | 58 | 1 | 0 | 193 |
| 12 | PZ10_Dokholyan_9 | 0 | 0 | 133 | 60 | 0 | 0 | 193 |
| 13 | PZ10_Dokholyan_1 | 0 | 0 | 137 | 58 | 0 | 0 | 195 |
| 14 | PZ10_Dokholyan_2 | 0 | 1 | 138 | 66 | 0 | 0 | 205 |
| 15 | PZ10_Dokholyan_8 | 0 | 1 | 144 | 66 | 0 | 0 | 211 |
| 16 | PZ10_Dokholyan_10 | 0 | 0 | 142 | 72 | 1 | 0 | 215 |
| 17 | PZ10_Chen_1 | 0 | 35 | 167 | 38 | 2 | 1 | 243 |
| 18 | PZ10_Bujnicki_4 | 0 | 43 | 211 | 47 | 17 | 0 | 318 |
| 19 | PZ10_Bujnicki_3 | 0 | 49 | 212 | 37 | 26 | 0 | 324 |
| 20 | PZ10_Bujnicki_9 | 0 | 49 | 221 | 41 | 18 | 0 | 329 |
| 21 | PZ10_Bujnicki_2 | 0 | 47 | 216 | 48 | 19 | 0 | 330 |
| 22 | PZ10_Bujnicki_1 | 0 | 48 | 220 | 44 | 20 | 0 | 332 |

| | | | | | | | | |
|---|---|---|---|---|---|---|---|---|
| 23 | PZ10_Bujnicki_10 | 0 | 40 | 222 | 51 | 25 | 0 | 338 |
| 24 | PZ10_Bujnicki_6 | 0 | 48 | 227 | 45 | 18 | 0 | 338 |
| 25 | PZ10_Bujnicki_5 | 0 | 43 | 225 | 53 | 18 | 0 | 339 |
| 26 | PZ10_Bujnicki_8 | 0 | 46 | 224 | 56 | 13 | 0 | 339 |
| 27 | PZ10_Bujnicki_7 | 0 | 47 | 251 | 60 | 16 | 0 | 374 |
| Average | | 0.74 | 19.30 | 140.22 | 41.96 | 7.19 | 0.48 | 209.89 |
| Standard deviation | | 2.88 | 21.93 | 81.97 | 24.37 | 9.53 | 0.98 | 122.06 |
| Median | | 0 | 4 | 138 | 51 | 0 | 0 | 205 |

**Table S12**. MAXIT report on structures in Puzzle 11.

| No. | 3D RNA model | Close con-tacts | Bond lengths | Bond angles | Plana-rity | Chira-lity | Poly-mer lin-kage | Total |
|---|---|---|---|---|---|---|---|---|
| 1 | PZ11_Adamiak_10 | 0 | 0 | 0 | 0 | 0 | 0 | 0 |
| 2 | PZ11_Adamiak_1 | 0 | 0 | 0 | 0 | 0 | 0 | 0 |
| 3 | PZ11_Adamiak_2 | 0 | 0 | 0 | 0 | 0 | 0 | 0 |
| 4 | PZ11_Adamiak_3 | 0 | 0 | 0 | 0 | 0 | 0 | 0 |
| 5 | PZ11_Adamiak_4 | 0 | 0 | 0 | 0 | 0 | 0 | 0 |
| 6 | PZ11_Adamiak_5 | 0 | 0 | 0 | 0 | 0 | 0 | 0 |
| 7 | PZ11_Adamiak_6 | 0 | 0 | 0 | 0 | 0 | 0 | 0 |
| 8 | PZ11_Adamiak_7 | 0 | 0 | 0 | 0 | 0 | 0 | 0 |
| 9 | PZ11_Adamiak_8 | 0 | 0 | 0 | 0 | 0 | 0 | 0 |
| 10 | PZ11_Adamiak_9 | 0 | 0 | 0 | 0 | 0 | 0 | 0 |
| 11 | PZ11_Chen_8 | 0 | 0 | 0 | 0 | 0 | 0 | 0 |
| 12 | PZ11_Das_10 | 0 | 0 | 0 | 0 | 0 | 0 | 0 |
| 13 | PZ11_Chen_10 | 0 | 0 | 1 | 0 | 0 | 0 | 1 |
| 14 | PZ11_Das_9 | 1 | 0 | 0 | 0 | 0 | 0 | 1 |
| 15 | PZ11_Chen_6 | 0 | 0 | 0 | 0 | 0 | 1 | 1 |
| 16 | PZ11_Chen_9 | 0 | 0 | 0 | 0 | 0 | 1 | 1 |
| 17 | PZ11_refstructure | 0 | 0 | 1 | 0 | 0 | 0 | 1 |
| 18 | PZ11_Chen_7 | 0 | 0 | 1 | 0 | 0 | 1 | 2 |
| 19 | PZ11_Das_6 | 0 | 2 | 2 | 0 | 0 | 1 | 5 |
| 20 | PZ11_Das_8 | 0 | 2 | 2 | 0 | 0 | 1 | 5 |
| 21 | PZ11_Das_3 | 0 | 2 | 2 | 0 | 0 | 1 | 5 |
| 22 | PZ11_Das_5 | 0 | 2 | 2 | 0 | 0 | 1 | 5 |
| 23 | PZ11_Das_4 | 0 | 2 | 3 | 0 | 0 | 1 | 6 |
| 24 | PZ11_Das_1 | 0 | 3 | 3 | 0 | 0 | 1 | 7 |
| 25 | PZ11_Das_7 | 1 | 4 | 4 | 0 | 0 | 1 | 10 |
| 26 | PZ11_Das_2 | 4 | 2 | 3 | 0 | 0 | 1 | 10 |
| 27 | PZ11_Bujnicki_1 | 0 | 3 | 4 | 12 | 2 | 0 | 21 |
| 28 | PZ11_Bujnicki_5 | 0 | 1 | 7 | 12 | 2 | 0 | 22 |
| 29 | PZ11_Bujnicki_9 | 0 | 3 | 8 | 9 | 4 | 0 | 24 |
| 30 | PZ11_Bujnicki_3 | 0 | 5 | 5 | 9 | 5 | 0 | 24 |
| 31 | PZ11_Bujnicki_2 | 0 | 5 | 8 | 10 | 3 | 0 | 26 |
| 32 | PZ11_Bujnicki_7 | 0 | 1 | 9 | 16 | 2 | 0 | 28 |

| | | | | | | | | |
|---|---|---|---|---|---|---|---|---|
| 33 | PZ11_Bujnicki_10 | 0 | 3 | 10 | 10 | 7 | 0 | 30 |
| 34 | PZ11_Bujnicki_8 | 0 | 4 | 12 | 17 | 3 | 0 | 36 |
| 35 | PZ11_Bujnicki_6 | 0 | 5 | 11 | 18 | 3 | 0 | 37 |
| 36 | PZ11_Chen_4 | 0 | 0 | 34 | 5 | 0 | 0 | 39 |
| 37 | PZ11_Chen_3 | 0 | 3 | 37 | 6 | 0 | 0 | 46 |
| 38 | PZ11_Chen_1 | 0 | 2 | 42 | 6 | 0 | 0 | 50 |
| 39 | PZ11_Chen_2 | 0 | 5 | 40 | 7 | 0 | 0 | 52 |
| 40 | PZ11_Chen_5 | 0 | 1 | 42 | 9 | 0 | 0 | 52 |
| 41 | PZ11_Ding_3 | 0 | 0 | 37 | 15 | 0 | 0 | 52 |
| 42 | PZ11_Ding_6 | 0 | 0 | 38 | 19 | 0 | 0 | 57 |
| 43 | PZ11_Ding_1 | 0 | 0 | 41 | 19 | 0 | 0 | 60 |
| 44 | PZ11_Ding_8 | 0 | 0 | 44 | 16 | 0 | 0 | 60 |
| 45 | PZ11_Ding_9 | 0 | 0 | 43 | 20 | 0 | 0 | 63 |
| 46 | PZ11_Ding_5 | 0 | 0 | 48 | 20 | 0 | 0 | 68 |
| 47 | PZ11_Ding_7 | 0 | 0 | 49 | 23 | 0 | 0 | 72 |
| 48 | PZ11_Ding_10 | 0 | 0 | 55 | 18 | 0 | 0 | 73 |
| 49 | PZ11_Ding_4 | 0 | 0 | 54 | 21 | 0 | 0 | 75 |
| 50 | PZ11_Ding_2 | 0 | 0 | 52 | 24 | 0 | 0 | 76 |
| 51 | PZ11_Xiao_3 | 0 | 0 | 68 | 28 | 0 | 0 | 96 |
| 52 | PZ11_Xiao_2 | 0 | 0 | 75 | 29 | 0 | 0 | 104 |
| 53 | PZ11_Xiao_1 | 0 | 0 | 80 | 28 | 0 | 0 | 108 |
| 54 | PZ11_Bujnicki_4 | 0 | 78 | 254 | 37 | 5 | 0 | 374 |
| Average | | 0.11 | 2.56 | 22.80 | 8.57 | 0.67 | 0.20 | 34.91 |
| Standard deviation | | 0.57 | 10.59 | 39.60 | 10.12 | 1.57 | 0.41 | 56.38 |
| Median | | 0 | 0 | 4.5 | 5.5 | 0 | 0 | 21.5 |

**Table S13**. MAXIT report on structures in Puzzle 12.

| No. | 3D RNA model | Close con-tacts | Bond lengths | Bond angles | Plana-rity | Chira-lity | Poly-mer lin-kage | Total |
|---|---|---|---|---|---|---|---|---|
| 1 | PZ12_Adamiak_1 | 0 | 0 | 0 | 0 | 0 | 0 | 0 |
| 2 | PZ12_Adamiak_2 | 0 | 0 | 0 | 0 | 0 | 0 | 0 |
| 3 | PZ12_Adamiak_3 | 0 | 0 | 0 | 0 | 0 | 0 | 0 |
| 4 | PZ12_Bujnicki_6 | 1 | 0 | 0 | 0 | 0 | 0 | 1 |
| 5 | PZ12_Das_5 | 1 | 0 | 1 | 0 | 0 | 0 | 2 |
| 6 | PZ12_Das_9 | 0 | 1 | 1 | 0 | 0 | 0 | 2 |
| 7 | PZ12_Das_8 | 0 | 1 | 1 | 0 | 0 | 0 | 2 |
| 8 | PZ12_Bujnicki_7 | 2 | 0 | 0 | 0 | 0 | 0 | 2 |
| 9 | PZ12_Bujnicki_8 | 3 | 0 | 0 | 0 | 0 | 0 | 3 |
| 10 | PZ12_Bujnicki_10 | 4 | 0 | 0 | 0 | 0 | 0 | 4 |
| 11 | PZ12_Das_3 | 0 | 1 | 3 | 0 | 0 | 0 | 4 |
| 12 | PZ12_Das_10 | 0 | 1 | 2 | 0 | 0 | 1 | 4 |
| 13 | PZ12_Bujnicki_1 | 4 | 0 | 2 | 0 | 0 | 0 | 6 |
| 14 | PZ12_Das_1 | 0 | 1 | 4 | 0 | 0 | 1 | 6 |
| 15 | PZ12_Bujnicki_5 | 1 | 0 | 5 | 0 | 0 | 0 | 6 |

| | | | | | | | | |
|---|---|---|---|---|---|---|---|---|
| 16 | PZ12_Bujnicki_4 | 3 | 0 | 3 | 0 | 0 | 0 | 6 |
| 17 | PZ12_Das_4 | 2 | 3 | 1 | 0 | 0 | 1 | 7 |
| 18 | PZ12_Bujnicki_3 | 5 | 0 | 2 | 0 | 0 | 0 | 7 |
| 19 | PZ12_Bujnicki_9 | 3 | 0 | 5 | 0 | 0 | 0 | 8 |
| 20 | PZ12_refstructure | 0 | 0 | 7 | 0 | 0 | 2 | 9 |
| 21 | PZ12_Das_2 | 0 | 2 | 6 | 0 | 0 | 1 | 9 |
| 22 | PZ12_Das_7 | 0 | 3 | 4 | 0 | 0 | 2 | 9 |
| 23 | PZ12_Das_6 | 1 | 3 | 5 | 0 | 0 | 2 | 11 |
| 24 | PZ12_Bujnicki_2 | 0 | 0 | 24 | 0 | 0 | 0 | 24 |
| 25 | PZ12_Chen_1 | 0 | 2 | 61 | 14 | 0 | 0 | 77 |
| 26 | PZ12_Chen_3 | 0 | 5 | 66 | 19 | 0 | 0 | 90 |
| 27 | PZ12_Chen_2 | 0 | 0 | 61 | 27 | 2 | 0 | 90 |
| 28 | PZ12_Chen_4 | 0 | 11 | 81 | 10 | 1 | 0 | 103 |
| 29 | PZ12_Chen_7 | 0 | 8 | 81 | 14 | 1 | 0 | 104 |
| 30 | PZ12_Chen_6 | 0 | 17 | 81 | 20 | 0 | 0 | 118 |
| 31 | PZ12_Chen_5 | 0 | 17 | 98 | 11 | 0 | 0 | 126 |
| 32 | PZ12_Ding_6 | 0 | 2 | 88 | 46 | 0 | 0 | 136 |
| 33 | PZ12_Ding_1 | 0 | 0 | 89 | 48 | 0 | 0 | 137 |
| 34 | PZ12_Ding_10 | 0 | 1 | 87 | 54 | 0 | 0 | 142 |
| 35 | PZ12_Ding_5 | 0 | 1 | 98 | 44 | 0 | 0 | 143 |
| 36 | PZ12_Ding_3 | 7 | 0 | 88 | 50 | 0 | 0 | 145 |
| 37 | PZ12_Ding_11 | 7 | 0 | 108 | 38 | 0 | 0 | 153 |
| 38 | PZ12_Ding_4 | 0 | 0 | 102 | 51 | 0 | 0 | 153 |
| 39 | PZ12_Ding_7 | 0 | 1 | 98 | 54 | 1 | 0 | 154 |
| 40 | PZ12_Ding_12 | 0 | 0 | 110 | 51 | 0 | 0 | 161 |
| 41 | PZ12_Ding_2 | 0 | 0 | 113 | 48 | 0 | 0 | 161 |
| 42 | PZ12_Ding_9 | 0 | 1 | 112 | 50 | 0 | 0 | 163 |
| 43 | PZ12_Chen_9 | 0 | 29 | 120 | 15 | 2 | 1 | 167 |
| 44 | PZ12_Ding_8 | 0 | 0 | 117 | 57 | 0 | 0 | 174 |
| 45 | PZ12_Weeks_2 | 0 | 0 | 94 | 89 | 0 | 0 | 183 |
| 46 | PZ12_Chen_10 | 0 | 41 | 123 | 21 | 3 | 2 | 190 |
| 47 | PZ12_Weeks_1 | 0 | 1 | 105 | 86 | 0 | 0 | 192 |
| 48 | PZ12_Chen_8 | 0 | 36 | 143 | 13 | 0 | 1 | 193 |
| 49 | PZ12_Weeks_3 | 0 | 4 | 127 | 87 | 0 | 0 | 218 |
| 50 | PZ12_Xiao_3 | 0 | 29 | 252 | 51 | 6 | 0 | 338 |
| 51 | PZ12_Xiao_2 | 3 | 68 | 376 | 59 | 6 | 0 | 512 |
| 52 | PZ12_Xiao_1 | 4 | 67 | 384 | 59 | 1 | 1 | 516 |
| Average | | 0.98 | 6.87 | 68.06 | 22.81 | 0.44 | 0.29 | 99.44 |
| Standard deviation | | 1.82 | 15.45 | 84.33 | 27.20 | 1.27 | 0.61 | 117.18 |
| Median | | 0 | 1 | 63.5 | 12 | 0 | 0 | 90 |

**Table S14**. MAXIT report on structures in Puzzle 13.

| No. | 3D RNA model | Close con-tacts | Bond lengths | Bond angles | Plana-rity | Chira-lity | Poly-mer lin-kage | Total |
|---|---|---|---|---|---|---|---|---|
| 1 | PZ13_Adamiak_1 | 0 | 0 | 0 | 0 | 0 | 0 | 0 |
| 2 | PZ13_Das_10 | 0 | 0 | 0 | 0 | 0 | 0 | 0 |
| 3 | PZ13_Das_4 | 0 | 0 | 0 | 0 | 0 | 0 | 0 |
| 4 | PZ13_Das_1 | 0 | 0 | 0 | 0 | 0 | 0 | 0 |
| 5 | PZ13_Das_6 | 0 | 0 | 0 | 0 | 0 | 0 | 0 |
| 6 | PZ13_Das_9 | 0 | 0 | 0 | 0 | 0 | 0 | 0 |
| 7 | PZ13_Das_2 | 0 | 0 | 0 | 0 | 0 | 0 | 0 |
| 8 | PZ13_Das_5 | 0 | 0 | 0 | 0 | 0 | 0 | 0 |
| 9 | PZ13_refstructure | 0 | 0 | 1 | 0 | 0 | 1 | 2 |
| 10 | PZ13_Das_8 | 0 | 1 | 1 | 0 | 0 | 0 | 2 |
| 11 | PZ13_Das_7 | 0 | 0 | 3 | 0 | 1 | 0 | 4 |
| 12 | PZ13_Das_3 | 0 | 2 | 2 | 0 | 6 | 1 | 11 |
| 13 | PZ13_Bujnicki_8 | 0 | 9 | 12 | 10 | 0 | 0 | 31 |
| 14 | PZ13_Bujnicki_10 | 0 | 11 | 17 | 8 | 4 | 0 | 40 |
| 15 | PZ13_Chen_1 | 0 | 0 | 37 | 7 | 2 | 0 | 46 |
| 16 | PZ13_Chen_4 | 0 | 0 | 43 | 5 | 2 | 0 | 50 |
| 17 | PZ13_Ding_7 | 0 | 0 | 37 | 14 | 0 | 0 | 51 |
| 18 | PZ13_Chen_6 | 0 | 3 | 37 | 11 | 1 | 0 | 52 |
| 19 | PZ13_Chen_3 | 0 | 1 | 42 | 16 | 1 | 0 | 60 |
| 20 | PZ13_Ding_5 | 0 | 0 | 38 | 23 | 0 | 0 | 61 |
| 21 | PZ13_Ding_3 | 0 | 0 | 43 | 21 | 0 | 0 | 64 |
| 22 | PZ13_Bujnicki_9 | 0 | 11 | 37 | 15 | 5 | 0 | 68 |
| 23 | PZ13_Ding_8 | 0 | 0 | 49 | 20 | 0 | 0 | 69 |
| 24 | PZ13_Ding_10 | 0 | 0 | 43 | 27 | 0 | 0 | 70 |
| 25 | PZ13_Ding_4 | 0 | 0 | 50 | 21 | 0 | 0 | 71 |
| 26 | PZ13_Chen_8 | 0 | 2 | 53 | 16 | 1 | 0 | 72 |
| 27 | PZ13_Ding_1 | 0 | 0 | 46 | 26 | 0 | 0 | 72 |
| 28 | PZ13_Ding_9 | 0 | 0 | 58 | 20 | 0 | 0 | 78 |
| 29 | PZ13_Chen_7 | 0 | 2 | 65 | 11 | 1 | 0 | 79 |
| 30 | PZ13_Ding_6 | 0 | 0 | 53 | 26 | 0 | 0 | 79 |
| 31 | PZ13_Bujnicki_6 | 0 | 13 | 52 | 14 | 0 | 0 | 79 |
| 32 | PZ13_Chen_9 | 0 | 13 | 48 | 15 | 4 | 0 | 80 |
| 33 | PZ13_Chen_5 | 0 | 2 | 61 | 14 | 3 | 0 | 80 |
| 34 | PZ13_Ding_2 | 0 | 0 | 63 | 27 | 0 | 0 | 90 |
| 35 | PZ13_Chen_2 | 0 | 7 | 69 | 12 | 5 | 0 | 93 |
| 36 | PZ13_Dokholyan_2 | 0 | 4 | 65 | 50 | 0 | 0 | 119 |
| 37 | PZ13_Bujnicki_7 | 0 | 16 | 92 | 12 | 2 | 0 | 122 |
| 38 | PZ13_Bujnicki_5 | 0 | 14 | 94 | 17 | 2 | 0 | 127 |
| 39 | PZ13_Xiao_10 | 0 | 0 | 100 | 26 | 1 | 0 | 127 |
| 40 | PZ13_Dokholyan_3 | 0 | 7 | 74 | 47 | 0 | 0 | 128 |
| 41 | PZ13_Dokholyan_1 | 0 | 6 | 79 | 45 | 0 | 0 | 130 |
| 42 | PZ13_Xiao_6 | 0 | 1 | 112 | 22 | 2 | 0 | 137 |

| | | | | | | | | |
|---|---|---|---|---|---|---|---|---|
| 43 | PZ13_Dokholyan_4 | 0 | 1 | 87 | 49 | 2 | 0 | 139 |
| 44 | PZ13_Bujnicki_2 | 0 | 20 | 101 | 18 | 1 | 0 | 140 |
| 45 | PZ13_Xiao_8 | 0 | 1 | 117 | 24 | 3 | 0 | 145 |
| 46 | PZ13_Xiao_9 | 0 | 2 | 116 | 32 | 3 | 0 | 153 |
| 47 | PZ13_Dokholyan_5 | 3 | 2 | 100 | 52 | 0 | 0 | 157 |
| 48 | PZ13_Bujnicki_3 | 0 | 25 | 120 | 16 | 2 | 0 | 163 |
| 49 | PZ13_Xiao_7 | 4 | 37 | 204 | 28 | 1 | 1 | 275 |
| 50 | PZ13_Bujnicki_4 | 0 | 48 | 220 | 22 | 0 | 0 | 290 |
| 51 | PZ13_Bujnicki_1 | 0 | 55 | 224 | 21 | 3 | 0 | 303 |
| 52 | PZ13_Xiao_1 | 1 | 98 | 269 | 37 | 2 | 1 | 408 |
| 53 | PZ13_Xiao_4 | 0 | 93 | 281 | 39 | 1 | 0 | 414 |
| 54 | PZ13_Xiao_2 | 5 | 112 | 279 | 33 | 6 | 1 | 436 |
| 55 | PZ13_Xiao_5 | 0 | 117 | 288 | 38 | 1 | 0 | 444 |
| 56 | PZ13_Xiao_3 | 4 | 119 | 313 | 30 | 5 | 1 | 472 |
| Average | | 0.30 | 15.27 | 78.48 | 18.52 | 1.30 | 0.11 | 113.98 |
| Standard deviation | | 1.06 | 31.51 | 83.43 | 14.55 | 1.72 | 0.31 | 122.69 |
| Median | | 0 | 1.5 | 52.5 | 16.5 | 1 | 0 | 78.5 |

**Table S15**. MAXIT report on structures in Puzzle 14 (bound).

| **No.** | **3D RNA model** | **Close contacts** | **Bond le-ngths** | **Bond angles** | **Plana-rity** | **Chira-lity** | **Polymer linkage** | **Total** |
|---|---|---|---|---|---|---|---|---|
| 1 | PZ14_AdamiakPostExp_1 | 0 | 0 | 0 | 0 | 0 | 0 | 0 |
| 2 | PZ14_AdamiakPostExp_2 | 0 | 0 | 0 | 0 | 0 | 0 | 0 |
| 3 | PZ14_DasPostExp_9 | 0 | 0 | 0 | 0 | 0 | 0 | 0 |
| 4 | PZ14_AdamiakPreExp_1 | 0 | 0 | 0 | 0 | 0 | 0 | 0 |
| 5 | PZ14_DasPreExp_10 | 0 | 0 | 0 | 0 | 0 | 0 | 0 |
| 6 | PZ14_AdamiakPreExp_2 | 0 | 0 | 0 | 0 | 0 | 0 | 0 |
| 7 | PZ14_DasPreExp_1 | 0 | 0 | 0 | 0 | 0 | 0 | 0 |
| 8 | PZ14_DasPreExp_6 | 0 | 0 | 0 | 0 | 0 | 0 | 0 |
| 9 | PZ14_DasPostExp_6 | 0 | 0 | 0 | 0 | 0 | 0 | 0 |
| 10 | PZ14_ChenPostExp_2 | 0 | 0 | 0 | 0 | 0 | 0 | 0 |
| 11 | PZ14_ChenPostExp_10 | 0 | 0 | 0 | 0 | 0 | 0 | 0 |
| 12 | PZ14_ChenPostExp_1 | 0 | 0 | 0 | 0 | 0 | 0 | 0 |
| 13 | PZ14_ChenPostExp_4 | 0 | 0 | 0 | 0 | 0 | 0 | 0 |
| 14 | PZ14_ChenPostExp_5 | 0 | 0 | 0 | 0 | 0 | 0 | 0 |
| 15 | PZ14_ChenPostExp_8 | 0 | 0 | 0 | 0 | 0 | 0 | 0 |
| 16 | PZ14_ChenPostExp_7 | 0 | 0 | 0 | 0 | 0 | 0 | 0 |
| 17 | PZ14_ChenPostExp_9 | 0 | 0 | 0 | 0 | 0 | 0 | 0 |
| 18 | PZ14_ChenPostExp_6 | 0 | 0 | 0 | 0 | 0 | 0 | 0 |
| 19 | PZ14_ChenPostExp_3 | 0 | 0 | 0 | 0 | 0 | 0 | 0 |
| 20 | PZ14_DasPostExp_4 | 0 | 0 | 1 | 0 | 0 | 0 | 1 |
| 21 | PZ14_DasPostExp_1 | 0 | 0 | 2 | 0 | 0 | 0 | 2 |
| 22 | PZ14_DasPreExp_5 | 4 | 0 | 0 | 0 | 0 | 0 | 4 |
| 23 | PZ14_DasPreExp_7 | 1 | 1 | 2 | 0 | 0 | 0 | 4 |

| | | | | | | | | |
|---|---|---|---|---|---|---|---|---|
| 24 | PZ14_DasPostExp_10 | 4 | 0 | 0 | 0 | 0 | 0 | 4 |
| 25 | PZ14_DasPostExp_2 | 4 | 0 | 0 | 0 | 0 | 0 | 4 |
| 26 | PZ14_DasPostExp_5 | 4 | 0 | 0 | 0 | 0 | 0 | 4 |
| 27 | PZ14_refstructure_bound | 7 | 0 | 1 | 0 | 0 | 0 | 8 |
| 28 | PZ14_DasPostExp_3 | 5 | 1 | 2 | 0 | 0 | 0 | 8 |
| 29 | PZ14_DasPreExp_4 | 8 | 0 | 1 | 0 | 0 | 0 | 9 |
| 30 | PZ14_DasPostExp_7 | 7 | 0 | 4 | 0 | 0 | 0 | 11 |
| 31 | PZ14_DasPreExp_2 | 7 | 0 | 4 | 0 | 0 | 0 | 11 |
| 32 | PZ14_DasPostExp_8 | 10 | 0 | 2 | 0 | 0 | 0 | 12 |
| 33 | PZ14_DasPreExp_3 | 10 | 0 | 2 | 0 | 0 | 0 | 12 |
| 34 | PZ14_DasPreExp_8 | 8 | 0 | 5 | 0 | 0 | 0 | 13 |
| 35 | PZ14_DasPreExp_9 | 11 | 0 | 3 | 0 | 0 | 0 | 14 |
| 36 | PZ14_BujnickiPreExp_4 | 9 | 4 | 7 | 1 | 2 | 0 | 23 |
| 37 | PZ14_BujnickiPreExp_5 | 0 | 15 | 8 | 3 | 2 | 0 | 28 |
| 38 | PZ14_DingPreExp_1 | 0 | 0 | 45 | 17 | 0 | 0 | 62 |
| 39 | PZ14_DingPostExp_3 | 0 | 1 | 47 | 18 | 0 | 0 | 66 |
| 40 | PZ14_DingPreExp_5 | 0 | 0 | 43 | 24 | 0 | 0 | 67 |
| 41 | PZ14_DingPostExp_10 | 0 | 0 | 46 | 24 | 0 | 0 | 70 |
| 42 | PZ14_DingPostExp_6 | 0 | 1 | 45 | 25 | 0 | 0 | 71 |
| 43 | PZ14_DingPostExp_5 | 0 | 0 | 48 | 26 | 0 | 0 | 74 |
| 44 | PZ14_DingPreExp_3 | 0 | 1 | 49 | 24 | 0 | 0 | 74 |
| 45 | PZ14_DingPostExp_4 | 0 | 1 | 44 | 30 | 0 | 0 | 75 |
| 46 | PZ14_DingPostExp_1 | 0 | 0 | 53 | 24 | 0 | 0 | 77 |
| 47 | PZ14_DingPostExp_9 | 0 | 0 | 53 | 25 | 0 | 0 | 78 |
| 48 | PZ14_BujnickiPreExp_6 | 0 | 16 | 53 | 5 | 5 | 0 | 79 |
| 49 | PZ14_DingPreExp_8 | 0 | 0 | 57 | 23 | 0 | 0 | 80 |
| 50 | PZ14_DingPreExp_4 | 0 | 1 | 56 | 24 | 0 | 0 | 81 |
| 51 | PZ14_DingPreExp_6 | 0 | 0 | 57 | 25 | 0 | 0 | 82 |
| 52 | PZ14_DingPreExp_2 | 0 | 0 | 55 | 27 | 0 | 0 | 82 |
| 53 | PZ14_DingPostExp_2 | 0 | 2 | 56 | 25 | 0 | 0 | 83 |
| 54 | PZ14_DingPreExp_7 | 0 | 0 | 60 | 24 | 0 | 0 | 84 |
| 55 | PZ14_DingPostExp_7 | 0 | 2 | 57 | 29 | 0 | 0 | 88 |
| 56 | PZ14_DingPostExp_8 | 0 | 0 | 63 | 24 | 0 | 1 | 88 |
| 57 | PZ14_BujnickiPreExp_7 | 0 | 14 | 62 | 13 | 10 | 0 | 99 |
| 58 | PZ14_BujnickiPostExp_3 | 0 | 23 | 73 | 11 | 9 | 0 | 116 |
| 59 | PZ14_BujnickiPostExp_5 | 2 | 41 | 104 | 24 | 15 | 0 | 186 |
| 60 | PZ14_BujnickiPostExp_1 | 0 | 48 | 125 | 13 | 5 | 0 | 191 |
| 61 | PZ14_BujnickiPostExp_4 | 3 | 54 | 159 | 10 | 3 | 0 | 229 |
| 62 | PZ14_BujnickiPostExp_2 | 3 | 50 | 160 | 24 | 5 | 0 | 242 |
| Average | | 1.73 | 4.45 | 27.65 | 8.74 | 0.90 | 0.02 | 43.48 |
| Standard deviation | | 3.15 | 12.42 | 38.67 | 11.33 | 2.71 | 0.13 | 57.82 |
| Median | | 0 | 0 | 2.5 | 0 | 0 | 0 | 11.5 |

**Table S16**. MAXIT report on structures in Puzzle 14 (free).

| No. | 3D RNA model | Close con-tacts | Bond lengths | Bond angles | Plana-rity | Chira-lity | Poly-mer linkage | Total |
|---|---|---|---|---|---|---|---|---|
| 1 | PZ14_DasPreExp_5 | 0 | 0 | 0 | 0 | 0 | 0 | 0 |
| 2 | PZ14_DasPostExp_10 | 0 | 0 | 0 | 0 | 0 | 0 | 0 |
| 3 | PZ14_DasPostExp_3 | 0 | 0 | 0 | 0 | 0 | 0 | 0 |
| 4 | PZ14_DasPostExp_2 | 0 | 0 | 0 | 0 | 0 | 0 | 0 |
| 5 | PZ14_DasPreExp_1 | 0 | 0 | 0 | 0 | 0 | 0 | 0 |
| 6 | PZ14_DasPostExp_6 | 0 | 0 | 0 | 0 | 0 | 0 | 0 |
| 7 | PZ14_DasPostExp_9 | 0 | 0 | 1 | 0 | 0 | 0 | 1 |
| 8 | PZ14_DasPreExp_10 | 0 | 0 | 1 | 0 | 0 | 0 | 1 |
| 9 | PZ14_DasPostExp_4 | 1 | 0 | 0 | 0 | 0 | 0 | 1 |
| 10 | PZ14_DasPostExp_5 | 0 | 0 | 2 | 0 | 0 | 0 | 2 |
| 11 | PZ14_DasPreExp_2 | 0 | 0 | 3 | 0 | 0 | 0 | 3 |
| 12 | PZ14_DasPostExp_7 | 0 | 0 | 3 | 0 | 0 | 0 | 3 |
| 13 | PZ14_DasPostExp_8 | 0 | 0 | 3 | 0 | 0 | 0 | 3 |
| 14 | PZ14_DasPreExp_4 | 0 | 0 | 3 | 0 | 0 | 0 | 3 |
| 15 | PZ14_DasPreExp_6 | 0 | 1 | 2 | 0 | 0 | 1 | 4 |
| 16 | PZ14_DasPostExp_1 | 0 | 1 | 2 | 0 | 0 | 1 | 4 |
| 17 | PZ14_DasPreExp_9 | 0 | 0 | 4 | 0 | 0 | 0 | 4 |
| 18 | PZ14_DasPreExp_3 | 5 | 0 | 0 | 0 | 0 | 0 | 5 |
| 19 | PZ14_DasPreExp_8 | 0 | 0 | 5 | 0 | 0 | 0 | 5 |
| 20 | PZ14_DasPreExp_7 | 6 | 0 | 0 | 0 | 0 | 0 | 6 |
| 21 | PZ14_refstructure_free | 21 | 0 | 4 | 0 | 0 | 0 | 26 |
| 22 | PZ14_ChenPostExp_2 | 0 | 4 | 33 | 6 | 2 | 1 | 46 |
| 23 | PZ14_ChenPostExp_4 | 0 | 7 | 40 | 7 | 2 | 1 | 57 |
| 24 | PZ14_ChenPostExp_8 | 0 | 7 | 39 | 10 | 0 | 1 | 57 |
| 25 | PZ14_ChenPostExp_1 | 0 | 6 | 45 | 7 | 1 | 0 | 59 |
| 26 | PZ14_ChenPostExp_5 | 0 | 6 | 47 | 4 | 1 | 1 | 59 |
| 27 | PZ14_DingPostExp_1 | 0 | 1 | 34 | 26 | 0 | 0 | 61 |
| 28 | PZ14_DingPreExp_4 | 0 | 0 | 41 | 20 | 0 | 0 | 61 |
| 29 | PZ14_DingPostExp_2 | 0 | 0 | 39 | 23 | 0 | 0 | 62 |
| 30 | PZ14_ChenPostExp_10 | 0 | 15 | 41 | 4 | 2 | 1 | 63 |
| 31 | PZ14_ChenPostExp_6 | 0 | 8 | 44 | 11 | 0 | 1 | 64 |
| 32 | PZ14_ChenPostExp_7 | 0 | 11 | 44 | 10 | 0 | 1 | 66 |
| 33 | PZ14_DingPostExp_5 | 0 | 1 | 43 | 23 | 0 | 0 | 67 |
| 34 | PZ14_ChenPostExp_9 | 0 | 10 | 43 | 12 | 2 | 1 | 68 |
| 35 | PZ14_DingPreExp_1 | 0 | 0 | 46 | 25 | 0 | 0 | 71 |
| 36 | PZ14_DingPostExp_7 | 0 | 0 | 51 | 24 | 0 | 0 | 75 |
| 37 | PZ14_DingPostExp_10 | 0 | 0 | 61 | 14 | 0 | 0 | 75 |
| 38 | PZ14_DingPostExp_3 | 0 | 1 | 49 | 26 | 0 | 0 | 76 |
| 39 | PZ14_DingPostExp_9 | 0 | 0 | 56 | 20 | 0 | 0 | 76 |
| 40 | PZ14_DingPostExp_8 | 0 | 0 | 55 | 22 | 0 | 0 | 77 |
| 41 | PZ14_DingPostExp_4 | 0 | 0 | 53 | 24 | 1 | 0 | 78 |
| 42 | PZ14_DingPostExp_6 | 0 | 0 | 53 | 26 | 0 | 0 | 79 |

| | | | | | | | | |
|---|---|---|---|---|---|---|---|---|
| 43 | PZ14_DingPreExp_3 | 0 | 0 | 52 | 27 | 0 | 0 | 79 |
| 44 | PZ14_BujnickiPreExp_4 | 0 | 13 | 52 | 11 | 6 | 0 | 82 |
| 45 | PZ14_BujnickiPreExp_2 | 0 | 12 | 53 | 8 | 10 | 0 | 83 |
| 46 | PZ14_BujnickiPreExp_3 | 0 | 14 | 53 | 12 | 8 | 0 | 87 |
| 47 | PZ14_DingPreExp_2 | 0 | 1 | 67 | 24 | 0 | 0 | 92 |
| 48 | PZ14_ChenPostExp_3 | 0 | 22 | 61 | 9 | 1 | 1 | 94 |
| 49 | PZ14_BujnickiPostExp_1 | 0 | 19 | 65 | 6 | 4 | 0 | 94 |
| 50 | PZ14_BujnickiPostExp_2 | 0 | 21 | 76 | 10 | 5 | 0 | 112 |
| 51 | PZ14_BujnickiPreExp_1 | 0 | 16 | 96 | 6 | 7 | 0 | 125 |
| 52 | PZ14_BujnickiPostExp_3 | 1 | 26 | 85 | 17 | 3 | 0 | 132 |
| 53 | PZ14_BujnickiPostExp_4 | 0 | 35 | 99 | 23 | 14 | 1 | 172 |
| Average | | 0.64 | 4.87 | 33.00 | 9.38 | 1.30 | 0.23 | 49.43 |
| Standard deviation | | 3.05 | 8.08 | 28.60 | 9.82 | 2.86 | 0.42 | 42.64 |
| Median | | 0 | 0 | 41 | 7 | 0 | 0 | 61 |

**Table S17**. MAXIT report on structures in Puzzle 15.

| No. | 3D RNA model | Close con-tacts | Bond lengths | Bond angles | Plana-rity | Chira-lity | Poly-mer linkage | Total |
|---|---|---|---|---|---|---|---|---|
| 1 | PZ15_FARFAR2_6 | 0 | 0 | 0 | 0 | 0 | 0 | 0 |
| 2 | PZ15_FARFAR2_8 | 0 | 0 | 0 | 0 | 0 | 0 | 0 |
| 3 | PZ15_RNAComposer1_1 | 0 | 0 | 0 | 0 | 0 | 0 | 0 |
| 4 | PZ15_RNAComposer1_2 | 0 | 0 | 0 | 0 | 0 | 0 | 0 |
| 5 | PZ15_RNAComposer2_1 | 0 | 0 | 0 | 0 | 0 | 0 | 0 |
| 6 | PZ15_RNAComposer2_2 | 0 | 0 | 0 | 0 | 0 | 0 | 0 |
| 7 | PZ15_Adamiak_10 | 0 | 0 | 0 | 0 | 0 | 0 | 0 |
| 8 | PZ15_Adamiak_1 | 0 | 0 | 0 | 0 | 0 | 0 | 0 |
| 9 | PZ15_Adamiak_2 | 0 | 0 | 0 | 0 | 0 | 0 | 0 |
| 10 | PZ15_Adamiak_3 | 0 | 0 | 0 | 0 | 0 | 0 | 0 |
| 11 | PZ15_Adamiak_4 | 0 | 0 | 0 | 0 | 0 | 0 | 0 |
| 12 | PZ15_Adamiak_5 | 0 | 0 | 0 | 0 | 0 | 0 | 0 |
| 13 | PZ15_Adamiak_6 | 0 | 0 | 0 | 0 | 0 | 0 | 0 |
| 14 | PZ15_Adamiak_7 | 0 | 0 | 0 | 0 | 0 | 0 | 0 |
| 15 | PZ15_Adamiak_8 | 0 | 0 | 0 | 0 | 0 | 0 | 0 |
| 16 | PZ15_Adamiak_9 | 0 | 0 | 0 | 0 | 0 | 0 | 0 |
| 17 | PZ15_FARFAR2_1 | 0 | 0 | 0 | 0 | 0 | 0 | 0 |
| 18 | PZ15_FARFAR2_4 | 0 | 0 | 0 | 0 | 0 | 0 | 0 |
| 19 | PZ15_refstructure | 0 | 0 | 0 | 0 | 0 | 0 | 0 |
| 20 | PZ15_FARFAR2_7 | 0 | 1 | 0 | 0 | 0 | 0 | 1 |
| 21 | PZ15_FARFAR2_2 | 0 | 0 | 1 | 0 | 0 | 0 | 1 |
| 22 | PZ15_FARFAR2_3 | 0 | 0 | 1 | 0 | 0 | 0 | 1 |
| 23 | PZ15_FARFAR1_8 | 0 | 1 | 1 | 0 | 0 | 0 | 2 |
| 24 | PZ15_FARFAR2_10 | 0 | 0 | 2 | 0 | 0 | 0 | 2 |
| 25 | PZ15_FARFAR2_5 | 0 | 1 | 1 | 0 | 0 | 0 | 2 |
| 26 | PZ15_FARFAR2_9 | 0 | 1 | 2 | 0 | 0 | 0 | 3 |

| | | | | | | | | |
|---|---|---|---|---|---|---|---|---|
| 27 | PZ15_FARFAR1_10 | 0 | 1 | 3 | 0 | 0 | 0 | 4 |
| 28 | PZ15_FARFAR1_1 | 0 | 1 | 3 | 0 | 0 | 0 | 4 |
| 29 | PZ15_FARFAR1_2 | 0 | 1 | 2 | 0 | 0 | 0 | 3 |
| 30 | PZ15_FARFAR1_3 | 0 | 1 | 2 | 0 | 0 | 0 | 3 |
| 31 | PZ15_FARFAR1_5 | 0 | 1 | 3 | 0 | 0 | 0 | 4 |
| 32 | PZ15_FARFAR1_6 | 0 | 1 | 2 | 0 | 0 | 0 | 3 |
| 33 | PZ15_FARFAR1_7 | 0 | 2 | 2 | 0 | 0 | 0 | 4 |
| 34 | PZ15_FARFAR1_9 | 0 | 2 | 2 | 0 | 0 | 0 | 4 |
| 35 | PZ15_FARFAR1_4 | 1 | 2 | 3 | 0 | 0 | 0 | 6 |
| 36 | PZ15_Chen_3 | 0 | 1 | 38 | 8 | 0 | 0 | 47 |
| 37 | PZ15_Chen_2 | 0 | 0 | 40 | 8 | 0 | 0 | 48 |
| 38 | PZ15_Chen_1 | 0 | 2 | 43 | 11 | 0 | 0 | 56 |
| 39 | PZ15_Chen_4 | 0 | 4 | 55 | 10 | 0 | 0 | 69 |
| 40 | PZ15_Chen_6 | 0 | 5 | 54 | 11 | 0 | 0 | 70 |
| 41 | PZ15_Chen_5 | 0 | 6 | 60 | 12 | 0 | 0 | 78 |
| 42 | PZ15_Chen_9 | 0 | 5 | 55 | 16 | 3 | 0 | 79 |
| 43 | PZ15_Chen_8 | 0 | 16 | 57 | 17 | 2 | 0 | 92 |
| 44 | PZ15_Chen_7 | 0 | 12 | 70 | 18 | 0 | 0 | 100 |
| 45 | PZ15_Chen_10 | 0 | 17 | 77 | 13 | 2 | 0 | 109 |
| 46 | PZ15_SimRNA2_10 | 13 | 42 | 122 | 22 | 0 | 12 | 211 |
| 47 | PZ15_SimRNA2_2 | 7 | 49 | 125 | 17 | 0 | 18 | 216 |
| 48 | PZ15_SimRNA1_7 | 10 | 53 | 129 | 15 | 0 | 12 | 219 |
| 49 | PZ15_SimRNA2_4 | 8 | 47 | 145 | 17 | 0 | 13 | 230 |
| 50 | PZ15_SimRNA1_6 | 10 | 49 | 141 | 16 | 1 | 18 | 235 |
| 51 | PZ15_SimRNA1_1 | 21 | 49 | 133 | 14 | 1 | 17 | 235 |
| 52 | PZ15_SimRNA1_10 | 10 | 51 | 139 | 15 | 3 | 18 | 236 |
| 53 | PZ15_SimRNA1_8 | 12 | 55 | 132 | 21 | 2 | 15 | 237 |
| 54 | PZ15_SimRNA1_5 | 14 | 52 | 139 | 16 | 2 | 16 | 239 |
| 55 | PZ15_SimRNA1_2 | 14 | 45 | 152 | 22 | 1 | 13 | 247 |
| 56 | PZ15_SimRNA1_9 | 24 | 52 | 136 | 18 | 1 | 20 | 251 |
| 57 | PZ15_SimRNA1_3 | 18 | 51 | 146 | 21 | 0 | 16 | 252 |
| 58 | PZ15_SimRNA2_9 | 15 | 56 | 148 | 19 | 3 | 16 | 257 |
| 59 | PZ15_SimRNA2_5 | 33 | 54 | 139 | 18 | 0 | 13 | 257 |
| 60 | PZ15_SimRNA2_7 | 35 | 49 | 137 | 19 | 1 | 19 | 260 |
| 61 | PZ15_3dRNA2_8 | 2 | 19 | 183 | 35 | 22 | 1 | 262 |
| 62 | PZ15_SimRNA2_6 | 20 | 56 | 150 | 18 | 1 | 18 | 263 |
| 63 | PZ15_SimRNA2_8 | 6 | 56 | 156 | 19 | 3 | 24 | 264 |
| 64 | PZ15_SimRNA2_1 | 7 | 51 | 162 | 25 | 2 | 17 | 264 |
| 65 | PZ15_3dRNA2_10 | 1 | 21 | 186 | 35 | 22 | 1 | 266 |
| 66 | PZ15_3dRNA2_1 | 2 | 20 | 193 | 33 | 22 | 1 | 271 |
| 67 | PZ15_3dRNA2_2 | 2 | 21 | 191 | 35 | 22 | 1 | 272 |
| 68 | PZ15_3dRNA2_4 | 2 | 18 | 192 | 37 | 22 | 1 | 272 |
| 69 | PZ15_3dRNA2_7 | 2 | 22 | 189 | 36 | 22 | 2 | 273 |
| 70 | PZ15_3dRNA2_6 | 3 | 23 | 193 | 32 | 22 | 2 | 275 |
| 71 | PZ15_3dRNA2_3 | 2 | 22 | 191 | 37 | 22 | 2 | 276 |

| | | | | | | | | |
|---|---|---|---|---|---|---|---|---|
| 72 | PZ15_3dRNA2_9 | 2 | 17 | 199 | 36 | 22 | 1 | 277 |
| 73 | PZ15_SimRNA1_4 | 26 | 57 | 144 | 28 | 1 | 24 | 280 |
| 74 | PZ15_SimRNA2_3 | 33 | 56 | 147 | 23 | 1 | 21 | 281 |
| 75 | PZ15_3dRNA2_5 | 2 | 21 | 199 | 35 | 22 | 2 | 281 |
| Average | | 4.76 | 17.57 | 70.89 | 11.44 | 3.33 | 4.72 | 112.72 |
| Standard deviation | | 8.69 | 21.81 | 75.53 | 12.61 | 7.42 | 7.68 | 120.51 |
| Median | | 0 | 2 | 43 | 10 | 0 | 0 | 56 |

**Table S18**. MAXIT report on structures in Puzzle 17.

| **No.** | **3D RNA model** | **Close con-tacts** | **Bond lengths** | **Bond angles** | **Plana-rity** | **Chira-lity** | **Poly-mer linkage** | **Total** |
|---|---|---|---|---|---|---|---|---|
| 1 | PZ17_Adamiak_1 | 0 | 0 | 0 | 0 | 0 | 0 | 0 |
| 2 | PZ17_Adamiak_2 | 0 | 0 | 0 | 0 | 0 | 0 | 0 |
| 3 | PZ17_Adamiak_3 | 0 | 0 | 0 | 0 | 0 | 0 | 0 |
| 4 | PZ17_RNAComposer1_10 | 0 | 0 | 0 | 0 | 0 | 0 | 0 |
| 5 | PZ17_RNAComposer1_1 | 0 | 0 | 0 | 0 | 0 | 0 | 0 |
| 6 | PZ17_RNAComposer1_2 | 0 | 0 | 0 | 0 | 0 | 0 | 0 |
| 7 | PZ17_RNAComposer1_3 | 0 | 0 | 0 | 0 | 0 | 0 | 0 |
| 8 | PZ17_RNAComposer1_4 | 0 | 0 | 0 | 0 | 0 | 0 | 0 |
| 9 | PZ17_RNAComposer1_5 | 0 | 0 | 0 | 0 | 0 | 0 | 0 |
| 10 | PZ17_RNAComposer1_6 | 0 | 0 | 0 | 0 | 0 | 0 | 0 |
| 11 | PZ17_RNAComposer1_7 | 0 | 0 | 0 | 0 | 0 | 0 | 0 |
| 12 | PZ17_RNAComposer1_8 | 0 | 0 | 0 | 0 | 0 | 0 | 0 |
| 13 | PZ17_RNAComposer1_9 | 0 | 0 | 0 | 0 | 0 | 0 | 0 |
| 14 | PZ17_RNAComposer2_10 | 0 | 0 | 0 | 0 | 0 | 0 | 0 |
| 15 | PZ17_Das_10 | 0 | 0 | 0 | 0 | 0 | 0 | 0 |
| 16 | PZ17_RNAComposer2_9 | 0 | 0 | 0 | 0 | 0 | 0 | 0 |
| 17 | PZ17_Das_4 | 0 | 0 | 0 | 0 | 0 | 0 | 0 |
| 18 | PZ17_Das_5 | 0 | 0 | 0 | 0 | 0 | 0 | 0 |
| 19 | PZ17_Das_6 | 0 | 0 | 0 | 0 | 0 | 0 | 0 |
| 20 | PZ17_Das_7 | 0 | 0 | 0 | 0 | 0 | 0 | 0 |
| 21 | PZ17_Das_8 | 0 | 0 | 0 | 0 | 0 | 0 | 0 |
| 22 | PZ17_Das_9 | 0 | 0 | 0 | 0 | 0 | 0 | 0 |
| 23 | PZ17_DasExtraInfo_1 | 0 | 0 | 0 | 0 | 0 | 0 | 0 |
| 24 | PZ17_RNAComposer2_5 | 1 | 0 | 0 | 0 | 0 | 0 | 1 |
| 25 | PZ17_Das_2 | 1 | 0 | 0 | 0 | 0 | 0 | 1 |
| 26 | PZ17_Das_3 | 1 | 0 | 0 | 0 | 0 | 0 | 1 |
| 27 | PZ17_Das_1 | 0 | 1 | 2 | 0 | 0 | 0 | 3 |
| 28 | PZ17_DasExtraInfo_3 | 0 | 1 | 1 | 0 | 0 | 1 | 3 |
| 29 | PZ17_RNAComposer2_2 | 4 | 0 | 0 | 0 | 0 | 0 | 4 |
| 30 | PZ17_RNAComposer2_8 | 5 | 0 | 0 | 0 | 0 | 0 | 5 |
| 31 | PZ17_RNAComposer2_6 | 6 | 0 | 0 | 0 | 0 | 0 | 6 |
| 32 | PZ17_RNAComposer2_7 | 6 | 0 | 0 | 0 | 0 | 0 | 6 |
| 33 | PZ17_DasExtraInfo_2 | 0 | 1 | 3 | 0 | 1 | 1 | 6 |

| | | | | | | | | |
|---|---|---|---|---|---|---|---|---|
| 34 | PZ17_RNAComposer2_4 | 9 | 0 | 0 | 0 | 0 | 0 | 9 |
| 35 | PZ17_refstructure | 0 | 0 | 10 | 0 | 0 | 1 | 11 |
| 36 | PZ17_RNAComposer2_1 | 12 | 0 | 0 | 0 | 0 | 0 | 12 |
| 37 | PZ17_Chen_10 | 0 | 0 | 21 | 3 | 1 | 0 | 25 |
| 38 | PZ17_Chen_3 | 0 | 0 | 34 | 11 | 2 | 0 | 47 |
| 39 | PZ17_Ding_2 | 0 | 0 | 32 | 22 | 0 | 0 | 54 |
| 40 | PZ17_Ding_4 | 0 | 0 | 39 | 20 | 0 | 0 | 59 |
| 41 | PZ17_Bujnicki_4 | 0 | 0 | 50 | 8 | 1 | 0 | 59 |
| 42 | PZ17_Ding_8 | 0 | 1 | 39 | 20 | 0 | 0 | 60 |
| 43 | PZ17_Ding_1 | 0 | 0 | 43 | 19 | 0 | 0 | 62 |
| 44 | PZ17_Ding_5 | 0 | 0 | 39 | 24 | 0 | 0 | 63 |
| 45 | PZ17_Ding_3 | 0 | 0 | 45 | 19 | 0 | 0 | 64 |
| 46 | PZ17_Bujnicki_6 | 0 | 0 | 54 | 12 | 0 | 0 | 66 |
| 47 | PZ17_Ding_10 | 0 | 0 | 46 | 24 | 0 | 0 | 70 |
| 48 | PZ17_Ding_6 | 0 | 0 | 50 | 20 | 0 | 0 | 70 |
| 49 | PZ17_Bujnicki_10 | 0 | 0 | 56 | 16 | 1 | 0 | 73 |
| 50 | PZ17_Bujnicki_2 | 0 | 0 | 56 | 16 | 1 | 0 | 73 |
| 51 | PZ17_Bujnicki_3 | 0 | 0 | 53 | 20 | 1 | 0 | 74 |
| 52 | PZ17_Major_2 | 0 | 1 | 57 | 16 | 0 | 0 | 74 |
| 53 | PZ17_Bujnicki_8 | 0 | 13 | 63 | 1 | 0 | 0 | 77 |
| 54 | PZ17_Ding_9 | 0 | 0 | 55 | 24 | 0 | 0 | 79 |
| 55 | PZ17_Bujnicki_7 | 0 | 13 | 65 | 3 | 0 | 0 | 81 |
| 56 | PZ17_Ding_7 | 0 | 0 | 64 | 18 | 0 | 0 | 82 |
| 57 | PZ17_Major_3 | 0 | 0 | 61 | 18 | 3 | 0 | 82 |
| 58 | PZ17_Major_1 | 0 | 0 | 62 | 25 | 0 | 0 | 87 |
| 59 | PZ17_Major_4 | 0 | 0 | 70 | 17 | 0 | 0 | 87 |
| 60 | PZ17_Major_6 | 0 | 0 | 65 | 21 | 5 | 0 | 91 |
| 61 | PZ17_Major_8 | 0 | 0 | 65 | 26 | 4 | 0 | 95 |
| 62 | PZ17_Major_5 | 0 | 0 | 72 | 23 | 0 | 0 | 95 |
| 63 | PZ17_Chen_7 | 0 | 20 | 64 | 20 | 3 | 0 | 107 |
| 64 | PZ17_Major_7 | 0 | 0 | 82 | 24 | 4 | 0 | 110 |
| 65 | PZ17_Major_9 | 4 | 8 | 85 | 13 | 9 | 0 | 119 |
| 66 | PZ17_Dohkolyan_1 | 0 | 4 | 68 | 48 | 0 | 0 | 120 |
| 67 | PZ17_Chen_8 | 0 | 28 | 74 | 17 | 3 | 0 | 122 |
| 68 | PZ17_Dohkolyan_2 | 2 | 2 | 73 | 47 | 0 | 0 | 124 |
| 69 | PZ17_Dohkolyan_3 | 0 | 0 | 75 | 49 | 0 | 0 | 124 |
| 70 | PZ17_Bujnicki_1 | 0 | 15 | 105 | 13 | 4 | 0 | 137 |
| 71 | PZ17_Bujnicki_9 | 0 | 15 | 105 | 13 | 4 | 0 | 137 |
| 72 | PZ17_Major_10 | 4 | 7 | 102 | 27 | 5 | 0 | 145 |
| 73 | PZ17_Xiao_4 | 0 | 1 | 100 | 22 | 27 | 0 | 150 |
| 74 | PZ17_Xiao_2 | 0 | 1 | 92 | 28 | 36 | 0 | 157 |
| 75 | PZ17_SimRNA2_2 | 5 | 37 | 95 | 11 | 1 | 14 | 163 |
| 76 | PZ17_Xiao_3 | 0 | 1 | 93 | 32 | 40 | 0 | 166 |
| 77 | PZ17_Xiao_1 | 0 | 10 | 110 | 29 | 17 | 0 | 166 |
| 78 | PZ17_Xiao_7 | 0 | 1 | 100 | 37 | 31 | 0 | 169 |

| | | | | | | | | |
|---|---|---|---|---|---|---|---|---|
| 79 | PZ17_Xiao_10 | 0 | 1 | 93 | 33 | 45 | 0 | 172 |
| 80 | PZ17_SimRNA1_2 | 19 | 39 | 91 | 14 | 0 | 12 | 175 |
| 81 | PZ17_SimRNA1_8 | 19 | 28 | 105 | 14 | 1 | 9 | 176 |
| 82 | PZ17_SimRNA2_4 | 7 | 46 | 100 | 12 | 1 | 13 | 179 |
| 83 | PZ17_Bujnicki_5 | 0 | 20 | 138 | 16 | 6 | 0 | 180 |
| 84 | PZ17_SimRNA1_7 | 6 | 44 | 95 | 20 | 0 | 18 | 183 |
| 85 | PZ17_SimRNA2_8 | 12 | 35 | 95 | 20 | 1 | 21 | 184 |
| 86 | PZ17_SimRNA2_3 | 13 | 47 | 99 | 13 | 1 | 13 | 186 |
| 87 | PZ17_SimRNA1_1 | 33 | 35 | 100 | 9 | 1 | 17 | 195 |
| 88 | PZ17_SimRNA1_3 | 23 | 41 | 101 | 16 | 0 | 14 | 195 |
| 89 | PZ17_SimRNA1_6 | 18 | 45 | 99 | 17 | 1 | 16 | 196 |
| 90 | PZ17_Chen_9 | 28 | 29 | 116 | 4 | 0 | 20 | 197 |
| 91 | PZ17_SimRNA1_9 | 7 | 41 | 117 | 18 | 0 | 16 | 199 |
| 92 | PZ17_Xiao_5 | 4 | 20 | 129 | 27 | 20 | 0 | 200 |
| 93 | PZ17_SimRNA2_5 | 10 | 44 | 111 | 19 | 2 | 19 | 205 |
| 94 | PZ17_SimRNA2_1 | 25 | 44 | 112 | 15 | 1 | 10 | 207 |
| 95 | PZ17_Chen_2 | 29 | 36 | 120 | 1 | 0 | 21 | 207 |
| 96 | PZ17_SimRNA2_7 | 20 | 43 | 112 | 17 | 0 | 18 | 210 |
| 97 | PZ17_Xiao_8 | 4 | 13 | 132 | 31 | 38 | 0 | 218 |
| 98 | PZ17_Chen_1 | 46 | 28 | 133 | 0 | 0 | 12 | 219 |
| 99 | PZ17_Xiao_6 | 5 | 23 | 143 | 35 | 16 | 0 | 222 |
| 100 | PZ17_Xiao_9 | 0 | 12 | 135 | 40 | 36 | 0 | 223 |
| 101 | PZ17_SimRNA2_9 | 31 | 47 | 113 | 13 | 2 | 17 | 223 |
| 102 | PZ17_SimRNA2_6 | 15 | 46 | 133 | 22 | 2 | 12 | 230 |
| 103 | PZ17_SimRNA1_5 | 44 | 42 | 114 | 15 | 0 | 20 | 235 |
| 104 | PZ17_SimRNA1_4 | 38 | 46 | 134 | 17 | 0 | 19 | 254 |
| 105 | PZ17_Chen_6 | 82 | 33 | 119 | 2 | 0 | 21 | 257 |
| 106 | PZ17_Chen_4 | 58 | 39 | 139 | 3 | 0 | 20 | 259 |
| 107 | PZ17_Chen_5 | 100 | 26 | 125 | 3 | 1 | 24 | 279 |
| 108 | PZ17_RNAComposer2_3 | 5 | 142 | 144 | 33 | 0 | 0 | 324 |
| Average | | 7.05 | 12.19 | 58.26 | 12.73 | 3.51 | 3.69 | 97.43 |
| Standard deviation | | 16.11 | 20.71 | 48.69 | 12.56 | 9.30 | 7.12 | 87.64 |
| Median | | 0 | 0 | 61.5 | 13 | 0 | 0 | 80 |

**Table S19**. MAXIT report on structures in Puzzle 18.

| No. | 3D RNA model | Close con-tacts | Bond lengths | Bond angles | Plana-rity | Chira-lity | Poly-mer linkage | Total |
|---|---|---|---|---|---|---|---|---|
| 1 | PZ18_RNAComposer_3 | 0 | 0 | 0 | 0 | 0 | 0 | 0 |
| 2 | PZ18_RNAComposer_4 | 0 | 0 | 0 | 0 | 0 | 0 | 0 |
| 3 | PZ18_RNAComposer_5 | 0 | 0 | 0 | 0 | 0 | 0 | 0 |
| 4 | PZ18_FARFAR_10 | 0 | 0 | 0 | 0 | 0 | 0 | 0 |
| 5 | PZ18_FARFAR_1 | 0 | 0 | 0 | 0 | 0 | 0 | 0 |
| 6 | PZ18_FARFAR_3 | 0 | 0 | 0 | 0 | 0 | 0 | 0 |
| 7 | PZ18_FARFAR_4 | 0 | 0 | 0 | 0 | 0 | 0 | 0 |
| 8 | PZ18_FARFAR_7 | 0 | 0 | 0 | 0 | 0 | 0 | 0 |
| 9 | PZ18_FARFAR_8 | 0 | 0 | 0 | 0 | 0 | 0 | 0 |
| 10 | PZ18_RNAComposer_1 | 0 | 0 | 0 | 0 | 0 | 0 | 0 |
| 11 | PZ18_RNAComposer_2 | 0 | 0 | 0 | 0 | 0 | 0 | 0 |
| 12 | PZ18_refstructure | 0 | 0 | 1 | 0 | 0 | 0 | 1 |
| 13 | PZ18_FARFAR_5 | 0 | 0 | 1 | 0 | 0 | 0 | 1 |
| 14 | PZ18_FARFAR_6 | 1 | 0 | 0 | 0 | 0 | 0 | 1 |
| 15 | PZ18_FARFAR_9 | 0 | 1 | 0 | 0 | 0 | 0 | 1 |
| 16 | PZ18_YagoubAli_1 | 1 | 1 | 1 | 0 | 0 | 0 | 3 |
| 17 | PZ18_FARFAR_2 | 4 | 0 | 0 | 0 | 0 | 0 | 4 |
| 18 | PZ18_Lee_2 | 0 | 1 | 2 | 5 | 0 | 0 | 8 |
| 19 | PZ18_LeeServer_2 | 0 | 1 | 2 | 5 | 0 | 0 | 8 |
| 20 | PZ18_Lee_1 | 0 | 1 | 2 | 7 | 0 | 0 | 10 |
| 21 | PZ18_LeeServer_4 | 2 | 1 | 1 | 7 | 0 | 0 | 11 |
| 22 | PZ18_LeeServer_5 | 2 | 1 | 2 | 7 | 0 | 0 | 12 |
| 23 | PZ18_Lee_3 | 0 | 1 | 2 | 10 | 0 | 0 | 13 |
| 24 | PZ18_Lee_4 | 2 | 1 | 2 | 8 | 0 | 0 | 13 |
| 25 | PZ18_LeeServer_1 | 0 | 1 | 2 | 10 | 0 | 0 | 13 |
| 26 | PZ18_Lee_5 | 1 | 1 | 2 | 10 | 0 | 0 | 14 |
| 27 | PZ18_LeeServer_3 | 0 | 1 | 3 | 10 | 0 | 0 | 14 |
| 28 | PZ18_Das_3 | 0 | 4 | 11 | 0 | 0 | 0 | 15 |
| 29 | PZ18_Das_4 | 0 | 4 | 11 | 0 | 0 | 0 | 15 |
| 30 | PZ18_Das_5 | 0 | 5 | 12 | 0 | 0 | 0 | 17 |
| 31 | PZ18_Das_2 | 2 | 5 | 10 | 0 | 0 | 1 | 18 |
| 32 | PZ18_Das_1 | 1 | 5 | 14 | 0 | 0 | 0 | 20 |
| 33 | PZ18_Chen_5 | 0 | 13 | 34 | 5 | 0 | 0 | 52 |
| 34 | PZ18_Chen_1 | 0 | 2 | 44 | 13 | 0 | 0 | 59 |
| 35 | PZ18_Chen_2 | 0 | 3 | 45 | 13 | 0 | 0 | 61 |
| 36 | PZ18_Ding_2 | 0 | 0 | 40 | 26 | 0 | 0 | 66 |
| 37 | PZ18_Chen_4 | 0 | 6 | 52 | 11 | 0 | 0 | 69 |
| 38 | PZ18_Chen_3 | 0 | 3 | 51 | 16 | 1 | 0 | 71 |
| 39 | PZ18_Ding_4 | 0 | 0 | 50 | 24 | 0 | 0 | 74 |
| 40 | PZ18_Ding_1 | 0 | 0 | 47 | 30 | 0 | 0 | 77 |
| 41 | PZ18_Ding_5 | 0 | 0 | 60 | 25 | 0 | 0 | 85 |
| 42 | PZ18_Ding_3 | 0 | 1 | 57 | 30 | 0 | 0 | 88 |

| | | | | | | | | |
|---|---|---|---|---|---|---|---|---|
| 43 | PZ18_3dRNA_2 | 0 | 1 | 105 | 22 | 3 | 0 | 131 |
| 44 | PZ18_Dokholyan_2 | 0 | 7 | 85 | 52 | 0 | 0 | 144 |
| 45 | PZ18_Dokholyan_3 | 0 | 7 | 89 | 51 | 0 | 0 | 147 |
| 46 | PZ18_Dokholyan_1 | 0 | 6 | 97 | 51 | 0 | 0 | 154 |
| 47 | PZ18_3dRNA_4 | 0 | 1 | 121 | 18 | 19 | 0 | 159 |
| 48 | PZ18_3dRNA_5 | 0 | 1 | 129 | 24 | 16 | 0 | 170 |
| 49 | PZ18_3dRNA_3 | 0 | 11 | 145 | 25 | 9 | 0 | 190 |
| 50 | PZ18_SimRNA_1 | 12 | 47 | 95 | 19 | 1 | 16 | 190 |
| 51 | PZ18_SimRNA_3 | 8 | 51 | 107 | 13 | 0 | 19 | 198 |
| 52 | PZ18_3dRNA_1 | 1 | 10 | 154 | 26 | 13 | 0 | 204 |
| 53 | PZ18_SimRNA_2 | 5 | 47 | 128 | 13 | 1 | 21 | 215 |
| Average | | 0.79 | 4.75 | 34.26 | 11.06 | 1.19 | 1.08 | 53.13 |
| Standard deviation | | 2.13 | 11.20 | 46.11 | 13.79 | 3.93 | 4.38 | 67.99 |
| Median | | 0 | 1 | 3 | 7 | 0 | 0 | 14 |

**Table S20**. MAXIT report on structures in Puzzle 19.

| No. | 3D RNA model | Close con-tacts | Bond lengths | Bond angles | Plana-rity | Chira-lity | Poly-mer linkage | Total |
|---|---|---|---|---|---|---|---|---|
| 1 | PZ19_RNAComposer_1 | 0 | 0 | 0 | 0 | 0 | 0 | 0 |
| 2 | PZ19_RNAComposer_2 | 0 | 0 | 0 | 0 | 0 | 0 | 0 |
| 3 | PZ19_RNAComposer_3 | 0 | 0 | 0 | 0 | 0 | 0 | 0 |
| 4 | PZ19_RNAComposer_4 | 0 | 0 | 0 | 0 | 0 | 0 | 0 |
| 5 | PZ19_RNAComposer_5 | 0 | 0 | 0 | 0 | 0 | 0 | 0 |
| 6 | PZ19_Adamiak_1 | 0 | 0 | 0 | 0 | 0 | 0 | 0 |
| 7 | PZ19_Adamiak_2 | 0 | 0 | 0 | 0 | 0 | 0 | 0 |
| 8 | PZ19_Adamiak_3 | 0 | 0 | 0 | 0 | 0 | 0 | 0 |
| 9 | PZ19_Adamiak_4 | 0 | 0 | 0 | 0 | 0 | 0 | 0 |
| 10 | PZ19_Adamiak_5 | 0 | 0 | 0 | 0 | 0 | 0 | 0 |
| 11 | PZ19_FARFAR_1 | 1 | 0 | 0 | 0 | 0 | 0 | 1 |
| 12 | PZ19_FARFAR_2 | 1 | 0 | 0 | 0 | 0 | 0 | 1 |
| 13 | PZ19_FARFAR_7 | 1 | 0 | 0 | 0 | 0 | 0 | 1 |
| 14 | PZ19_FARFAR_9 | 1 | 0 | 0 | 0 | 0 | 0 | 1 |
| 15 | PZ19_FARFAR_10 | 1 | 1 | 0 | 0 | 0 | 0 | 2 |
| 16 | PZ19_FARFAR_3 | 1 | 0 | 1 | 0 | 0 | 0 | 2 |
| 17 | PZ19_FARFAR_4 | 1 | 0 | 1 | 0 | 0 | 0 | 2 |
| 18 | PZ19_FARFAR_5 | 1 | 0 | 1 | 0 | 0 | 0 | 2 |
| 19 | PZ19_FARFAR_6 | 1 | 1 | 0 | 0 | 0 | 0 | 2 |
| 20 | PZ19_Das_1 | 0 | 1 | 3 | 0 | 0 | 0 | 4 |
| 21 | PZ19_Das_5 | 0 | 2 | 2 | 0 | 0 | 0 | 4 |
| 22 | PZ19_FARFAR_8 | 1 | 1 | 2 | 0 | 0 | 0 | 4 |
| 23 | PZ19_Das_4 | 1 | 1 | 4 | 0 | 0 | 0 | 6 |
| 24 | PZ19_Das_3 | 0 | 3 | 4 | 0 | 0 | 0 | 7 |
| 25 | PZ19_Das_2 | 0 | 3 | 5 | 0 | 0 | 0 | 8 |
| 26 | PZ19_LeeServer_5 | 0 | 2 | 4 | 12 | 0 | 0 | 18 |

| | | | | | | | | |
|---|---|---|---|---|---|---|---|---|
| 27 | PZ19_refstructure | 0 | 7 | 16 | 0 | 0 | 0 | 23 |
| 28 | PZ19_LeeServer_4 | 0 | 2 | 5 | 17 | 0 | 0 | 24 |
| 29 | PZ19_Chen_2 | 0 | 2 | 37 | 11 | 0 | 0 | 50 |
| 30 | PZ19_Chen_4 | 0 | 7 | 43 | 5 | 0 | 0 | 55 |
| 31 | PZ19_Ding_4 | 0 | 1 | 41 | 19 | 0 | 0 | 61 |
| 32 | PZ19_Chen_3 | 0 | 1 | 44 | 16 | 0 | 0 | 61 |
| 33 | PZ19_Ding_2 | 0 | 0 | 38 | 26 | 0 | 0 | 64 |
| 34 | PZ19_LeeServer_1 | 0 | 2 | 4 | 14 | 50 | 0 | 70 |
| 35 | PZ19_Ding_5 | 0 | 1 | 45 | 25 | 0 | 0 | 71 |
| 36 | PZ19_LeeServer_2 | 0 | 2 | 5 | 13 | 51 | 0 | 71 |
| 37 | PZ19_LeeServer_3 | 0 | 2 | 5 | 15 | 49 | 0 | 71 |
| 38 | PZ19_Ding_3 | 0 | 0 | 49 | 26 | 0 | 0 | 75 |
| 39 | PZ19_Ding_1 | 0 | 1 | 52 | 26 | 0 | 0 | 79 |
| 40 | PZ19_Bujnicki_1 | 0 | 8 | 60 | 9 | 2 | 0 | 79 |
| 41 | PZ19_Bujnicki_2 | 0 | 6 | 84 | 9 | 5 | 0 | 104 |
| 42 | PZ19_Dokholyan_1 | 0 | 0 | 61 | 46 | 0 | 0 | 107 |
| 43 | PZ19_Bujnicki_5 | 0 | 10 | 80 | 9 | 11 | 0 | 110 |
| 44 | PZ19_Chen_1 | 0 | 18 | 89 | 20 | 1 | 0 | 128 |
| 45 | PZ19_Bujnicki_3 | 0 | 6 | 102 | 19 | 4 | 0 | 131 |
| 46 | PZ19_SimRNA_1 | 14 | 38 | 91 | 12 | 1 | 11 | 167 |
| 47 | PZ19_Chen_5 | 0 | 30 | 110 | 23 | 4 | 0 | 167 |
| 48 | PZ19_3dRNA_5 | 1 | 13 | 117 | 27 | 11 | 0 | 169 |
| 49 | PZ19_3dRNA_2 | 1 | 13 | 118 | 33 | 11 | 0 | 176 |
| 50 | PZ19_Bujnicki_4 | 0 | 12 | 132 | 22 | 10 | 0 | 176 |
| 51 | PZ19_3dRNA_4 | 1 | 12 | 121 | 29 | 15 | 0 | 178 |
| 52 | PZ19_3dRNA_3 | 1 | 8 | 119 | 36 | 15 | 0 | 179 |
| 53 | PZ19_3dRNA_1 | 1 | 13 | 128 | 33 | 12 | 0 | 187 |
| 54 | PZ19_SimRNA_3 | 10 | 50 | 110 | 11 | 2 | 14 | 197 |
| 55 | PZ19_SimRNA_2 | 8 | 43 | 117 | 17 | 0 | 17 | 202 |
| Average | | 0.87 | 5.87 | 37.27 | 10.55 | 4.62 | 0.76 | 59.95 |
| Standard deviation | | 2.49 | 10.86 | 46.13 | 12.21 | 11.75 | 3.26 | 68.27 |
| Median | | 0 | 1 | 5 | 9 | 0 | 0 | 24 |

**Table S21**. MAXIT report on structures in Puzzle 20.

| No. | 3D RNA model | Close con-tacts | Bond lengths | Bond angles | Plana-rity | Chira-lity | Poly-mer linkage | Total |
|---|---|---|---|---|---|---|---|---|
| 1 | PZ20_RNAComposer_5 | 0 | 0 | 0 | 0 | 0 | 0 | 0 |
| 2 | PZ20_FARFAR_8 | 0 | 0 | 0 | 0 | 0 | 0 | 0 |
| 3 | PZ20_Adamiak_1 | 0 | 0 | 0 | 0 | 0 | 0 | 0 |
| 4 | PZ20_FARFAR_9 | 0 | 0 | 0 | 0 | 0 | 0 | 0 |
| 5 | PZ20_Adamiak_2 | 0 | 0 | 0 | 0 | 0 | 0 | 0 |
| 6 | PZ20_Adamiak_3 | 0 | 0 | 0 | 0 | 0 | 0 | 0 |
| 7 | PZ20_Adamiak_4 | 0 | 0 | 0 | 0 | 0 | 0 | 0 |
| 8 | PZ20_Adamiak_5 | 0 | 0 | 0 | 0 | 0 | 0 | 0 |

| | | | | | | | | |
|---|---|---|---|---|---|---|---|---|
| 9 | PZ20_FARFAR_1 | 0 | 0 | 0 | 0 | 0 | 0 | 0 |
| 10 | PZ20_refstructure | 0 | 0 | 0 | 0 | 0 | 0 | 0 |
| 11 | PZ20_FARFAR_5 | 0 | 0 | 0 | 0 | 0 | 0 | 0 |
| 12 | PZ20_RNAComposer_3 | 0 | 0 | 0 | 0 | 0 | 0 | 0 |
| 13 | PZ20_FARFAR_6 | 0 | 0 | 0 | 0 | 0 | 0 | 0 |
| 14 | PZ20_RNAComposer_4 | 0 | 0 | 0 | 0 | 0 | 0 | 0 |
| 15 | PZ20_FARFAR_2 | 1 | 0 | 0 | 0 | 0 | 0 | 1 |
| 16 | PZ20_FARFAR_3 | 0 | 0 | 1 | 0 | 0 | 0 | 1 |
| 17 | PZ20_FARFAR_4 | 0 | 0 | 1 | 0 | 0 | 0 | 1 |
| 18 | PZ20_RNAComposer_2 | 1 | 0 | 0 | 0 | 0 | 0 | 1 |
| 19 | PZ20_FARFAR_10 | 0 | 0 | 2 | 0 | 0 | 0 | 2 |
| 20 | PZ20_FARFAR_7 | 0 | 0 | 1 | 0 | 1 | 0 | 2 |
| 21 | PZ20_RNAComposer_1 | 4 | 0 | 0 | 0 | 0 | 0 | 4 |
| 22 | PZ20_Das_2 | 0 | 1 | 4 | 0 | 0 | 0 | 5 |
| 23 | PZ20_Das_4 | 3 | 1 | 2 | 0 | 0 | 0 | 6 |
| 24 | PZ20_Das_5 | 2 | 1 | 2 | 0 | 0 | 1 | 6 |
| 25 | PZ20_Das_1 | 2 | 1 | 4 | 0 | 0 | 0 | 7 |
| 26 | PZ20_Das_3 | 1 | 3 | 3 | 0 | 0 | 1 | 8 |
| 27 | PZ20_Bujnicki_5 | 0 | 9 | 80 | 17 | 3 | 0 | 109 |
| 28 | PZ20_Bujnicki_1 | 3 | 13 | 115 | 14 | 1 | 2 | 148 |
| 29 | PZ20_Xiao3_5 | 1 | 1 | 115 | 34 | 34 | 0 | 185 |
| 30 | PZ20_SimRNA_5 | 12 | 45 | 122 | 19 | 0 | 10 | 208 |
| 31 | PZ20_SimRNA_2 | 25 | 48 | 107 | 17 | 1 | 21 | 219 |
| 32 | PZ20_Xiao3_2 | 1 | 9 | 133 | 39 | 37 | 0 | 219 |
| 33 | PZ20_Xiao3_4 | 1 | 9 | 120 | 43 | 50 | 0 | 223 |
| 34 | PZ20_SimRNA_4 | 5 | 58 | 124 | 16 | 0 | 22 | 225 |
| 35 | PZ20_SimRNA_1 | 18 | 54 | 119 | 17 | 1 | 17 | 226 |
| 36 | PZ20_SimRNA_3 | 15 | 52 | 133 | 22 | 2 | 14 | 238 |
| 37 | PZ20_Xiao3_1 | 1 | 19 | 142 | 42 | 53 | 0 | 257 |
| 38 | PZ20_Xiao3_3 | 1 | 28 | 165 | 38 | 25 | 0 | 257 |
| 39 | PZ20_Bujnicki_2 | 3 | 33 | 207 | 15 | 3 | 2 | 263 |
| 40 | PZ20_Bujnicki_4 | 4 | 39 | 206 | 14 | 1 | 3 | 267 |
| 41 | PZ20_Bujnicki_3 | 5 | 56 | 205 | 12 | 2 | 3 | 283 |
| Average | | 2.66 | 11.71 | 51.54 | 8.76 | 5.22 | 2.34 | 82.22 |
| Standard deviation | | 5.37 | 19.37 | 71.48 | 13.53 | 13.58 | 5.73 | 110.79 |
| Median | | 1 | 0 | 2 | 0 | 0 | 0 | 4 |

**Table S22**. MAXIT report on structures in Puzzle 21.

| **No.** | **3D RNA model** | **Close con-tacts** | **Bond lengths** | **Bond angles** | **Plana-rity** | **Chira-lity** | **Poly-mer linkage** | **Total** |
|---|---|---|---|---|---|---|---|---|
| 1 | PZ21_FARFAR_10 | 0 | 0 | 0 | 0 | 0 | 0 | 0 |
| 2 | PZ21_FARFAR_1 | 0 | 0 | 0 | 0 | 0 | 0 | 0 |
| 3 | PZ21_FARFAR_3 | 0 | 0 | 0 | 0 | 0 | 0 | 0 |
| 4 | PZ21_FARFAR_4 | 0 | 0 | 0 | 0 | 0 | 0 | 0 |
| 5 | PZ21_Adamiak_1 | 0 | 0 | 0 | 0 | 0 | 0 | 0 |

| | | | | | | | | |
|---|---|---|---|---|---|---|---|---|
| 6 | PZ21_Das_1 | 0 | 0 | 0 | 0 | 0 | 0 | 0 |
| 7 | PZ21_FARFAR_5 | 0 | 0 | 0 | 0 | 0 | 0 | 0 |
| 8 | PZ21_Adamiak_2 | 0 | 0 | 0 | 0 | 0 | 0 | 0 |
| 9 | PZ21_Das_2 | 0 | 0 | 0 | 0 | 0 | 0 | 0 |
| 10 | PZ21_FARFAR_6 | 0 | 0 | 0 | 0 | 0 | 0 | 0 |
| 11 | PZ21_Adamiak_3 | 0 | 0 | 0 | 0 | 0 | 0 | 0 |
| 12 | PZ21_FARFAR_7 | 0 | 0 | 0 | 0 | 0 | 0 | 0 |
| 13 | PZ21_Adamiak_4 | 0 | 0 | 0 | 0 | 0 | 0 | 0 |
| 14 | PZ21_Das_4 | 0 | 0 | 0 | 0 | 0 | 0 | 0 |
| 15 | PZ21_FARFAR_8 | 0 | 0 | 0 | 0 | 0 | 0 | 0 |
| 16 | PZ21_Adamiak_5 | 0 | 0 | 0 | 0 | 0 | 0 | 0 |
| 17 | PZ21_FARFAR_9 | 0 | 0 | 0 | 0 | 0 | 0 | 0 |
| 18 | PZ21_DasLORES_1 | 0 | 0 | 0 | 0 | 0 | 0 | 0 |
| 19 | PZ21_DasLORES_4 | 0 | 0 | 0 | 0 | 0 | 0 | 0 |
| 20 | PZ21_RNAComposer_1 | 0 | 0 | 0 | 0 | 0 | 0 | 0 |
| 21 | PZ21_RNAComposer_2 | 0 | 0 | 0 | 0 | 0 | 0 | 0 |
| 22 | PZ21_RNAComposer_4 | 0 | 0 | 0 | 0 | 0 | 0 | 0 |
| 23 | PZ21_RNAComposer_5 | 0 | 0 | 0 | 0 | 0 | 0 | 0 |
| 24 | PZ21_DasLORES_2 | 1 | 0 | 0 | 0 | 0 | 0 | 1 |
| 25 | PZ21_DasLORES_5 | 1 | 0 | 0 | 0 | 0 | 0 | 1 |
| 26 | PZ21_RNAComposer_3 | 1 | 0 | 0 | 0 | 0 | 0 | 1 |
| 27 | PZ21_FARFAR_2 | 0 | 1 | 1 | 0 | 0 | 0 | 2 |
| 28 | PZ21_Das_3 | 0 | 0 | 3 | 0 | 0 | 0 | 3 |
| 29 | PZ21_DasLORES_3 | 0 | 1 | 2 | 0 | 0 | 0 | 3 |
| 30 | PZ21_Das_5 | 0 | 0 | 5 | 0 | 0 | 0 | 5 |
| 31 | PZ21_refstructure | 2 | 15 | 9 | 0 | 0 | 1 | 27 |
| 32 | PZ21_Bujnicki_4 | 0 | 7 | 40 | 5 | 4 | 0 | 56 |
| 33 | PZ21_Sanbonmatsu_1 | 6 | 14 | 25 | 0 | 12 | 0 | 57 |
| 34 | PZ21_Sanbonmatsu_2 | 6 | 14 | 25 | 0 | 12 | 0 | 57 |
| 35 | PZ21_Bujnicki_2 | 2 | 13 | 34 | 3 | 2 | 5 | 59 |
| 36 | PZ21_Bujnicki_1 | 0 | 8 | 52 | 4 | 4 | 0 | 68 |
| 37 | PZ21_Bujnicki_3 | 0 | 8 | 50 | 4 | 6 | 0 | 68 |
| 38 | PZ21_Sanbonmatsu_3 | 23 | 14 | 29 | 0 | 8 | 0 | 74 |
| 39 | PZ21_Sanbonmatsu_4 | 23 | 14 | 29 | 0 | 8 | 0 | 74 |
| 40 | PZ21_Bujnicki_5 | 0 | 6 | 59 | 6 | 4 | 0 | 75 |
| 41 | PZ21_3dRNA_5 | 0 | 0 | 75 | 12 | 22 | 0 | 109 |
| 42 | PZ21_3dRNA_3 | 0 | 0 | 69 | 22 | 21 | 0 | 112 |
| 43 | PZ21_SimRNA_2 | 2 | 25 | 66 | 11 | 2 | 10 | 116 |
| 44 | PZ21_3dRNA_4 | 0 | 1 | 79 | 11 | 26 | 0 | 117 |
| 45 | PZ21_SimRNA_4 | 8 | 27 | 76 | 12 | 0 | 10 | 133 |
| 46 | PZ21_SimRNA_1 | 7 | 30 | 79 | 9 | 0 | 13 | 138 |
| 47 | PZ21_3dRNA_1 | 0 | 19 | 95 | 23 | 13 | 0 | 150 |
| 48 | PZ21_SimRNA_3 | 12 | 36 | 89 | 15 | 1 | 13 | 166 |
| 49 | PZ21_SimRNA_5 | 12 | 36 | 89 | 15 | 1 | 13 | 166 |
| 50 | PZ21_3dRNA_2 | 0 | 21 | 107 | 25 | 22 | 0 | 175 |

| | | | | | | | | |
|---|---|---|---|---|---|---|---|---|
| 51 | PZ21_ChenHighLig_2 | 9 | 23 | 153 | 2 | 0 | 5 | 192 |
| 52 | PZ21_ChenLowLig_2 | 9 | 23 | 154 | 2 | 0 | 5 | 193 |
| 53 | PZ21_ChenLowLig_5 | 33 | 25 | 123 | 1 | 0 | 12 | 194 |
| 54 | PZ21_ChenHighLig_5 | 33 | 25 | 123 | 1 | 0 | 12 | 194 |
| 55 | PZ21_ChenLowLig_3 | 30 | 32 | 135 | 2 | 0 | 9 | 208 |
| 56 | PZ21_ChenHighLig_3 | 30 | 32 | 135 | 2 | 0 | 9 | 208 |
| 57 | PZ21_ChenHighLig_1 | 36 | 31 | 130 | 1 | 0 | 15 | 213 |
| 58 | PZ21_ChenLowLig_1 | 36 | 31 | 131 | 1 | 0 | 15 | 214 |
| 59 | PZ21_ChenHighLig_4 | 49 | 37 | 134 | 4 | 0 | 13 | 237 |
| 60 | PZ21_ChenLowLig_4 | 49 | 37 | 135 | 4 | 0 | 13 | 238 |
| Average | | 7.00 | 10.10 | 42.33 | 3.28 | 2.80 | 2.88 | 68.40 |
| Standard deviation | | 13.11 | 12.98 | 52.13 | 6.11 | 6.23 | 5.08 | 81.87 |
| Median | | 0 | 0.5 | 7 | 0 | 0 | 0 | 16 |

**Table S23**. MAXIT report on structures in Puzzle 24.

| No. | 3D RNA model | Close con-tacts | Bond lengths | Bond angles | Plana-rity | Chira-lity | Poly-mer linkage | Total |
|---|---|---|---|---|---|---|---|---|
| 1 | PZ24_RNAComposer_1 | 0 | 0 | 0 | 0 | 0 | 0 | 0 |
| 2 | PZ24_RNAComposer_2 | 0 | 0 | 0 | 0 | 0 | 0 | 0 |
| 3 | PZ24_RNAComposer_3 | 0 | 0 | 0 | 0 | 0 | 0 | 0 |
| 4 | PZ24_RNAComposer_4 | 0 | 0 | 0 | 0 | 0 | 0 | 0 |
| 5 | PZ24_RNAComposer_5 | 0 | 0 | 0 | 0 | 0 | 0 | 0 |
| 6 | PZ24_Adamiak_1 | 0 | 0 | 0 | 0 | 0 | 0 | 0 |
| 7 | PZ24_Adamiak_2 | 0 | 0 | 0 | 0 | 0 | 0 | 0 |
| 8 | PZ24_Adamiak_3 | 0 | 0 | 0 | 0 | 0 | 0 | 0 |
| 9 | PZ24_Das_1 | 0 | 0 | 0 | 0 | 0 | 0 | 0 |
| 10 | PZ24_Adamiak_4 | 0 | 0 | 0 | 0 | 0 | 0 | 0 |
| 11 | PZ24_Das_2 | 0 | 0 | 0 | 0 | 0 | 0 | 0 |
| 12 | PZ24_Adamiak_5 | 0 | 0 | 0 | 0 | 0 | 0 | 0 |
| 13 | PZ24_Das_3 | 0 | 0 | 0 | 0 | 0 | 0 | 0 |
| 14 | PZ24_Das_4 | 0 | 0 | 0 | 0 | 0 | 0 | 0 |
| 15 | PZ24_FARFAR2_5 | 0 | 0 | 0 | 0 | 0 | 0 | 0 |
| 16 | PZ24_Das_5 | 0 | 0 | 0 | 0 | 0 | 0 | 0 |
| 17 | PZ24_FARFAR2_7 | 0 | 0 | 0 | 0 | 0 | 0 | 0 |
| 18 | PZ24_Das_8 | 0 | 0 | 0 | 0 | 0 | 0 | 0 |
| 19 | PZ24_FARFAR2_9 | 0 | 0 | 0 | 0 | 0 | 0 | 0 |
| 20 | PZ24_DasTFN_10 | 0 | 0 | 0 | 0 | 0 | 0 | 0 |
| 21 | PZ24_DasTFN_4 | 0 | 0 | 0 | 0 | 0 | 0 | 0 |
| 22 | PZ24_FARFAR2_10 | 1 | 0 | 0 | 0 | 0 | 0 | 1 |
| 23 | PZ24_Das_10 | 0 | 0 | 0 | 1 | 0 | 0 | 1 |
| 24 | PZ24_FARFAR2_1 | 0 | 0 | 1 | 0 | 0 | 0 | 1 |
| 25 | PZ24_FARFAR2_2 | 0 | 0 | 0 | 1 | 0 | 0 | 1 |
| 26 | PZ24_FARFAR2_6 | 0 | 1 | 0 | 0 | 0 | 0 | 1 |
| 27 | PZ24_FARFAR2_8 | 0 | 0 | 1 | 0 | 0 | 0 | 1 |
| 28 | PZ24_Das_9 | 0 | 0 | 0 | 1 | 0 | 0 | 1 |

| | | | | | | | | |
|---|---|---|---|---|---|---|---|---|
| 29 | PZ24_DasTFN_3 | 0 | 0 | 1 | 0 | 0 | 0 | 1 |
| 30 | PZ24_DasTFN_7 | 0 | 0 | 1 | 0 | 0 | 0 | 1 |
| 31 | PZ24_DasTFN_9 | 0 | 1 | 0 | 0 | 0 | 0 | 1 |
| 32 | PZ24_FARFAR2_3 | 0 | 0 | 0 | 2 | 0 | 0 | 2 |
| 33 | PZ24_Das_6 | 0 | 0 | 0 | 2 | 0 | 0 | 2 |
| 34 | PZ24_Das_7 | 0 | 0 | 0 | 2 | 0 | 0 | 2 |
| 35 | PZ24_FARFAR2_4 | 0 | 0 | 1 | 2 | 0 | 0 | 3 |
| 36 | PZ24_DasTFN_2 | 1 | 0 | 1 | 1 | 0 | 0 | 3 |
| 37 | PZ24_DasTFN_6 | 0 | 1 | 1 | 1 | 0 | 0 | 3 |
| 38 | PZ24_DasTFN_8 | 0 | 0 | 3 | 0 | 0 | 0 | 3 |
| 39 | PZ24_DasTFN_5 | 0 | 0 | 2 | 2 | 0 | 0 | 4 |
| 40 | PZ24_DasTFN_1 | 2 | 0 | 2 | 2 | 0 | 0 | 6 |
| 41 | PZ24_refstructure | 6 | 0 | 0 | 0 | 0 | 1 | 7 |
| 42 | PZ24_Vfold3D_4 | 0 | 2 | 50 | 17 | 2 | 0 | 71 |
| 43 | PZ24_Vfold3D_2 | 0 | 0 | 67 | 21 | 0 | 0 | 88 |
| 44 | PZ24_Vfold3D_5 | 0 | 4 | 66 | 20 | 0 | 0 | 90 |
| 45 | PZ24_Vfold3D_1 | 0 | 4 | 69 | 19 | 0 | 0 | 92 |
| 46 | PZ24_Vfold3D_3 | 0 | 10 | 67 | 24 | 0 | 0 | 101 |
| 47 | PZ24_3dRNA_4 | 0 | 0 | 106 | 17 | 0 | 0 | 123 |
| 48 | PZ24_3dRNA_2 | 0 | 0 | 109 | 19 | 0 | 0 | 128 |
| 49 | PZ24_3dRNA_3 | 0 | 0 | 115 | 18 | 1 | 0 | 134 |
| 50 | PZ24_3dRNA_1 | 0 | 0 | 119 | 18 | 0 | 0 | 137 |
| 51 | PZ24_3dRNA_5 | 0 | 0 | 113 | 27 | 0 | 0 | 140 |
| 52 | PZ24_Bujnicki_4 | 0 | 11 | 124 | 15 | 6 | 0 | 156 |
| 53 | PZ24_Bujnicki_2 | 0 | 12 | 124 | 17 | 6 | 0 | 159 |
| 54 | PZ24_Bujnicki_1 | 0 | 14 | 120 | 15 | 16 | 0 | 165 |
| 55 | PZ24_Ding_9 | 0 | 3 | 92 | 74 | 0 | 0 | 169 |
| 56 | PZ24_Bujnicki_3 | 0 | 12 | 130 | 25 | 9 | 0 | 176 |
| 57 | PZ24_Ding_7 | 0 | 0 | 111 | 78 | 0 | 0 | 189 |
| 58 | PZ24_Ding_4 | 0 | 1 | 112 | 78 | 0 | 0 | 191 |
| 59 | PZ24_Ding_6 | 0 | 1 | 120 | 71 | 0 | 0 | 192 |
| 60 | PZ24_Ding_10 | 0 | 0 | 119 | 76 | 0 | 0 | 195 |
| 61 | PZ24_Ding_5 | 0 | 1 | 121 | 74 | 0 | 0 | 196 |
| 62 | PZ24_Ding_1 | 0 | 2 | 118 | 77 | 0 | 0 | 197 |
| 63 | PZ24_iFoldRNA_4 | 0 | 1 | 151 | 48 | 0 | 0 | 200 |
| 64 | PZ24_Ding_8 | 0 | 3 | 115 | 86 | 0 | 0 | 204 |
| 65 | PZ24_Ding_2 | 0 | 2 | 124 | 79 | 0 | 0 | 205 |
| 66 | PZ24_Ding_3 | 0 | 0 | 121 | 84 | 0 | 0 | 205 |
| 67 | PZ24_iFoldRNA_2 | 0 | 1 | 159 | 49 | 0 | 0 | 209 |
| 68 | PZ24_iFoldRNA_1 | 0 | 1 | 168 | 48 | 0 | 0 | 217 |
| 69 | PZ24_iFoldRNA_5 | 0 | 1 | 158 | 59 | 0 | 0 | 218 |
| 70 | PZ24_iFoldRNA_3 | 0 | 1 | 172 | 45 | 0 | 0 | 218 |
| 71 | PZ24_Bujnicki_5 | 0 | 15 | 183 | 25 | 7 | 0 | 230 |
| 72 | PZ24_Kollmann_7 | 22 | 41 | 151 | 24 | 4 | 9 | 251 |
| 73 | PZ24_Kollmann_6 | 33 | 46 | 158 | 27 | 1 | 16 | 281 |

| | | | | | | | | |
|---|---|---|---|---|---|---|---|---|
| 74 | PZ24_Kollmann_2 | 69 | 44 | 141 | 16 | 2 | 10 | 282 |
| 75 | PZ24_Kollmann_9 | 26 | 55 | 183 | 26 | 3 | 12 | 305 |
| 76 | PZ24_Kollmann_4 | 67 | 51 | 152 | 24 | 2 | 15 | 311 |
| 77 | PZ24_Kollmann_10 | 50 | 47 | 174 | 29 | 1 | 12 | 313 |
| 78 | PZ24_Kollmann_8 | 62 | 46 | 176 | 26 | 0 | 9 | 319 |
| 79 | PZ24_Kollmann_5 | 46 | 62 | 173 | 27 | 1 | 15 | 324 |
| 80 | PZ24_SimRNA_2 | 23 | 72 | 185 | 25 | 0 | 24 | 329 |
| 81 | PZ24_Kollmann_3 | 47 | 58 | 183 | 29 | 1 | 13 | 331 |
| 82 | PZ24_Kollmann_1 | 67 | 61 | 188 | 31 | 4 | 19 | 370 |
| 83 | PZ24_SimRNA_5 | 27 | 84 | 200 | 30 | 3 | 33 | 377 |
| 84 | PZ24_SimRNA_3 | 25 | 95 | 222 | 32 | 0 | 24 | 398 |
| 85 | PZ24_SimRNA_4 | 12 | 98 | 247 | 27 | 1 | 32 | 417 |
| 86 | PZ24_SimRNA_1 | 45 | 83 | 229 | 39 | 2 | 26 | 424 |
| 87 | PZ24_VfoldLA_4 | 61 | 70 | 284 | 0 | 0 | 71 | 486 |
| 88 | PZ24_VfoldLA_1 | 124 | 51 | 252 | 7 | 0 | 61 | 495 |
| 89 | PZ24_VfoldLA_3 | 119 | 58 | 262 | 8 | 0 | 65 | 512 |
| 90 | PZ24_VfoldLA_5 | 98 | 77 | 277 | 7 | 1 | 61 | 521 |
| 91 | PZ24_VfoldLA_2 | 135 | 69 | 271 | 7 | 0 | 57 | 539 |
| Average | | 12.84 | 15.09 | 84.01 | 19.57 | 0.80 | 6.43 | 138.74 |
| Standard deviation | | 29.22 | 27.12 | 87.30 | 24.99 | 2.29 | 15.61 | 155.32 |
| Median | | 0 | 0 | 69 | 8 | 0 | 0 | 101 |

**Table S24**. Total number of errors of different types by cluster and the average number over all types of errors (ATN) per model in every cluster.

| No. | Clus-ter | Total number of models | Close con-tacts | Bond lengths | Bond angles | Plana-rity | Chira-lity | Poly-mer lin-kage | Total | ATN |
|---|---|---|---|---|---|---|---|---|---|---|
| 1 | H1 | 64 | 0 | 0 | 0 | 0 | 0 | 0 | 0 | 0.00 |
| 2 | H2 | 131 | 118 | 2,426 | 12,444 | 2,231 | 584 | 91 | 17,894 | 136.60 |
| 3 | H3 | 145 | 1,854 | 2,817 | 12,188 | 1,712 | 110 | 732 | 19,413 | 133.88 |
| 4 | H4 | 188 | 414 | 254 | 642 | 27 | 8 | 93 | 1,446 | 7.69 |
| 5 | H5 | 124 | 14 | 44 | 8,710 | 4,241 | 2 | 1 | 13,012 | 104.94 |
| 6 | H6 | 52 | 5 | 107 | 6,036 | 3,313 | 9 | 1 | 9,471 | 182.13 |
| 7 | H7 | 10 | 489 | 511 | 1,679 | 259 | 19 | 130 | 3,087 | 308.70 |
| 8 | H8 | 5 | 3 | 5 | 10 | 40 | 0 | 0 | 58 | 11.60 |
| 9 | H9 | 32 | 11 | 1,112 | 7,364 | 1,869 | 107 | 28 | 10,491 | 327.84 |
| 10 | H10 | 1 | 26 | 15 | 24 | 6 | 0 | 1 | 72 | 72.00 |
| 11 | H11 | 4 | 58 | 56 | 108 | 0 | 40 | 0 | 262 | 65.50 |
| 12 | H12 | 3 | 6 | 102 | 99 | 0 | 2 | 9 | 218 | 72.67 |
| 13 | H13 | 3 | 0 | 5 | 326 | 262 | 0 | 0 | 593 | 197.67 |
| 14 | H14 | 1 | 271 | 39 | 210 | 0 | 0 | 0 | 520 | 520.00 |
| 15 | H15 | 33 | 39 | 952 | 5,815 | 1,215 | 551 | 9 | 8,581 | 260.03 |
| 16 | H16 | 1 | 1 | 1 | 1 | 0 | 0 | 0 | 3 | 3.00 |
| 17 | W1 | 30 | 26 | 328 | 4,160 | 816 | 443 | 14 | 5,787 | 192.90 |
| 18 | W2 | 70 | 18 | 22 | 45 | 5 | 1 | 0 | 91 | 1.30 |
| 19 | W3 | 5 | 0 | 5 | 808 | 249 | 0 | 0 | 1,062 | 212.40 |

| | | | | | | | | | | |
|---|---|---|---|---|---|---|---|---|---|---|
| 20 | W4 | 10 | 4 | 15 | 33 | 110 | 150 | 0 | 312 | 31.20 |
| 21 | W5 | 49 | 54 | 142 | 144 | 33 | 0 | 0 | 373 | 7.61 |
| 22 | W6 | 59 | 986 | 2,899 | 7,483 | 1,056 | 56 | 998 | 13,478 | 228.44 |
| 23 | W7 | 10 | 537 | 345 | 1,665 | 130 | 3 | 315 | 2,995 | 299.50 |

BARNABA reports

BBRMSD - RMSD is calculated using backbone atoms only.
ERMSD - eRMSD measures the distance between structures by considering only the relative positions and orientations of nucleobases.

**Table S25**. BARNABA report on structures in Puzzle 01.

| No. | 3D RNA model | BBRMSD [nm] | ERMSD [nm] | BBRMSD [Å] | ERMSD [Å] |
|---|---|---|---|---|---|
| 1 | PZ01_Das_3 | 0.35 | 0.55 | 3.50 | 5.53 |
| 2 | PZ01_Santalucia_1 | 0.62 | 0.71 | 6.19 | 7.08 |
| 3 | PZ01_Das_1 | 0.39 | 0.73 | 3.93 | 7.30 |
| 4 | PZ01_Das_4 | 0.37 | 0.82 | 3.73 | 8.18 |
| 5 | PZ01_Das_2 | 0.45 | 0.91 | 4.55 | 9.05 |
| 6 | PZ01_Bujnicki_1 | 0.59 | 0.93 | 5.87 | 9.27 |
| 7 | PZ01_Major_1 | 0.43 | 0.94 | 4.31 | 9.44 |
| 8 | PZ01_Chen_1 | 0.43 | 0.95 | 4.35 | 9.47 |
| 9 | PZ01_Bujnicki_3 | 0.54 | 0.97 | 5.42 | 9.66 |
| 10 | PZ01_Das_5 | 0.43 | 0.98 | 4.35 | 9.80 |
| 11 | PZ01_Dokholyan_1 | 0.73 | 1.02 | 7.34 | 10.19 |
| 12 | PZ01_Bujnicki_2 | 0.64 | 1.02 | 6.42 | 10.23 |
| 13 | PZ01_Bujnicki_5 | 0.53 | 1.38 | 5.25 | 13.82 |
| 14 | PZ01_Bujnicki_4 | 0.51 | 1.44 | 5.09 | 14.40 |
| | Average | 0.50 | 0.95 | 5.02 | 9.53 |
| | Standard deviation | 0.35 | 0.55 | 3.50 | 5.53 |
| | Median | 0.73 | 1.44 | 7.34 | 14.40 |

**Table S26**. BARNABA report on structures in Puzzle 02.

| No. | 3D RNA model | BBRMSD [nm] | ERMSD [nm] | BBRMSD [Å] | ERMSD [Å] |
|---|---|---|---|---|---|
| 1 | PZ02_Santalucia_1 | 0.38 | 0.83 | 3.79 | 8.25 |
| 2 | PZ02_Das_1 | 0.23 | 0.86 | 2.28 | 8.57 |
| 3 | PZ02_Bujnicki_1 | 0.23 | 0.87 | 2.34 | 8.65 |
| 4 | PZ02_Bujnicki_3 | 0.21 | 0.88 | 2.09 | 8.76 |
| 5 | PZ02_Bujnicki_2 | 0.20 | 0.88 | 2.04 | 8.84 |
| 6 | PZ02_Dokholyan_1 | 0.23 | 0.89 | 2.35 | 8.94 |
| 7 | PZ02_Das_2 | 0.29 | 0.97 | 2.86 | 9.74 |
| 8 | PZ02_Chen_1 | 0.27 | 1.00 | 2.69 | 10.00 |
| 9 | PZ02_Das_4 | 0.26 | 1.01 | 2.56 | 10.11 |
| 10 | PZ02_Das_3 | 0.30 | 1.04 | 2.96 | 10.36 |
| 11 | PZ02_Das_5 | 0.35 | 1.05 | 3.48 | 10.52 |
| 12 | PZ02_Wildauer_1 | 0.32 | 1.13 | 3.17 | 11.34 |
| 13 | PZ02_Major_1 | 2.51 | 2.31 | 25.14 | 23.10 |
| | Average | 0.44 | 1.06 | 4.44 | 10.55 |
| | Standard deviation | 0.20 | 0.83 | 2.04 | 8.25 |
| | Median | 2.51 | 2.31 | 25.14 | 23.10 |

**Table S27**. BARNABA report on structures in Puzzle 03.

| No. | 3D RNA model | BBRMSD [nm] | ERMSD [nm] | BBRMSD [Å] | ERMSD [Å] |
|---|---|---|---|---|---|
| 1 | PZ03_Das_5 | 1.26 | 1.33 | 12.57 | 13.34 |
| 2 | PZ03_Das_2 | 1.27 | 1.36 | 12.73 | 13.58 |
| 3 | PZ03_Chen_1 | 0.77 | 1.37 | 7.75 | 13.72 |
| 4 | PZ03_Das_1 | 1.75 | 1.37 | 17.47 | 13.73 |
| 5 | PZ03_Das_3 | 1.86 | 1.42 | 18.56 | 14.21 |
| 6 | PZ03_Das_4 | 1.85 | 1.44 | 18.49 | 14.39 |
| 7 | PZ03_Dokholyan_2 | 1.18 | 1.47 | 11.81 | 14.71 |
| 8 | PZ03_Bujnicki_1 | 1.22 | 1.49 | 12.21 | 14.90 |
| 9 | PZ03_Bujnicki_2 | 1.50 | 1.50 | 14.97 | 14.99 |
| 10 | PZ03_Dokholyan_1 | 1.81 | 1.50 | 18.10 | 15.02 |
| 11 | PZ03_Major_2 | 1.41 | 1.60 | 14.15 | 16.05 |
| 12 | PZ03_Major_1 | 2.53 | 2.03 | 25.30 | 20.27 |
| | Average | 1.53 | 1.49 | 15.34 | 14.91 |
| | Standard deviation | 0.77 | 1.33 | 7.75 | 13.34 |
| | Median | 0.73 | 1.44 | 7.34 | 14.40 |

**Table S28**. BARNABA report on structures in Puzzle 04.

| No. | 3D RNA model | BBRMSD [nm] | ERMSD [nm] | BBRMSD [Å] | ERMSD [Å] |
|---|---|---|---|---|---|
| 1 | PZ04_Das_1 | 0.47 | 0.50 | 4.73 | 5.00 |
| 2 | PZ04_Das_4 | 0.47 | 0.50 | 4.73 | 5.02 |
| 3 | PZ04_Das_3 | 0.47 | 0.50 | 4.73 | 5.02 |
| 4 | PZ04_Das_2 | 0.47 | 0.50 | 4.73 | 5.03 |
| 5 | PZ04_Das_5 | 0.47 | 0.50 | 4.73 | 5.03 |
| 6 | PZ04_Chen_9 | 0.35 | 0.55 | 3.45 | 5.51 |
| 7 | PZ04_Chen_2 | 0.35 | 0.55 | 3.45 | 5.52 |
| 8 | PZ04_Chen_4 | 0.35 | 0.55 | 3.47 | 5.53 |
| 9 | PZ04_Chen_8 | 0.35 | 0.56 | 3.47 | 5.56 |
| 10 | PZ04_Chen_10 | 0.35 | 0.56 | 3.46 | 5.59 |
| 11 | PZ04_Chen_7 | 0.35 | 0.56 | 3.46 | 5.60 |
| 12 | PZ04_Chen_3 | 0.35 | 0.56 | 3.47 | 5.63 |
| 13 | PZ04_Chen_6 | 0.35 | 0.57 | 3.47 | 5.70 |
| 14 | PZ04_Chen_5 | 0.35 | 0.58 | 3.47 | 5.77 |
| 15 | PZ04_Major_1 | 0.46 | 0.59 | 4.65 | 5.89 |
| 16 | PZ04_Chen_1 | 0.35 | 0.59 | 3.45 | 5.90 |
| 17 | PZ04_Dokholyan_1 | 0.56 | 0.63 | 5.59 | 6.31 |
| 18 | PZ04_Dokholyan_2 | 0.56 | 0.63 | 5.59 | 6.31 |
| 19 | PZ04_Santalucia_1 | 0.42 | 0.64 | 4.24 | 6.35 |
| 20 | PZ04_Bujnicki_1 | 0.43 | 0.73 | 4.27 | 7.34 |
| 21 | PZ04_Bujnicki_3 | 0.43 | 0.74 | 4.27 | 7.36 |
| 22 | PZ04_Bujnicki_5 | 0.43 | 0.74 | 4.27 | 7.37 |
| 23 | PZ04_Bujnicki_2 | 0.43 | 0.74 | 4.27 | 7.40 |
| 24 | PZ04_Bujnicki_4 | 0.43 | 0.74 | 4.27 | 7.40 |
| 25 | PZ04_Adamiak_3 | 0.44 | 0.85 | 4.39 | 8.53 |
| 26 | PZ04_Adamiak_2 | 0.46 | 0.87 | 4.58 | 8.70 |
| 27 | PZ04_Adamiak_1 | 0.47 | 0.87 | 4.67 | 8.74 |
| 28 | PZ04_Adamiak_4 | 0.67 | 1.12 | 6.72 | 11.24 |

| | | | | | |
|---|---|---|---|---|---|
| 29 | PZ04_Adamiak_5 | 0.47 | 1.19 | 4.72 | 11.94 |
| 30 | PZ04_Mikolajczak_1 | 1.34 | 1.34 | 13.40 | 13.40 |
| | Average | 0.46 | 0.69 | 4.61 | 6.86 |
| | Standard deviation | 0.35 | 0.50 | 3.45 | 5.00 |
| | Median | 1.34 | 1.34 | 13.40 | 13.40 |

**Table S29**. BARNABA report on structures in Puzzle 05.

| No. | 3D RNA model | BBRMSD [nm] | ERMSD [nm] | BBRMSD [Å] | ERMSD [Å] |
|---|---|---|---|---|---|
| 1 | PZ05_Das_2 | 0.94 | 1.30 | 9.41 | 13.00 |
| 2 | PZ05_Das_1 | 1.02 | 1.33 | 10.20 | 13.26 |
| 3 | PZ05_Bujnicki_2 | 2.11 | 1.44 | 21.11 | 14.38 |
| 4 | PZ05_Adamiak_1 | 1.71 | 1.48 | 17.12 | 14.75 |
| 5 | PZ05_Chen_5 | 3.13 | 1.48 | 31.30 | 14.85 |
| 6 | PZ05_Chen_2 | 2.63 | 1.50 | 26.30 | 15.02 |
| 7 | PZ05_Ding_3 | 2.34 | 1.50 | 23.39 | 15.03 |
| 8 | PZ05_Chen_1 | 2.69 | 1.52 | 26.92 | 15.15 |
| 9 | PZ05_Ding_5 | 2.29 | 1.52 | 22.93 | 15.22 |
| 10 | PZ05_Bujnicki_5 | 2.41 | 1.53 | 24.12 | 15.27 |
| 11 | PZ05_Chen_4 | 3.22 | 1.53 | 32.25 | 15.30 |
| 12 | PZ05_Chen_6 | 3.22 | 1.53 | 32.24 | 15.34 |
| 13 | PZ05_Dokholyan_1 | 2.21 | 1.54 | 22.07 | 15.36 |
| 14 | PZ05_Ding_1 | 2.74 | 1.54 | 27.40 | 15.40 |
| 15 | PZ05_Chen_7 | 2.67 | 1.54 | 26.74 | 15.42 |
| 16 | PZ05_Dokholyan_3 | 2.53 | 1.54 | 25.30 | 15.44 |
| 17 | PZ05_Ding_2 | 1.96 | 1.55 | 19.59 | 15.52 |
| 18 | PZ05_Ding_4 | 1.96 | 1.55 | 19.59 | 15.52 |
| 19 | PZ05_Dokholyan_2 | 2.27 | 1.56 | 22.72 | 15.58 |
| 20 | PZ05_Xiao_1 | 3.65 | 1.57 | 36.48 | 15.66 |
| 21 | PZ05_Bujnicki_1 | 2.53 | 1.58 | 25.32 | 15.75 |
| 22 | PZ05_Xiao_2 | 3.29 | 1.58 | 32.89 | 15.81 |
| 23 | PZ05_Bujnicki_4 | 2.29 | 1.60 | 22.92 | 15.99 |
| 24 | PZ05_Chen_3 | 2.99 | 1.64 | 29.90 | 16.40 |
| 25 | PZ05_Bujnicki_3 | 2.25 | 1.79 | 22.48 | 17.90 |
| | Average | 2.44 | 1.53 | 24.43 | 15.29 |
| | Standard deviation | 0.94 | 1.30 | 9.41 | 13.00 |
| | Median | 3.65 | 1.79 | 36.48 | 17.90 |

**Table S30**. BARNABA report on structures in Puzzle 06.

| No. | 3D RNA model | BBRMSD [nm] | ERMSD [nm] | BBRMSD [Å] | ERMSD [Å] |
|---|---|---|---|---|---|
| 1 | PZ06_Das_1 | 1.45 | 1.17 | 14.51 | 11.74 |
| 2 | PZ06_Das_6 | 1.27 | 1.18 | 12.71 | 11.78 |
| 3 | PZ06_Das_4 | 1.21 | 1.19 | 12.13 | 11.88 |
| 4 | PZ06_Das_2 | 1.41 | 1.20 | 14.05 | 12.02 |
| 5 | PZ06_Das_9 | 1.54 | 1.21 | 15.43 | 12.07 |
| 6 | PZ06_Das_5 | 1.60 | 1.22 | 16.01 | 12.21 |
| 7 | PZ06_Das_8 | 1.84 | 1.24 | 18.40 | 12.37 |
| 8 | PZ06_Das_7 | 1.84 | 1.25 | 18.43 | 12.55 |

| | | | | | |
|---|---|---|---|---|---|
| 9 | PZ06_Das_3 | 1.60 | 1.27 | 16.00 | 12.66 |
| 10 | PZ06_Das_10 | 2.98 | 1.31 | 29.77 | 13.12 |
| 11 | PZ06_Major_6 | 2.45 | 1.34 | 24.48 | 13.38 |
| 12 | PZ06_Major_2 | 2.29 | 1.36 | 22.93 | 13.60 |
| 13 | PZ06_Major_7 | 2.36 | 1.37 | 23.59 | 13.69 |
| 14 | PZ06_Major_1 | 2.36 | 1.38 | 23.62 | 13.76 |
| 15 | PZ06_Major_4 | 2.34 | 1.39 | 23.38 | 13.88 |
| 16 | PZ06_Major_3 | 2.25 | 1.40 | 22.49 | 13.96 |
| 17 | PZ06_Major_5 | 2.51 | 1.45 | 25.12 | 14.45 |
| 18 | PZ06_Bujnicki_1 | 3.76 | 1.46 | 37.59 | 14.60 |
| 19 | PZ06_Dokholyan_3 | 2.70 | 1.48 | 26.98 | 14.80 |
| 20 | PZ06_Bujnicki_2 | 3.15 | 1.49 | 31.51 | 14.88 |
| 21 | PZ06_Chen_4 | 2.28 | 1.50 | 22.84 | 15.02 |
| 22 | PZ06_Dokholyan_5 | 2.33 | 1.51 | 23.30 | 15.13 |
| 23 | PZ06_Dokholyan_4 | 2.59 | 1.52 | 25.88 | 15.21 |
| 24 | PZ06_Chen_1 | 2.51 | 1.52 | 25.05 | 15.21 |
| 25 | PZ06_Chen_3 | 2.42 | 1.52 | 24.22 | 15.22 |
| 26 | PZ06_Dokholyan_1 | 2.71 | 1.52 | 27.09 | 15.24 |
| 27 | PZ06_Dokholyan_2 | 2.74 | 1.52 | 27.39 | 15.24 |
| 28 | PZ06_Chen_6 | 3.65 | 1.54 | 36.54 | 15.40 |
| 29 | PZ06_Chen_2 | 2.29 | 1.54 | 22.91 | 15.40 |
| 30 | PZ06_Chen_5 | 2.40 | 1.55 | 24.04 | 15.47 |
| 31 | PZ06_Bujnicki_4 | 3.26 | 1.56 | 32.57 | 15.63 |
| 32 | PZ06_Chen_7 | 3.14 | 1.58 | 31.44 | 15.77 |
| 33 | PZ06_Dokholyan_6 | 3.81 | 1.58 | 38.14 | 15.83 |
| 34 | PZ06_Bujnicki_3 | 3.25 | 1.63 | 32.50 | 16.27 |
| | Average | 2.42 | 1.41 | 24.21 | 14.10 |
| | Standard deviation | 1.21 | 1.17 | 12.13 | 11.74 |
| | Median | 3.81 | 1.63 | 38.14 | 16.27 |

**Table S31**. BARNABA report on structures in Puzzle 07.

| No. | 3D RNA model | BBRMSD [nm] | ERMSD [nm] | BBRMSD [Å] | ERMSD [Å] |
|---|---|---|---|---|---|
| 1 | PZ07_Das_1 | 2.10 | 1.27 | 20.99 | 12.69 |
| 2 | PZ07_Chen_8 | 3.08 | 1.28 | 30.77 | 12.84 |
| 3 | PZ07_Chen_10 | 3.24 | 1.32 | 32.41 | 13.22 |
| 4 | PZ07_Chen_2 | 3.50 | 1.33 | 34.99 | 13.25 |
| 5 | PZ07_Das_6 | 2.58 | 1.33 | 25.81 | 13.30 |
| 6 | PZ07_Chen_9 | 3.40 | 1.34 | 34.01 | 13.42 |
| 7 | PZ07_Chen_5 | 2.42 | 1.35 | 24.15 | 13.49 |
| 8 | PZ07_Das_2 | 2.42 | 1.36 | 24.16 | 13.56 |
| 9 | PZ07_Chen_4 | 2.85 | 1.36 | 28.48 | 13.59 |
| 10 | PZ07_Bujnicki_3 | 2.70 | 1.36 | 27.04 | 13.64 |
| 11 | PZ07_Chen_3 | 3.05 | 1.37 | 30.50 | 13.67 |
| 12 | PZ07_Das_3 | 2.51 | 1.37 | 25.08 | 13.68 |
| 13 | PZ07_Das_5 | 2.52 | 1.37 | 25.23 | 13.71 |
| 14 | PZ07_Chen_6 | 2.98 | 1.37 | 29.78 | 13.71 |
| 15 | PZ07_Bujnicki_6 | 2.71 | 1.37 | 27.07 | 13.75 |
| 16 | PZ07_Chen_7 | 2.98 | 1.38 | 29.78 | 13.75 |
| 17 | PZ07_Das_8 | 2.74 | 1.40 | 27.37 | 13.99 |

| | | | | | |
|---|---|---|---|---|---|
| 18 | PZ07_Das_4 | 2.58 | 1.40 | 25.83 | 14.02 |
| 19 | PZ07_Das_7 | 2.57 | 1.41 | 25.68 | 14.08 |
| 20 | PZ07_Bujnicki_4 | 2.58 | 1.41 | 25.82 | 14.08 |
| 21 | PZ07_Bujnicki_5 | 2.58 | 1.41 | 25.82 | 14.09 |
| 22 | PZ07_Bujnicki_7 | 2.78 | 1.43 | 27.83 | 14.32 |
| 23 | PZ07_Adamiak_5 | 2.93 | 1.48 | 29.33 | 14.77 |
| 24 | PZ07_Dokholyan_1 | 3.09 | 1.48 | 30.90 | 14.82 |
| 25 | PZ07_Ding_4 | 3.27 | 1.50 | 32.66 | 14.98 |
| 26 | PZ07_Ding_6 | 2.99 | 1.51 | 29.92 | 15.09 |
| 27 | PZ07_Ding_1 | 2.98 | 1.51 | 29.76 | 15.10 |
| 28 | PZ07_Ding_10 | 3.73 | 1.52 | 37.26 | 15.17 |
| 29 | PZ07_Ding_8 | 2.80 | 1.52 | 28.05 | 15.20 |
| 30 | PZ07_Ding_9 | 2.75 | 1.53 | 27.47 | 15.26 |
| 31 | PZ07_Ding_7 | 3.05 | 1.53 | 30.54 | 15.28 |
| 32 | PZ07_Adamiak_3 | 3.15 | 1.55 | 31.53 | 15.47 |
| 33 | PZ07_Adamiak_1 | 2.97 | 1.55 | 29.75 | 15.50 |
| 34 | PZ07_Ding_2 | 3.34 | 1.56 | 33.42 | 15.61 |
| 35 | PZ07_Ding_3 | 3.25 | 1.56 | 32.46 | 15.63 |
| 36 | PZ07_Ding_5 | 3.02 | 1.58 | 30.25 | 15.79 |
| 37 | PZ07_Bujnicki_2 | 6.11 | 1.58 | 61.11 | 15.82 |
| 38 | PZ07_Adamiak_2 | 2.97 | 1.58 | 29.66 | 15.84 |
| 39 | PZ07_Major_7 | 2.90 | 1.58 | 29.04 | 15.84 |
| 40 | PZ07_Major_5 | 2.90 | 1.58 | 29.03 | 15.84 |
| 41 | PZ07_Major_2 | 2.88 | 1.59 | 28.78 | 15.87 |
| 42 | PZ07_Adamiak_4 | 3.07 | 1.59 | 30.74 | 15.87 |
| 43 | PZ07_Major_9 | 2.80 | 1.59 | 28.01 | 15.93 |
| 44 | PZ07_Major_6 | 2.84 | 1.60 | 28.36 | 15.98 |
| 45 | PZ07_Major_1 | 2.96 | 1.60 | 29.60 | 16.05 |
| 46 | PZ07_Major_8 | 2.82 | 1.61 | 28.19 | 16.08 |
| 47 | PZ07_Major_3 | 2.76 | 1.61 | 27.61 | 16.12 |
| 48 | PZ07_Major_4 | 2.82 | 1.62 | 28.17 | 16.23 |
| 49 | PZ07_Major_10 | 3.22 | 1.62 | 32.16 | 16.25 |
| 50 | PZ07_Bujnicki_1 | 4.30 | 1.67 | 43.04 | 16.68 |
| 51 | PZ07_Chen_1 | 2.90 | 1.80 | 29.01 | 18.04 |
| 52 | PZ07_Dokholyan_2 | 3.74 | 1.82 | 37.37 | 18.17 |
| | Average | 3.00 | 1.49 | 30.03 | 14.89 |
| | Standard deviation | 2.10 | 1.27 | 20.99 | 12.69 |
| | Median | 6.11 | 1.82 | 61.11 | 18.17 |

**Table S32**. BARNABA report on structures in Puzzle 08

| No. | 3D RNA model | BBRMSD [nm] | ERMSD [nm] | BBRMSD [Å] | ERMSD [Å] |
|---|---|---|---|---|---|
| 1 | PZ08_Das_2 | 1.13 | 1.10 | 11.30 | 11.04 |
| 2 | PZ08_Das_1 | 0.64 | 1.14 | 6.40 | 11.40 |
| 3 | PZ08_Das_3 | 0.47 | 1.15 | 4.73 | 11.51 |
| 4 | PZ08_Bujnicki_4 | 1.29 | 1.17 | 12.95 | 11.66 |
| 5 | PZ08_Das_6 | 1.12 | 1.17 | 11.21 | 11.68 |
| 6 | PZ08_Das_5 | 1.16 | 1.18 | 11.62 | 11.78 |
| 7 | PZ08_Bujnicki_2 | 1.27 | 1.19 | 12.66 | 11.85 |
| 8 | PZ08_Das_4 | 1.12 | 1.19 | 11.20 | 11.87 |

| | | | | | |
|---|---|---|---|---|---|
| 9 | PZ08_Bujnicki_5 | 1.16 | 1.23 | 11.57 | 12.32 |
| 10 | PZ08_Bujnicki_1 | 1.31 | 1.24 | 13.11 | 12.44 |
| 11 | PZ08_Bujnicki_8 | 1.20 | 1.25 | 12.02 | 12.46 |
| 12 | PZ08_Bujnicki_9 | 0.68 | 1.25 | 6.82 | 12.55 |
| 13 | PZ08_Chen_1 | 1.33 | 1.29 | 13.30 | 12.85 |
| 14 | PZ08_Bujnicki_7 | 0.67 | 1.29 | 6.67 | 12.92 |
| 15 | PZ08_Bujnicki_6 | 1.27 | 1.30 | 12.65 | 12.97 |
| 16 | PZ08_Bujnicki_3 | 1.20 | 1.31 | 12.03 | 13.09 |
| 17 | PZ08_Chen_2 | 1.30 | 1.31 | 13.01 | 13.14 |
| 18 | PZ08_Chen_3 | 1.14 | 1.32 | 11.39 | 13.23 |
| 19 | PZ08_Chen_5 | 1.18 | 1.33 | 11.83 | 13.26 |
| 20 | PZ08_Chen_10 | 1.24 | 1.33 | 12.38 | 13.30 |
| 21 | PZ08_Ding_4 | 1.59 | 1.35 | 15.88 | 13.52 |
| 22 | PZ08_Chen_4 | 1.30 | 1.37 | 13.05 | 13.70 |
| 23 | PZ08_Ding_7 | 1.33 | 1.37 | 13.32 | 13.73 |
| 24 | PZ08_Ding_5 | 1.32 | 1.39 | 13.18 | 13.88 |
| 25 | PZ08_Ding_8 | 1.58 | 1.40 | 15.75 | 13.98 |
| 26 | PZ08_Ding_10 | 1.21 | 1.40 | 12.07 | 14.04 |
| 27 | PZ08_Bujnicki_10 | 1.13 | 1.41 | 11.30 | 14.09 |
| 28 | PZ08_Chen_7 | 1.29 | 1.42 | 12.88 | 14.21 |
| 29 | PZ08_Adamiak_2 | 1.51 | 1.42 | 15.13 | 14.23 |
| 30 | PZ08_Ding_6 | 1.46 | 1.43 | 14.64 | 14.28 |
| 31 | PZ08_Adamiak_1 | 1.46 | 1.44 | 14.62 | 14.37 |
| 32 | PZ08_Ding_1 | 1.26 | 1.44 | 12.65 | 14.40 |
| 33 | PZ08_Ding_3 | 1.16 | 1.45 | 11.62 | 14.47 |
| 34 | PZ08_Ding_2 | 1.42 | 1.46 | 14.23 | 14.62 |
| 35 | PZ08_Ding_9 | 1.71 | 1.47 | 17.06 | 14.67 |
| 36 | PZ08_Chen_6 | 1.18 | 1.49 | 11.82 | 14.92 |
| 37 | PZ08_Chen_8 | 1.60 | 1.49 | 15.96 | 14.93 |
| 38 | PZ08_Chen_9 | 1.13 | 1.55 | 11.34 | 15.46 |
| 39 | PZ08_Dokholyan_1 | 2.46 | 1.60 | 24.56 | 15.97 |
| 40 | PZ08_Dokholyan_3 | 2.57 | 1.67 | 25.73 | 16.67 |
| 41 | PZ08_Dokholyan_2 | 1.87 | 1.71 | 18.70 | 17.10 |
| 42 | PZ08_Dokholyan_4 | 3.17 | 2.06 | 31.68 | 20.57 |
| | Average | 1.35 | 1.37 | 13.48 | 13.69 |
| | Standard deviation | 0.47 | 1.10 | 4.73 | 11.04 |
| | Median | 3.17 | 2.06 | 31.68 | 20.57 |

**Table S33**. BARNABA report on structures in Puzzle 9.

| No. | 3D RNA model | BBRMSD [nm] | ERMSD [nm] | BBRMSD [Å] | ERMSD [Å] |
|---|---|---|---|---|---|
| 1 | PZ09_Das_1 | 0.90 | 1.20 | 9.03 | 11.95 |
| 2 | PZ09_Das_4 | 1.00 | 1.24 | 9.95 | 12.42 |
| 3 | PZ09_Das_5 | 0.80 | 1.25 | 7.99 | 12.49 |
| 4 | PZ09_Das_2 | 0.81 | 1.29 | 8.13 | 12.90 |
| 5 | PZ09_Das_8 | 1.12 | 1.34 | 11.22 | 13.36 |
| 6 | PZ09_Das_3 | 0.88 | 1.35 | 8.84 | 13.54 |
| 7 | PZ09_Das_7 | 0.94 | 1.35 | 9.43 | 13.55 |
| 8 | PZ09_Das_6 | 0.91 | 1.36 | 9.07 | 13.56 |
| 9 | PZ09_Chen_1 | 0.64 | 1.36 | 6.40 | 13.59 |

| | | | | | |
|---|---|---|---|---|---|
| 10 | PZ09_Chen_8 | 0.65 | 1.39 | 6.46 | 13.90 |
| 11 | PZ09_Das_9 | 1.00 | 1.43 | 10.01 | 14.33 |
| 12 | PZ09_Chen_5 | 0.63 | 1.44 | 6.33 | 14.42 |
| 13 | PZ09_Chen_6 | 0.65 | 1.44 | 6.52 | 14.44 |
| 14 | PZ09_Chen_4 | 0.64 | 1.45 | 6.40 | 14.52 |
| 15 | PZ09_Chen_2 | 0.63 | 1.46 | 6.32 | 14.56 |
| 16 | PZ09_Chen_3 | 0.63 | 1.46 | 6.32 | 14.60 |
| 17 | PZ09_Chen_7 | 0.59 | 1.51 | 5.90 | 15.10 |
| 18 | PZ09_Ding_7 | 0.98 | 1.54 | 9.77 | 15.44 |
| 19 | PZ09_Bujnicki_3 | 0.60 | 1.57 | 5.98 | 15.69 |
| 20 | PZ09_Bujnicki_4 | 0.60 | 1.57 | 5.98 | 15.69 |
| 21 | PZ09_Bujnicki_5 | 0.60 | 1.57 | 5.98 | 15.69 |
| 22 | PZ09_Ding_5 | 0.93 | 1.68 | 9.25 | 16.76 |
| 23 | PZ09_Ding_10 | 1.16 | 1.68 | 11.61 | 16.81 |
| 24 | PZ09_Ding_4 | 0.99 | 1.68 | 9.94 | 16.83 |
| 25 | PZ09_Ding_3 | 0.93 | 1.69 | 9.31 | 16.90 |
| 26 | PZ09_Ding_9 | 1.01 | 1.69 | 10.13 | 16.95 |
| 27 | PZ09_Ding_6 | 1.25 | 1.70 | 12.49 | 17.03 |
| 28 | PZ09_Dokholyan_1 | 2.06 | 1.71 | 20.56 | 17.07 |
| 29 | PZ09_Dokholyan_2 | 2.61 | 1.71 | 26.11 | 17.11 |
| 30 | PZ09_Ding_8 | 1.15 | 1.71 | 11.53 | 17.13 |
| 31 | PZ09_Ding_1 | 1.01 | 1.75 | 10.06 | 17.47 |
| 32 | PZ09_Ding_2 | 1.02 | 1.75 | 10.24 | 17.48 |
| 33 | PZ09_Dokholyan_4 | 2.00 | 1.78 | 19.99 | 17.75 |
| 34 | PZ09_Dokholyan_3 | 2.28 | 1.78 | 22.81 | 17.80 |
| | Average | 1.02 | 1.53 | 10.18 | 15.26 |
| | Standard deviation | 0.59 | 1.20 | 5.90 | 11.95 |
| | Average | 1.02 | 1.53 | 10.18 | 15.26 |

**Table S34**. BARNABA report on structures in Puzzle 10.

| No. | 3D RNA model | BBRMSD [nm] | ERMSD [nm] | BBRMSD [Å] | ERMSD [Å] |
|---|---|---|---|---|---|
| 1 | PZ10_Das_5 | 1.08 | 0.82 | 10.77 | 8.16 |
| 2 | PZ10_Das_4 | 0.73 | 0.89 | 7.31 | 8.92 |
| 3 | PZ10_Das_2 | 1.08 | 0.91 | 10.81 | 9.07 |
| 4 | PZ10_Das_1 | 0.79 | 0.91 | 7.86 | 9.14 |
| 5 | PZ10_Das_3 | 0.70 | 0.92 | 7.03 | 9.16 |
| 6 | PZ10_Bujnicki_8 | 1.05 | 1.00 | 10.50 | 10.01 |
| 7 | PZ10_Bujnicki_1 | 1.05 | 1.01 | 10.54 | 10.05 |
| 8 | PZ10_Bujnicki_9 | 1.06 | 1.04 | 10.60 | 10.40 |
| 9 | PZ10_Bujnicki_7 | 1.05 | 1.05 | 10.54 | 10.49 |
| 10 | PZ10_Bujnicki_3 | 1.06 | 1.08 | 10.60 | 10.76 |
| 11 | PZ10_Bujnicki_5 | 1.06 | 1.08 | 10.56 | 10.83 |
| 12 | PZ10_Bujnicki_10 | 1.07 | 1.09 | 10.66 | 10.89 |
| 13 | PZ10_Bujnicki_2 | 1.07 | 1.10 | 10.67 | 10.96 |
| 14 | PZ10_Bujnicki_6 | 1.06 | 1.12 | 10.63 | 11.20 |
| 15 | PZ10_Bujnicki_4 | 0.96 | 1.22 | 9.55 | 12.21 |
| 16 | PZ10_Chen_1 | 1.17 | 1.25 | 11.73 | 12.47 |
| 17 | PZ10_Ding_4 | 1.36 | 1.43 | 13.64 | 14.35 |
| 18 | PZ10_Ding_3 | 1.32 | 1.45 | 13.23 | 14.51 |

| | | | | | |
|---|---|---|---|---|---|
| 19 | PZ10_Ding_8 | 1.61 | 1.46 | 16.13 | 14.59 |
| 20 | PZ10_Ding_6 | 1.82 | 1.46 | 18.21 | 14.65 |
| 21 | PZ10_Ding_1 | 1.34 | 1.48 | 13.39 | 14.83 |
| 22 | PZ10_Ding_10 | 1.35 | 1.49 | 13.52 | 14.88 |
| 23 | PZ10_Ding_7 | 1.68 | 1.50 | 16.80 | 15.00 |
| 24 | PZ10_Ding_2 | 1.45 | 1.51 | 14.54 | 15.13 |
| 25 | PZ10_Ding_5 | 1.72 | 1.51 | 17.25 | 15.14 |
| 26 | PZ10_Ding_9 | 1.72 | 1.55 | 17.22 | 15.46 |
| | Average | 1.21 | 1.20 | 12.09 | 12.05 |
| | Standard deviation | 0.70 | 0.82 | 7.03 | 8.16 |
| | Median | 1.82 | 1.55 | 18.21 | 15.46 |

**Table S35**. BARNABA report on structures in Puzzle 11.

| No. | 3D RNA model | BBRMSD [nm] | ERMSD [nm] | BBRMSD [Å] | ERMSD [Å] |
|---|---|---|---|---|---|
| 1 | PZ11_Das_2 | 0.87 | 0.99 | 8.72 | 9.94 |
| 2 | PZ11_Das_4 | 0.69 | 1.03 | 6.91 | 10.28 |
| 3 | PZ11_Chen_1 | 0.56 | 1.07 | 5.60 | 10.65 |
| 4 | PZ11_Chen_3 | 0.49 | 1.07 | 4.91 | 10.69 |
| 5 | PZ11_Das_1 | 0.57 | 1.08 | 5.71 | 10.78 |
| 6 | PZ11_Das_8 | 0.98 | 1.09 | 9.76 | 10.89 |
| 7 | PZ11_Chen_9 | 0.78 | 1.09 | 7.76 | 10.90 |
| 8 | PZ11_Chen_5 | 0.88 | 1.09 | 8.79 | 10.93 |
| 9 | PZ11_Bujnicki_9 | 0.52 | 1.09 | 5.18 | 10.95 |
| 10 | PZ11_Chen_2 | 0.74 | 1.09 | 7.37 | 10.95 |
| 11 | PZ11_Das_7 | 0.82 | 1.10 | 8.18 | 11.01 |
| 12 | PZ11_Chen_10 | 0.84 | 1.10 | 8.37 | 11.02 |
| 13 | PZ11_Adamiak_5 | 0.95 | 1.11 | 9.50 | 11.15 |
| 14 | PZ11_Chen_7 | 0.76 | 1.12 | 7.56 | 11.24 |
| 15 | PZ11_Chen_8 | 0.81 | 1.14 | 8.06 | 11.40 |
| 16 | PZ11_Das_6 | 0.66 | 1.15 | 6.64 | 11.45 |
| 17 | PZ11_Bujnicki_1 | 0.66 | 1.15 | 6.56 | 11.54 |
| 18 | PZ11_Ding_1 | 0.57 | 1.16 | 5.67 | 11.56 |
| 19 | PZ11_Chen_4 | 0.65 | 1.16 | 6.54 | 11.58 |
| 20 | PZ11_Das_10 | 0.97 | 1.17 | 9.72 | 11.69 |
| 21 | PZ11_Das_3 | 0.81 | 1.17 | 8.10 | 11.72 |
| 22 | PZ11_Adamiak_7 | 0.74 | 1.18 | 7.42 | 11.76 |
| 23 | PZ11_Bujnicki_3 | 0.69 | 1.19 | 6.86 | 11.94 |
| 24 | PZ11_Chen_6 | 0.70 | 1.21 | 7.02 | 12.08 |
| 25 | PZ11_Ding_6 | 0.71 | 1.21 | 7.06 | 12.15 |
| 26 | PZ11_Ding_7 | 0.57 | 1.22 | 5.69 | 12.18 |
| 27 | PZ11_Das_5 | 0.77 | 1.23 | 7.68 | 12.35 |
| 28 | PZ11_Ding_10 | 1.13 | 1.24 | 11.32 | 12.41 |
| 29 | PZ11_Bujnicki_4 | 0.66 | 1.24 | 6.64 | 12.43 |
| 30 | PZ11_Das_9 | 1.93 | 1.25 | 19.34 | 12.46 |
| 31 | PZ11_Xiao_3 | 1.13 | 1.25 | 11.32 | 12.49 |
| 32 | PZ11_Ding_2 | 1.13 | 1.27 | 11.26 | 12.73 |
| 33 | PZ11_Ding_3 | 1.06 | 1.28 | 10.64 | 12.78 |
| 34 | PZ11_Ding_9 | 0.83 | 1.30 | 8.28 | 13.01 |
| 35 | PZ11_Xiao_2 | 1.32 | 1.33 | 13.16 | 13.31 |

| | | | | | |
|---|---|---|---|---|---|
| 36 | PZ11_Ding_8 | 0.66 | 1.33 | 6.61 | 13.33 |
| 37 | PZ11_Ding_5 | 0.66 | 1.34 | 6.62 | 13.39 |
| 38 | PZ11_Adamiak_6 | 0.67 | 1.34 | 6.67 | 13.43 |
| 39 | PZ11_Ding_4 | 0.64 | 1.35 | 6.44 | 13.46 |
| 40 | PZ11_Adamiak_4 | 0.93 | 1.35 | 9.25 | 13.50 |
| 41 | PZ11_Adamiak_9 | 0.87 | 1.36 | 8.74 | 13.65 |
| 42 | PZ11_Bujnicki_6 | 1.29 | 1.38 | 12.89 | 13.81 |
| 43 | PZ11_Adamiak_2 | 1.11 | 1.39 | 11.14 | 13.91 |
| 44 | PZ11_Adamiak_10 | 1.03 | 1.39 | 10.34 | 13.95 |
| 45 | PZ11_Adamiak_1 | 0.89 | 1.40 | 8.94 | 13.96 |
| 46 | PZ11_Bujnicki_10 | 0.81 | 1.42 | 8.12 | 14.24 |
| 47 | PZ11_Adamiak_8 | 1.10 | 1.44 | 10.97 | 14.37 |
| 48 | PZ11_Bujnicki_7 | 0.78 | 1.44 | 7.75 | 14.41 |
| 49 | PZ11_Bujnicki_5 | 0.99 | 1.45 | 9.87 | 14.47 |
| 50 | PZ11_Adamiak_3 | 1.01 | 1.45 | 10.07 | 14.51 |
| 51 | PZ11_Bujnicki_8 | 1.22 | 1.46 | 12.22 | 14.57 |
| 52 | PZ11_Bujnicki_2 | 1.92 | 1.48 | 19.23 | 14.81 |
| 53 | PZ11_Xiao_1 | 0.97 | 1.53 | 9.72 | 15.28 |
| | Average | 0.88 | 1.24 | 8.77 | 12.44 |
| | Standard deviation | 0.49 | 0.99 | 4.91 | 9.94 |
| | Median | 1.93 | 1.53 | 19.34 | 15.28 |

**Table S36**. BARNABA report on structures in Puzzle 12.

| No. | 3D RNA model | BBRMSD [nm] | ERMSD [nm] | BBRMSD [Å] | ERMSD [Å] |
|---|---|---|---|---|---|
| 1 | PZ12_Das_2 | 1.40 | 1.16 | 13.97 | 11.62 |
| 2 | PZ12_Das_5 | 1.47 | 1.28 | 14.65 | 12.80 |
| 3 | PZ12_Das_1 | 1.47 | 1.30 | 14.69 | 13.03 |
| 4 | PZ12_Das_8 | 1.35 | 1.31 | 13.52 | 13.10 |
| 5 | PZ12_Das_10 | 1.80 | 1.32 | 17.98 | 13.17 |
| 6 | PZ12_Das_7 | 1.33 | 1.34 | 13.27 | 13.38 |
| 7 | PZ12_Das_4 | 1.83 | 1.34 | 18.35 | 13.43 |
| 8 | PZ12_Das_3 | 1.44 | 1.35 | 14.40 | 13.45 |
| 9 | PZ12_Adamiak_1 | 1.75 | 1.37 | 17.54 | 13.74 |
| 10 | PZ12_Bujnicki_10 | 1.41 | 1.39 | 14.10 | 13.89 |
| 11 | PZ12_Bujnicki_4 | 1.61 | 1.40 | 16.10 | 13.98 |
| 12 | PZ12_Bujnicki_3 | 1.24 | 1.42 | 12.41 | 14.24 |
| 13 | PZ12_Adamiak_3 | 1.64 | 1.43 | 16.44 | 14.31 |
| 14 | PZ12_Das_9 | 1.42 | 1.44 | 14.17 | 14.35 |
| 15 | PZ12_Das_6 | 1.41 | 1.44 | 14.11 | 14.41 |
| 16 | PZ12_Adamiak_2 | 1.63 | 1.44 | 16.31 | 14.41 |
| 17 | PZ12_Chen_1 | 2.34 | 1.44 | 23.41 | 14.45 |
| 18 | PZ12_Chen_8 | 1.86 | 1.45 | 18.65 | 14.47 |
| 19 | PZ12_Bujnicki_5 | 1.30 | 1.46 | 13.04 | 14.56 |
| 20 | PZ12_Chen_7 | 2.09 | 1.47 | 20.92 | 14.73 |
| 21 | PZ12_Bujnicki_6 | 1.61 | 1.47 | 16.14 | 14.75 |
| 22 | PZ12_Bujnicki_1 | 2.24 | 1.48 | 22.35 | 14.80 |
| 23 | PZ12_Bujnicki_8 | 1.14 | 1.48 | 11.38 | 14.85 |
| 24 | PZ12_Chen_6 | 2.05 | 1.50 | 20.54 | 14.97 |
| 25 | PZ12_Chen_9 | 1.90 | 1.51 | 18.98 | 15.08 |

| | | | | | |
|---|---|---|---|---|---|
| 26 | PZ12_Ding_12 | 1.06 | 1.51 | 10.60 | 15.09 |
| 27 | PZ12_Bujnicki_2 | 1.92 | 1.51 | 19.25 | 15.14 |
| 28 | PZ12_Ding_2 | 2.11 | 1.52 | 21.12 | 15.18 |
| 29 | PZ12_Ding_11 | 1.07 | 1.53 | 10.74 | 15.25 |
| 30 | PZ12_Chen_2 | 1.95 | 1.53 | 19.52 | 15.27 |
| 31 | PZ12_Chen_3 | 1.98 | 1.53 | 19.76 | 15.29 |
| 32 | PZ12_Ding_3 | 1.82 | 1.53 | 18.20 | 15.32 |
| 33 | PZ12_Ding_10 | 1.94 | 1.54 | 19.35 | 15.37 |
| 34 | PZ12_Chen_10 | 2.01 | 1.54 | 20.11 | 15.41 |
| 35 | PZ12_Ding_5 | 2.18 | 1.55 | 21.76 | 15.48 |
| 36 | PZ12_Ding_8 | 1.34 | 1.55 | 13.41 | 15.53 |
| 37 | PZ12_Bujnicki_7 | 1.68 | 1.56 | 16.83 | 15.56 |
| 38 | PZ12_Chen_4 | 1.68 | 1.56 | 16.80 | 15.58 |
| 39 | PZ12_Weeks_3 | 2.06 | 1.56 | 20.57 | 15.64 |
| 40 | PZ12_Ding_4 | 1.42 | 1.57 | 14.24 | 15.73 |
| 41 | PZ12_Ding_7 | 1.85 | 1.57 | 18.55 | 15.74 |
| 42 | PZ12_Chen_5 | 2.09 | 1.58 | 20.93 | 15.76 |
| 43 | PZ12_Ding_6 | 1.61 | 1.58 | 16.07 | 15.79 |
| 44 | PZ12_Bujnicki_9 | 1.41 | 1.58 | 14.08 | 15.84 |
| 45 | PZ12_Ding_9 | 1.52 | 1.60 | 15.25 | 16.05 |
| 46 | PZ12_Weeks_2 | 2.09 | 1.61 | 20.92 | 16.11 |
| 47 | PZ12_Weeks_1 | 2.12 | 1.61 | 21.23 | 16.12 |
| 48 | PZ12_Ding_1 | 1.35 | 1.63 | 13.52 | 16.25 |
| 49 | PZ12_Xiao_3 | 3.63 | 1.87 | 36.26 | 18.67 |
| 50 | PZ12_Xiao_1 | 3.01 | 2.11 | 30.14 | 21.09 |
| 51 | PZ12_Xiao_2 | 2.97 | 2.12 | 29.72 | 21.20 |
| | Average | 1.78 | 1.51 | 17.77 | 15.09 |
| | Standard deviation | 1.06 | 1.16 | 10.60 | 11.62 |
| | Median | 3.63 | 2.12 | 36.26 | 21.20 |

**Table S37**. BARNABA report on structures in Puzzle 13.

| No. | 3D RNA model | BBRMSD [nm] | ERMSD [nm] | BBRMSD [Å] | ERMSD [Å] |
|---|---|---|---|---|---|
| 1 | PZ13_Chen_6 | 1.68 | 1.23 | 16.79 | 12.30 |
| 2 | PZ13_Chen_8 | 1.53 | 1.27 | 15.35 | 12.69 |
| 3 | PZ13_Das_1 | 0.78 | 1.27 | 7.79 | 12.75 |
| 4 | PZ13_Das_2 | 0.55 | 1.28 | 5.53 | 12.78 |
| 5 | PZ13_Das_4 | 1.17 | 1.31 | 11.67 | 13.13 |
| 6 | PZ13_Adamiak_1 | 1.00 | 1.32 | 10.04 | 13.23 |
| 7 | PZ13_Das_7 | 0.56 | 1.34 | 5.63 | 13.40 |
| 8 | PZ13_Chen_3 | 1.00 | 1.34 | 10.00 | 13.44 |
| 9 | PZ13_Chen_9 | 1.60 | 1.37 | 16.03 | 13.66 |
| 10 | PZ13_Chen_7 | 1.58 | 1.37 | 15.77 | 13.67 |
| 11 | PZ13_Chen_5 | 0.62 | 1.37 | 6.17 | 13.74 |
| 12 | PZ13_Ding_3 | 1.40 | 1.37 | 14.02 | 13.75 |
| 13 | PZ13_Das_10 | 1.33 | 1.38 | 13.30 | 13.81 |
| 14 | PZ13_Das_8 | 1.42 | 1.38 | 14.24 | 13.82 |
| 15 | PZ13_Das_3 | 0.90 | 1.38 | 9.03 | 13.84 |
| 16 | PZ13_Chen_2 | 0.97 | 1.39 | 9.74 | 13.86 |
| 17 | PZ13_Bujnicki_8 | 1.30 | 1.39 | 13.01 | 13.90 |

| | | | | | |
|---|---|---|---|---|---|
| 18 | PZ13_Bujnicki_5 | 1.27 | 1.40 | 12.74 | 13.96 |
| 19 | PZ13_Ding_5 | 1.54 | 1.41 | 15.44 | 14.11 |
| 20 | PZ13_Das_5 | 1.32 | 1.42 | 13.24 | 14.19 |
| 21 | PZ13_Ding_7 | 1.69 | 1.43 | 16.88 | 14.27 |
| 22 | PZ13_Das_6 | 1.52 | 1.44 | 15.22 | 14.42 |
| 23 | PZ13_Ding_6 | 1.51 | 1.44 | 15.10 | 14.43 |
| 24 | PZ13_Bujnicki_6 | 1.59 | 1.44 | 15.92 | 14.43 |
| 25 | PZ13_Bujnicki_7 | 1.33 | 1.46 | 13.30 | 14.59 |
| 26 | PZ13_Ding_10 | 1.74 | 1.46 | 17.42 | 14.59 |
| 27 | PZ13_Ding_1 | 1.69 | 1.46 | 16.89 | 14.62 |
| 28 | PZ13_Chen_1 | 1.56 | 1.46 | 15.64 | 14.64 |
| 29 | PZ13_Ding_4 | 1.46 | 1.46 | 14.63 | 14.65 |
| 30 | PZ13_Bujnicki_2 | 1.43 | 1.47 | 14.28 | 14.66 |
| 31 | PZ13_Ding_2 | 1.71 | 1.47 | 17.11 | 14.71 |
| 32 | PZ13_Ding_8 | 1.53 | 1.48 | 15.31 | 14.75 |
| 33 | PZ13_Bujnicki_3 | 0.88 | 1.49 | 8.75 | 14.92 |
| 34 | PZ13_Ding_9 | 1.82 | 1.49 | 18.16 | 14.94 |
| 35 | PZ13_Das_9 | 1.26 | 1.50 | 12.56 | 15.04 |
| 36 | PZ13_Bujnicki_1 | 0.96 | 1.59 | 9.57 | 15.89 |
| 37 | PZ13_Bujnicki_4 | 1.59 | 1.59 | 15.88 | 15.92 |
| 38 | PZ13_Chen_4 | 1.60 | 1.60 | 15.98 | 15.97 |
| 39 | PZ13_Xiao_5 | 2.93 | 1.61 | 29.33 | 16.12 |
| 40 | PZ13_Dokholyan_2 | 2.64 | 1.61 | 26.40 | 16.14 |
| 41 | PZ13_Dokholyan_1 | 2.28 | 1.78 | 22.81 | 17.79 |
| 42 | PZ13_Dokholyan_5 | 1.88 | 1.80 | 18.80 | 18.01 |
| 43 | PZ13_Bujnicki_10 | 1.56 | 1.95 | 15.64 | 19.53 |
| 44 | PZ13_Bujnicki_9 | 1.75 | 1.97 | 17.51 | 19.65 |
| 45 | PZ13_Dokholyan_3 | 2.35 | 1.98 | 23.52 | 19.75 |
| 46 | PZ13_Xiao_1 | 1.76 | 2.03 | 17.63 | 20.34 |
| 47 | PZ13_Xiao_6 | 2.82 | 2.05 | 28.24 | 20.46 |
| 48 | PZ13_Xiao_4 | 2.85 | 2.07 | 28.51 | 20.67 |
| 49 | PZ13_Xiao_10 | 2.81 | 2.07 | 28.07 | 20.72 |
| 50 | PZ13_Xiao_9 | 3.15 | 2.08 | 31.54 | 20.76 |
| 51 | PZ13_Xiao_8 | 2.93 | 2.08 | 29.25 | 20.80 |
| 52 | PZ13_Dokholyan_4 | 2.32 | 2.11 | 23.20 | 21.07 |
| 53 | PZ13_Xiao_7 | 2.78 | 2.18 | 27.85 | 21.77 |
| 54 | PZ13_Xiao_3 | 1.76 | 2.31 | 17.56 | 23.09 |
| 55 | PZ13_Xiao_2 | 2.51 | 2.34 | 25.08 | 23.37 |
| | Average | 1.66 | 1.60 | 16.64 | 15.95 |
| | Standard deviation | 0.55 | 1.23 | 5.53 | 12.30 |
| | Median | 3.15 | 2.34 | 31.54 | 23.37 |

**Table S38**. BARNABA report on structures in Puzzle 14a.

| No. | 3D RNA model | BBRMSD [nm] | ERMSD [nm] | BBRMSD [Å] | ERMSD [Å] |
|---|---|---|---|---|---|
| 1 | PZ14_ChenPostExp_8 | 1.58 | 1.01 | 15.79 | 10.05 |
| 2 | PZ14_DasPreExp_9 | 0.80 | 1.05 | 8.02 | 10.49 |
| 3 | PZ14_DasPostExp_2 | 1.23 | 1.05 | 12.34 | 10.53 |
| 4 | PZ14_DasPreExp_6 | 0.80 | 1.08 | 7.97 | 10.78 |
| 5 | PZ14_DasPostExp_5 | 1.25 | 1.10 | 12.54 | 11.02 |

| | | | | | |
|---|---|---|---|---|---|
| 6 | PZ14_ChenPostExp_3 | 1.71 | 1.10 | 17.15 | 11.04 |
| 7 | PZ14_AdamiakPreExp_1 | 1.69 | 1.13 | 16.85 | 11.30 |
| 8 | PZ14_BujnickiPreExp_5 | 0.78 | 1.14 | 7.77 | 11.36 |
| 9 | PZ14_DasPreExp_7 | 1.28 | 1.14 | 12.80 | 11.38 |
| 10 | PZ14_AdamiakPostExp_2 | 1.53 | 1.14 | 15.28 | 11.44 |
| 11 | PZ14_BujnickiPreExp_6 | 1.15 | 1.15 | 11.50 | 11.48 |
| 12 | PZ14_AdamiakPostExp_1 | 1.60 | 1.15 | 15.97 | 11.49 |
| 13 | PZ14_DasPostExp_4 | 1.34 | 1.15 | 13.38 | 11.53 |
| 14 | PZ14_DasPreExp_8 | 0.86 | 1.15 | 8.57 | 11.53 |
| 15 | PZ14_BujnickiPreExp_4 | 0.61 | 1.15 | 6.09 | 11.54 |
| 16 | PZ14_DasPostExp_10 | 1.02 | 1.16 | 10.18 | 11.58 |
| 17 | PZ14_DasPreExp_5 | 1.02 | 1.16 | 10.18 | 11.58 |
| 18 | PZ14_DasPostExp_6 | 1.28 | 1.16 | 12.83 | 11.61 |
| 19 | PZ14_DasPreExp_1 | 1.28 | 1.16 | 12.83 | 11.61 |
| 20 | PZ14_DasPostExp_8 | 1.26 | 1.16 | 12.61 | 11.63 |
| 21 | PZ14_DasPreExp_3 | 1.26 | 1.16 | 12.61 | 11.63 |
| 22 | PZ14_ChenPostExp_4 | 1.63 | 1.16 | 16.32 | 11.65 |
| 23 | PZ14_AdamiakPreExp_2 | 1.71 | 1.18 | 17.10 | 11.82 |
| 24 | PZ14_ChenPostExp_10 | 1.71 | 1.18 | 17.12 | 11.85 |
| 25 | PZ14_ChenPostExp_5 | 0.86 | 1.21 | 8.56 | 12.08 |
| 26 | PZ14_DasPostExp_7 | 1.23 | 1.22 | 12.27 | 12.22 |
| 27 | PZ14_DasPreExp_2 | 1.23 | 1.22 | 12.27 | 12.22 |
| 28 | PZ14_ChenPostExp_9 | 1.11 | 1.22 | 11.11 | 12.23 |
| 29 | PZ14_DasPreExp_4 | 1.78 | 1.23 | 17.81 | 12.29 |
| 30 | PZ14_ChenPostExp_6 | 0.76 | 1.23 | 7.61 | 12.29 |
| 31 | PZ14_BujnickiPostExp_1 | 1.46 | 1.24 | 14.61 | 12.36 |
| 32 | PZ14_DasPostExp_1 | 1.50 | 1.26 | 14.95 | 12.58 |
| 33 | PZ14_ChenPostExp_2 | 0.80 | 1.27 | 7.99 | 12.69 |
| 34 | PZ14_DasPostExp_3 | 1.47 | 1.28 | 14.74 | 12.81 |
| 35 | PZ14_BujnickiPostExp_5 | 1.21 | 1.33 | 12.12 | 13.30 |
| 36 | PZ14_ChenPostExp_7 | 1.18 | 1.37 | 11.81 | 13.65 |
| 37 | PZ14_ChenPostExp_1 | 1.83 | 1.37 | 18.34 | 13.71 |
| 38 | PZ14_DingPostExp_8 | 1.22 | 1.37 | 12.24 | 13.73 |
| 39 | PZ14_DasPostExp_9 | 1.73 | 1.39 | 17.25 | 13.85 |
| 40 | PZ14_DasPreExp_10 | 1.73 | 1.39 | 17.25 | 13.85 |
| 41 | PZ14_DingPostExp_2 | 1.25 | 1.39 | 12.49 | 13.88 |
| 42 | PZ14_DingPreExp_5 | 1.79 | 1.45 | 17.93 | 14.49 |
| 43 | PZ14_DingPostExp_3 | 1.55 | 1.45 | 15.54 | 14.51 |
| 44 | PZ14_DingPostExp_6 | 1.84 | 1.47 | 18.40 | 14.73 |
| 45 | PZ14_DingPostExp_9 | 1.35 | 1.48 | 13.46 | 14.76 |
| 46 | PZ14_DingPreExp_3 | 1.38 | 1.48 | 13.85 | 14.80 |
| 47 | PZ14_DingPostExp_1 | 1.15 | 1.48 | 11.52 | 14.84 |
| 48 | PZ14_BujnickiPostExp_4 | 1.33 | 1.52 | 13.35 | 15.20 |
| 49 | PZ14_DingPreExp_1 | 1.45 | 1.52 | 14.54 | 15.20 |
| 50 | PZ14_DingPreExp_6 | 1.65 | 1.53 | 16.55 | 15.29 |
| 51 | PZ14_DingPreExp_8 | 1.68 | 1.54 | 16.75 | 15.35 |
| 52 | PZ14_BujnickiPostExp_2 | 1.42 | 1.54 | 14.20 | 15.41 |
| 53 | PZ14_DingPostExp_4 | 1.68 | 1.56 | 16.79 | 15.59 |
| 54 | PZ14_DingPreExp_4 | 1.57 | 1.56 | 15.70 | 15.60 |
| 55 | PZ14_DingPreExp_7 | 1.54 | 1.56 | 15.38 | 15.61 |
| 56 | PZ14_BujnickiPostExp_3 | 1.64 | 1.57 | 16.37 | 15.65 |

| | | | | | |
|---|---|---|---|---|---|
| 57 | PZ14_DingPreExp_2 | 1.37 | 1.57 | 13.75 | 15.73 |
| 58 | PZ14_DingPostExp_10 | 1.64 | 1.59 | 16.40 | 15.86 |
| 59 | PZ14_DingPostExp_7 | 1.63 | 1.59 | 16.33 | 15.89 |
| 60 | PZ14_DingPostExp_5 | 1.37 | 1.59 | 13.70 | 15.92 |
| 61 | PZ14_BujnickiPreExp_7 | 1.58 | 1.61 | 15.76 | 16.08 |
| | Average | 1.37 | 1.31 | 13.66 | 13.08 |
| | Standard deviation | 0.61 | 1.01 | 6.09 | 10.05 |
| | Median | 1.84 | 1.61 | 18.40 | 16.08 |

**Table S39**. BARNABA report on structures in Puzzle 14b.

| No. | 3D RNA model | BBRMSD [nm] | ERMSD [nm] | BBRMSD [Å] | ERMSD [Å] |
|---|---|---|---|---|---|
| 1 | PZ14_BujnickiPreExp_1 | 1.34 | 0.72 | 13.41 | 7.17 |
| 2 | PZ14_DasPostExp_6 | 0.74 | 0.82 | 7.38 | 8.24 |
| 3 | PZ14_DasPreExp_1 | 0.74 | 0.82 | 7.38 | 8.24 |
| 4 | PZ14_BujnickiPreExp_3 | 1.26 | 0.87 | 12.62 | 8.70 |
| 5 | PZ14_BujnickiPreExp_2 | 1.80 | 0.87 | 17.95 | 8.72 |
| 6 | PZ14_DasPostExp_3 | 0.73 | 0.89 | 7.34 | 8.89 |
| 7 | PZ14_ChenPostExp_3 | 2.37 | 0.90 | 23.69 | 8.97 |
| 8 | PZ14_DasPreExp_9 | 0.89 | 0.91 | 8.87 | 9.11 |
| 9 | PZ14_ChenPostExp_8 | 1.93 | 0.95 | 19.27 | 9.49 |
| 10 | PZ14_DasPostExp_2 | 0.74 | 0.95 | 7.40 | 9.51 |
| 11 | PZ14_DasPostExp_10 | 1.03 | 0.96 | 10.29 | 9.65 |
| 12 | PZ14_DasPreExp_5 | 1.03 | 0.96 | 10.29 | 9.65 |
| 13 | PZ14_DasPostExp_4 | 1.24 | 0.97 | 12.35 | 9.67 |
| 14 | PZ14_DasPostExp_7 | 1.20 | 0.99 | 12.01 | 9.87 |
| 15 | PZ14_DasPreExp_2 | 1.20 | 0.99 | 12.01 | 9.87 |
| 16 | PZ14_DasPreExp_8 | 1.65 | 0.99 | 16.48 | 9.93 |
| 17 | PZ14_ChenPostExp_4 | 1.94 | 1.01 | 19.35 | 10.06 |
| 18 | PZ14_DasPostExp_5 | 0.91 | 1.02 | 9.13 | 10.22 |
| 19 | PZ14_DasPostExp_8 | 2.05 | 1.03 | 20.47 | 10.35 |
| 20 | PZ14_DasPreExp_4 | 2.05 | 1.03 | 20.47 | 10.35 |
| 21 | PZ14_ChenPostExp_10 | 1.61 | 1.04 | 16.12 | 10.40 |
| 22 | PZ14_DasPreExp_3 | 1.11 | 1.05 | 11.11 | 10.46 |
| 23 | PZ14_ChenPostExp_5 | 1.08 | 1.06 | 10.84 | 10.55 |
| 24 | PZ14_DasPostExp_1 | 1.41 | 1.06 | 14.05 | 10.56 |
| 25 | PZ14_DasPreExp_6 | 1.41 | 1.06 | 14.05 | 10.56 |
| 26 | PZ14_ChenPostExp_6 | 1.18 | 1.09 | 11.76 | 10.92 |
| 27 | PZ14_DasPreExp_7 | 1.77 | 1.10 | 17.69 | 10.97 |
| 28 | PZ14_ChenPostExp_9 | 1.96 | 1.10 | 19.63 | 11.03 |
| 29 | PZ14_BujnickiPostExp_2 | 1.58 | 1.14 | 15.81 | 11.35 |
| 30 | PZ14_ChenPostExp_2 | 0.98 | 1.14 | 9.78 | 11.38 |
| 31 | PZ14_ChenPostExp_7 | 1.39 | 1.15 | 13.87 | 11.45 |
| 32 | PZ14_BujnickiPostExp_4 | 1.87 | 1.19 | 18.75 | 11.88 |
| 33 | PZ14_BujnickiPostExp_1 | 1.38 | 1.19 | 13.76 | 11.93 |
| 34 | PZ14_DasPostExp_9 | 1.25 | 1.19 | 12.52 | 11.93 |
| 35 | PZ14_DasPreExp_10 | 1.25 | 1.19 | 12.52 | 11.93 |
| 36 | PZ14_ChenPostExp_1 | 1.79 | 1.23 | 17.91 | 12.31 |
| 37 | PZ14_DingPostExp_8 | 0.70 | 1.31 | 6.99 | 13.11 |
| 38 | PZ14_DingPostExp_2 | 1.88 | 1.32 | 18.76 | 13.17 |

| | | | | | |
|---|---|---|---|---|---|
| 39 | PZ14_DingPostExp_1 | 1.69 | 1.32 | 16.94 | 13.25 |
| 40 | PZ14_DingPreExp_4 | 0.92 | 1.33 | 9.17 | 13.33 |
| 41 | PZ14_DingPostExp_5 | 1.76 | 1.37 | 17.60 | 13.72 |
| 42 | PZ14_DingPostExp_3 | 1.29 | 1.37 | 12.90 | 13.73 |
| 43 | PZ14_DingPostExp_10 | 1.12 | 1.38 | 11.19 | 13.82 |
| 44 | PZ14_BujnickiPreExp_4 | 1.41 | 1.40 | 14.11 | 13.96 |
| 45 | PZ14_DingPreExp_1 | 0.96 | 1.41 | 9.59 | 14.11 |
| 46 | PZ14_DingPostExp_7 | 1.75 | 1.43 | 17.48 | 14.26 |
| 47 | PZ14_DingPostExp_9 | 0.83 | 1.46 | 8.33 | 14.62 |
| 48 | PZ14_BujnickiPostExp_3 | 1.63 | 1.47 | 16.31 | 14.74 |
| 49 | PZ14_DingPostExp_4 | 0.88 | 1.48 | 8.79 | 14.77 |
| 50 | PZ14_DingPreExp_2 | 1.89 | 1.48 | 18.94 | 14.80 |
| 51 | PZ14_DingPreExp_3 | 2.09 | 1.52 | 20.90 | 15.15 |
| 52 | PZ14_DingPostExp_6 | 0.83 | 1.54 | 8.29 | 15.39 |
| | Average | 1.37 | 1.14 | 13.74 | 11.39 |
| | Standard deviation | 0.70 | 0.72 | 6.99 | 7.17 |
| | Median | 1.84 | 1.61 | 18.40 | 16.08 |

**Table S40**. BARNABA report on structures in Puzzle 15.

| No. | 3D RNA model | BBRMSD [nm] | ERMSD [nm] | BBRMSD [Å] | ERMSD [Å] |
|---|---|---|---|---|---|
| 1 | PZ15_Adamiak_2 | 0.85 | 1.04 | 8.52 | 10.44 |
| 2 | PZ15_RNAComposer1_1 | 0.85 | 1.04 | 8.52 | 10.44 |
| 3 | PZ15_Adamiak_3 | 0.88 | 1.05 | 8.82 | 10.51 |
| 4 | PZ15_RNAComposer1_2 | 0.88 | 1.05 | 8.82 | 10.51 |
| 5 | PZ15_Chen_3 | 0.92 | 1.08 | 9.22 | 10.76 |
| 6 | PZ15_Adamiak_6 | 0.92 | 1.28 | 9.25 | 12.85 |
| 7 | PZ15_Adamiak_10 | 0.71 | 1.29 | 7.06 | 12.86 |
| 8 | PZ15_Adamiak_7 | 0.87 | 1.29 | 8.72 | 12.87 |
| 9 | PZ15_Adamiak_8 | 0.69 | 1.30 | 6.88 | 13.01 |
| 10 | PZ15_Adamiak_4 | 0.91 | 1.35 | 9.07 | 13.46 |
| 11 | PZ15_RNAComposer2_1 | 0.91 | 1.35 | 9.07 | 13.46 |
| 12 | PZ15_Adamiak_9 | 0.82 | 1.36 | 8.25 | 13.56 |
| 13 | PZ15_Chen_1 | 1.00 | 1.36 | 9.99 | 13.62 |
| 14 | PZ15_Chen_4 | 1.09 | 1.38 | 10.86 | 13.78 |
| 15 | PZ15_Adamiak_5 | 0.90 | 1.38 | 8.97 | 13.82 |
| 16 | PZ15_RNAComposer2_2 | 0.90 | 1.38 | 8.97 | 13.82 |
| 17 | PZ15_Adamiak_1 | 0.94 | 1.38 | 9.42 | 13.83 |
| 18 | PZ15_FARFAR2_1 | 1.28 | 1.40 | 12.77 | 13.98 |
| 19 | PZ15_FARFAR2_10 | 1.50 | 1.40 | 15.04 | 13.99 |
| 20 | PZ15_Chen_2 | 1.66 | 1.43 | 16.57 | 14.28 |
| 21 | PZ15_FARFAR2_9 | 1.56 | 1.43 | 15.64 | 14.34 |
| 22 | PZ15_SimRNA2_2 | 1.94 | 1.45 | 19.38 | 14.46 |
| 23 | PZ15_Chen_5 | 1.70 | 1.46 | 16.95 | 14.58 |
| 24 | PZ15_Chen_6 | 0.89 | 1.46 | 8.94 | 14.59 |
| 25 | PZ15_FARFAR2_6 | 1.84 | 1.46 | 18.41 | 14.61 |
| 26 | PZ15_FARFAR1_5 | 1.44 | 1.46 | 14.40 | 14.61 |
| 27 | PZ15_FARFAR1_1 | 1.92 | 1.47 | 19.16 | 14.67 |
| 28 | PZ15_SimRNA2_1 | 1.58 | 1.47 | 15.79 | 14.71 |
| 29 | PZ15_SimRNA2_4 | 1.22 | 1.47 | 12.25 | 14.74 |

| | | | | | |
|---|---|---|---|---|---|
| 30 | PZ15_Chen_9 | 1.45 | 1.48 | 14.48 | 14.75 |
| 31 | PZ15_3dRNA2_10 | 2.55 | 1.48 | 25.49 | 14.85 |
| 32 | PZ15_3dRNA2_1 | 2.55 | 1.49 | 25.49 | 14.86 |
| 33 | PZ15_3dRNA2_3 | 2.54 | 1.49 | 25.45 | 14.86 |
| 34 | PZ15_3dRNA2_8 | 2.56 | 1.49 | 25.56 | 14.86 |
| 35 | PZ15_3dRNA2_9 | 2.54 | 1.49 | 25.44 | 14.88 |
| 36 | PZ15_3dRNA2_7 | 2.54 | 1.49 | 25.42 | 14.89 |
| 37 | PZ15_3dRNA2_5 | 2.54 | 1.49 | 25.43 | 14.89 |
| 38 | PZ15_3dRNA2_4 | 2.54 | 1.49 | 25.42 | 14.90 |
| 39 | PZ15_3dRNA2_2 | 2.54 | 1.49 | 25.41 | 14.91 |
| 40 | PZ15_FARFAR1_3 | 1.65 | 1.49 | 16.53 | 14.94 |
| 41 | PZ15_FARFAR1_10 | 1.94 | 1.50 | 19.39 | 14.98 |
| 42 | PZ15_3dRNA2_6 | 2.55 | 1.50 | 25.49 | 14.98 |
| 43 | PZ15_FARFAR1_9 | 1.61 | 1.50 | 16.11 | 15.04 |
| 44 | PZ15_FARFAR1_7 | 1.64 | 1.52 | 16.41 | 15.15 |
| 45 | PZ15_FARFAR1_4 | 1.60 | 1.52 | 16.02 | 15.21 |
| 46 | PZ15_SimRNA2_10 | 2.02 | 1.53 | 20.20 | 15.29 |
| 47 | PZ15_FARFAR2_5 | 1.12 | 1.53 | 11.24 | 15.30 |
| 48 | PZ15_FARFAR1_6 | 1.65 | 1.54 | 16.47 | 15.37 |
| 49 | PZ15_FARFAR1_8 | 2.13 | 1.54 | 21.33 | 15.39 |
| 50 | PZ15_SimRNA2_8 | 2.32 | 1.54 | 23.21 | 15.44 |
| 51 | PZ15_FARFAR1_2 | 1.70 | 1.55 | 17.03 | 15.45 |
| 52 | PZ15_SimRNA2_9 | 1.51 | 1.55 | 15.10 | 15.49 |
| 53 | PZ15_FARFAR2_3 | 1.69 | 1.55 | 16.93 | 15.53 |
| 54 | PZ15_FARFAR2_2 | 1.71 | 1.56 | 17.09 | 15.56 |
| 55 | PZ15_SimRNA2_3 | 2.03 | 1.56 | 20.33 | 15.59 |
| 56 | PZ15_SimRNA2_6 | 2.12 | 1.56 | 21.18 | 15.63 |
| 57 | PZ15_FARFAR2_8 | 1.45 | 1.57 | 14.55 | 15.70 |
| 58 | PZ15_SimRNA2_5 | 2.05 | 1.57 | 20.47 | 15.71 |
| 59 | PZ15_Chen_7 | 1.22 | 1.57 | 12.19 | 15.73 |
| 60 | PZ15_Chen_8 | 1.27 | 1.58 | 12.73 | 15.78 |
| 61 | PZ15_SimRNA1_10 | 2.04 | 1.58 | 20.43 | 15.80 |
| 62 | PZ15_SimRNA1_1 | 0.73 | 1.58 | 7.33 | 15.82 |
| 63 | PZ15_FARFAR2_4 | 1.05 | 1.59 | 10.52 | 15.91 |
| 64 | PZ15_Chen_10 | 1.00 | 1.60 | 9.99 | 15.97 |
| 65 | PZ15_SimRNA1_7 | 1.87 | 1.60 | 18.74 | 15.99 |
| 66 | PZ15_FARFAR2_7 | 1.40 | 1.63 | 14.02 | 16.27 |
| 67 | PZ15_SimRNA1_9 | 1.98 | 1.64 | 19.84 | 16.39 |
| 68 | PZ15_SimRNA1_8 | 2.09 | 1.64 | 20.89 | 16.40 |
| 69 | PZ15_SimRNA2_7 | 0.98 | 1.67 | 9.77 | 16.65 |
| 70 | PZ15_SimRNA1_5 | 2.17 | 1.79 | 21.74 | 17.93 |
| 71 | PZ15_SimRNA1_2 | 2.18 | 1.83 | 21.75 | 18.35 |
| 72 | PZ15_SimRNA1_4 | 2.47 | 2.04 | 24.75 | 20.39 |
| 73 | PZ15_SimRNA1_6 | 2.27 | 2.09 | 22.66 | 20.88 |
| 74 | PZ15_SimRNA1_3 | 1.76 | 2.14 | 17.63 | 21.42 |
| | Average | 1.61 | 1.49 | 16.07 | 14.91 |
| | Standard deviation | 0.69 | 1.04 | 6.88 | 10.44 |
| | Median | 1.84 | 1.61 | 18.40 | 16.08 |

**Table S41**. BARNABA report on structures in Puzzle 17.

| No. | 3D RNA model | BBRMSD [nm] | ERMSD [nm] | BBRMSD [Å] | ERMSD [Å] |
|---|---|---|---|---|---|
| 1 | PZ17_Das_1 | 0.90 | 1.36 | 8.95 | 13.61 |
| 2 | PZ17_Das_8 | 0.92 | 1.51 | 9.19 | 15.09 |
| 3 | PZ17_Bujnicki_8 | 1.31 | 1.52 | 13.10 | 15.20 |
| 4 | PZ17_Bujnicki_7 | 1.15 | 1.53 | 11.53 | 15.34 |
| 5 | PZ17_DasExtraInfo_2 | 1.31 | 1.58 | 13.07 | 15.84 |
| 6 | PZ17_DasExtraInfo_3 | 1.40 | 1.58 | 13.98 | 15.85 |
| 7 | PZ17_DasExtraInfo_1 | 1.17 | 1.59 | 11.72 | 15.86 |
| 8 | PZ17_Das_7 | 1.14 | 1.59 | 11.38 | 15.89 |
| 9 | PZ17_Bujnicki_6 | 1.55 | 1.60 | 15.50 | 15.96 |
| 10 | PZ17_Chen_2 | 1.24 | 1.60 | 12.37 | 16.04 |
| 11 | PZ17_Das_4 | 0.68 | 1.61 | 6.83 | 16.07 |
| 12 | PZ17_Das_3 | 1.52 | 1.61 | 15.16 | 16.08 |
| 13 | PZ17_Chen_8 | 1.75 | 1.61 | 17.49 | 16.11 |
| 14 | PZ17_Das_9 | 0.93 | 1.62 | 9.26 | 16.17 |
| 15 | PZ17_Das_5 | 1.62 | 1.62 | 16.20 | 16.18 |
| 16 | PZ17_Das_6 | 1.22 | 1.62 | 12.24 | 16.21 |
| 17 | PZ17_SimRNA2_1 | 0.52 | 1.62 | 5.23 | 16.23 |
| 18 | PZ17_Bujnicki_1 | 1.48 | 1.63 | 14.83 | 16.29 |
| 19 | PZ17_Bujnicki_9 | 1.48 | 1.63 | 14.83 | 16.29 |
| 20 | PZ17_Das_2 | 1.15 | 1.65 | 11.54 | 16.45 |
| 21 | PZ17_Das_10 | 1.84 | 1.65 | 18.45 | 16.47 |
| 22 | PZ17_Bujnicki_5 | 1.49 | 1.65 | 14.92 | 16.50 |
| 23 | PZ17_SimRNA2_8 | 1.36 | 1.67 | 13.60 | 16.69 |
| 24 | PZ17_Chen_7 | 1.80 | 1.67 | 17.99 | 16.74 |
| 25 | PZ17_SimRNA2_3 | 1.66 | 1.68 | 16.64 | 16.78 |
| 26 | PZ17_Xiao_2 | 1.76 | 1.68 | 17.63 | 16.81 |
| 27 | PZ17_Ding_7 | 1.45 | 1.69 | 14.47 | 16.88 |
| 28 | PZ17_Dohkolyan_3 | 1.55 | 1.70 | 15.49 | 16.95 |
| 29 | PZ17_Adamiak_3 | 1.25 | 1.70 | 12.49 | 16.98 |
| 30 | PZ17_Bujnicki_4 | 1.71 | 1.70 | 17.08 | 17.02 |
| 31 | PZ17_Adamiak_1 | 1.28 | 1.71 | 12.77 | 17.07 |
| 32 | PZ17_Bujnicki_10 | 1.54 | 1.71 | 15.41 | 17.07 |
| 33 | PZ17_Bujnicki_2 | 1.54 | 1.71 | 15.41 | 17.07 |
| 34 | PZ17_Ding_8 | 1.61 | 1.71 | 16.06 | 17.11 |
| 35 | PZ17_Adamiak_2 | 1.22 | 1.71 | 12.19 | 17.13 |
| 36 | PZ17_RNAComposer2_8 | 1.40 | 1.72 | 13.98 | 17.22 |
| 37 | PZ17_Chen_3 | 1.83 | 1.74 | 18.33 | 17.39 |
| 38 | PZ17_Bujnicki_3 | 1.56 | 1.74 | 15.55 | 17.40 |
| 39 | PZ17_Xiao_3 | 1.64 | 1.74 | 16.40 | 17.43 |
| 40 | PZ17_Ding_10 | 1.49 | 1.75 | 14.92 | 17.48 |
| 41 | PZ17_SimRNA2_7 | 0.81 | 1.75 | 8.07 | 17.48 |
| 42 | PZ17_RNAComposer2_7 | 1.31 | 1.75 | 13.14 | 17.50 |
| 43 | PZ17_Xiao_9 | 1.44 | 1.76 | 14.42 | 17.56 |
| 44 | PZ17_Ding_9 | 1.75 | 1.76 | 17.52 | 17.58 |
| 45 | PZ17_SimRNA2_2 | 1.67 | 1.77 | 16.71 | 17.68 |
| 46 | PZ17_Xiao_5 | 1.54 | 1.77 | 15.40 | 17.69 |
| 47 | PZ17_Xiao_8 | 1.66 | 1.77 | 16.58 | 17.72 |
| 48 | PZ17_SimRNA2_4 | 1.59 | 1.77 | 15.93 | 17.73 |

| | | | | | |
|---|---|---|---|---|---|
| 49 | PZ17_RNAComposer2_2 | 1.27 | 1.78 | 12.67 | 17.75 |
| 50 | PZ17_RNAComposer2_4 | 1.41 | 1.78 | 14.13 | 17.78 |
| 51 | PZ17_Chen_1 | 1.75 | 1.78 | 17.45 | 17.84 |
| 52 | PZ17_Xiao_1 | 1.54 | 1.79 | 15.35 | 17.85 |
| 53 | PZ17_RNAComposer2_1 | 1.29 | 1.79 | 12.95 | 17.86 |
| 54 | PZ17_RNAComposer2_6 | 1.34 | 1.79 | 13.39 | 17.94 |
| 55 | PZ17_Ding_6 | 1.80 | 1.80 | 17.99 | 17.96 |
| 56 | PZ17_SimRNA2_6 | 1.24 | 1.80 | 12.39 | 17.97 |
| 57 | PZ17_SimRNA2_5 | 1.93 | 1.81 | 19.30 | 18.09 |
| 58 | PZ17_Chen_6 | 1.56 | 1.81 | 15.57 | 18.13 |
| 59 | PZ17_RNAComposer2_5 | 1.32 | 1.81 | 13.19 | 18.14 |
| 60 | PZ17_Chen_9 | 1.65 | 1.82 | 16.50 | 18.18 |
| 61 | PZ17_Xiao_10 | 1.62 | 1.82 | 16.17 | 18.19 |
| 62 | PZ17_Ding_5 | 1.69 | 1.82 | 16.94 | 18.24 |
| 63 | PZ17_Ding_3 | 1.53 | 1.84 | 15.27 | 18.37 |
| 64 | PZ17_Xiao_6 | 1.78 | 1.84 | 17.78 | 18.43 |
| 65 | PZ17_SimRNA2_9 | 1.50 | 1.85 | 14.97 | 18.54 |
| 66 | PZ17_Ding_1 | 1.59 | 1.87 | 15.90 | 18.67 |
| 67 | PZ17_RNAComposer2_10 | 1.63 | 1.87 | 16.33 | 18.70 |
| 68 | PZ17_Xiao_4 | 1.83 | 1.87 | 18.26 | 18.70 |
| 69 | PZ17_RNAComposer2_3 | 1.25 | 1.90 | 12.46 | 18.97 |
| 70 | PZ17_RNAComposer2_9 | 1.68 | 1.90 | 16.82 | 18.97 |
| 71 | PZ17_Ding_2 | 1.70 | 1.92 | 17.02 | 19.18 |
| 72 | PZ17_Ding_4 | 1.67 | 1.92 | 16.72 | 19.23 |
| 73 | PZ17_Major_5 | 1.98 | 1.93 | 19.80 | 19.28 |
| 74 | PZ17_Major_3 | 1.86 | 1.93 | 18.55 | 19.30 |
| 75 | PZ17_Major_1 | 1.79 | 1.93 | 17.91 | 19.34 |
| 76 | PZ17_Dohkolyan_2 | 1.73 | 1.94 | 17.32 | 19.45 |
| 77 | PZ17_Xiao_7 | 1.41 | 1.96 | 14.15 | 19.57 |
| 78 | PZ17_RNAComposer1_3 | 1.79 | 1.96 | 17.92 | 19.62 |
| 79 | PZ17_Major_2 | 1.93 | 1.96 | 19.32 | 19.63 |
| 80 | PZ17_Major_4 | 1.95 | 1.96 | 19.52 | 19.64 |
| 81 | PZ17_RNAComposer1_1 | 1.96 | 1.98 | 19.61 | 19.77 |
| 82 | PZ17_RNAComposer1_4 | 2.05 | 2.01 | 20.48 | 20.07 |
| 83 | PZ17_RNAComposer1_6 | 1.93 | 2.01 | 19.29 | 20.09 |
| 84 | PZ17_RNAComposer1_5 | 2.03 | 2.01 | 20.28 | 20.10 |
| 85 | PZ17_RNAComposer1_8 | 1.79 | 2.01 | 17.92 | 20.14 |
| 86 | PZ17_SimRNA1_6 | 1.58 | 2.02 | 15.81 | 20.18 |
| 87 | PZ17_SimRNA1_8 | 1.73 | 2.02 | 17.27 | 20.19 |
| 88 | PZ17_SimRNA1_2 | 1.72 | 2.02 | 17.21 | 20.20 |
| 89 | PZ17_RNAComposer1_9 | 2.00 | 2.03 | 19.97 | 20.28 |
| 90 | PZ17_SimRNA1_5 | 1.77 | 2.04 | 17.66 | 20.39 |
| 91 | PZ17_Major_7 | 1.72 | 2.04 | 17.23 | 20.45 |
| 92 | PZ17_Major_6 | 1.64 | 2.05 | 16.43 | 20.53 |
| 93 | PZ17_RNAComposer1_10 | 1.89 | 2.06 | 18.91 | 20.63 |
| 94 | PZ17_Major_8 | 1.76 | 2.06 | 17.63 | 20.65 |
| 95 | PZ17_Chen_5 | 1.01 | 2.07 | 10.11 | 20.74 |
| 96 | PZ17_SimRNA1_4 | 1.73 | 2.09 | 17.33 | 20.92 |
| 97 | PZ17_SimRNA1_3 | 1.77 | 2.11 | 17.74 | 21.09 |
| 98 | PZ17_Major_9 | 1.75 | 2.12 | 17.45 | 21.18 |
| 99 | PZ17_SimRNA1_1 | 1.61 | 2.12 | 16.10 | 21.21 |

| | | | | | |
|---|---|---|---|---|---|
| 100 | PZ17_Chen_4 | 1.56 | 2.15 | 15.62 | 21.53 |
| 101 | PZ17_Major_10 | 1.42 | 2.21 | 14.22 | 22.08 |
| 102 | PZ17_RNAComposer1_2 | 2.24 | 2.22 | 22.38 | 22.24 |
| 103 | PZ17_RNAComposer1_7 | 2.13 | 2.23 | 21.25 | 22.31 |
| 104 | PZ17_Dohkolyan_1 | 2.23 | 2.24 | 22.30 | 22.41 |
| 105 | PZ17_Chen_10 | 2.00 | 2.26 | 20.03 | 22.64 |
| 106 | PZ17_SimRNA1_7 | 1.96 | 2.33 | 19.57 | 23.28 |
| 107 | PZ17_SimRNA1_9 | 2.10 | 2.37 | 21.04 | 23.65 |
| | Average | 1.57 | 1.83 | 15.73 | 18.34 |
| | Standard deviation | 0.52 | 1.36 | 5.23 | 13.61 |
| | Median | 2.37 | 1.54 | 23.69 | 15.39 |

**Table S42**. BARNABA report on structures in Puzzle 18.

| No. | 3D RNA model | BBRMSD [nm] | ERMSD [nm] | BBRMSD [Å] | ERMSD [Å] |
|---|---|---|---|---|---|
| 1 | PZ18_Das_1 | 0.55 | 0.78 | 5.52 | 7.85 |
| 2 | PZ18_Das_3 | 0.57 | 0.81 | 5.69 | 8.14 |
| 3 | PZ18_Das_2 | 0.40 | 0.88 | 3.95 | 8.79 |
| 4 | PZ18_Das_4 | 0.56 | 0.94 | 5.63 | 9.35 |
| 5 | PZ18_Chen_1 | 0.35 | 0.98 | 3.53 | 9.80 |
| 6 | PZ18_Chen_2 | 0.69 | 0.99 | 6.91 | 9.89 |
| 7 | PZ18_Das_5 | 1.10 | 1.10 | 11.05 | 10.98 |
| 8 | PZ18_FARFAR_7 | 1.53 | 1.43 | 15.33 | 14.32 |
| 9 | PZ18_Chen_3 | 0.89 | 1.44 | 8.87 | 14.42 |
| 10 | PZ18_Chen_4 | 0.89 | 1.45 | 8.90 | 14.45 |
| 11 | PZ18_FARFAR_1 | 1.65 | 1.45 | 16.55 | 14.50 |
| 12 | PZ18_3dRNA_2 | 1.94 | 1.46 | 19.35 | 14.56 |
| 13 | PZ18_Dokholyan_2 | 1.19 | 1.46 | 11.93 | 14.58 |
| 14 | PZ18_FARFAR_4 | 1.18 | 1.46 | 11.76 | 14.62 |
| 15 | PZ18_FARFAR_2 | 0.97 | 1.46 | 9.74 | 14.64 |
| 16 | PZ18_Dokholyan_1 | 0.87 | 1.48 | 8.66 | 14.76 |
| 17 | PZ18_FARFAR_10 | 1.34 | 1.48 | 13.35 | 14.80 |
| 18 | PZ18_FARFAR_3 | 1.27 | 1.48 | 12.68 | 14.84 |
| 19 | PZ18_FARFAR_8 | 1.58 | 1.48 | 15.76 | 14.85 |
| 20 | PZ18_FARFAR_9 | 1.78 | 1.50 | 17.78 | 14.96 |
| 21 | PZ18_Chen_5 | 1.58 | 1.50 | 15.80 | 15.03 |
| 22 | PZ18_FARFAR_6 | 1.01 | 1.50 | 10.10 | 15.04 |
| 23 | PZ18_FARFAR_5 | 1.33 | 1.52 | 13.34 | 15.22 |
| 24 | PZ18_RNAComposer_2 | 2.16 | 1.52 | 21.56 | 15.24 |
| 25 | PZ18_Ding_2 | 2.08 | 1.53 | 20.80 | 15.34 |
| 26 | PZ18_Dokholyan_3 | 0.77 | 1.54 | 7.72 | 15.40 |
| 27 | PZ18_3dRNA_1 | 1.91 | 1.54 | 19.13 | 15.42 |
| 28 | PZ18_RNAComposer_5 | 1.55 | 1.55 | 15.46 | 15.51 |
| 29 | PZ18_3dRNA_4 | 1.24 | 1.57 | 12.40 | 15.73 |
| 30 | PZ18_RNAComposer_4 | 2.13 | 1.57 | 21.32 | 15.74 |
| 31 | PZ18_Ding_5 | 1.62 | 1.58 | 16.19 | 15.81 |
| 32 | PZ18_RNAComposer_1 | 2.11 | 1.58 | 21.14 | 15.82 |
| 33 | PZ18_YagoubAli_1 | 1.90 | 1.58 | 19.04 | 15.83 |
| 34 | PZ18_Lee_2 | 2.16 | 1.60 | 21.64 | 16.04 |
| 35 | PZ18_3dRNA_3 | 2.03 | 1.61 | 20.28 | 16.06 |

| | | | | | |
|---|---|---|---|---|---|
| 36 | PZ18_Ding_1 | 1.99 | 1.61 | 19.89 | 16.10 |
| 37 | PZ18_SimRNA_2 | 1.10 | 1.63 | 11.01 | 16.32 |
| 38 | PZ18_LeeServer_2 | 2.30 | 1.64 | 23.02 | 16.38 |
| 39 | PZ18_3dRNA_5 | 2.41 | 1.67 | 24.05 | 16.68 |
| 40 | PZ18_Ding_3 | 1.64 | 1.69 | 16.40 | 16.90 |
| 41 | PZ18_LeeServer_5 | 1.88 | 1.69 | 18.83 | 16.93 |
| 42 | PZ18_RNAComposer_3 | 2.10 | 1.70 | 21.02 | 16.97 |
| 43 | PZ18_Lee_5 | 2.44 | 1.71 | 24.38 | 17.12 |
| 44 | PZ18_Lee_1 | 2.23 | 1.73 | 22.34 | 17.30 |
| 45 | PZ18_LeeServer_1 | 2.26 | 1.75 | 22.63 | 17.45 |
| 46 | PZ18_Lee_4 | 2.25 | 1.76 | 22.52 | 17.57 |
| 47 | PZ18_LeeServer_4 | 1.96 | 1.76 | 19.56 | 17.58 |
| 48 | PZ18_SimRNA_3 | 2.48 | 1.81 | 24.77 | 18.12 |
| 49 | PZ18_Ding_4 | 2.49 | 1.98 | 24.94 | 19.83 |
| 50 | PZ18_SimRNA_1 | 2.30 | 2.04 | 23.03 | 20.44 |
| 51 | PZ18_Lee_3 | 2.15 | 2.05 | 21.46 | 20.52 |
| 52 | PZ18_LeeServer_3 | 2.12 | 2.10 | 21.20 | 21.00 |
| | Average | 1.60 | 1.52 | 15.96 | 15.22 |
| | Standard deviation | 0.35 | 0.78 | 3.53 | 7.85 |
| | Median | 2.49 | 2.10 | 24.94 | 21.00 |

**Table S43**. BARNABA report on structures in Puzzle 19.

| No. | 3D RNA model | BBRMSD [nm] | ERMSD [nm] | BBRMSD [Å] | ERMSD [Å] |
|---|---|---|---|---|---|
| 1 | PZ19_Chen_4 | 1.05 | 1.30 | 10.54 | 12.98 |
| 2 | PZ19_Bujnicki_3 | 1.60 | 1.32 | 16.05 | 13.15 |
| 3 | PZ19_SimRNA_2 | 1.53 | 1.32 | 15.30 | 13.21 |
| 4 | PZ19_Chen_1 | 0.58 | 1.35 | 5.81 | 13.52 |
| 5 | PZ19_Das_2 | 1.20 | 1.39 | 11.98 | 13.87 |
| 6 | PZ19_Das_4 | 1.53 | 1.39 | 15.26 | 13.87 |
| 7 | PZ19_RNAComposer_5 | 1.21 | 1.39 | 12.05 | 13.90 |
| 8 | PZ19_Das_3 | 1.52 | 1.40 | 15.17 | 14.02 |
| 9 | PZ19_Chen_3 | 1.59 | 1.41 | 15.85 | 14.11 |
| 10 | PZ19_RNAComposer_2 | 1.13 | 1.41 | 11.34 | 14.14 |
| 11 | PZ19_Bujnicki_1 | 1.51 | 1.42 | 15.08 | 14.21 |
| 12 | PZ19_3dRNA_5 | 1.62 | 1.44 | 16.22 | 14.39 |
| 13 | PZ19_3dRNA_2 | 1.47 | 1.45 | 14.66 | 14.48 |
| 14 | PZ19_Das_1 | 1.91 | 1.45 | 19.07 | 14.48 |
| 15 | PZ19_Das_5 | 1.17 | 1.45 | 11.72 | 14.53 |
| 16 | PZ19_Chen_2 | 0.76 | 1.45 | 7.63 | 14.54 |
| 17 | PZ19_Ding_3 | 1.84 | 1.47 | 18.45 | 14.67 |
| 18 | PZ19_Ding_1 | 1.61 | 1.47 | 16.06 | 14.69 |
| 19 | PZ19_RNAComposer_1 | 1.62 | 1.47 | 16.22 | 14.70 |
| 20 | PZ19_3dRNA_1 | 1.63 | 1.47 | 16.29 | 14.72 |
| 21 | PZ19_Dokholyan_1 | 1.77 | 1.48 | 17.71 | 14.77 |
| 22 | PZ19_Ding_2 | 1.72 | 1.49 | 17.20 | 14.89 |
| 23 | PZ19_3dRNA_3 | 1.66 | 1.50 | 16.64 | 15.02 |
| 24 | PZ19_LeeServer_1 | 1.53 | 1.51 | 15.35 | 15.14 |
| 25 | PZ19_Adamiak_1 | 1.17 | 1.52 | 11.67 | 15.16 |
| 26 | PZ19_RNAComposer_3 | 1.00 | 1.53 | 9.98 | 15.31 |

| | | | | | |
|---|---|---|---|---|---|
| 27 | PZ19_Bujnicki_2 | 1.08 | 1.54 | 10.80 | 15.37 |
| 28 | PZ19_Bujnicki_5 | 1.76 | 1.54 | 17.62 | 15.44 |
| 29 | PZ19_Ding_4 | 1.87 | 1.54 | 18.73 | 15.44 |
| 30 | PZ19_SimRNA_1 | 1.48 | 1.55 | 14.82 | 15.54 |
| 31 | PZ19_3dRNA_4 | 1.41 | 1.56 | 14.12 | 15.56 |
| 32 | PZ19_RNAComposer_4 | 1.30 | 1.58 | 13.00 | 15.79 |
| 33 | PZ19_Adamiak_4 | 1.31 | 1.59 | 13.05 | 15.92 |
| 34 | PZ19_LeeServer_3 | 2.02 | 1.59 | 20.15 | 15.93 |
| 35 | PZ19_Ding_5 | 2.13 | 1.60 | 21.26 | 16.01 |
| 36 | PZ19_Adamiak_5 | 1.02 | 1.60 | 10.22 | 16.04 |
| 37 | PZ19_LeeServer_2 | 1.16 | 1.60 | 11.62 | 16.04 |
| 38 | PZ19_LeeServer_5 | 1.61 | 1.62 | 16.08 | 16.16 |
| 39 | PZ19_Bujnicki_4 | 1.54 | 1.62 | 15.39 | 16.18 |
| 40 | PZ19_Adamiak_2 | 1.75 | 1.63 | 17.49 | 16.26 |
| 41 | PZ19_LeeServer_4 | 1.26 | 1.65 | 12.62 | 16.52 |
| 42 | PZ19_Adamiak_3 | 1.51 | 1.78 | 15.11 | 17.79 |
| 43 | PZ19_SimRNA_3 | 1.34 | 1.96 | 13.40 | 19.56 |
| 44 | PZ19_FARFAR_8 | 1.78 | 2.11 | 17.79 | 21.13 |
| 45 | PZ19_FARFAR_9 | 1.87 | 2.12 | 18.72 | 21.19 |
| 46 | PZ19_FARFAR_5 | 1.62 | 2.12 | 16.20 | 21.22 |
| 47 | PZ19_FARFAR_1 | 1.75 | 2.13 | 17.54 | 21.26 |
| 48 | PZ19_FARFAR_7 | 1.84 | 2.13 | 18.42 | 21.26 |
| 49 | PZ19_Chen_5 | 2.23 | 2.13 | 22.27 | 21.30 |
| 50 | PZ19_FARFAR_2 | 1.71 | 2.13 | 17.08 | 21.30 |
| 51 | PZ19_FARFAR_10 | 1.88 | 2.13 | 18.82 | 21.32 |
| 52 | PZ19_FARFAR_3 | 1.83 | 2.16 | 18.33 | 21.59 |
| 53 | PZ19_FARFAR_6 | 1.75 | 2.16 | 17.47 | 21.64 |
| 54 | PZ19_FARFAR_4 | 1.64 | 2.20 | 16.41 | 22.00 |
| | Average | 1.53 | 1.64 | 15.26 | 16.36 |
| | Standard deviation | 0.58 | 1.30 | 5.81 | 12.98 |
| | Median | 2.23 | 2.20 | 22.27 | 22.00 |

**Table S44**. BARNABA report on structures in Puzzle 20.

| No. | 3D RNA model | BBRMSD [nm] | ERMSD [nm] | BBRMSD [Å] | ERMSD [Å] |
|---|---|---|---|---|---|
| 1 | PZ20_Bujnicki_3 | 0.68 | 0.98 | 6.81 | 9.80 |
| 2 | PZ20_Bujnicki_2 | 0.71 | 0.98 | 7.14 | 9.81 |
| 3 | PZ20_Bujnicki_1 | 0.55 | 1.00 | 5.49 | 9.99 |
| 4 | PZ20_Adamiak_5 | 0.71 | 1.02 | 7.14 | 10.22 |
| 5 | PZ20_Bujnicki_4 | 0.47 | 1.03 | 4.74 | 10.35 |
| 6 | PZ20_RNAComposer_5 | 1.37 | 1.07 | 13.66 | 10.69 |
| 7 | PZ20_RNAComposer_4 | 1.19 | 1.12 | 11.91 | 11.18 |
| 8 | PZ20_Bujnicki_5 | 0.53 | 1.13 | 5.34 | 11.28 |
| 9 | PZ20_Das_3 | 0.72 | 1.16 | 7.15 | 11.60 |
| 10 | PZ20_RNAComposer_1 | 1.78 | 1.16 | 17.85 | 11.62 |
| 11 | PZ20_RNAComposer_2 | 1.73 | 1.18 | 17.34 | 11.79 |
| 12 | PZ20_RNAComposer_3 | 1.36 | 1.22 | 13.59 | 12.18 |
| 13 | PZ20_Adamiak_2 | 1.65 | 1.24 | 16.55 | 12.44 |
| 14 | PZ20_Adamiak_4 | 1.46 | 1.30 | 14.62 | 13.04 |
| 15 | PZ20_Das_5 | 1.70 | 1.31 | 17.01 | 13.05 |

| | | | | | |
|---|---|---|---|---|---|
| 16 | PZ20_FARFAR_6 | 2.01 | 1.31 | 20.06 | 13.08 |
| 17 | PZ20_Adamiak_1 | 1.23 | 1.31 | 12.25 | 13.09 |
| 18 | PZ20_FARFAR_3 | 1.74 | 1.32 | 17.42 | 13.21 |
| 19 | PZ20_FARFAR_5 | 1.99 | 1.32 | 19.92 | 13.22 |
| 20 | PZ20_Adamiak_3 | 1.79 | 1.32 | 17.88 | 13.25 |
| 21 | PZ20_FARFAR_1 | 1.86 | 1.35 | 18.56 | 13.55 |
| 22 | PZ20_FARFAR_9 | 1.73 | 1.36 | 17.35 | 13.58 |
| 23 | PZ20_FARFAR_8 | 1.72 | 1.36 | 17.18 | 13.64 |
| 24 | PZ20_FARFAR_7 | 1.77 | 1.37 | 17.69 | 13.68 |
| 25 | PZ20_FARFAR_10 | 1.79 | 1.38 | 17.93 | 13.84 |
| 26 | PZ20_Xiao3_3 | 1.96 | 1.39 | 19.56 | 13.87 |
| 27 | PZ20_FARFAR_4 | 1.78 | 1.39 | 17.84 | 13.88 |
| 28 | PZ20_Das_2 | 0.61 | 1.39 | 6.08 | 13.93 |
| 29 | PZ20_FARFAR_2 | 1.86 | 1.39 | 18.58 | 13.95 |
| 30 | PZ20_Xiao3_4 | 2.03 | 1.40 | 20.34 | 14.02 |
| 31 | PZ20_Das_4 | 1.00 | 1.42 | 9.97 | 14.24 |
| 32 | PZ20_Xiao3_5 | 1.28 | 1.43 | 12.76 | 14.28 |
| 33 | PZ20_Das_1 | 0.68 | 1.43 | 6.81 | 14.30 |
| 34 | PZ20_Xiao3_1 | 1.86 | 1.44 | 18.61 | 14.39 |
| 35 | PZ20_Xiao3_2 | 2.03 | 1.44 | 20.26 | 14.41 |
| 36 | PZ20_SimRNA_4 | 2.08 | 2.07 | 20.79 | 20.65 |
| 37 | PZ20_SimRNA_2 | 2.44 | 2.18 | 24.36 | 21.76 |
| 38 | PZ20_SimRNA_5 | 2.13 | 2.23 | 21.30 | 22.31 |
| 39 | PZ20_SimRNA_3 | 2.25 | 2.41 | 22.50 | 24.08 |
| 40 | PZ20_SimRNA_1 | 1.89 | 2.43 | 18.95 | 24.30 |
| | Average | 1.50 | 1.39 | 15.03 | 13.94 |
| | Standard deviation | 0.47 | 0.98 | 4.74 | 9.80 |
| | Median | 2.44 | 2.43 | 24.36 | 24.30 |

**Table S45**. BARNABA report on structures in Puzzle 21.

| **No.** | **3D RNA model** | **BBRMSD [nm]** | **ERMSD [nm]** | **BBRMSD [Å]** | **ERMSD [Å]** |
|---|---|---|---|---|---|
| 1 | PZ21_DasLORES_1 | 0.33 | 1.34 | 3.28 | 13.41 |
| 2 | PZ21_Das_5 | 0.64 | 1.61 | 6.35 | 16.12 |
| 3 | PZ21_Adamiak_4 | 0.45 | 1.65 | 4.50 | 16.52 |
| 4 | PZ21_Adamiak_5 | 0.54 | 1.69 | 5.36 | 16.90 |
| 5 | PZ21_Adamiak_3 | 0.44 | 1.70 | 4.38 | 16.98 |
| 6 | PZ21_ChenHighLig_2 | 0.38 | 1.70 | 3.78 | 17.03 |
| 7 | PZ21_ChenLowLig_2 | 0.38 | 1.70 | 3.78 | 17.03 |
| 8 | PZ21_DasLORES_5 | 0.92 | 1.71 | 9.15 | 17.12 |
| 9 | PZ21_Bujnicki_4 | 0.65 | 1.73 | 6.46 | 17.29 |
| 10 | PZ21_RNAComposer_4 | 1.46 | 1.73 | 14.55 | 17.34 |
| 11 | PZ21_RNAComposer_3 | 1.36 | 1.74 | 13.61 | 17.35 |
| 12 | PZ21_DasLORES_4 | 0.62 | 1.74 | 6.23 | 17.36 |
| 13 | PZ21_Adamiak_2 | 0.44 | 1.74 | 4.42 | 17.42 |
| 14 | PZ21_DasLORES_2 | 0.38 | 1.75 | 3.85 | 17.46 |
| 15 | PZ21_Adamiak_1 | 0.44 | 1.76 | 4.41 | 17.63 |
| 16 | PZ21_ChenHighLig_5 | 0.73 | 1.81 | 7.32 | 18.08 |
| 17 | PZ21_ChenLowLig_5 | 0.73 | 1.81 | 7.32 | 18.08 |
| 18 | PZ21_Bujnicki_2 | 0.98 | 1.82 | 9.81 | 18.17 |

| | | | | | |
|---|---|---|---|---|---|
| 19 | PZ21_ChenHighLig_1 | 0.42 | 1.84 | 4.16 | 18.41 |
| 20 | PZ21_ChenLowLig_1 | 0.42 | 1.84 | 4.16 | 18.41 |
| 21 | PZ21_Das_3 | 0.49 | 1.84 | 4.92 | 18.43 |
| 22 | PZ21_Das_4 | 0.55 | 1.84 | 5.54 | 18.44 |
| 23 | PZ21_DasLORES_3 | 1.53 | 1.88 | 15.34 | 18.78 |
| 24 | PZ21_Bujnicki_1 | 0.73 | 1.88 | 7.26 | 18.81 |
| 25 | PZ21_ChenHighLig_4 | 0.51 | 1.88 | 5.08 | 18.82 |
| 26 | PZ21_ChenLowLig_4 | 0.51 | 1.88 | 5.08 | 18.82 |
| 27 | PZ21_ChenHighLig_3 | 0.53 | 1.89 | 5.30 | 18.86 |
| 28 | PZ21_ChenLowLig_3 | 0.53 | 1.89 | 5.30 | 18.86 |
| 29 | PZ21_Das_1 | 0.57 | 1.89 | 5.71 | 18.91 |
| 30 | PZ21_Das_2 | 0.56 | 1.93 | 5.64 | 19.34 |
| 31 | PZ21_3dRNA_5 | 1.77 | 1.98 | 17.66 | 19.76 |
| 32 | PZ21_3dRNA_1 | 1.28 | 2.01 | 12.83 | 20.06 |
| 33 | PZ21_3dRNA_4 | 1.82 | 2.02 | 18.20 | 20.16 |
| 34 | PZ21_3dRNA_3 | 1.28 | 2.03 | 12.82 | 20.29 |
| 35 | PZ21_FARFAR_9 | 1.85 | 2.03 | 18.52 | 20.32 |
| 36 | PZ21_FARFAR_3 | 1.81 | 2.05 | 18.12 | 20.50 |
| 37 | PZ21_FARFAR_10 | 1.60 | 2.06 | 16.01 | 20.58 |
| 38 | PZ21_RNAComposer_2 | 1.85 | 2.07 | 18.48 | 20.67 |
| 39 | PZ21_FARFAR_6 | 1.67 | 2.07 | 16.66 | 20.72 |
| 40 | PZ21_FARFAR_4 | 1.87 | 2.09 | 18.73 | 20.85 |
| 41 | PZ21_Sanbonmatsu_1 | 1.52 | 2.09 | 15.24 | 20.89 |
| 42 | PZ21_Sanbonmatsu_2 | 1.52 | 2.09 | 15.24 | 20.89 |
| 43 | PZ21_RNAComposer_1 | 1.50 | 2.09 | 15.02 | 20.92 |
| 44 | PZ21_FARFAR_8 | 1.82 | 2.09 | 18.17 | 20.94 |
| 45 | PZ21_RNAComposer_5 | 1.76 | 2.11 | 17.56 | 21.06 |
| 46 | PZ21_FARFAR_7 | 1.86 | 2.12 | 18.63 | 21.17 |
| 47 | PZ21_FARFAR_2 | 1.66 | 2.12 | 16.64 | 21.20 |
| 48 | PZ21_FARFAR_1 | 1.84 | 2.13 | 18.35 | 21.32 |
| 49 | PZ21_3dRNA_2 | 1.32 | 2.16 | 13.23 | 21.63 |
| 50 | PZ21_FARFAR_5 | 1.86 | 2.17 | 18.64 | 21.67 |
| 51 | PZ21_SimRNA_4 | 1.16 | 2.18 | 11.58 | 21.76 |
| 52 | PZ21_Sanbonmatsu_3 | 1.48 | 2.18 | 14.81 | 21.84 |
| 53 | PZ21_Sanbonmatsu_4 | 1.48 | 2.18 | 14.81 | 21.84 |
| 54 | PZ21_SimRNA_2 | 1.26 | 2.19 | 12.59 | 21.94 |
| 55 | PZ21_Bujnicki_3 | 1.26 | 2.21 | 12.58 | 22.09 |
| 56 | PZ21_SimRNA_3 | 1.41 | 2.23 | 14.08 | 22.26 |
| 57 | PZ21_SimRNA_5 | 1.41 | 2.23 | 14.08 | 22.26 |
| 58 | PZ21_SimRNA_1 | 1.24 | 2.24 | 12.41 | 22.36 |
| 59 | PZ21_Bujnicki_5 | 2.19 | 2.38 | 21.93 | 23.76 |
| | Average | 1.11 | 1.95 | 11.11 | 19.45 |
| | Standard deviation | 0.33 | 1.34 | 3.28 | 13.41 |
| | Median | 2.19 | 2.38 | 21.93 | 23.76 |

**Table 45**. BARNABA report on structures in Puzzle 24.

| No. | 3D RNA model | BBRMSD [nm] | ERMSD [nm] | BBRMSD [Å] | ERMSD [Å] |
|---|---|---|---|---|---|
| 1 | PZ24_Bujnicki_4 | 1.90 | 0.89 | 18.95 | 8.87 |
| 2 | PZ24_Bujnicki_5 | 1.51 | 0.91 | 15.07 | 9.12 |

| | | | | | |
|---|---|---|---|---|---|
| 3 | PZ24_Vfold3D_1 | 1.54 | 0.92 | 15.43 | 9.18 |
| 4 | PZ24_FARFAR2_8 | 1.34 | 0.94 | 13.45 | 9.44 |
| 5 | PZ24_Bujnicki_2 | 1.03 | 0.94 | 10.27 | 9.44 |
| 6 | PZ24_FARFAR2_10 | 1.05 | 0.97 | 10.51 | 9.70 |
| 7 | PZ24_DasTFN_1 | 1.12 | 0.97 | 11.24 | 9.71 |
| 8 | PZ24_DasTFN_9 | 0.92 | 0.97 | 9.24 | 9.73 |
| 9 | PZ24_FARFAR2_6 | 0.92 | 0.97 | 9.24 | 9.73 |
| 10 | PZ24_Das_3 | 1.74 | 0.99 | 17.42 | 9.89 |
| 11 | PZ24_Das_2 | 1.77 | 0.99 | 17.72 | 9.90 |
| 12 | PZ24_Das_10 | 1.69 | 0.99 | 16.88 | 9.90 |
| 13 | PZ24_Das_6 | 1.49 | 0.99 | 14.94 | 9.90 |
| 14 | PZ24_Das_8 | 1.56 | 0.99 | 15.59 | 9.91 |
| 15 | PZ24_Das_5 | 1.48 | 0.99 | 14.82 | 9.92 |
| 16 | PZ24_Das_1 | 1.64 | 0.99 | 16.37 | 9.92 |
| 17 | PZ24_DasTFN_8 | 1.15 | 0.99 | 11.49 | 9.94 |
| 18 | PZ24_Das_4 | 1.55 | 0.99 | 15.55 | 9.94 |
| 19 | PZ24_Das_9 | 1.45 | 0.99 | 14.52 | 9.94 |
| 20 | PZ24_Das_7 | 1.41 | 1.00 | 14.07 | 9.97 |
| 21 | PZ24_FARFAR2_3 | 2.05 | 1.00 | 20.47 | 9.98 |
| 22 | PZ24_Vfold3D_3 | 1.65 | 1.00 | 16.54 | 9.99 |
| 23 | PZ24_FARFAR2_9 | 0.80 | 1.00 | 8.00 | 10.00 |
| 24 | PZ24_Bujnicki_3 | 1.14 | 1.00 | 11.35 | 10.01 |
| 25 | PZ24_FARFAR2_7 | 1.27 | 1.02 | 12.70 | 10.17 |
| 26 | PZ24_Vfold3D_5 | 1.17 | 1.02 | 11.70 | 10.24 |
| 27 | PZ24_DasTFN_7 | 1.72 | 1.02 | 17.21 | 10.24 |
| 28 | PZ24_DasTFN_6 | 0.70 | 1.03 | 6.99 | 10.28 |
| 29 | PZ24_Adamiak_4 | 1.54 | 1.04 | 15.39 | 10.43 |
| 30 | PZ24_DasTFN_10 | 0.49 | 1.04 | 4.88 | 10.44 |
| 31 | PZ24_RNAComposer_2 | 1.09 | 1.04 | 10.90 | 10.45 |
| 32 | PZ24_Vfold3D_2 | 1.33 | 1.05 | 13.32 | 10.45 |
| 33 | PZ24_FARFAR2_4 | 1.43 | 1.05 | 14.27 | 10.49 |
| 34 | PZ24_Bujnicki_1 | 1.55 | 1.05 | 15.49 | 10.51 |
| 35 | PZ24_DasTFN_3 | 1.75 | 1.06 | 17.46 | 10.58 |
| 36 | PZ24_FARFAR2_1 | 1.75 | 1.06 | 17.46 | 10.58 |
| 37 | PZ24_Vfold3D_4 | 1.49 | 1.06 | 14.93 | 10.63 |
| 38 | PZ24_Adamiak_5 | 1.02 | 1.07 | 10.16 | 10.67 |
| 39 | PZ24_RNAComposer_1 | 1.38 | 1.07 | 13.77 | 10.73 |
| 40 | PZ24_Adamiak_3 | 1.47 | 1.08 | 14.69 | 10.76 |
| 41 | PZ24_DasTFN_5 | 2.08 | 1.09 | 20.77 | 10.95 |
| 42 | PZ24_3dRNA_5 | 1.95 | 1.11 | 19.46 | 11.14 |
| 43 | PZ24_3dRNA_4 | 1.63 | 1.12 | 16.25 | 11.16 |
| 44 | PZ24_DasTFN_4 | 2.24 | 1.12 | 22.38 | 11.17 |
| 45 | PZ24_Adamiak_2 | 1.12 | 1.12 | 11.22 | 11.19 |
| 46 | PZ24_DasTFN_2 | 1.41 | 1.12 | 14.08 | 11.21 |
| 47 | PZ24_3dRNA_3 | 1.90 | 1.12 | 19.00 | 11.21 |
| 48 | PZ24_FARFAR2_5 | 2.01 | 1.12 | 20.06 | 11.24 |
| 49 | PZ24_Adamiak_1 | 1.07 | 1.13 | 10.71 | 11.27 |
| 50 | PZ24_3dRNA_2 | 2.09 | 1.13 | 20.86 | 11.32 |
| 51 | PZ24_3dRNA_1 | 1.66 | 1.15 | 16.55 | 11.45 |
| 52 | PZ24_RNAComposer_4 | 1.32 | 1.15 | 13.17 | 11.51 |
| 53 | PZ24_FARFAR2_2 | 1.79 | 1.16 | 17.88 | 11.55 |

| | | | | | |
|---|---|---|---|---|---|
| 54 | PZ24_iFoldRNA_1 | 1.98 | 1.16 | 19.82 | 11.64 |
| 55 | PZ24_VfoldLA_4 | 1.53 | 1.18 | 15.26 | 11.77 |
| 56 | PZ24_iFoldRNA_2 | 1.06 | 1.20 | 10.61 | 11.98 |
| 57 | PZ24_RNAComposer_3 | 1.34 | 1.22 | 13.38 | 12.19 |
| 58 | PZ24_RNAComposer_5 | 1.76 | 1.25 | 17.58 | 12.55 |
| 59 | PZ24_VfoldLA_1 | 1.56 | 1.32 | 15.57 | 13.16 |
| 60 | PZ24_VfoldLA_2 | 1.61 | 1.36 | 16.06 | 13.63 |
| 61 | PZ24_VfoldLA_3 | 1.52 | 1.37 | 15.21 | 13.72 |
| 62 | PZ24_VfoldLA_5 | 1.43 | 1.38 | 14.30 | 13.79 |
| 63 | PZ24_Ding_4 | 2.89 | 1.51 | 28.86 | 15.11 |
| 64 | PZ24_Ding_3 | 2.15 | 1.52 | 21.50 | 15.17 |
| 65 | PZ24_Ding_1 | 2.08 | 1.54 | 20.85 | 15.38 |
| 66 | PZ24_iFoldRNA_3 | 2.20 | 1.54 | 22.03 | 15.42 |
| 67 | PZ24_Ding_7 | 2.68 | 1.54 | 26.80 | 15.43 |
| 68 | PZ24_Ding_5 | 2.03 | 1.55 | 20.30 | 15.50 |
| 69 | PZ24_iFoldRNA_4 | 2.84 | 1.55 | 28.38 | 15.52 |
| 70 | PZ24_Ding_10 | 2.10 | 1.56 | 20.97 | 15.62 |
| 71 | PZ24_Ding_9 | 2.18 | 1.56 | 21.78 | 15.65 |
| 72 | PZ24_Ding_8 | 2.49 | 1.59 | 24.91 | 15.93 |
| 73 | PZ24_Ding_6 | 2.02 | 1.60 | 20.23 | 16.00 |
| 74 | PZ24_Ding_2 | 2.17 | 1.62 | 21.71 | 16.17 |
| 75 | PZ24_SimRNA_5 | 1.67 | 1.66 | 16.72 | 16.57 |
| 76 | PZ24_SimRNA_3 | 1.50 | 1.67 | 15.03 | 16.70 |
| 77 | PZ24_iFoldRNA_5 | 2.42 | 1.68 | 24.18 | 16.78 |
| 78 | PZ24_Kollmann_5 | 3.35 | 1.84 | 33.48 | 18.38 |
| 79 | PZ24_Kollmann_1 | 3.08 | 1.84 | 30.79 | 18.38 |
| 80 | PZ24_SimRNA_1 | 2.22 | 1.87 | 22.23 | 18.69 |
| 81 | PZ24_Kollmann_6 | 3.15 | 1.93 | 31.51 | 19.33 |
| 82 | PZ24_SimRNA_4 | 2.67 | 1.94 | 26.68 | 19.37 |
| 83 | PZ24_Kollmann_3 | 3.34 | 2.00 | 33.36 | 19.96 |
| 84 | PZ24_Kollmann_8 | 3.35 | 2.06 | 33.49 | 20.60 |
| 85 | PZ24_Kollmann_2 | 2.60 | 2.10 | 26.00 | 21.01 |
| 86 | PZ24_Kollmann_4 | 3.30 | 2.12 | 33.01 | 21.17 |
| 87 | PZ24_SimRNA_2 | 2.55 | 2.13 | 25.50 | 21.31 |
| 88 | PZ24_Kollmann_7 | 2.96 | 2.18 | 29.64 | 21.80 |
| 89 | PZ24_Kollmann_10 | 3.02 | 2.26 | 30.15 | 22.63 |
| 90 | PZ24_Kollmann_9 | 3.05 | 2.28 | 30.52 | 22.82 |
| | Average | 1.81 | 1.29 | 18.12 | 12.94 |
| | Standard deviation | 0.49 | 0.89 | 4.88 | 8.87 |
| | Median | 3.35 | 2.28 | 33.49 | 22.82 |

X3DNA reports

**Table S46**. X3DNA report on reference structures.

| | | | Base pairs | | Dinucleotide steps | | | | | |
|---|---|---|---|---|---|---|---|---|---|---|
| | | | | | **Right-handed** | | **Left-handed** | **Other** | | |
| **No.** | **3D RNA model** | **#Helices** | **#all bps** | **#canonical / wobble bps** | **#A-RNA** | **#B-RNA** | **#Z-RNA** | **#Unclassified steps** | **#Steps without a continuous backbone** | **Total** |
| 1 | PZ01_refstructure | 1 | 21 | 19 | 10 | 0 | 0 | 7 | 3 | 20 |
| 2 | PZ02_refstructure | 4 | 38 | 36 | 14 | 0 | 0 | 12 | 8 | 34 |
| 3 | PZ03_refstructure | 2 | 32 | 25 | 13 | 0 | 0 | 11 | 6 | 30 |
| 4 | PZ04_refstructure | 4 | 51 | 46 | 20 | 0 | 0 | 22 | 5 | 47 |
| 5 | PZ05_refstructure | 7 | 71 | 59 | 37 | 0 | 0 | 18 | 9 | 64 |
| 6 | PZ06_refstructure | 7 | 61 | 49 | 16 | 0 | 0 | 25 | 13 | 54 |
| 7 | PZ07_refstructure | 4 | 73 | 61 | 30 | 0 | 0 | 25 | 14 | 69 |
| 8 | PZ08_refstructure | 3 | 36 | 31 | 19 | 0 | 0 | 9 | 5 | 33 |
| 9 | PZ09_refstructure | 3 | 31 | 20 | 12 | 0 | 0 | 6 | 10 | 28 |
| 10 | PZ10_refstructure | 4 | 57 | 44 | 19 | 0 | 0 | 25 | 16 | 60 |
| 11 | PZ11_refstructure | 1 | 59 | 45 | 3 | 0 | 0 | 16 | 4 | 23 |
| 12 | PZ12_refstructure | 4 | 61 | 46 | 19 | 0 | 0 | 17 | 6 | 42 |
| 13 | PZ13_refstructure | 2 | 63 | 47 | 14 | 0 | 0 | 7 | 3 | 24 |
| 14 | PZ14_refstructure_bound | 2 | 65 | 48 | 10 | 0 | 0 | 7 | 3 | 20 |
| 15 | PZ14_refstructure_free | 2 | 67 | 49 | 9 | 0 | 0 | 8 | 2 | 19 |
| 16 | PZ15_refstructure | 2 | 69 | 50 | 13 | 0 | 0 | 2 | 4 | 19 |
| 17 | PZ17_refstructure | 2 | 71 | 51 | 8 | 0 | 0 | 9 | 4 | 21 |
| 18 | PZ18_refstructure | 2 | 73 | 52 | 12 | 0 | 0 | 9 | 6 | 27 |
| 19 | PZ19_refstructure | 2 | 75 | 53 | 12 | 0 | 0 | 6 | 7 | 25 |
| 20 | PZ20_refstructure | 2 | 78 | 54 | 17 | 0 | 0 | 4 | 7 | 28 |
| 21 | PZ21_refstructure | 2 | 80 | 56 | 6 | 0 | 0 | 2 | 3 | 11 |
| 22 | PZ24_refstructure | 3 | 82 | 57 | 32 | 0 | 0 | 11 | 3 | 46 |
| | Average | 2.95 | 84 | 58 | 15.68 | 0.00 | 0.00 | 11.73 | 6.41 | 33.82 |
| | Standard deviation | 1.62 | 86 | 59 | 8.35 | 0.00 | 0.00 | 7.36 | 3.87 | 16.27 |
| | Median | 2.00 | 88 | 60 | 13.50 | 0.00 | 0.00 | 9.00 | 5.50 | 28.00 |

**Table S47**. X3DNA report on structures in Puzzle 01.

| | | | Base pairs | | Dinucleotide steps | | | | | |
|---|---|---|---|---|---|---|---|---|---|---|
| | | | | | Right-handed | | Left-handed | Other | | |
| **No.** | **3D RNA model** | **#Helices** | **#all bps** | **#canonical / wobble bps** | **#A-RNA** | **#B-RNA** | **#Z-RNA** | **#Unclassified steps** | **#Steps without a continuous backbone** | **Total** |
| 1 | PZ01_Bujnicki_1 | 1 | 22 | 20 | 13 | 0 | 0 | 6 | 2 | 21 |
| 2 | PZ01_Bujnicki_2 | 1 | 21 | 18 | 7 | 0 | 0 | 10 | 3 | 20 |
| 3 | PZ01_Bujnicki_3 | 1 | 20 | 17 | 9 | 0 | 0 | 7 | 3 | 19 |
| 4 | PZ01_Bujnicki_4 | 1 | 19 | 13 | 9 | 0 | 0 | 4 | 5 | 18 |
| 5 | PZ01_Bujnicki_5 | 1 | 19 | 13 | 11 | 0 | 0 | 3 | 4 | 18 |
| 6 | PZ01_Chen_1 | 1 | 21 | 16 | 7 | 0 | 0 | 9 | 4 | 20 |
| 7 | PZ01_Das_1 | 1 | 22 | 20 | 11 | 0 | 0 | 8 | 2 | 21 |
| 8 | PZ01_Das_2 | 1 | 22 | 20 | 11 | 0 | 0 | 8 | 2 | 21 |
| 9 | PZ01_Das_3 | 1 | 22 | 20 | 11 | 0 | 0 | 8 | 2 | 21 |
| 10 | PZ01_Das_4 | 1 | 22 | 20 | 11 | 0 | 0 | 8 | 2 | 21 |
| 11 | PZ01_Das_5 | 1 | 21 | 20 | 11 | 0 | 0 | 6 | 3 | 20 |
| 12 | PZ01_Dokholyan_1 | 1 | 22 | 20 | 9 | 0 | 0 | 9 | 3 | 21 |
| 13 | PZ01_Major_1 | 1 | 21 | 18 | 5 | 0 | 0 | 12 | 3 | 20 |
| 14 | PZ01_Santalucia_1 | 1 | 21 | 20 | 14 | 0 | 0 | 3 | 3 | 20 |
| 15 | PZ01_refstructure | 1 | 21 | 19 | 10 | 0 | 0 | 7 | 3 | 20 |
| | Average | 1.00 | 21.07 | 18.27 | 9.93 | 0.00 | 0.00 | 7.20 | 2.93 | 20.07 |
| | Standard deviation | 0.00 | 1.03 | 2.49 | 2.34 | 0.00 | 0.00 | 2.51 | 0.88 | 1.03 |
| | Median | 1.00 | 21.00 | 20.00 | 11.00 | 0.00 | 0.00 | 8.00 | 3.00 | 20.00 |

**Table S49**. X3DNA report on structures in Puzzle 02.

| | | | Base pairs | | Dinucleotide steps | | | | | |
|---|---|---|---|---|---|---|---|---|---|---|
| | | | | | **Right-handed** | | **Left-handed** | **Other** | | |
| **No.** | **3D RNA model** | **#Helices** | **#all bps** | **#canonical / wobble bps** | **#A-RNA** | **#B-RNA** | **#Z-RNA** | **#Unclassified steps** | **#Steps without a continuous backbone** | **Total** |
| 1 | PZ02_refstructure | 4 | 38 | 36 | 14 | 0 | 0 | 12 | 8 | 34 |
| 2 | PZ02_Bujnicki_1 | 4 | 38 | 37 | 5 | 0 | 0 | 21 | 8 | 34 |
| 3 | PZ02_Bujnicki_2 | 3 | 40 | 38 | 0 | 1 | 0 | 26 | 10 | 37 |
| 4 | PZ02_Bujnicki_3 | 3 | 40 | 38 | 0 | 1 | 0 | 26 | 10 | 37 |
| 5 | PZ02_Chen_1 | 4 | 36 | 35 | 9 | 0 | 0 | 14 | 9 | 32 |
| 6 | PZ02_Das_1 | 4 | 40 | 38 | 14 | 0 | 0 | 12 | 10 | 36 |
| 7 | PZ02_Das_2 | 3 | 39 | 37 | 15 | 0 | 0 | 11 | 10 | 36 |
| 8 | PZ02_Das_3 | 4 | 40 | 37 | 14 | 0 | 0 | 12 | 10 | 36 |
| 9 | PZ02_Das_4 | 4 | 40 | 37 | 16 | 0 | 0 | 10 | 10 | 36 |
| 10 | PZ02_Das_5 | 4 | 42 | 37 | 17 | 0 | 0 | 10 | 11 | 38 |
| 11 | PZ02_Dokholyan_1 | 4 | 39 | 37 | 17 | 0 | 0 | 9 | 9 | 35 |
| 12 | PZ02_Major_1 | 4 | 37 | 37 | 22 | 0 | 0 | 11 | 0 | 33 |
| 13 | PZ02_Santalucia_1 | 4 | 40 | 40 | 28 | 0 | 0 | 0 | 8 | 36 |
| 14 | PZ02_Wildauer_1 | 4 | 37 | 35 | 0 | 0 | 0 | 25 | 8 | 33 |
| | Average | 3.79 | 39.00 | 37.07 | 12.21 | 0.14 | 0.00 | 14.21 | 8.64 | 35.21 |
| | Standard deviation | 0.43 | 1.62 | 1.27 | 8.47 | 0.36 | 0.00 | 7.55 | 2.68 | 1.76 |
| | Median | 4.00 | 39.50 | 37.00 | 14.00 | 0.00 | 0.00 | 12.00 | 9.50 | 36.00 |

**Table S50**. X3DNA report on structures in Puzzle 03.

| | | | Base pairs | | Dinucleotide steps | | | | | |
|---|---|---|---|---|---|---|---|---|---|---|
| | | | | | **Right-handed** | | **Left-handed** | **Other** | | |
| **No.** | **3D RNA model** | **#Helices** | **#all bps** | **#canonical / wobble bps** | **#A-RNA** | **#B-RNA** | **#Z-RNA** | **#Unclassified steps** | **#Steps without a continuous backbone** | **Total** |
| 1 | PZ03_Bujnicki_1 | 2 | 33 | 23 | 6 | 0 | 0 | 21 | 4 | 31 |
| 2 | PZ03_Bujnicki_2 | 2 | 34 | 22 | 12 | 0 | 0 | 15 | 5 | 32 |

| | | | | | | | | | | |
|---|---|---|---|---|---|---|---|---|---|---|
| 3 | PZ03_Chen_1 | 3 | 31 | 21 | 3 | 0 | 0 | 22 | 3 | 28 |
| 4 | PZ03_Das_1 | 2 | 37 | 23 | 16 | 0 | 0 | 13 | 6 | 35 |
| 5 | PZ03_Das_2 | 2 | 37 | 23 | 15 | 0 | 0 | 16 | 4 | 35 |
| 6 | PZ03_Das_3 | 2 | 37 | 24 | 15 | 0 | 0 | 14 | 6 | 35 |
| 7 | PZ03_Das_4 | 2 | 38 | 23 | 16 | 0 | 0 | 13 | 7 | 36 |
| 8 | PZ03_Das_5 | 2 | 37 | 24 | 17 | 0 | 0 | 14 | 4 | 35 |
| 9 | PZ03_Dokholyan_1 | 3 | 27 | 23 | 15 | 0 | 0 | 6 | 3 | 24 |
| 10 | PZ03_Dokholyan_2 | 4 | 28 | 22 | 13 | 0 | 0 | 9 | 2 | 24 |
| 11 | PZ03_Major_1 | 4 | 28 | 21 | 4 | 0 | 0 | 16 | 4 | 24 |
| 12 | PZ03_Major_2 | 3 | 32 | 19 | 1 | 0 | 0 | 26 | 2 | 29 |
| 13 | PZ03_refstructure | 2 | 32 | 25 | 13 | 0 | 0 | 11 | 6 | 30 |
| | Average | 2.54 | 33.15 | 22.54 | 11.23 | 0.00 | 0.00 | 15.08 | 4.31 | 30.62 |
| | Standard deviation | 0.78 | 3.89 | 1.56 | 5.63 | 0.00 | 0.00 | 5.41 | 1.60 | 4.56 |
| | Median | 2.00 | 33.00 | 23.00 | 13.00 | 0.00 | 0.00 | 14.00 | 4.00 | 31.00 |

**Table S51**. X3DNA report on structures in Puzzle 04.

| | | | Base pairs | | Dinucleotide steps | | | | | |
|---|---|---|---|---|---|---|---|---|---|---|
| | | | | | **Right-handed** | | **Left-handed** | **Other** | | |
| **No.** | **3D RNA model** | **#Helices** | **#all bps** | **#canonical / wobble bps** | **#A-RNA** | **#B-RNA** | **#Z-RNA** | **#Unclassified steps** | **#Steps without a continuous backbone** | **Total** |
| 1 | PZ04_refstructure | 4 | 51 | 46 | 20 | 0 | 0 | 22 | 5 | 47 |
| 2 | PZ04_Adamiak_1 | 4 | 49 | 44 | 9 | 0 | 0 | 32 | 4 | 45 |
| 3 | PZ04_Adamiak_2 | 5 | 51 | 44 | 19 | 0 | 0 | 24 | 3 | 46 |
| 4 | PZ04_Adamiak_3 | 5 | 52 | 44 | 20 | 0 | 0 | 23 | 4 | 47 |
| 5 | PZ04_Adamiak_4 | 4 | 48 | 26 | 0 | 0 | 0 | 35 | 9 | 44 |
| 6 | PZ04_Adamiak_5 | 4 | 42 | 18 | 0 | 0 | 0 | 30 | 8 | 38 |
| 7 | PZ04_Bujnicki_1 | 4 | 51 | 46 | 32 | 0 | 0 | 10 | 5 | 47 |
| 8 | PZ04_Bujnicki_2 | 4 | 51 | 46 | 32 | 0 | 0 | 10 | 5 | 47 |
| 9 | PZ04_Bujnicki_3 | 4 | 51 | 46 | 32 | 0 | 0 | 10 | 5 | 47 |
| 10 | PZ04_Bujnicki_4 | 4 | 51 | 46 | 32 | 0 | 0 | 10 | 5 | 47 |
| 11 | PZ04_Bujnicki_5 | 4 | 51 | 46 | 32 | 0 | 0 | 10 | 5 | 47 |

| | | | | | | | | | | |
|---|---|---|---|---|---|---|---|---|---|---|
| 12 | PZ04_Chen_1 | 3 | 51 | 45 | 32 | 0 | 0 | 10 | 6 | 48 |
| 13 | PZ04_Chen_10 | 3 | 51 | 45 | 31 | 0 | 0 | 11 | 6 | 48 |
| 14 | PZ04_Chen_2 | 2 | 52 | 45 | 31 | 0 | 0 | 11 | 8 | 50 |
| 15 | PZ04_Chen_3 | 4 | 51 | 45 | 31 | 0 | 0 | 11 | 5 | 47 |
| 16 | PZ04_Chen_4 | 2 | 52 | 45 | 29 | 0 | 0 | 13 | 8 | 50 |
| 17 | PZ04_Chen_5 | 4 | 51 | 45 | 31 | 0 | 0 | 11 | 5 | 47 |
| 18 | PZ04_Chen_6 | 3 | 51 | 45 | 31 | 0 | 0 | 11 | 6 | 48 |
| 19 | PZ04_Chen_7 | 3 | 51 | 45 | 31 | 0 | 0 | 11 | 6 | 48 |
| 20 | PZ04_Chen_8 | 3 | 51 | 45 | 31 | 0 | 0 | 11 | 6 | 48 |
| 21 | PZ04_Chen_9 | 3 | 52 | 45 | 31 | 0 | 0 | 11 | 7 | 49 |
| 22 | PZ04_Das_1 | 3 | 52 | 46 | 28 | 0 | 0 | 15 | 6 | 49 |
| 23 | PZ04_Das_2 | 3 | 52 | 46 | 28 | 0 | 0 | 15 | 6 | 49 |
| 24 | PZ04_Das_3 | 3 | 52 | 46 | 28 | 0 | 0 | 15 | 6 | 49 |
| 25 | PZ04_Das_4 | 3 | 52 | 46 | 28 | 0 | 0 | 15 | 6 | 49 |
| 26 | PZ04_Das_5 | 3 | 52 | 46 | 28 | 0 | 0 | 15 | 6 | 49 |
| 27 | PZ04_Dokholyan_1 | 3 | 53 | 46 | 24 | 0 | 0 | 19 | 7 | 50 |
| 28 | PZ04_Dokholyan_2 | 3 | 53 | 46 | 24 | 0 | 0 | 19 | 7 | 50 |
| 29 | PZ04_Major_1 | 4 | 52 | 47 | 16 | 0 | 0 | 27 | 5 | 48 |
| 30 | PZ04_Mikolajczak_1 | 5 | 39 | 28 | 15 | 0 | 0 | 11 | 8 | 34 |
| 31 | PZ04_Santalucia_1 | 4 | 52 | 46 | 29 | 0 | 0 | 14 | 5 | 48 |
| | Average | 3.55 | 50.65 | 43.39 | 25.32 | 0.00 | 0.00 | 15.87 | 5.90 | 47.10 |
| | Standard deviation | 0.77 | 2.90 | 6.63 | 9.00 | 0.00 | 0.00 | 7.21 | 1.35 | 3.32 |
| | Median | 4.00 | 51.00 | 45.00 | 29.00 | 0.00 | 0.00 | 13.00 | 6.00 | 48.00 |

**Table S52**. X3DNA report on structures in Puzzle 05.

| | | | Base pairs | | Dinucleotide steps | | | | | |
|---|---|---|---|---|---|---|---|---|---|---|
| | | | | | **Right-handed** | | **Left-handed** | **Other** | | |
| **No.** | **3D RNA model** | **#Helices** | **#all bps** | **#canonical / wobble bps** | **#A-RNA** | **#B-RNA** | **#Z-RNA** | **#Unclassified steps** | **#Steps without a continuous backbone** | **Total** |
| 1 | PZ05_refstructure | 7 | 71 | 59 | 37 | 0 | 0 | 18 | 9 | 64 |
| 2 | PZ05_Adamiak_1 | 9 | 67 | 63 | 23 | 0 | 0 | 29 | 6 | 58 |

| | | | | | | | | | | |
|---|---|---|---|---|---|---|---|---|---|---|
| 3 | PZ05_Bujnicki_1 | 9 | 65 | 57 | 11 | 0 | 0 | 39 | 6 | 56 |
| 4 | PZ05_Bujnicki_2 | 7 | 68 | 62 | 16 | 0 | 0 | 36 | 9 | 61 |
| 5 | PZ05_Bujnicki_3 | 10 | 58 | 47 | 15 | 0 | 0 | 23 | 10 | 48 |
| 6 | PZ05_Bujnicki_4 | 8 | 65 | 56 | 17 | 0 | 0 | 33 | 7 | 57 |
| 7 | PZ05_Bujnicki_5 | 7 | 63 | 59 | 25 | 0 | 0 | 21 | 10 | 56 |
| 8 | PZ05_Chen_1 | 9 | 60 | 47 | 30 | 0 | 0 | 16 | 5 | 51 |
| 9 | PZ05_Chen_2 | 9 | 60 | 47 | 20 | 1 | 0 | 24 | 6 | 51 |
| 10 | PZ05_Chen_3 | 9 | 59 | 51 | 28 | 0 | 0 | 18 | 4 | 50 |
| 11 | PZ05_Chen_4 | 9 | 60 | 51 | 25 | 0 | 0 | 22 | 4 | 51 |
| 12 | PZ05_Chen_5 | 8 | 63 | 47 | 31 | 0 | 0 | 16 | 8 | 55 |
| 13 | PZ05_Chen_6 | 11 | 60 | 49 | 25 | 0 | 0 | 22 | 2 | 49 |
| 14 | PZ05_Chen_7 | 8 | 55 | 42 | 18 | 1 | 0 | 22 | 6 | 47 |
| 15 | PZ05_Das_1 | 6 | 77 | 67 | 35 | 0 | 0 | 25 | 11 | 71 |
| 16 | PZ05_Das_2 | 6 | 76 | 64 | 31 | 0 | 0 | 27 | 12 | 70 |
| 17 | PZ05_Dokholyan_1 | 6 | 67 | 63 | 23 | 0 | 0 | 28 | 10 | 61 |
| 18 | PZ05_Dokholyan_2 | 6 | 67 | 62 | 20 | 0 | 0 | 31 | 10 | 61 |
| 19 | PZ05_Dokholyan_3 | 8 | 68 | 63 | 21 | 0 | 0 | 32 | 7 | 60 |
| 20 | PZ05_Dokholyan_4 | 6 | 67 | 62 | 20 | 0 | 0 | 31 | 10 | 61 |
| 21 | PZ05_Dokholyan_5 | 6 | 69 | 62 | 26 | 0 | 0 | 28 | 9 | 63 |
| 22 | PZ05_Dokholyan_6 | 7 | 71 | 62 | 28 | 0 | 0 | 28 | 8 | 64 |
| 23 | PZ05_Dokholyan_7 | 7 | 68 | 62 | 22 | 0 | 0 | 32 | 7 | 61 |
| 24 | PZ05_Dokholyan_8 | 7 | 66 | 61 | 17 | 0 | 0 | 33 | 9 | 59 |
| 25 | PZ05_Xiao_1 | 9 | 61 | 52 | 6 | 0 | 0 | 43 | 3 | 52 |
| 26 | PZ05_Xiao_2 | 9 | 60 | 52 | 7 | 0 | 0 | 40 | 4 | 51 |
| | Average | 7.81 | 65.04 | 56.50 | 22.19 | 0.08 | 0.00 | 27.58 | 7.38 | 57.23 |
| | Standard deviation | 1.41 | 5.41 | 7.03 | 7.75 | 0.27 | 0.00 | 7.34 | 2.65 | 6.51 |
| | Median | 8.00 | 65.50 | 59.00 | 22.50 | 0.00 | 0.00 | 28.00 | 7.50 | 57.50 |

**Table S53**. X3DNA report on structures in Puzzle 06.

| | | | Base pairs | Dinucleotide steps | | | |
|---|---|---|---|---|---|---|---|
| | | | | **Right-handed** | **Left-handed** | **Other** | **Total** |

| No. | 3D RNA model | #Helices | #all bps | #canonical / wobble bps | #A-RNA | #B-RNA | #Z-RNA | #Unclassified steps | #Steps without a continuous backbone | |
|---|---|---|---|---|---|---|---|---|---|---|
| 1 | PZ06_refstructure | 7 | 61 | 49 | 16 | 0 | 0 | 25 | 13 | 54 |
| 2 | PZ06_Bujnicki_1 | 5 | 60 | 54 | 13 | 0 | 0 | 33 | 9 | 55 |
| 3 | PZ06_Bujnicki_2 | 6 | 59 | 54 | 24 | 0 | 0 | 24 | 5 | 53 |
| 4 | PZ06_Bujnicki_3 | 5 | 62 | 51 | 18 | 0 | 0 | 32 | 7 | 57 |
| 5 | PZ06_Bujnicki_4 | 4 | 57 | 50 | 15 | 0 | 0 | 27 | 11 | 53 |
| 6 | PZ06_Chen_1 | 7 | 59 | 45 | 16 | 0 | 0 | 28 | 8 | 52 |
| 7 | PZ06_Chen_2 | 9 | 57 | 46 | 7 | 0 | 0 | 38 | 3 | 48 |
| 8 | PZ06_Chen_3 | 6 | 56 | 47 | 8 | 1 | 0 | 35 | 6 | 50 |
| 9 | PZ06_Chen_4 | 5 | 61 | 47 | 32 | 0 | 0 | 15 | 9 | 56 |
| 10 | PZ06_Chen_5 | 7 | 57 | 47 | 27 | 0 | 0 | 17 | 6 | 50 |
| 11 | PZ06_Chen_6 | 5 | 61 | 46 | 37 | 0 | 0 | 10 | 9 | 56 |
| 12 | PZ06_Chen_7 | 6 | 57 | 46 | 14 | 1 | 0 | 30 | 6 | 51 |
| 13 | PZ06_Das_1 | 7 | 66 | 56 | 31 | 0 | 0 | 20 | 8 | 59 |
| 14 | PZ06_Das_10 | 5 | 66 | 54 | 31 | 0 | 0 | 19 | 11 | 61 |
| 15 | PZ06_Das_2 | 7 | 63 | 55 | 32 | 0 | 0 | 18 | 6 | 56 |
| 16 | PZ06_Das_3 | 7 | 62 | 56 | 28 | 0 | 0 | 22 | 5 | 55 |
| 17 | PZ06_Das_4 | 7 | 67 | 56 | 31 | 0 | 0 | 20 | 9 | 60 |
| 18 | PZ06_Das_5 | 7 | 67 | 55 | 26 | 0 | 0 | 24 | 6 | 56 |
| 19 | PZ06_Das_6 | 7 | 67 | 56 | 31 | 0 | 0 | 20 | 8 | 59 |
| 20 | PZ06_Das_7 | 7 | 67 | 55 | 27 | 0 | 0 | 23 | 5 | 55 |
| 21 | PZ06_Das_8 | 7 | 67 | 55 | 26 | 0 | 0 | 24 | 6 | 56 |
| 22 | PZ06_Das_9 | 7 | 67 | 55 | 32 | 0 | 0 | 18 | 6 | 56 |
| 23 | PZ06_Dokholyan_1 | 6 | 67 | 53 | 16 | 0 | 0 | 34 | 5 | 55 |
| 24 | PZ06_Dokholyan_2 | 6 | 67 | 54 | 14 | 0 | 0 | 34 | 6 | 54 |
| 25 | PZ06_Dokholyan_3 | 7 | 67 | 53 | 22 | 0 | 0 | 23 | 8 | 53 |
| 26 | PZ06_Dokholyan_4 | 6 | 67 | 53 | 14 | 0 | 0 | 35 | 5 | 54 |
| 27 | PZ06_Dokholyan_5 | 5 | 67 | 53 | 20 | 0 | 0 | 28 | 6 | 54 |
| 28 | PZ06_Dokholyan_6 | 4 | 67 | 54 | 15 | 0 | 0 | 35 | 6 | 56 |
| 29 | PZ06_Major_1 | 6 | 67 | 56 | 10 | 1 | 0 | 38 | 8 | 57 |

| | | | | | | | | | | |
|---|---|---|---|---|---|---|---|---|---|---|
| 30 | PZ06_Major_2 | 6 | 67 | 54 | 4 | 0 | 0 | 45 | 7 | 56 |
| 31 | PZ06_Major_3 | 6 | 67 | 56 | 6 | 1 | 0 | 42 | 7 | 56 |
| 32 | PZ06_Major_4 | 5 | 67 | 57 | 7 | 0 | 0 | 44 | 8 | 59 |
| 33 | PZ06_Major_5 | 6 | 67 | 56 | 7 | 1 | 0 | 41 | 7 | 56 |
| 34 | PZ06_Major_6 | 5 | 67 | 57 | 9 | 1 | 0 | 41 | 8 | 59 |
| 35 | PZ06_Major_7 | 6 | 67 | 56 | 15 | 0 | 0 | 36 | 7 | 58 |
| | Average | 6.11 | 67 | 52.77 | 19.46 | 0.17 | 0.00 | 28.51 | 7.14 | 55.29 |
| | Standard deviation | 1.05 | 67 | 3.74 | 9.48 | 0.38 | 0.00 | 9.12 | 2.00 | 2.92 |
| | Median | 6.00 | 67 | 54.00 | 16.00 | 0.00 | 0.00 | 28.00 | 7.00 | 56.00 |

**Table S54**. X3DNA report on structures in Puzzle 07.

| | | | Base pairs | | Dinucleotide steps | | | | | |
|---|---|---|---|---|---|---|---|---|---|---|
| | | | | | **Right-handed** | | **Left-handed** | **Other** | | |
| **No.** | **3D RNA model** | **#Helices** | **#all bps** | **#canonical / wobble bps** | **#A-RNA** | **#B-RNA** | **#Z-RNA** | **#Unclassified steps** | **#Steps without a continuous backbone** | **Total** |
| 1 | PZ07_refstructure | 4 | 73 | 61 | 30 | 0 | 0 | 25 | 14 | 69 |
| 2 | PZ07_Adamiak_1 | 4 | 76 | 62 | 29 | 0 | 0 | 30 | 13 | 72 |
| 3 | PZ07_Adamiak_2 | 6 | 73 | 61 | 23 | 0 | 0 | 34 | 10 | 67 |
| 4 | PZ07_Adamiak_3 | 6 | 72 | 60 | 26 | 0 | 0 | 29 | 11 | 66 |
| 5 | PZ07_Adamiak_4 | 3 | 76 | 65 | 26 | 0 | 0 | 34 | 13 | 73 |
| 6 | PZ07_Adamiak_5 | 5 | 74 | 65 | 29 | 0 | 0 | 30 | 10 | 69 |
| 7 | PZ07_Bujnicki_1 | 3 | 41 | 32 | 16 | 0 | 0 | 14 | 8 | 38 |
| 8 | PZ07_Bujnicki_2 | 3 | 37 | 33 | 13 | 0 | 0 | 17 | 4 | 34 |
| 9 | PZ07_Bujnicki_3 | 3 | 77 | 68 | 40 | 0 | 0 | 21 | 13 | 74 |
| 10 | PZ07_Bujnicki_4 | 3 | 76 | 68 | 31 | 0 | 0 | 30 | 12 | 73 |
| 11 | PZ07_Bujnicki_5 | 3 | 76 | 68 | 31 | 0 | 0 | 30 | 12 | 73 |
| 12 | PZ07_Bujnicki_6 | 3 | 77 | 68 | 40 | 0 | 0 | 21 | 13 | 74 |
| 13 | PZ07_Bujnicki_7 | 3 | 77 | 68 | 36 | 0 | 0 | 25 | 13 | 74 |
| 14 | PZ07_Chen_1 | 8 | 44 | 30 | 0 | 0 | 0 | 30 | 6 | 36 |
| 15 | PZ07_Chen_10 | 5 | 70 | 59 | 43 | 0 | 0 | 11 | 11 | 65 |
| 16 | PZ07_Chen_2 | 7 | 71 | 58 | 41 | 0 | 0 | 10 | 13 | 64 |

| | | | | | | | | | | |
|---|---|---|---|---|---|---|---|---|---|---|
| 17 | PZ07_Chen_3 | 5 | 68 | 58 | 41 | 0 | 0 | 10 | 12 | 63 |
| 18 | PZ07_Chen_4 | 4 | 69 | 57 | 41 | 0 | 0 | 12 | 12 | 65 |
| 19 | PZ07_Chen_5 | 5 | 69 | 58 | 41 | 0 | 0 | 10 | 13 | 64 |
| 20 | PZ07_Chen_6 | 5 | 69 | 58 | 41 | 0 | 0 | 9 | 14 | 64 |
| 21 | PZ07_Chen_7 | 6 | 69 | 57 | 40 | 0 | 0 | 11 | 12 | 63 |
| 22 | PZ07_Chen_8 | 5 | 71 | 59 | 43 | 0 | 0 | 11 | 12 | 66 |
| 23 | PZ07_Chen_9 | 6 | 70 | 58 | 40 | 0 | 0 | 12 | 12 | 64 |
| 24 | PZ07_Das_1 | 4 | 75 | 62 | 39 | 0 | 0 | 21 | 11 | 71 |
| 25 | PZ07_Das_2 | 4 | 75 | 63 | 41 | 0 | 0 | 19 | 11 | 71 |
| 26 | PZ07_Das_3 | 4 | 74 | 61 | 40 | 0 | 0 | 20 | 10 | 70 |
| 27 | PZ07_Das_4 | 4 | 73 | 61 | 34 | 0 | 0 | 24 | 11 | 69 |
| 28 | PZ07_Das_5 | 5 | 80 | 65 | 38 | 0 | 0 | 25 | 12 | 75 |
| 29 | PZ07_Das_6 | 3 | 75 | 62 | 37 | 0 | 0 | 24 | 11 | 72 |
| 30 | PZ07_Das_7 | 5 | 71 | 62 | 36 | 0 | 0 | 20 | 10 | 66 |
| 31 | PZ07_Das_8 | 4 | 73 | 62 | 39 | 0 | 0 | 15 | 15 | 69 |
| 32 | PZ07_Ding_1 | 5 | 66 | 63 | 18 | 0 | 0 | 35 | 8 | 61 |
| 33 | PZ07_Ding_10 | 4 | 68 | 63 | 24 | 0 | 0 | 29 | 11 | 64 |
| 34 | PZ07_Ding_2 | 6 | 67 | 63 | 24 | 0 | 0 | 29 | 8 | 61 |
| 35 | PZ07_Ding_3 | 5 | 70 | 63 | 13 | 0 | 0 | 41 | 11 | 65 |
| 36 | PZ07_Ding_4 | 5 | 68 | 63 | 19 | 0 | 0 | 32 | 12 | 63 |
| 37 | PZ07_Ding_5 | 5 | 66 | 63 | 18 | 0 | 0 | 31 | 12 | 61 |
| 38 | PZ07_Ding_6 | 5 | 69 | 63 | 21 | 0 | 0 | 34 | 9 | 64 |
| 39 | PZ07_Ding_7 | 5 | 66 | 63 | 28 | 0 | 0 | 24 | 9 | 61 |
| 40 | PZ07_Ding_8 | 5 | 65 | 61 | 27 | 0 | 0 | 23 | 10 | 60 |
| 41 | PZ07_Ding_9 | 6 | 65 | 63 | 22 | 0 | 0 | 29 | 8 | 59 |
| 42 | PZ07_Dokholyan_1 | 7 | 68 | 61 | 14 | 0 | 0 | 40 | 7 | 61 |
| 43 | PZ07_Dokholyan_2 | 6 | 60 | 56 | 17 | 0 | 0 | 30 | 7 | 54 |
| 44 | PZ07_Major_1 | 6 | 72 | 61 | 0 | 2 | 0 | 53 | 11 | 66 |
| 45 | PZ07_Major_10 | 6 | 72 | 60 | 1 | 1 | 0 | 51 | 13 | 66 |
| 46 | PZ07_Major_2 | 6 | 72 | 61 | 0 | 1 | 0 | 53 | 12 | 66 |
| 47 | PZ07_Major_3 | 6 | 73 | 62 | 1 | 0 | 0 | 53 | 13 | 67 |
| 48 | PZ07_Major_4 | 6 | 72 | 60 | 9 | 0 | 0 | 45 | 12 | 66 |
| 49 | PZ07_Major_5 | 6 | 70 | 60 | 8 | 0 | 0 | 43 | 13 | 64 |

| | | | | | | | | | | |
|---|---|---|---|---|---|---|---|---|---|---|
| 50 | PZ07_Major_6 | 6 | 72 | 62 | 3 | 0 | 0 | 51 | 12 | 66 |
| 51 | PZ07_Major_7 | 6 | 70 | 60 | 8 | 0 | 0 | 43 | 13 | 64 |
| 52 | PZ07_Major_8 | 6 | 73 | 60 | 0 | 1 | 0 | 53 | 13 | 67 |
| 53 | PZ07_Major_9 | 6 | 72 | 61 | 0 | 0 | 0 | 55 | 11 | 66 |
| | Average | 4.94 | 69.57 | 60.02 | 24.91 | 0.09 | 0.00 | 28.51 | 11.11 | 64.62 |
| | Standard deviation | 1.23 | 8.09 | 7.58 | 14.41 | 0.35 | 0.00 | 13.29 | 2.19 | 8.34 |
| | Median | 5.00 | 71.00 | 61.00 | 27.00 | 0.00 | 0.00 | 29.00 | 12.00 | 66.00 |

**Table S55**. X3DNA report on structures in Puzzle 08.

| | | | Base pairs | | Dinucleotide steps | | | | | |
|---|---|---|---|---|---|---|---|---|---|---|
| | | | | | **Right-handed** | | **Left-handed** | **Other** | | |
| **No.** | **3D RNA model** | **#Helices** | **#all bps** | **#canonical / wobble bps** | **#A-RNA** | **#B-RNA** | **#Z-RNA** | **#Unclassified steps** | **#Steps without a continuous backbone** | **Total** |
| 1 | PZ08_refstructure | 3 | 36 | 31 | 19 | 0 | 0 | 9 | 5 | 33 |
| 2 | PZ08_Adamiak_1 | 5 | 35 | 32 | 16 | 0 | 0 | 11 | 3 | 30 |
| 3 | PZ08_Adamiak_2 | 4 | 35 | 32 | 17 | 0 | 0 | 10 | 4 | 31 |
| 4 | PZ08_Bujnicki_1 | 4 | 37 | 32 | 17 | 0 | 0 | 11 | 5 | 33 |
| 5 | PZ08_Bujnicki_10 | 5 | 33 | 30 | 19 | 0 | 0 | 6 | 3 | 28 |
| 6 | PZ08_Bujnicki_2 | 2 | 35 | 31 | 15 | 0 | 0 | 11 | 7 | 33 |
| 7 | PZ08_Bujnicki_3 | 3 | 35 | 30 | 11 | 0 | 0 | 15 | 6 | 32 |
| 8 | PZ08_Bujnicki_4 | 2 | 34 | 30 | 17 | 0 | 0 | 8 | 7 | 32 |
| 9 | PZ08_Bujnicki_5 | 3 | 36 | 32 | 16 | 0 | 0 | 11 | 6 | 33 |
| 10 | PZ08_Bujnicki_6 | 3 | 37 | 32 | 16 | 0 | 0 | 12 | 6 | 34 |
| 11 | PZ08_Bujnicki_7 | 5 | 35 | 31 | 11 | 0 | 0 | 15 | 4 | 30 |
| 12 | PZ08_Bujnicki_8 | 4 | 37 | 30 | 21 | 0 | 0 | 8 | 4 | 33 |
| 13 | PZ08_Bujnicki_9 | 6 | 33 | 29 | 15 | 0 | 0 | 9 | 3 | 27 |
| 14 | PZ08_Chen_1 | 4 | 35 | 31 | 12 | 0 | 0 | 15 | 4 | 31 |
| 15 | PZ08_Chen_10 | 4 | 33 | 29 | 14 | 0 | 0 | 11 | 4 | 29 |
| 16 | PZ08_Chen_2 | 5 | 33 | 29 | 13 | 0 | 0 | 12 | 3 | 28 |
| 17 | PZ08_Chen_3 | 3 | 34 | 30 | 12 | 0 | 0 | 13 | 6 | 31 |
| 18 | PZ08_Chen_4 | 5 | 33 | 29 | 14 | 0 | 0 | 11 | 3 | 28 |

| | | | | | | | | | | |
|---|---|---|---|---|---|---|---|---|---|---|
| 19 | PZ08_Chen_5 | 3 | 35 | 31 | 14 | 0 | 0 | 13 | 5 | 32 |
| 20 | PZ08_Chen_6 | 4 | 35 | 30 | 4 | 0 | 0 | 22 | 5 | 31 |
| 21 | PZ08_Chen_7 | 4 | 32 | 28 | 12 | 0 | 0 | 12 | 4 | 28 |
| 22 | PZ08_Chen_8 | 5 | 31 | 26 | 5 | 0 | 0 | 18 | 3 | 26 |
| 23 | PZ08_Chen_9 | 5 | 33 | 28 | 3 | 0 | 0 | 20 | 5 | 28 |
| 24 | PZ08_Das_1 | 3 | 37 | 32 | 16 | 0 | 0 | 13 | 5 | 34 |
| 25 | PZ08_Das_2 | 3 | 36 | 32 | 19 | 0 | 0 | 9 | 5 | 33 |
| 26 | PZ08_Das_3 | 3 | 36 | 31 | 20 | 0 | 0 | 8 | 5 | 33 |
| 27 | PZ08_Das_4 | 2 | 35 | 32 | 15 | 0 | 0 | 12 | 6 | 33 |
| 28 | PZ08_Das_5 | 3 | 37 | 32 | 15 | 0 | 0 | 12 | 7 | 34 |
| 29 | PZ08_Das_6 | 2 | 35 | 32 | 16 | 0 | 0 | 11 | 6 | 33 |
| 30 | PZ08_Ding_1 | 3 | 33 | 32 | 8 | 0 | 0 | 15 | 7 | 30 |
| 31 | PZ08_Ding_10 | 4 | 33 | 31 | 8 | 0 | 0 | 17 | 4 | 29 |
| 32 | PZ08_Ding_2 | 4 | 30 | 30 | 5 | 0 | 0 | 17 | 4 | 26 |
| 33 | PZ08_Ding_3 | 5 | 31 | 30 | 6 | 0 | 0 | 17 | 3 | 26 |
| 34 | PZ08_Ding_4 | 4 | 32 | 30 | 8 | 0 | 0 | 16 | 4 | 28 |
| 35 | PZ08_Ding_5 | 4 | 32 | 29 | 9 | 0 | 0 | 15 | 4 | 28 |
| 36 | PZ08_Ding_6 | 5 | 31 | 29 | 12 | 0 | 0 | 10 | 4 | 26 |
| 37 | PZ08_Ding_7 | 2 | 35 | 29 | 10 | 0 | 0 | 16 | 7 | 33 |
| 38 | PZ08_Ding_8 | 3 | 34 | 30 | 12 | 0 | 0 | 12 | 7 | 31 |
| 39 | PZ08_Ding_9 | 3 | 35 | 32 | 5 | 0 | 0 | 22 | 5 | 32 |
| 40 | PZ08_Dokholyan_1 | 4 | 32 | 29 | 10 | 0 | 0 | 14 | 4 | 28 |
| 41 | PZ08_Dokholyan_2 | 4 | 32 | 29 | 1 | 0 | 0 | 23 | 4 | 28 |
| 42 | PZ08_Dokholyan_3 | 5 | 32 | 29 | 9 | 0 | 0 | 15 | 3 | 27 |
| 43 | PZ08_Dokholyan_4 | 3 | 35 | 32 | 8 | 0 | 0 | 19 | 5 | 32 |
| | Average | 3.72 | 34.07 | 30.35 | 12.21 | 0.00 | 0.00 | 13.40 | 4.74 | 30.35 |
| | Standard deviation | 1.03 | 1.88 | 1.45 | 4.96 | 0.00 | 0.00 | 4.02 | 1.31 | 2.57 |
| | Median | 4.00 | 35.00 | 30.00 | 12.00 | 0.00 | 0.00 | 12.00 | 5.00 | 31.00 |

**Table S56**. X3DNA report on structures in Puzzle 09.

| | | | Base pairs | | Dinucleotide steps | | | | | |
|---|---|---|---|---|---|---|---|---|---|---|
| | | | | | **Right-handed** | | **Left-handed** | **Other** | | **Total** |

| No. | 3D RNA model | #Helices | #all bps | #canonical / wobble bps | #A-RNA | #B-RNA | #Z-RNA | #Unclassified steps | #Steps without a continuous backbone | |
|---|---|---|---|---|---|---|---|---|---|---|
| 1 | PZ09_refstructure | 3 | 31 | 20 | 12 | 0 | 0 | 6 | 10 | 28 |
| 2 | PZ09_Bujnicki_3 | 4 | 30 | 15 | 10 | 0 | 0 | 10 | 6 | 26 |
| 3 | PZ09_Bujnicki_4 | 4 | 30 | 15 | 10 | 0 | 0 | 10 | 6 | 26 |
| 4 | PZ09_Bujnicki_5 | 4 | 30 | 15 | 10 | 0 | 0 | 10 | 6 | 26 |
| 5 | PZ09_Chen_1 | 3 | 27 | 18 | 11 | 0 | 0 | 7 | 6 | 24 |
| 6 | PZ09_Chen_2 | 4 | 27 | 18 | 9 | 0 | 0 | 8 | 6 | 23 |
| 7 | PZ09_Chen_3 | 5 | 26 | 18 | 10 | 0 | 0 | 6 | 5 | 21 |
| 8 | PZ09_Chen_4 | 4 | 27 | 18 | 9 | 0 | 0 | 7 | 7 | 23 |
| 9 | PZ09_Chen_5 | 5 | 26 | 18 | 9 | 0 | 0 | 7 | 5 | 21 |
| 10 | PZ09_Chen_6 | 4 | 24 | 17 | 8 | 0 | 0 | 8 | 4 | 20 |
| 11 | PZ09_Chen_7 | 5 | 25 | 17 | 9 | 0 | 0 | 8 | 3 | 20 |
| 12 | PZ09_Chen_8 | 3 | 27 | 17 | 8 | 0 | 0 | 7 | 9 | 24 |
| 13 | PZ09_Das_1 | 3 | 29 | 19 | 12 | 0 | 0 | 8 | 6 | 26 |
| 14 | PZ09_Das_2 | 3 | 29 | 19 | 12 | 0 | 0 | 6 | 8 | 26 |
| 15 | PZ09_Das_3 | 3 | 28 | 18 | 3 | 0 | 0 | 15 | 7 | 25 |
| 16 | PZ09_Das_4 | 3 | 30 | 20 | 12 | 0 | 0 | 10 | 5 | 27 |
| 17 | PZ09_Das_5 | 4 | 26 | 18 | 12 | 0 | 0 | 7 | 3 | 22 |
| 18 | PZ09_Das_6 | 3 | 29 | 20 | 13 | 0 | 0 | 10 | 3 | 26 |
| 19 | PZ09_Das_7 | 4 | 27 | 18 | 12 | 0 | 0 | 8 | 3 | 23 |
| 20 | PZ09_Das_8 | 4 | 25 | 18 | 12 | 0 | 0 | 7 | 2 | 21 |
| 21 | PZ09_Das_9 | 4 | 28 | 18 | 13 | 0 | 0 | 7 | 4 | 24 |
| 22 | PZ09_Ding_1 | 3 | 24 | 23 | 8 | 0 | 0 | 8 | 5 | 21 |
| 23 | PZ09_Ding_10 | 3 | 22 | 21 | 5 | 0 | 0 | 11 | 3 | 19 |
| 24 | PZ09_Ding_2 | 2 | 24 | 23 | 8 | 0 | 0 | 8 | 6 | 22 |
| 25 | PZ09_Ding_3 | 2 | 24 | 23 | 7 | 0 | 0 | 9 | 6 | 22 |
| 26 | PZ09_Ding_4 | 2 | 26 | 23 | 7 | 0 | 0 | 11 | 6 | 24 |
| 27 | PZ09_Ding_5 | 2 | 27 | 24 | 10 | 0 | 0 | 7 | 8 | 25 |
| 28 | PZ09_Ding_6 | 2 | 25 | 22 | 9 | 0 | 0 | 8 | 6 | 23 |
| 29 | PZ09_Ding_7 | 3 | 20 | 18 | 8 | 0 | 0 | 7 | 2 | 17 |

|  |  |  |  |  |  |  |  |  |  |  |
|---|---|---|---|---|---|---|---|---|---|---|
| 30 | PZ09_Ding_8 | 2 | 24 | 23 | 12 | 0 | 0 | 5 | 5 | 22 |
| 31 | PZ09_Ding_9 | 2 | 23 | 22 | 9 | 0 | 0 | 8 | 4 | 21 |
| 32 | PZ09_Dokholyan_1 | 3 | 22 | 20 | 7 | 0 | 0 | 11 | 1 | 19 |
| 33 | PZ09_Dokholyan_2 | 3 | 21 | 20 | 5 | 0 | 0 | 12 | 1 | 18 |
| 34 | PZ09_Dokholyan_3 | 3 | 23 | 22 | 4 | 0 | 0 | 13 | 3 | 20 |
| 35 | PZ09_Dokholyan_4 | 3 | 26 | 23 | 7 | 0 | 0 | 14 | 2 | 23 |
|  | Average | 3.26 | 26.06 | 19.46 | 9.20 | 0.00 | 0.00 | 8.69 | 4.91 | 22.80 |
|  | Standard deviation | 0.89 | 2.79 | 2.55 | 2.58 | 0.00 | 0.00 | 2.35 | 2.19 | 2.71 |
|  | Median | 3.00 | 26.00 | 19.00 | 9.00 | 0.00 | 0.00 | 8.00 | 5.00 | 23.00 |

**Table S57**. X3DNA report on structures in Puzzle 10.

|  |  |  | Base pairs |  | Dinucleotide steps |  |  |  |  |  |
|---|---|---|---|---|---|---|---|---|---|---|
|  |  |  |  |  | **Right-handed** |  | **Left-handed** | **Other** |  | **Total** |
| **No.** | **3D RNA model** | **#Helices** | **#all bps** | **#canonical / wobble bps** | **#A-RNA** | **#B-RNA** | **#Z-RNA** | **#Unclassified steps** | **#Steps without a continuous backbone** |  |
| 1 | PZ10_refstructure | 4 | 64 | 47 | 19 | 0 | 0 | 25 | 16 | 60 |
| 2 | PZ10_Bujnicki_1 | 5 | 63 | 47 | 23 | 0 | 0 | 17 | 18 | 58 |
| 3 | PZ10_Bujnicki_10 | 7 | 61 | 44 | 18 | 0 | 0 | 23 | 13 | 54 |
| 4 | PZ10_Bujnicki_2 | 8 | 64 | 47 | 22 | 0 | 0 | 19 | 15 | 56 |
| 5 | PZ10_Bujnicki_3 | 4 | 61 | 46 | 26 | 0 | 0 | 15 | 16 | 57 |
| 6 | PZ10_Bujnicki_4 | 5 | 62 | 49 | 26 | 0 | 0 | 14 | 17 | 57 |
| 7 | PZ10_Bujnicki_5 | 6 | 61 | 46 | 22 | 0 | 0 | 17 | 16 | 55 |
| 8 | PZ10_Bujnicki_6 | 5 | 60 | 47 | 26 | 0 | 0 | 13 | 16 | 55 |
| 9 | PZ10_Bujnicki_7 | 6 | 64 | 48 | 21 | 0 | 0 | 20 | 17 | 58 |
| 10 | PZ10_Bujnicki_8 | 6 | 65 | 48 | 25 | 0 | 0 | 19 | 15 | 59 |
| 11 | PZ10_Bujnicki_9 | 6 | 62 | 45 | 26 | 0 | 0 | 16 | 14 | 56 |
| 12 | PZ10_Chen_1 | 6 | 50 | 37 | 3 | 0 | 0 | 24 | 17 | 44 |
| 13 | PZ10_Das_1 | 4 | 65 | 47 | 29 | 0 | 0 | 14 | 18 | 61 |
| 14 | PZ10_Das_2 | 4 | 64 | 49 | 30 | 0 | 0 | 14 | 16 | 60 |
| 15 | PZ10_Das_3 | 4 | 62 | 46 | 29 | 0 | 0 | 14 | 15 | 58 |
| 16 | PZ10_Das_4 | 4 | 62 | 47 | 30 | 0 | 0 | 13 | 15 | 58 |

| | | | | | | | | | | |
|---|---|---|---|---|---|---|---|---|---|---|
| 17 | PZ10_Das_5 | 3 | 64 | 47 | 29 | 0 | 0 | 14 | 18 | 61 |
| 18 | PZ10_Dokholyan_1 | 4 | 52 | 46 | 16 | 0 | 0 | 22 | 10 | 48 |
| 19 | PZ10_Dokholyan_10 | 4 | 48 | 46 | 13 | 0 | 0 | 24 | 7 | 44 |
| 20 | PZ10_Dokholyan_2 | 5 | 52 | 45 | 14 | 0 | 0 | 23 | 10 | 47 |
| 21 | PZ10_Dokholyan_3 | 5 | 54 | 47 | 17 | 0 | 0 | 22 | 10 | 49 |
| 22 | PZ10_Dokholyan_4 | 7 | 49 | 47 | 14 | 0 | 0 | 23 | 5 | 42 |
| 23 | PZ10_Dokholyan_5 | 6 | 49 | 45 | 17 | 0 | 0 | 21 | 5 | 43 |
| 24 | PZ10_Dokholyan_6 | 5 | 52 | 47 | 14 | 0 | 0 | 23 | 10 | 47 |
| 25 | PZ10_Dokholyan_7 | 6 | 52 | 47 | 15 | 0 | 0 | 23 | 8 | 46 |
| 26 | PZ10_Dokholyan_8 | 5 | 52 | 47 | 17 | 0 | 0 | 22 | 8 | 47 |
| 27 | PZ10_Dokholyan_9 | 5 | 54 | 48 | 13 | 1 | 0 | 27 | 8 | 49 |
| | Average | 5.15 | 58.07 | 46.37 | 20.52 | 0.04 | 0.00 | 19.30 | 13.07 | 52.93 |
| | Standard deviation | 1.17 | 6.00 | 2.20 | 6.77 | 0.19 | 0.00 | 4.34 | 4.21 | 6.26 |
| | Median | 5.00 | 61.00 | 47.00 | 21.00 | 0.00 | 0.00 | 20.00 | 15.00 | 55.00 |

**Table S58**. X3DNA report on structures in Puzzle 11.

| | | | Base pairs | | Dinucleotide steps | | | | | |
|---|---|---|---|---|---|---|---|---|---|---|
| | | | | | **Right-handed** | | **Left-handed** | **Other** | | |
| **No.** | **3D RNA model** | **#Helices** | **#all bps** | **#canonical / wobble bps** | **#A-RNA** | **#B-RNA** | **#Z-RNA** | **#Unclassified steps** | **#Steps without a continuous backbone** | **Total** |
| 1 | PZ11_Adamiak_1 | 2 | 22 | 20 | 6 | 0 | 0 | 11 | 3 | 20 |
| 2 | PZ11_Adamiak_10 | 1 | 25 | 21 | 7 | 0 | 0 | 13 | 4 | 24 |
| 3 | PZ11_Adamiak_2 | 3 | 21 | 20 | 6 | 0 | 0 | 11 | 1 | 18 |
| 4 | PZ11_Adamiak_3 | 1 | 25 | 22 | 6 | 0 | 0 | 13 | 5 | 24 |
| 5 | PZ11_Adamiak_4 | 2 | 20 | 19 | 7 | 0 | 0 | 9 | 2 | 18 |
| 6 | PZ11_Adamiak_5 | 1 | 23 | 22 | 4 | 0 | 0 | 14 | 4 | 22 |
| 7 | PZ11_Adamiak_6 | 1 | 24 | 19 | 7 | 0 | 0 | 11 | 5 | 23 |
| 8 | PZ11_Adamiak_7 | 1 | 24 | 21 | 6 | 0 | 0 | 12 | 5 | 23 |
| 9 | PZ11_Adamiak_8 | 1 | 25 | 18 | 5 | 0 | 0 | 15 | 4 | 24 |
| 10 | PZ11_Adamiak_9 | 1 | 25 | 20 | 6 | 0 | 0 | 15 | 3 | 24 |
| 11 | PZ11_Bujnicki_1 | 1 | 24 | 22 | 6 | 0 | 0 | 12 | 5 | 23 |

| | | | | | | | | | | |
|---|---|---|---|---|---|---|---|---|---|---|
| 12 | PZ11_Bujnicki_10 | 1 | 24 | 22 | 6 | 0 | 0 | 12 | 5 | 23 |
| 13 | PZ11_Bujnicki_2 | 3 | 21 | 20 | 9 | 0 | 0 | 4 | 5 | 18 |
| 14 | PZ11_Bujnicki_3 | 1 | 23 | 23 | 11 | 0 | 0 | 6 | 5 | 22 |
| 15 | PZ11_Bujnicki_4 | 1 | 23 | 22 | 0 | 2 | 0 | 16 | 4 | 22 |
| 16 | PZ11_Bujnicki_5 | 1 | 24 | 23 | 5 | 0 | 0 | 13 | 5 | 23 |
| 17 | PZ11_Bujnicki_6 | 1 | 24 | 24 | 9 | 0 | 0 | 8 | 6 | 23 |
| 18 | PZ11_Bujnicki_7 | 1 | 23 | 23 | 8 | 0 | 0 | 9 | 5 | 22 |
| 19 | PZ11_Bujnicki_8 | 1 | 24 | 23 | 7 | 0 | 0 | 12 | 4 | 23 |
| 20 | PZ11_Bujnicki_9 | 1 | 24 | 24 | 9 | 0 | 0 | 9 | 5 | 23 |
| 21 | PZ11_Chen_1 | 1 | 23 | 21 | 8 | 0 | 0 | 9 | 5 | 22 |
| 22 | PZ11_Chen_10 | 1 | 25 | 24 | 8 | 0 | 0 | 9 | 7 | 24 |
| 23 | PZ11_Chen_2 | 1 | 23 | 22 | 9 | 0 | 0 | 8 | 5 | 22 |
| 24 | PZ11_Chen_3 | 1 | 24 | 21 | 9 | 0 | 0 | 8 | 6 | 23 |
| 25 | PZ11_Chen_4 | 1 | 23 | 21 | 6 | 0 | 0 | 10 | 6 | 22 |
| 26 | PZ11_Chen_5 | 1 | 23 | 22 | 15 | 0 | 0 | 2 | 5 | 22 |
| 27 | PZ11_Chen_6 | 1 | 25 | 23 | 7 | 0 | 0 | 12 | 5 | 24 |
| 28 | PZ11_Chen_7 | 1 | 24 | 23 | 8 | 0 | 0 | 9 | 6 | 23 |
| 29 | PZ11_Chen_8 | 1 | 24 | 23 | 8 | 0 | 0 | 10 | 5 | 23 |
| 30 | PZ11_Chen_9 | 1 | 24 | 23 | 9 | 0 | 0 | 9 | 5 | 23 |
| 31 | PZ11_Das_1 | 1 | 24 | 21 | 10 | 0 | 0 | 9 | 4 | 23 |
| 32 | PZ11_Das_10 | 2 | 24 | 22 | 8 | 0 | 0 | 10 | 4 | 22 |
| 33 | PZ11_Das_2 | 1 | 24 | 21 | 9 | 0 | 0 | 10 | 4 | 23 |
| 34 | PZ11_Das_3 | 1 | 25 | 22 | 10 | 0 | 0 | 9 | 5 | 24 |
| 35 | PZ11_Das_4 | 1 | 24 | 22 | 11 | 0 | 0 | 8 | 4 | 23 |
| 36 | PZ11_Das_5 | 1 | 25 | 19 | 8 | 0 | 0 | 12 | 4 | 24 |
| 37 | PZ11_Das_6 | 1 | 24 | 21 | 8 | 0 | 0 | 10 | 5 | 23 |
| 38 | PZ11_Das_7 | 1 | 24 | 20 | 9 | 0 | 0 | 10 | 4 | 23 |
| 39 | PZ11_Das_8 | 2 | 24 | 22 | 10 | 0 | 0 | 8 | 4 | 22 |
| 40 | PZ11_Das_9 | 2 | 22 | 20 | 8 | 0 | 0 | 9 | 3 | 20 |
| 41 | PZ11_Ding_1 | 1 | 21 | 18 | 4 | 0 | 0 | 10 | 6 | 20 |
| 42 | PZ11_Ding_10 | 2 | 18 | 15 | 0 | 0 | 0 | 12 | 4 | 16 |
| 43 | PZ11_Ding_2 | 2 | 23 | 20 | 3 | 0 | 0 | 13 | 5 | 21 |
| 44 | PZ11_Ding_3 | 3 | 20 | 19 | 3 | 0 | 0 | 12 | 2 | 17 |

| | | | | | | | | | | |
|---|---|---|---|---|---|---|---|---|---|---|
| 45 | PZ11_Ding_4 | 2 | 20 | 18 | 1 | 0 | 0 | 14 | 3 | 18 |
| 46 | PZ11_Ding_5 | 2 | 15 | 14 | 0 | 0 | 0 | 11 | 2 | 13 |
| 47 | PZ11_Ding_6 | 2 | 19 | 18 | 4 | 0 | 0 | 9 | 4 | 17 |
| 48 | PZ11_Ding_7 | 2 | 22 | 21 | 2 | 0 | 0 | 13 | 5 | 20 |
| 49 | PZ11_Ding_8 | 1 | 20 | 19 | 1 | 0 | 0 | 12 | 6 | 19 |
| 50 | PZ11_Ding_9 | 2 | 20 | 18 | 3 | 0 | 0 | 9 | 6 | 18 |
| 51 | PZ11_Xiao_1 | 2 | 25 | 20 | 1 | 0 | 0 | 20 | 2 | 23 |
| 52 | PZ11_Xiao_2 | 2 | 23 | 18 | 1 | 0 | 0 | 16 | 4 | 21 |
| 53 | PZ11_Xiao_3 | 2 | 25 | 19 | 3 | 0 | 0 | 17 | 3 | 23 |
| 54 | PZ11_refstructure | 1 | 24 | 22 | 3 | 0 | 0 | 16 | 4 | 23 |
| | Average | 1.39 | 22.98 | 20.74 | 6.24 | 0.04 | 0.00 | 10.93 | 4.39 | 21.59 |
| | Standard deviation | 0.60 | 2.06 | 2.12 | 3.28 | 0.27 | 0.00 | 3.15 | 1.22 | 2.45 |
| | Median | 1.00 | 24.00 | 21.00 | 7.00 | 0.00 | 0.00 | 10.50 | 5.00 | 23.00 |

**Table S59**. X3DNA report on structures in Puzzle 12.

| | | | Base pairs | | Dinucleotide steps | | | | | |
|---|---|---|---|---|---|---|---|---|---|---|
| | | | | | **Right-handed** | | **Left-handed** | **Other** | | |
| **No.** | **3D RNA model** | **#Helices** | **#all bps** | **#canonical / wobble bps** | **#A-RNA** | **#B-RNA** | **#Z-RNA** | **#Unclassified steps** | **#Steps without a continuous backbone** | **Total** |
| 1 | PZ12_refstructure | 4 | 46 | 38 | 19 | 0 | 0 | 17 | 6 | 42 |
| 2 | PZ12_Adamiak_1 | 4 | 42 | 36 | 16 | 0 | 0 | 16 | 6 | 38 |
| 3 | PZ12_Adamiak_2 | 4 | 42 | 36 | 14 | 0 | 0 | 18 | 6 | 38 |
| 4 | PZ12_Adamiak_3 | 4 | 44 | 37 | 17 | 0 | 0 | 16 | 7 | 40 |
| 5 | PZ12_Bujnicki_1 | 6 | 40 | 35 | 11 | 0 | 0 | 20 | 3 | 34 |
| 6 | PZ12_Bujnicki_10 | 4 | 44 | 39 | 14 | 0 | 0 | 20 | 6 | 40 |
| 7 | PZ12_Bujnicki_2 | 5 | 44 | 37 | 1 | 0 | 0 | 32 | 6 | 39 |
| 8 | PZ12_Bujnicki_3 | 4 | 45 | 37 | 10 | 0 | 0 | 24 | 7 | 41 |
| 9 | PZ12_Bujnicki_4 | 5 | 45 | 39 | 12 | 0 | 0 | 21 | 7 | 40 |
| 10 | PZ12_Bujnicki_5 | 5 | 43 | 39 | 12 | 0 | 0 | 20 | 6 | 38 |
| 11 | PZ12_Bujnicki_6 | 5 | 47 | 38 | 10 | 0 | 0 | 21 | 11 | 42 |

| | | | | | | | | | | |
|---|---|---|---|---|---|---|---|---|---|---|
| 12 | PZ12_Bujnicki_7 | 3 | 43 | 38 | 13 | 0 | 0 | 21 | 6 | 40 |
| 13 | PZ12_Bujnicki_8 | 4 | 44 | 40 | 10 | 0 | 0 | 23 | 7 | 40 |
| 14 | PZ12_Bujnicki_9 | 5 | 46 | 41 | 9 | 0 | 0 | 24 | 8 | 41 |
| 15 | PZ12_Chen_1 | 6 | 42 | 36 | 14 | 0 | 0 | 19 | 3 | 36 |
| 16 | PZ12_Chen_10 | 7 | 40 | 38 | 20 | 0 | 0 | 10 | 3 | 33 |
| 17 | PZ12_Chen_2 | 5 | 40 | 35 | 13 | 0 | 0 | 16 | 6 | 35 |
| 18 | PZ12_Chen_3 | 6 | 40 | 37 | 20 | 0 | 0 | 9 | 5 | 34 |
| 19 | PZ12_Chen_4 | 6 | 38 | 35 | 20 | 0 | 0 | 8 | 4 | 32 |
| 20 | PZ12_Chen_5 | 7 | 40 | 35 | 23 | 0 | 0 | 7 | 3 | 33 |
| 21 | PZ12_Chen_6 | 5 | 39 | 36 | 19 | 0 | 0 | 8 | 7 | 34 |
| 22 | PZ12_Chen_7 | 7 | 37 | 35 | 19 | 0 | 0 | 10 | 1 | 30 |
| 23 | PZ12_Chen_8 | 6 | 38 | 35 | 18 | 0 | 0 | 12 | 2 | 32 |
| 24 | PZ12_Chen_9 | 6 | 39 | 35 | 19 | 0 | 0 | 10 | 4 | 33 |
| 25 | PZ12_Das_1 | 4 | 44 | 38 | 24 | 0 | 0 | 10 | 6 | 40 |
| 26 | PZ12_Das_10 | 6 | 44 | 38 | 26 | 0 | 0 | 9 | 3 | 38 |
| 27 | PZ12_Das_2 | 4 | 43 | 38 | 24 | 0 | 0 | 10 | 5 | 39 |
| 28 | PZ12_Das_3 | 4 | 46 | 40 | 23 | 0 | 0 | 12 | 7 | 42 |
| 29 | PZ12_Das_4 | 4 | 44 | 38 | 25 | 0 | 0 | 8 | 7 | 40 |
| 30 | PZ12_Das_5 | 4 | 41 | 37 | 23 | 0 | 0 | 9 | 5 | 37 |
| 31 | PZ12_Das_6 | 5 | 44 | 39 | 23 | 0 | 0 | 11 | 5 | 39 |
| 32 | PZ12_Das_7 | 4 | 44 | 38 | 21 | 0 | 0 | 14 | 5 | 40 |
| 33 | PZ12_Das_8 | 4 | 45 | 39 | 25 | 0 | 0 | 10 | 6 | 41 |
| 34 | PZ12_Das_9 | 4 | 45 | 38 | 24 | 0 | 0 | 12 | 5 | 41 |
| 35 | PZ12_Ding_1 | 4 | 38 | 38 | 19 | 0 | 0 | 10 | 5 | 34 |
| 36 | PZ12_Ding_10 | 4 | 39 | 38 | 14 | 0 | 0 | 15 | 6 | 35 |
| 37 | PZ12_Ding_11 | 4 | 40 | 38 | 23 | 0 | 0 | 8 | 5 | 36 |
| 38 | PZ12_Ding_12 | 4 | 39 | 38 | 18 | 0 | 0 | 12 | 5 | 35 |
| 39 | PZ12_Ding_2 | 4 | 40 | 38 | 18 | 0 | 0 | 12 | 6 | 36 |
| 40 | PZ12_Ding_3 | 4 | 40 | 36 | 16 | 0 | 0 | 14 | 6 | 36 |
| 41 | PZ12_Ding_4 | 4 | 41 | 38 | 16 | 0 | 0 | 15 | 6 | 37 |
| 42 | PZ12_Ding_5 | 4 | 40 | 38 | 16 | 0 | 0 | 14 | 6 | 36 |
| 43 | PZ12_Ding_6 | 4 | 38 | 38 | 13 | 0 | 0 | 16 | 5 | 34 |
| 44 | PZ12_Ding_7 | 4 | 40 | 38 | 14 | 0 | 0 | 17 | 5 | 36 |

| | | | | | | | | | | |
|---|---|---|---|---|---|---|---|---|---|---|
| 45 | PZ12_Ding_8 | 5 | 39 | 38 | 16 | 0 | 0 | 14 | 4 | 34 |
| 46 | PZ12_Ding_9 | 4 | 39 | 38 | 12 | 0 | 0 | 18 | 5 | 35 |
| 47 | PZ12_Weeks_1 | 5 | 45 | 40 | 20 | 0 | 0 | 13 | 7 | 40 |
| 48 | PZ12_Weeks_2 | 4 | 44 | 41 | 17 | 0 | 0 | 15 | 8 | 40 |
| 49 | PZ12_Weeks_3 | 7 | 40 | 38 | 14 | 0 | 0 | 16 | 3 | 33 |
| 50 | PZ12_Xiao_1 | 8 | 32 | 27 | 6 | 0 | 0 | 17 | 1 | 24 |
| 51 | PZ12_Xiao_2 | 8 | 32 | 28 | 7 | 0 | 0 | 16 | 1 | 24 |
| 52 | PZ12_Xiao_3 | 7 | 32 | 28 | 4 | 0 | 0 | 20 | 1 | 25 |
| | Average | 4.88 | 41.27 | 37.04 | 16.42 | 0.00 | 0.00 | 14.79 | 5.17 | 36.38 |
| | Standard deviation | 1.22 | 3.49 | 2.79 | 5.72 | 0.00 | 0.00 | 5.21 | 1.98 | 4.31 |
| | Median | 4.00 | 41.00 | 38.00 | 16.50 | 0.00 | 0.00 | 14.50 | 5.50 | 36.50 |

**Table S60**. X3DNA report on structures in Puzzle 13.

| | | | Base pairs | | Dinucleotide steps | | | | | |
|---|---|---|---|---|---|---|---|---|---|---|
| | | | | | **Right-handed** | | **Left-handed** | **Other** | | |
| **No.** | **3D RNA model** | **#Helices** | **#all bps** | **#canonical / wobble bps** | **#A-RNA** | **#B-RNA** | **#Z-RNA** | **#Unclassified steps** | **#Steps without a continuous backbone** | **Total** |
| 1 | PZ13_refstructure | 2 | 26 | 21 | 14 | 0 | 0 | 7 | 3 | 24 |
| 2 | PZ13_Adamiak_1 | 3 | 21 | 19 | 13 | 0 | 0 | 3 | 2 | 18 |
| 3 | PZ13_Bujnicki_1 | 2 | 24 | 21 | 7 | 0 | 0 | 11 | 4 | 22 |
| 4 | PZ13_Bujnicki_10 | 4 | 25 | 25 | 15 | 0 | 0 | 4 | 2 | 21 |
| 5 | PZ13_Bujnicki_2 | 2 | 24 | 21 | 10 | 0 | 0 | 8 | 4 | 22 |
| 6 | PZ13_Bujnicki_3 | 3 | 25 | 23 | 15 | 0 | 0 | 3 | 4 | 22 |
| 7 | PZ13_Bujnicki_4 | 2 | 23 | 21 | 10 | 0 | 0 | 8 | 3 | 21 |
| 8 | PZ13_Bujnicki_5 | 3 | 23 | 22 | 14 | 0 | 0 | 2 | 4 | 20 |
| 9 | PZ13_Bujnicki_6 | 4 | 23 | 21 | 12 | 0 | 0 | 6 | 1 | 19 |
| 10 | PZ13_Bujnicki_7 | 2 | 24 | 21 | 14 | 0 | 0 | 5 | 3 | 22 |
| 11 | PZ13_Bujnicki_8 | 3 | 24 | 22 | 14 | 0 | 0 | 4 | 3 | 21 |
| 12 | PZ13_Bujnicki_9 | 3 | 22 | 21 | 11 | 0 | 0 | 6 | 2 | 19 |
| 13 | PZ13_Chen_1 | 4 | 23 | 20 | 8 | 0 | 0 | 10 | 1 | 19 |
| 14 | PZ13_Chen_2 | 4 | 22 | 19 | 0 | 0 | 0 | 17 | 1 | 18 |

| | | | | | | | | | | |
|---|---|---|---|---|---|---|---|---|---|---|
| 15 | PZ13_Chen_3 | 3 | 20 | 19 | 11 | 0 | 0 | 5 | 1 | 17 |
| 16 | PZ13_Chen_4 | 4 | 24 | 20 | 7 | 0 | 0 | 11 | 2 | 20 |
| 17 | PZ13_Chen_5 | 3 | 20 | 19 | 5 | 0 | 0 | 11 | 1 | 17 |
| 18 | PZ13_Chen_6 | 4 | 25 | 20 | 11 | 0 | 0 | 9 | 1 | 21 |
| 19 | PZ13_Chen_7 | 4 | 19 | 18 | 6 | 0 | 0 | 9 | 0 | 15 |
| 20 | PZ13_Chen_8 | 4 | 19 | 18 | 10 | 0 | 0 | 5 | 0 | 15 |
| 21 | PZ13_Chen_9 | 4 | 21 | 18 | 12 | 0 | 0 | 4 | 1 | 17 |
| 22 | PZ13_Das_1 | 3 | 25 | 21 | 14 | 0 | 0 | 4 | 4 | 22 |
| 23 | PZ13_Das_10 | 4 | 24 | 21 | 14 | 0 | 0 | 3 | 3 | 20 |
| 24 | PZ13_Das_2 | 3 | 22 | 19 | 14 | 0 | 0 | 4 | 1 | 19 |
| 25 | PZ13_Das_3 | 4 | 22 | 19 | 13 | 0 | 0 | 5 | 0 | 18 |
| 26 | PZ13_Das_4 | 2 | 23 | 20 | 14 | 0 | 0 | 4 | 3 | 21 |
| 27 | PZ13_Das_5 | 3 | 24 | 20 | 15 | 0 | 0 | 3 | 3 | 21 |
| 28 | PZ13_Das_6 | 4 | 22 | 20 | 12 | 0 | 0 | 5 | 1 | 18 |
| 29 | PZ13_Das_7 | 3 | 23 | 20 | 14 | 0 | 0 | 4 | 2 | 20 |
| 30 | PZ13_Das_8 | 4 | 25 | 20 | 16 | 0 | 0 | 5 | 0 | 21 |
| 31 | PZ13_Das_9 | 3 | 22 | 19 | 12 | 0 | 0 | 6 | 1 | 19 |
| 32 | PZ13_Ding_1 | 3 | 21 | 19 | 4 | 0 | 0 | 11 | 3 | 18 |
| 33 | PZ13_Ding_10 | 4 | 21 | 20 | 12 | 0 | 0 | 5 | 0 | 17 |
| 34 | PZ13_Ding_2 | 3 | 22 | 20 | 6 | 0 | 0 | 12 | 1 | 19 |
| 35 | PZ13_Ding_3 | 3 | 24 | 20 | 7 | 0 | 0 | 13 | 1 | 21 |
| 36 | PZ13_Ding_4 | 4 | 20 | 19 | 4 | 0 | 0 | 12 | 0 | 16 |
| 37 | PZ13_Ding_5 | 2 | 22 | 20 | 10 | 0 | 0 | 7 | 3 | 20 |
| 38 | PZ13_Ding_6 | 4 | 21 | 19 | 7 | 0 | 0 | 10 | 0 | 17 |
| 39 | PZ13_Ding_7 | 3 | 22 | 20 | 9 | 0 | 0 | 8 | 2 | 19 |
| 40 | PZ13_Ding_8 | 3 | 24 | 20 | 7 | 0 | 0 | 11 | 3 | 21 |
| 41 | PZ13_Ding_9 | 3 | 23 | 20 | 10 | 0 | 0 | 8 | 2 | 20 |
| 42 | PZ13_Dokholyan_1 | 5 | 22 | 18 | 7 | 0 | 0 | 10 | 0 | 17 |
| 43 | PZ13_Dokholyan_2 | 3 | 21 | 18 | 9 | 0 | 0 | 9 | 0 | 18 |
| 44 | PZ13_Dokholyan_3 | 3 | 18 | 14 | 5 | 0 | 0 | 8 | 2 | 15 |
| 45 | PZ13_Dokholyan_4 | 2 | 14 | 13 | 3 | 0 | 0 | 7 | 2 | 12 |
| 46 | PZ13_Dokholyan_5 | 3 | 18 | 17 | 5 | 0 | 0 | 9 | 1 | 15 |
| 47 | PZ13_Xiao_1 | 2 | 12 | 9 | 4 | 0 | 0 | 4 | 2 | 10 |

| | | | | | | | | | | |
|---|---|---|---|---|---|---|---|---|---|---|
| 48 | PZ13_Xiao_10 | 4 | 18 | 15 | 3 | 0 | 0 | 11 | 0 | 14 |
| 49 | PZ13_Xiao_2 | 2 | 6 | 4 | 0 | 0 | 0 | 4 | 0 | 4 |
| 50 | PZ13_Xiao_3 | 1 | 5 | 2 | 1 | 0 | 0 | 2 | 1 | 4 |
| 51 | PZ13_Xiao_4 | 2 | 19 | 17 | 4 | 0 | 0 | 9 | 4 | 17 |
| 52 | PZ13_Xiao_5 | 3 | 21 | 18 | 8 | 0 | 0 | 10 | 0 | 18 |
| 53 | PZ13_Xiao_6 | 3 | 15 | 13 | 3 | 0 | 0 | 8 | 1 | 12 |
| 54 | PZ13_Xiao_7 | 5 | 16 | 15 | 4 | 0 | 0 | 7 | 0 | 11 |
| 55 | PZ13_Xiao_8 | 3 | 19 | 17 | 3 | 0 | 0 | 11 | 2 | 16 |
| 56 | PZ13_Xiao_9 | 3 | 16 | 13 | 4 | 0 | 0 | 9 | 0 | 13 |
| | Average | 3.14 | 20.88 | 18.36 | 8.86 | 0.00 | 0.00 | 7.25 | 1.63 | 17.73 |
| | Standard deviation | 0.84 | 4.19 | 4.03 | 4.45 | 0.00 | 0.00 | 3.27 | 1.33 | 4.03 |
| | Median | 3.00 | 22.00 | 19.50 | 9.50 | 0.00 | 0.00 | 7.00 | 1.00 | 18.50 |

**Table S61**. X3DNA report on structures in Puzzle 14a.

| | | | Base pairs | | Dinucleotide steps | | | | | |
|---|---|---|---|---|---|---|---|---|---|---|
| | | | | | **Right-handed** | | **Left-handed** | **Other** | | |
| **No.** | **3D RNA model** | **#Helices** | **#all bps** | **#canonical / wobble bps** | **#A-RNA** | **#B-RNA** | **#Z-RNA** | **#Unclassified steps** | **#Steps without a continuous backbone** | **Total** |
| 1 | PZ14_refstructure_bound | 2 | 22 | 17 | 10 | 0 | 0 | 7 | 3 | 20 |
| 2 | PZ14_AdamiakPostExp_1 | 2 | 21 | 16 | 11 | 0 | 0 | 6 | 2 | 19 |
| 3 | PZ14_AdamiakPostExp_2 | 2 | 19 | 16 | 4 | 0 | 0 | 10 | 3 | 17 |
| 4 | PZ14_AdamiakPreExp_1 | 2 | 21 | 16 | 10 | 0 | 0 | 7 | 2 | 19 |
| 5 | PZ14_AdamiakPreExp_2 | 3 | 21 | 16 | 6 | 0 | 0 | 9 | 3 | 18 |
| 6 | PZ14_BujnickiPostExp_1 | 2 | 19 | 16 | 9 | 0 | 0 | 6 | 2 | 17 |
| 7 | PZ14_BujnickiPostExp_2 | 3 | 20 | 18 | 12 | 0 | 0 | 3 | 2 | 17 |
| 8 | PZ14_BujnickiPostExp_3 | 4 | 24 | 21 | 11 | 0 | 0 | 5 | 4 | 20 |
| 9 | PZ14_BujnickiPostExp_4 | 4 | 23 | 19 | 12 | 0 | 0 | 7 | 0 | 19 |
| 10 | PZ14_BujnickiPostExp_5 | 2 | 20 | 16 | 10 | 0 | 0 | 6 | 2 | 18 |
| 11 | PZ14_BujnickiPreExp_4 | 2 | 26 | 17 | 13 | 0 | 0 | 8 | 3 | 24 |
| 12 | PZ14_BujnickiPreExp_5 | 2 | 27 | 17 | 11 | 0 | 0 | 11 | 3 | 25 |
| 13 | PZ14_BujnickiPreExp_6 | 2 | 23 | 18 | 10 | 0 | 0 | 8 | 3 | 21 |

| | | | | | | | | | | |
|---|---|---|---|---|---|---|---|---|---|---|
| 14 | PZ14_BujnickiPreExp_7 | 4 | 23 | 23 | 9 | 0 | 0 | 7 | 3 | 19 |
| 15 | PZ14_ChenPostExp_1 | 2 | 20 | 17 | 9 | 0 | 0 | 5 | 4 | 18 |
| 16 | PZ14_ChenPostExp_10 | 3 | 19 | 16 | 12 | 0 | 0 | 3 | 1 | 16 |
| 17 | PZ14_ChenPostExp_2 | 2 | 18 | 16 | 10 | 0 | 0 | 5 | 1 | 16 |
| 18 | PZ14_ChenPostExp_3 | 3 | 20 | 16 | 13 | 0 | 0 | 3 | 1 | 17 |
| 19 | PZ14_ChenPostExp_4 | 3 | 20 | 16 | 12 | 0 | 0 | 4 | 1 | 17 |
| 20 | PZ14_ChenPostExp_5 | 3 | 17 | 16 | 13 | 0 | 0 | 1 | 0 | 14 |
| 21 | PZ14_ChenPostExp_6 | 3 | 17 | 16 | 11 | 0 | 0 | 3 | 0 | 14 |
| 22 | PZ14_ChenPostExp_7 | 2 | 18 | 16 | 13 | 0 | 0 | 2 | 1 | 16 |
| 23 | PZ14_ChenPostExp_8 | 2 | 20 | 16 | 13 | 0 | 0 | 2 | 3 | 18 |
| 24 | PZ14_ChenPostExp_9 | 3 | 17 | 16 | 13 | 0 | 0 | 1 | 0 | 14 |
| 25 | PZ14_DasPostExp_1 | 3 | 25 | 16 | 9 | 0 | 0 | 12 | 1 | 22 |
| 26 | PZ14_DasPostExp_10 | 2 | 25 | 17 | 9 | 0 | 0 | 11 | 3 | 23 |
| 27 | PZ14_DasPostExp_2 | 2 | 25 | 17 | 9 | 0 | 0 | 11 | 3 | 23 |
| 28 | PZ14_DasPostExp_3 | 2 | 25 | 16 | 9 | 0 | 0 | 12 | 2 | 23 |
| 29 | PZ14_DasPostExp_4 | 2 | 24 | 16 | 10 | 0 | 0 | 10 | 2 | 22 |
| 30 | PZ14_DasPostExp_5 | 2 | 23 | 16 | 10 | 0 | 0 | 8 | 3 | 21 |
| 31 | PZ14_DasPostExp_6 | 2 | 25 | 17 | 9 | 0 | 0 | 10 | 4 | 23 |
| 32 | PZ14_DasPostExp_7 | 2 | 24 | 17 | 10 | 0 | 0 | 10 | 2 | 22 |
| 33 | PZ14_DasPostExp_8 | 1 | 24 | 17 | 9 | 0 | 0 | 9 | 5 | 23 |
| 34 | PZ14_DasPostExp_9 | 3 | 23 | 16 | 10 | 0 | 0 | 9 | 1 | 20 |
| 35 | PZ14_DasPreExp_1 | 2 | 25 | 17 | 9 | 0 | 0 | 10 | 4 | 23 |
| 36 | PZ14_DasPreExp_10 | 3 | 23 | 16 | 10 | 0 | 0 | 9 | 1 | 20 |
| 37 | PZ14_DasPreExp_2 | 2 | 24 | 17 | 10 | 0 | 0 | 10 | 2 | 22 |
| 38 | PZ14_DasPreExp_3 | 1 | 24 | 17 | 9 | 0 | 0 | 9 | 5 | 23 |
| 39 | PZ14_DasPreExp_4 | 4 | 26 | 17 | 8 | 0 | 0 | 13 | 1 | 22 |
| 40 | PZ14_DasPreExp_5 | 2 | 25 | 17 | 9 | 0 | 0 | 11 | 3 | 23 |
| 41 | PZ14_DasPreExp_6 | 2 | 26 | 17 | 9 | 0 | 0 | 12 | 3 | 24 |
| 42 | PZ14_DasPreExp_7 | 2 | 24 | 16 | 9 | 0 | 0 | 11 | 2 | 22 |
| 43 | PZ14_DasPreExp_8 | 2 | 24 | 17 | 10 | 0 | 0 | 10 | 2 | 22 |
| 44 | PZ14_DasPreExp_9 | 2 | 24 | 17 | 10 | 0 | 0 | 10 | 2 | 22 |
| 45 | PZ14_DingPostExp_1 | 2 | 18 | 16 | 8 | 0 | 0 | 7 | 1 | 16 |
| 46 | PZ14_DingPostExp_10 | 2 | 17 | 16 | 5 | 0 | 0 | 9 | 1 | 15 |

| | | | | | | | | | | |
|---|---|---|---|---|---|---|---|---|---|---|
| 47 | PZ14_DingPostExp_2 | 2 | 21 | 16 | 10 | 0 | 0 | 6 | 3 | 19 |
| 48 | PZ14_DingPostExp_3 | 2 | 18 | 16 | 7 | 0 | 0 | 6 | 3 | 16 |
| 49 | PZ14_DingPostExp_4 | 2 | 16 | 16 | 10 | 0 | 0 | 3 | 1 | 14 |
| 50 | PZ14_DingPostExp_5 | 2 | 19 | 16 | 7 | 0 | 0 | 6 | 4 | 17 |
| 51 | PZ14_DingPostExp_6 | 1 | 23 | 16 | 7 | 0 | 0 | 10 | 5 | 22 |
| 52 | PZ14_DingPostExp_7 | 2 | 18 | 16 | 7 | 0 | 0 | 7 | 2 | 16 |
| 53 | PZ14_DingPostExp_8 | 2 | 19 | 16 | 9 | 0 | 0 | 5 | 3 | 17 |
| 54 | PZ14_DingPostExp_9 | 2 | 19 | 16 | 4 | 0 | 0 | 10 | 3 | 17 |
| 55 | PZ14_DingPreExp_1 | 2 | 16 | 16 | 7 | 0 | 0 | 6 | 1 | 14 |
| 56 | PZ14_DingPreExp_2 | 3 | 17 | 16 | 5 | 0 | 0 | 8 | 1 | 14 |
| 57 | PZ14_DingPreExp_3 | 2 | 19 | 17 | 10 | 0 | 0 | 4 | 3 | 17 |
| 58 | PZ14_DingPreExp_4 | 2 | 16 | 16 | 6 | 0 | 0 | 7 | 1 | 14 |
| 59 | PZ14_DingPreExp_5 | 2 | 19 | 16 | 9 | 0 | 0 | 5 | 3 | 17 |
| 60 | PZ14_DingPreExp_6 | 2 | 20 | 16 | 8 | 0 | 0 | 7 | 3 | 18 |
| 61 | PZ14_DingPreExp_7 | 2 | 17 | 16 | 9 | 0 | 0 | 4 | 2 | 15 |
| 62 | PZ14_DingPreExp_8 | 3 | 18 | 16 | 2 | 0 | 0 | 11 | 2 | 15 |
| | Average | 2.29 | 21.18 | 16.60 | 9.26 | 0.00 | 0.00 | 7.37 | 2.26 | 18.89 |
| | Standard deviation | 0.66 | 3.12 | 1.19 | 2.33 | 0.00 | 0.00 | 3.06 | 1.24 | 3.21 |
| | Median | 2.00 | 21.00 | 16.00 | 9.00 | 0.00 | 0.00 | 7.00 | 2.00 | 18.50 |

**Table S62**. X3DNA report on structures in Puzzle 14b.

| | | | Base pairs | | Dinucleotide steps | | | | | |
|---|---|---|---|---|---|---|---|---|---|---|
| | | | | | **Right-handed** | | **Left-handed** | **Other** | | |
| **No.** | **3D RNA model** | **#Helices** | **#all bps** | **#canonical / wobble bps** | **#A-RNA** | **#B-RNA** | **#Z-RNA** | **#Unclassified steps** | **#Steps without a continuous backbone** | **Total** |
| 1 | PZ14_refstructure_free | 2 | 21 | 17 | 9 | 0 | 0 | 8 | 2 | 19 |
| 2 | PZ14_BujnickiPostExp_1 | 3 | 22 | 19 | 12 | 0 | 0 | 5 | 2 | 19 |
| 3 | PZ14_BujnickiPostExp_2 | 2 | 23 | 17 | 12 | 0 | 0 | 5 | 4 | 21 |
| 4 | PZ14_BujnickiPostExp_3 | 5 | 23 | 22 | 11 | 0 | 0 | 6 | 1 | 18 |
| 5 | PZ14_BujnickiPostExp_4 | 2 | 20 | 17 | 10 | 0 | 0 | 6 | 2 | 18 |
| 6 | PZ14_BujnickiPreExp_1 | 2 | 21 | 17 | 12 | 0 | 0 | 5 | 2 | 19 |
| 7 | PZ14_BujnickiPreExp_2 | 2 | 21 | 17 | 11 | 0 | 0 | 6 | 2 | 19 |
| 8 | PZ14_BujnickiPreExp_3 | 3 | 21 | 17 | 11 | 0 | 0 | 6 | 1 | 18 |
| 9 | PZ14_BujnickiPreExp_4 | 4 | 23 | 22 | 13 | 0 | 0 | 5 | 1 | 19 |
| 10 | PZ14_ChenPostExp_1 | 2 | 20 | 16 | 9 | 0 | 0 | 5 | 4 | 18 |
| 11 | PZ14_ChenPostExp_10 | 3 | 19 | 17 | 12 | 0 | 0 | 2 | 2 | 16 |
| 12 | PZ14_ChenPostExp_2 | 3 | 18 | 16 | 10 | 0 | 0 | 5 | 0 | 15 |
| 13 | PZ14_ChenPostExp_3 | 3 | 20 | 16 | 13 | 0 | 0 | 3 | 1 | 17 |
| 14 | PZ14_ChenPostExp_4 | 3 | 20 | 16 | 12 | 0 | 0 | 4 | 1 | 17 |
| 15 | PZ14_ChenPostExp_5 | 3 | 17 | 16 | 13 | 0 | 0 | 1 | 0 | 14 |
| 16 | PZ14_ChenPostExp_6 | 3 | 17 | 16 | 11 | 0 | 0 | 3 | 0 | 14 |
| 17 | PZ14_ChenPostExp_7 | 2 | 18 | 17 | 13 | 0 | 0 | 1 | 2 | 16 |
| 18 | PZ14_ChenPostExp_8 | 3 | 20 | 16 | 13 | 0 | 0 | 2 | 2 | 17 |
| 19 | PZ14_ChenPostExp_9 | 3 | 17 | 16 | 13 | 0 | 0 | 1 | 0 | 14 |
| 20 | PZ14_DasPostExp_1 | 2 | 22 | 17 | 7 | 0 | 0 | 11 | 2 | 20 |
| 21 | PZ14_DasPostExp_10 | 2 | 25 | 17 | 10 | 0 | 0 | 11 | 2 | 23 |
| 22 | PZ14_DasPostExp_2 | 2 | 24 | 16 | 9 | 0 | 0 | 11 | 2 | 22 |
| 23 | PZ14_DasPostExp_3 | 2 | 25 | 18 | 9 | 0 | 0 | 12 | 2 | 23 |
| 24 | PZ14_DasPostExp_4 | 2 | 25 | 17 | 9 | 0 | 0 | 12 | 2 | 23 |
| 25 | PZ14_DasPostExp_5 | 2 | 25 | 16 | 9 | 0 | 0 | 12 | 2 | 23 |
| 26 | PZ14_DasPostExp_6 | 2 | 25 | 17 | 9 | 0 | 0 | 12 | 2 | 23 |

| | | | | | | | | | | |
|---|---|---|---|---|---|---|---|---|---|---|
| 27 | PZ14_DasPostExp_7 | 2 | 23 | 17 | 9 | 0 | 0 | 10 | 2 | 21 |
| 28 | PZ14_DasPostExp_8 | 2 | 23 | 17 | 9 | 0 | 0 | 10 | 2 | 21 |
| 29 | PZ14_DasPostExp_9 | 2 | 25 | 17 | 10 | 0 | 0 | 11 | 2 | 23 |
| 30 | PZ14_DasPreExp_1 | 2 | 25 | 17 | 9 | 0 | 0 | 12 | 2 | 23 |
| 31 | PZ14_DasPreExp_10 | 2 | 25 | 17 | 10 | 0 | 0 | 11 | 2 | 23 |
| 32 | PZ14_DasPreExp_2 | 2 | 23 | 17 | 9 | 0 | 0 | 10 | 2 | 21 |
| 33 | PZ14_DasPreExp_3 | 2 | 25 | 17 | 8 | 0 | 0 | 13 | 2 | 23 |
| 34 | PZ14_DasPreExp_4 | 2 | 23 | 17 | 9 | 0 | 0 | 10 | 2 | 21 |
| 35 | PZ14_DasPreExp_5 | 2 | 25 | 17 | 10 | 0 | 0 | 11 | 2 | 23 |
| 36 | PZ14_DasPreExp_6 | 2 | 22 | 17 | 7 | 0 | 0 | 11 | 2 | 20 |
| 37 | PZ14_DasPreExp_7 | 2 | 24 | 17 | 9 | 0 | 0 | 11 | 2 | 22 |
| 38 | PZ14_DasPreExp_8 | 2 | 23 | 17 | 9 | 0 | 0 | 10 | 2 | 21 |
| 39 | PZ14_DasPreExp_9 | 2 | 23 | 17 | 9 | 0 | 0 | 10 | 2 | 21 |
| 40 | PZ14_DingPostExp_1 | 2 | 20 | 17 | 11 | 0 | 0 | 4 | 3 | 18 |
| 41 | PZ14_DingPostExp_10 | 2 | 17 | 17 | 9 | 0 | 0 | 5 | 1 | 15 |
| 42 | PZ14_DingPostExp_2 | 2 | 19 | 16 | 7 | 0 | 0 | 6 | 4 | 17 |
| 43 | PZ14_DingPostExp_3 | 2 | 20 | 18 | 8 | 0 | 0 | 7 | 3 | 18 |
| 44 | PZ14_DingPostExp_4 | 2 | 17 | 17 | 10 | 0 | 0 | 4 | 1 | 15 |
| 45 | PZ14_DingPostExp_5 | 2 | 17 | 17 | 7 | 0 | 0 | 7 | 1 | 15 |
| 46 | PZ14_DingPostExp_6 | 2 | 19 | 17 | 7 | 0 | 0 | 9 | 1 | 17 |
| 47 | PZ14_DingPostExp_7 | 3 | 21 | 17 | 10 | 0 | 0 | 5 | 3 | 18 |
| 48 | PZ14_DingPostExp_8 | 2 | 19 | 17 | 6 | 0 | 0 | 9 | 2 | 17 |
| 49 | PZ14_DingPostExp_9 | 2 | 17 | 17 | 5 | 0 | 0 | 9 | 1 | 15 |
| 50 | PZ14_DingPreExp_1 | 2 | 19 | 17 | 10 | 0 | 0 | 5 | 2 | 17 |
| 51 | PZ14_DingPreExp_2 | 2 | 18 | 17 | 10 | 0 | 0 | 4 | 2 | 16 |
| 52 | PZ14_DingPreExp_3 | 2 | 18 | 17 | 7 | 0 | 0 | 8 | 1 | 16 |
| 53 | PZ14_DingPreExp_4 | 2 | 21 | 17 | 6 | 0 | 0 | 11 | 2 | 19 |
| | Average | 2.30 | 21.19 | 17.06 | 9.66 | 0.00 | 0.00 | 7.42 | 1.81 | 18.89 |
| | Standard deviation | 0.61 | 2.74 | 1.13 | 2.00 | 0.00 | 0.00 | 3.48 | 0.88 | 2.89 |
| | Median | 2.00 | 21.00 | 17.00 | 9.00 | 0.00 | 0.00 | 7.00 | 2.00 | 19.00 |

**Table S63**. X3DNA report on structures in Puzzle 15.

| | | | Base pairs | | Dinucleotide steps | | | | | |
|---|---|---|---|---|---|---|---|---|---|---|
| | | | | | Right-handed | | Left-handed | Other | | |
| No. | 3D RNA model | #Helices | #all bps | #canonical / wobble bps | #A-RNA | #B-RNA | #Z-RNA | #Unclassified steps | #Steps without a continuous backbone | Total |
| 1 | PZ15_3dRNA2_1 | 3 | 23 | 22 | 7 | 0 | 0 | 11 | 2 | 20 |
| 2 | PZ15_3dRNA2_10 | 3 | 23 | 22 | 5 | 0 | 0 | 13 | 2 | 20 |
| 3 | PZ15_3dRNA2_2 | 3 | 23 | 22 | 7 | 0 | 0 | 11 | 2 | 20 |
| 4 | PZ15_3dRNA2_3 | 3 | 23 | 22 | 7 | 0 | 0 | 11 | 2 | 20 |
| 5 | PZ15_3dRNA2_4 | 3 | 23 | 22 | 6 | 0 | 0 | 12 | 2 | 20 |
| 6 | PZ15_3dRNA2_5 | 3 | 23 | 22 | 7 | 0 | 0 | 11 | 2 | 20 |
| 7 | PZ15_3dRNA2_6 | 3 | 23 | 22 | 7 | 0 | 0 | 11 | 2 | 20 |
| 8 | PZ15_3dRNA2_7 | 3 | 23 | 22 | 6 | 0 | 0 | 12 | 2 | 20 |
| 9 | PZ15_3dRNA2_8 | 3 | 23 | 22 | 5 | 0 | 0 | 13 | 2 | 20 |
| 10 | PZ15_3dRNA2_9 | 3 | 23 | 22 | 6 | 0 | 0 | 12 | 2 | 20 |
| 11 | PZ15_Adamiak_1 | 3 | 24 | 24 | 12 | 0 | 0 | 7 | 2 | 21 |
| 12 | PZ15_Adamiak_10 | 3 | 27 | 25 | 18 | 0 | 0 | 1 | 5 | 24 |
| 13 | PZ15_Adamiak_2 | 2 | 28 | 24 | 18 | 0 | 0 | 3 | 5 | 26 |
| 14 | PZ15_Adamiak_3 | 2 | 29 | 24 | 17 | 0 | 0 | 3 | 7 | 27 |
| 15 | PZ15_Adamiak_4 | 3 | 25 | 25 | 19 | 0 | 0 | 0 | 3 | 22 |
| 16 | PZ15_Adamiak_5 | 3 | 26 | 25 | 19 | 0 | 0 | 0 | 4 | 23 |
| 17 | PZ15_Adamiak_6 | 2 | 26 | 24 | 17 | 0 | 0 | 3 | 4 | 24 |
| 18 | PZ15_Adamiak_7 | 2 | 25 | 25 | 19 | 0 | 0 | 0 | 4 | 23 |
| 19 | PZ15_Adamiak_8 | 2 | 28 | 25 | 16 | 0 | 0 | 4 | 6 | 26 |
| 20 | PZ15_Adamiak_9 | 3 | 23 | 23 | 15 | 0 | 0 | 4 | 1 | 20 |
| 21 | PZ15_Chen_1 | 2 | 26 | 23 | 16 | 0 | 0 | 4 | 4 | 24 |
| 22 | PZ15_Chen_10 | 3 | 25 | 22 | 16 | 0 | 0 | 2 | 4 | 22 |
| 23 | PZ15_Chen_2 | 3 | 24 | 23 | 18 | 0 | 0 | 1 | 2 | 21 |
| 24 | PZ15_Chen_3 | 2 | 28 | 24 | 16 | 0 | 0 | 5 | 5 | 26 |
| 25 | PZ15_Chen_4 | 2 | 26 | 22 | 9 | 0 | 0 | 11 | 4 | 24 |
| 26 | PZ15_Chen_5 | 3 | 24 | 22 | 18 | 0 | 0 | 1 | 2 | 21 |

| | | | | | | | | | | |
|---|---|---|---|---|---|---|---|---|---|---|
| 27 | PZ15_Chen_6 | 4 | 24 | 22 | 13 | 0 | 0 | 5 | 2 | 20 |
| 28 | PZ15_Chen_7 | 5 | 22 | 21 | 15 | 0 | 0 | 2 | 0 | 17 |
| 29 | PZ15_Chen_8 | 2 | 24 | 21 | 14 | 0 | 0 | 3 | 5 | 22 |
| 30 | PZ15_Chen_9 | 4 | 22 | 19 | 6 | 0 | 0 | 10 | 2 | 18 |
| 31 | PZ15_FARFAR1_1 | 3 | 27 | 23 | 19 | 0 | 0 | 4 | 1 | 24 |
| 32 | PZ15_FARFAR1_10 | 2 | 28 | 23 | 19 | 0 | 0 | 4 | 3 | 26 |
| 33 | PZ15_FARFAR1_2 | 3 | 25 | 23 | 19 | 0 | 0 | 2 | 1 | 22 |
| 34 | PZ15_FARFAR1_3 | 2 | 25 | 24 | 19 | 0 | 0 | 1 | 3 | 23 |
| 35 | PZ15_FARFAR1_4 | 3 | 29 | 23 | 20 | 0 | 0 | 4 | 2 | 26 |
| 36 | PZ15_FARFAR1_5 | 2 | 27 | 23 | 19 | 0 | 0 | 3 | 3 | 25 |
| 37 | PZ15_FARFAR1_6 | 3 | 25 | 23 | 19 | 0 | 0 | 1 | 2 | 22 |
| 38 | PZ15_FARFAR1_7 | 2 | 27 | 23 | 19 | 0 | 0 | 3 | 3 | 25 |
| 39 | PZ15_FARFAR1_8 | 3 | 29 | 24 | 19 | 0 | 0 | 6 | 1 | 26 |
| 40 | PZ15_FARFAR1_9 | 2 | 27 | 23 | 19 | 0 | 0 | 4 | 2 | 25 |
| 41 | PZ15_FARFAR2_1 | 2 | 28 | 24 | 20 | 0 | 0 | 2 | 4 | 26 |
| 42 | PZ15_FARFAR2_10 | 2 | 27 | 24 | 19 | 0 | 0 | 3 | 3 | 25 |
| 43 | PZ15_FARFAR2_2 | 3 | 28 | 24 | 19 | 0 | 0 | 5 | 1 | 25 |
| 44 | PZ15_FARFAR2_3 | 2 | 29 | 24 | 21 | 0 | 0 | 4 | 2 | 27 |
| 45 | PZ15_FARFAR2_4 | 4 | 26 | 24 | 19 | 0 | 0 | 2 | 1 | 22 |
| 46 | PZ15_FARFAR2_5 | 2 | 27 | 24 | 19 | 0 | 0 | 4 | 2 | 25 |
| 47 | PZ15_FARFAR2_6 | 3 | 29 | 24 | 18 | 0 | 0 | 6 | 2 | 26 |
| 48 | PZ15_FARFAR2_7 | 3 | 27 | 24 | 20 | 0 | 0 | 3 | 1 | 24 |
| 49 | PZ15_FARFAR2_8 | 2 | 28 | 24 | 19 | 0 | 0 | 4 | 3 | 26 |
| 50 | PZ15_FARFAR2_9 | 3 | 29 | 24 | 19 | 0 | 0 | 4 | 3 | 26 |
| 51 | PZ15_RNAComposer1_1 | 2 | 28 | 24 | 18 | 0 | 0 | 3 | 5 | 26 |
| 52 | PZ15_RNAComposer1_2 | 2 | 29 | 24 | 17 | 0 | 0 | 3 | 7 | 27 |
| 53 | PZ15_RNAComposer2_1 | 3 | 25 | 25 | 19 | 0 | 0 | 0 | 3 | 22 |
| 54 | PZ15_RNAComposer2_2 | 3 | 26 | 25 | 19 | 0 | 0 | 0 | 4 | 23 |
| 55 | PZ15_SimRNA1_1 | 3 | 27 | 25 | 8 | 0 | 0 | 12 | 4 | 24 |
| 56 | PZ15_SimRNA1_10 | 3 | 29 | 24 | 14 | 0 | 0 | 10 | 2 | 26 |
| 57 | PZ15_SimRNA1_2 | 2 | 26 | 23 | 6 | 0 | 0 | 14 | 4 | 24 |
| 58 | PZ15_SimRNA1_3 | 2 | 27 | 25 | 6 | 0 | 0 | 11 | 8 | 25 |
| 59 | PZ15_SimRNA1_4 | 4 | 27 | 22 | 9 | 0 | 0 | 11 | 3 | 23 |

| | | | | | | | | | | |
|---|---|---|---|---|---|---|---|---|---|---|
| 60 | PZ15_SimRNA1_5 | 3 | 27 | 23 | 9 | 0 | 0 | 12 | 3 | 24 |
| 61 | PZ15_SimRNA1_6 | 1 | 26 | 24 | 6 | 0 | 0 | 11 | 8 | 25 |
| 62 | PZ15_SimRNA1_7 | 2 | 24 | 22 | 9 | 0 | 0 | 8 | 5 | 22 |
| 63 | PZ15_SimRNA1_8 | 2 | 28 | 23 | 11 | 0 | 0 | 12 | 3 | 26 |
| 64 | PZ15_SimRNA1_9 | 4 | 27 | 23 | 11 | 0 | 0 | 11 | 1 | 23 |
| 65 | PZ15_SimRNA2_1 | 2 | 26 | 24 | 10 | 0 | 0 | 12 | 2 | 24 |
| 66 | PZ15_SimRNA2_10 | 2 | 26 | 24 | 6 | 0 | 0 | 16 | 2 | 24 |
| 67 | PZ15_SimRNA2_2 | 2 | 27 | 24 | 13 | 0 | 0 | 9 | 3 | 25 |
| 68 | PZ15_SimRNA2_3 | 2 | 28 | 24 | 10 | 0 | 0 | 13 | 3 | 26 |
| 69 | PZ15_SimRNA2_4 | 3 | 26 | 24 | 7 | 0 | 0 | 15 | 1 | 23 |
| 70 | PZ15_SimRNA2_5 | 2 | 29 | 25 | 11 | 0 | 0 | 13 | 3 | 27 |
| 71 | PZ15_SimRNA2_6 | 3 | 28 | 24 | 11 | 0 | 0 | 13 | 1 | 25 |
| 72 | PZ15_SimRNA2_7 | 2 | 29 | 24 | 11 | 0 | 0 | 12 | 4 | 27 |
| 73 | PZ15_SimRNA2_8 | 3 | 29 | 24 | 10 | 0 | 0 | 14 | 2 | 26 |
| 74 | PZ15_SimRNA2_9 | 2 | 28 | 23 | 7 | 0 | 0 | 17 | 2 | 26 |
| 75 | PZ15_refstructure | 2 | 21 | 17 | 13 | 0 | 0 | 2 | 4 | 19 |
| | Average | 2.63 | 26.08 | 23.25 | 13.72 | 0.00 | 0.00 | 6.79 | 2.95 | 23.45 |
| | Standard deviation | 0.69 | 2.17 | 1.38 | 5.26 | 0.00 | 0.00 | 4.86 | 1.65 | 2.51 |
| | Median | 3.00 | 26.00 | 24.00 | 15.00 | 0.00 | 0.00 | 5.00 | 3.00 | 24.00 |

**Table S64**. X3DNA report on structures in Puzzle 17.

| | | | Base pairs | | Dinucleotide steps | | | | | |
|---|---|---|---|---|---|---|---|---|---|---|
| | | | | | **Right-handed** | | **Left-handed** | **Other** | | |
| **No.** | **3D RNA model** | **#Helices** | **#all bps** | **#canonical / wobble bps** | **#A-RNA** | **#B-RNA** | **#Z-RNA** | **#Unclassified steps** | **#Steps without a continuous backbone** | **Total** |
| 1 | PZ17_refstructure | 2 | 23 | 19 | 8 | 0 | 0 | 9 | 4 | 21 |
| 2 | PZ17_Adamiak_1 | 5 | 23 | 21 | 6 | 0 | 0 | 12 | 0 | 18 |
| 3 | PZ17_Adamiak_2 | 5 | 22 | 21 | 6 | 0 | 0 | 10 | 1 | 17 |
| 4 | PZ17_Adamiak_3 | 5 | 22 | 20 | 7 | 0 | 0 | 10 | 0 | 17 |
| 5 | PZ17_Bujnicki_1 | 2 | 23 | 23 | 15 | 0 | 0 | 2 | 4 | 21 |
| 6 | PZ17_Bujnicki_10 | 2 | 23 | 21 | 4 | 0 | 0 | 13 | 4 | 21 |

| | | | | | | | | | | |
|---|---|---|---|---|---|---|---|---|---|---|
| 7 | PZ17_Bujnicki_2 | 2 | 23 | 21 | 4 | 0 | 0 | 13 | 4 | 21 |
| 8 | PZ17_Bujnicki_3 | 2 | 22 | 21 | 6 | 0 | 0 | 10 | 4 | 20 |
| 9 | PZ17_Bujnicki_4 | 2 | 22 | 21 | 9 | 0 | 0 | 7 | 4 | 20 |
| 10 | PZ17_Bujnicki_5 | 2 | 22 | 22 | 11 | 0 | 0 | 5 | 4 | 20 |
| 11 | PZ17_Bujnicki_6 | 2 | 23 | 21 | 5 | 1 | 0 | 11 | 4 | 21 |
| 12 | PZ17_Bujnicki_7 | 3 | 20 | 20 | 12 | 0 | 0 | 4 | 1 | 17 |
| 13 | PZ17_Bujnicki_8 | 3 | 21 | 19 | 11 | 0 | 0 | 5 | 2 | 18 |
| 14 | PZ17_Bujnicki_9 | 2 | 23 | 23 | 15 | 0 | 0 | 2 | 4 | 21 |
| 15 | PZ17_Chen_1 | 2 | 19 | 18 | 5 | 0 | 0 | 10 | 2 | 17 |
| 16 | PZ17_Chen_10 | 1 | 24 | 16 | 5 | 0 | 0 | 14 | 4 | 23 |
| 17 | PZ17_Chen_2 | 2 | 22 | 20 | 5 | 0 | 0 | 12 | 3 | 20 |
| 18 | PZ17_Chen_3 | 3 | 16 | 16 | 3 | 0 | 0 | 9 | 1 | 13 |
| 19 | PZ17_Chen_4 | 1 | 20 | 17 | 6 | 0 | 0 | 9 | 4 | 19 |
| 20 | PZ17_Chen_5 | 6 | 21 | 19 | 5 | 0 | 0 | 9 | 1 | 15 |
| 21 | PZ17_Chen_6 | 3 | 20 | 17 | 2 | 0 | 0 | 11 | 4 | 17 |
| 22 | PZ17_Chen_7 | 3 | 19 | 19 | 12 | 0 | 0 | 2 | 2 | 16 |
| 23 | PZ17_Chen_8 | 2 | 19 | 18 | 12 | 0 | 0 | 3 | 2 | 17 |
| 24 | PZ17_Chen_9 | 3 | 23 | 20 | 7 | 0 | 0 | 7 | 6 | 20 |
| 25 | PZ17_Das_1 | 2 | 23 | 20 | 15 | 0 | 0 | 3 | 3 | 21 |
| 26 | PZ17_Das_10 | 4 | 23 | 21 | 14 | 0 | 0 | 3 | 2 | 19 |
| 27 | PZ17_Das_2 | 3 | 23 | 20 | 15 | 0 | 0 | 3 | 2 | 20 |
| 28 | PZ17_Das_3 | 3 | 20 | 19 | 12 | 0 | 0 | 3 | 2 | 17 |
| 29 | PZ17_Das_4 | 2 | 22 | 21 | 14 | 0 | 0 | 3 | 3 | 20 |
| 30 | PZ17_Das_5 | 2 | 21 | 21 | 14 | 0 | 0 | 2 | 3 | 19 |
| 31 | PZ17_Das_6 | 2 | 21 | 21 | 15 | 0 | 0 | 1 | 3 | 19 |
| 32 | PZ17_Das_7 | 4 | 23 | 19 | 14 | 0 | 0 | 3 | 2 | 19 |
| 33 | PZ17_Das_8 | 2 | 22 | 21 | 15 | 0 | 0 | 1 | 4 | 20 |
| 34 | PZ17_Das_9 | 4 | 24 | 20 | 15 | 0 | 0 | 2 | 3 | 20 |
| 35 | PZ17_DasExtraInfo_1 | 2 | 22 | 21 | 14 | 0 | 0 | 3 | 3 | 20 |
| 36 | PZ17_DasExtraInfo_2 | 2 | 23 | 21 | 14 | 0 | 0 | 4 | 3 | 21 |
| 37 | PZ17_DasExtraInfo_3 | 2 | 22 | 21 | 15 | 0 | 0 | 2 | 3 | 20 |
| 38 | PZ17_Ding_1 | 1 | 19 | 19 | 9 | 0 | 0 | 6 | 3 | 18 |
| 39 | PZ17_Ding_10 | 4 | 19 | 19 | 6 | 0 | 0 | 8 | 1 | 15 |

| | | | | | | | | | | |
|---|---|---|---|---|---|---|---|---|---|---|
| 40 | PZ17_Ding_2 | 1 | 19 | 19 | 3 | 0 | 0 | 12 | 3 | 18 |
| 41 | PZ17_Ding_3 | 1 | 19 | 19 | 7 | 0 | 0 | 8 | 3 | 18 |
| 42 | PZ17_Ding_4 | 1 | 20 | 18 | 10 | 0 | 0 | 5 | 4 | 19 |
| 43 | PZ17_Ding_5 | 1 | 19 | 18 | 9 | 0 | 0 | 6 | 3 | 18 |
| 44 | PZ17_Ding_6 | 4 | 22 | 20 | 5 | 0 | 0 | 11 | 2 | 18 |
| 45 | PZ17_Ding_7 | 3 | 21 | 20 | 8 | 0 | 0 | 6 | 4 | 18 |
| 46 | PZ17_Ding_8 | 4 | 23 | 20 | 8 | 0 | 0 | 8 | 3 | 19 |
| 47 | PZ17_Ding_9 | 2 | 22 | 21 | 7 | 0 | 0 | 8 | 5 | 20 |
| 48 | PZ17_Dohkolyan_1 | 2 | 17 | 17 | 8 | 0 | 0 | 6 | 1 | 15 |
| 49 | PZ17_Dohkolyan_2 | 3 | 20 | 17 | 4 | 0 | 0 | 10 | 3 | 17 |
| 50 | PZ17_Dohkolyan_3 | 3 | 22 | 20 | 8 | 0 | 0 | 9 | 2 | 19 |
| 51 | PZ17_Major_1 | 3 | 24 | 19 | 1 | 0 | 0 | 18 | 2 | 21 |
| 52 | PZ17_Major_10 | 2 | 13 | 6 | 0 | 0 | 0 | 5 | 6 | 11 |
| 53 | PZ17_Major_2 | 4 | 25 | 24 | 1 | 0 | 0 | 18 | 2 | 21 |
| 54 | PZ17_Major_3 | 3 | 25 | 23 | 1 | 0 | 0 | 18 | 3 | 22 |
| 55 | PZ17_Major_4 | 3 | 25 | 24 | 1 | 1 | 0 | 17 | 3 | 22 |
| 56 | PZ17_Major_5 | 3 | 24 | 21 | 0 | 1 | 0 | 17 | 3 | 21 |
| 57 | PZ17_Major_6 | 3 | 16 | 12 | 0 | 0 | 0 | 10 | 3 | 13 |
| 58 | PZ17_Major_7 | 4 | 12 | 8 | 0 | 0 | 0 | 8 | 0 | 8 |
| 59 | PZ17_Major_8 | 3 | 16 | 9 | 0 | 0 | 0 | 9 | 4 | 13 |
| 60 | PZ17_Major_9 | 2 | 8 | 5 | 1 | 0 | 0 | 4 | 1 | 6 |
| 61 | PZ17_RNAComposer1_1 | 2 | 18 | 17 | 10 | 0 | 0 | 5 | 1 | 16 |
| 62 | PZ17_RNAComposer1_10 | 2 | 17 | 16 | 7 | 0 | 0 | 7 | 1 | 15 |
| 63 | PZ17_RNAComposer1_2 | 1 | 24 | 17 | 3 | 0 | 0 | 18 | 2 | 23 |
| 64 | PZ17_RNAComposer1_3 | 1 | 19 | 14 | 7 | 0 | 0 | 8 | 3 | 18 |
| 65 | PZ17_RNAComposer1_4 | 2 | 17 | 16 | 7 | 0 | 0 | 7 | 1 | 15 |
| 66 | PZ17_RNAComposer1_5 | 1 | 21 | 16 | 6 | 0 | 0 | 10 | 4 | 20 |
| 67 | PZ17_RNAComposer1_6 | 3 | 17 | 15 | 8 | 0 | 0 | 6 | 0 | 14 |
| 68 | PZ17_RNAComposer1_7 | 2 | 22 | 17 | 5 | 0 | 0 | 12 | 3 | 20 |
| 69 | PZ17_RNAComposer1_8 | 3 | 17 | 15 | 8 | 0 | 0 | 6 | 0 | 14 |
| 70 | PZ17_RNAComposer1_9 | 2 | 16 | 16 | 7 | 0 | 0 | 6 | 1 | 14 |
| 71 | PZ17_RNAComposer2_1 | 5 | 22 | 21 | 6 | 0 | 0 | 11 | 0 | 17 |
| 72 | PZ17_RNAComposer2_10 | 3 | 15 | 15 | 6 | 0 | 0 | 6 | 0 | 12 |

| | | | | | | | | | | |
|---|---|---|---|---|---|---|---|---|---|---|
| 73 | PZ17_RNAComposer2_2 | 5 | 22 | 21 | 8 | 0 | 0 | 9 | 0 | 17 |
| 74 | PZ17_RNAComposer2_3 | 4 | 19 | 13 | 0 | 0 | 0 | 11 | 4 | 15 |
| 75 | PZ17_RNAComposer2_4 | 4 | 21 | 20 | 6 | 0 | 0 | 10 | 1 | 17 |
| 76 | PZ17_RNAComposer2_5 | 4 | 21 | 20 | 6 | 0 | 0 | 10 | 1 | 17 |
| 77 | PZ17_RNAComposer2_6 | 5 | 21 | 20 | 6 | 0 | 0 | 10 | 0 | 16 |
| 78 | PZ17_RNAComposer2_7 | 5 | 19 | 19 | 8 | 0 | 0 | 6 | 0 | 14 |
| 79 | PZ17_RNAComposer2_8 | 4 | 21 | 18 | 6 | 0 | 0 | 10 | 1 | 17 |
| 80 | PZ17_RNAComposer2_9 | 3 | 15 | 14 | 6 | 0 | 0 | 6 | 0 | 12 |
| 81 | PZ17_SimRNA1_1 | 1 | 22 | 19 | 8 | 0 | 0 | 8 | 5 | 21 |
| 82 | PZ17_SimRNA1_2 | 2 | 22 | 19 | 11 | 0 | 0 | 4 | 5 | 20 |
| 83 | PZ17_SimRNA1_3 | 3 | 22 | 19 | 9 | 0 | 0 | 7 | 3 | 19 |
| 84 | PZ17_SimRNA1_4 | 3 | 21 | 19 | 9 | 0 | 0 | 6 | 3 | 18 |
| 85 | PZ17_SimRNA1_5 | 3 | 20 | 19 | 10 | 0 | 0 | 5 | 2 | 17 |
| 86 | PZ17_SimRNA1_6 | 2 | 22 | 19 | 11 | 0 | 0 | 5 | 4 | 20 |
| 87 | PZ17_SimRNA1_7 | 1 | 24 | 20 | 9 | 0 | 0 | 7 | 7 | 23 |
| 88 | PZ17_SimRNA1_8 | 1 | 21 | 19 | 8 | 0 | 0 | 7 | 5 | 20 |
| 89 | PZ17_SimRNA1_9 | 2 | 20 | 19 | 5 | 0 | 0 | 9 | 4 | 18 |
| 90 | PZ17_SimRNA2_1 | 3 | 23 | 22 | 13 | 0 | 0 | 4 | 3 | 20 |
| 91 | PZ17_SimRNA2_2 | 3 | 25 | 24 | 12 | 0 | 0 | 6 | 4 | 22 |
| 92 | PZ17_SimRNA2_3 | 2 | 22 | 21 | 10 | 0 | 0 | 7 | 3 | 20 |
| 93 | PZ17_SimRNA2_4 | 4 | 26 | 24 | 9 | 0 | 0 | 9 | 4 | 22 |
| 94 | PZ17_SimRNA2_5 | 2 | 23 | 21 | 10 | 0 | 0 | 7 | 4 | 21 |
| 95 | PZ17_SimRNA2_6 | 3 | 23 | 20 | 8 | 0 | 0 | 9 | 3 | 20 |
| 96 | PZ17_SimRNA2_7 | 2 | 22 | 22 | 11 | 0 | 0 | 6 | 3 | 20 |
| 97 | PZ17_SimRNA2_8 | 2 | 24 | 20 | 9 | 0 | 0 | 8 | 5 | 22 |
| 98 | PZ17_SimRNA2_9 | 2 | 24 | 22 | 8 | 0 | 0 | 8 | 6 | 22 |
| 99 | PZ17_Xiao_1 | 3 | 15 | 12 | 1 | 0 | 0 | 8 | 3 | 12 |
| 100 | PZ17_Xiao_10 | 3 | 12 | 9 | 1 | 0 | 0 | 6 | 2 | 9 |
| 101 | PZ17_Xiao_2 | 4 | 15 | 12 | 1 | 0 | 0 | 9 | 1 | 11 |
| 102 | PZ17_Xiao_3 | 2 | 15 | 11 | 1 | 0 | 0 | 4 | 8 | 13 |
| 103 | PZ17_Xiao_4 | 4 | 11 | 6 | 0 | 0 | 0 | 2 | 5 | 7 |
| 104 | PZ17_Xiao_5 | 3 | 12 | 10 | 3 | 0 | 0 | 4 | 2 | 9 |
| 105 | PZ17_Xiao_6 | 3 | 14 | 10 | 0 | 0 | 0 | 8 | 3 | 11 |

| | | | | | | | | | | |
|---|---|---|---|---|---|---|---|---|---|---|
| 106 | PZ17_Xiao_7 | 3 | 12 | 9 | 0 | 0 | 0 | 5 | 4 | 9 |
| 107 | PZ17_Xiao_8 | 4 | 11 | 7 | 1 | 0 | 0 | 2 | 4 | 7 |
| 108 | PZ17_Xiao_9 | 3 | 13 | 10 | 0 | 0 | 0 | 6 | 4 | 10 |
| | Average | 2.71 | 20.13 | 17.97 | 7.11 | 0.03 | 0.00 | 7.49 | 2.79 | 17.42 |
| | Standard deviation | 1.12 | 3.69 | 4.26 | 4.48 | 0.17 | 0.00 | 3.89 | 1.64 | 3.91 |
| | Median | 3.00 | 21.00 | 19.00 | 7.00 | 0.00 | 0.00 | 7.00 | 3.00 | 18.00 |

**Table S65**. X3DNA report on structures in Puzzle 18.

| | | | Base pairs | | Dinucleotide steps | | | | | |
|---|---|---|---|---|---|---|---|---|---|---|
| | | | | | **Right-handed** | | **Left-handed** | **Other** | | |
| **No.** | **3D RNA model** | **#Helices** | **#all bps** | **#canonical / wobble bps** | **#A-RNA** | **#B-RNA** | **#Z-RNA** | **#Unclassified steps** | **#Steps without a continuous backbone** | **Total** |
| 1 | PZ18_refstructure | 2 | 29 | 25 | 12 | 0 | 0 | 9 | 6 | 27 |
| 2 | PZ18_3dRNA_1 | 2 | 29 | 25 | 4 | 0 | 0 | 20 | 3 | 27 |
| 3 | PZ18_3dRNA_2 | 3 | 29 | 25 | 4 | 0 | 0 | 20 | 2 | 26 |
| 4 | PZ18_3dRNA_3 | 3 | 27 | 24 | 5 | 0 | 0 | 17 | 2 | 24 |
| 5 | PZ18_3dRNA_4 | 2 | 28 | 25 | 1 | 0 | 0 | 22 | 3 | 26 |
| 6 | PZ18_3dRNA_5 | 2 | 29 | 25 | 2 | 0 | 0 | 22 | 3 | 27 |
| 7 | PZ18_Chen_1 | 2 | 28 | 27 | 0 | 1 | 0 | 21 | 4 | 26 |
| 8 | PZ18_Chen_2 | 2 | 28 | 25 | 1 | 1 | 0 | 20 | 4 | 26 |
| 9 | PZ18_Chen_3 | 2 | 29 | 25 | 4 | 0 | 0 | 20 | 3 | 27 |
| 10 | PZ18_Chen_4 | 2 | 28 | 24 | 2 | 0 | 0 | 20 | 4 | 26 |
| 11 | PZ18_Chen_5 | 3 | 29 | 25 | 20 | 0 | 0 | 4 | 2 | 26 |
| 12 | PZ18_Das_1 | 2 | 31 | 25 | 11 | 0 | 0 | 12 | 6 | 29 |
| 13 | PZ18_Das_2 | 2 | 30 | 25 | 11 | 0 | 0 | 11 | 6 | 28 |
| 14 | PZ18_Das_3 | 2 | 30 | 25 | 11 | 0 | 0 | 11 | 6 | 28 |
| 15 | PZ18_Das_4 | 2 | 30 | 26 | 11 | 0 | 0 | 13 | 4 | 28 |
| 16 | PZ18_Das_5 | 4 | 26 | 21 | 8 | 0 | 0 | 11 | 3 | 22 |
| 17 | PZ18_Ding_1 | 2 | 30 | 26 | 9 | 0 | 0 | 14 | 5 | 28 |
| 18 | PZ18_Ding_2 | 2 | 27 | 26 | 11 | 0 | 0 | 11 | 3 | 25 |

| | | | | | | | | | | |
|---|---|---|---|---|---|---|---|---|---|---|
| 19 | PZ18_Ding_3 | 2 | 29 | 26 | 10 | 0 | 0 | 12 | 5 | 27 |
| 20 | PZ18_Ding_4 | 1 | 27 | 23 | 9 | 0 | 0 | 12 | 5 | 26 |
| 21 | PZ18_Ding_5 | 2 | 28 | 26 | 10 | 0 | 0 | 13 | 3 | 26 |
| 22 | PZ18_Dokholyan_1 | 3 | 27 | 26 | 9 | 0 | 0 | 12 | 3 | 24 |
| 23 | PZ18_Dokholyan_2 | 4 | 28 | 26 | 8 | 0 | 0 | 14 | 2 | 24 |
| 24 | PZ18_Dokholyan_3 | 4 | 27 | 26 | 7 | 0 | 0 | 13 | 3 | 23 |
| 25 | PZ18_FARFAR_1 | 2 | 30 | 26 | 18 | 0 | 0 | 7 | 3 | 28 |
| 26 | PZ18_FARFAR_10 | 3 | 30 | 26 | 18 | 0 | 0 | 6 | 3 | 27 |
| 27 | PZ18_FARFAR_2 | 2 | 30 | 26 | 18 | 0 | 0 | 7 | 3 | 28 |
| 28 | PZ18_FARFAR_3 | 2 | 30 | 26 | 18 | 0 | 0 | 7 | 3 | 28 |
| 29 | PZ18_FARFAR_4 | 2 | 29 | 26 | 18 | 0 | 0 | 6 | 3 | 27 |
| 30 | PZ18_FARFAR_5 | 2 | 30 | 26 | 18 | 0 | 0 | 7 | 3 | 28 |
| 31 | PZ18_FARFAR_6 | 2 | 29 | 26 | 18 | 0 | 0 | 6 | 3 | 27 |
| 32 | PZ18_FARFAR_7 | 2 | 30 | 26 | 18 | 0 | 0 | 6 | 4 | 28 |
| 33 | PZ18_FARFAR_8 | 2 | 30 | 26 | 18 | 0 | 0 | 7 | 3 | 28 |
| 34 | PZ18_FARFAR_9 | 3 | 28 | 26 | 18 | 0 | 0 | 5 | 2 | 25 |
| 35 | PZ18_Lee_1 | 3 | 30 | 27 | 14 | 0 | 0 | 11 | 2 | 27 |
| 36 | PZ18_Lee_2 | 2 | 28 | 26 | 13 | 0 | 0 | 9 | 4 | 26 |
| 37 | PZ18_Lee_3 | 2 | 25 | 23 | 7 | 0 | 0 | 10 | 6 | 23 |
| 38 | PZ18_Lee_4 | 2 | 26 | 26 | 15 | 0 | 0 | 6 | 3 | 24 |
| 39 | PZ18_Lee_5 | 2 | 25 | 25 | 15 | 0 | 0 | 5 | 3 | 23 |
| 40 | PZ18_LeeServer_1 | 2 | 29 | 25 | 14 | 0 | 0 | 10 | 3 | 27 |
| 41 | PZ18_LeeServer_2 | 3 | 26 | 25 | 10 | 0 | 0 | 10 | 3 | 23 |
| 42 | PZ18_LeeServer_3 | 3 | 24 | 22 | 5 | 0 | 0 | 13 | 3 | 21 |
| 43 | PZ18_LeeServer_4 | 3 | 27 | 26 | 14 | 0 | 0 | 7 | 3 | 24 |
| 44 | PZ18_LeeServer_5 | 2 | 26 | 25 | 15 | 0 | 0 | 5 | 4 | 24 |
| 45 | PZ18_RNAComposer_1 | 2 | 29 | 26 | 15 | 0 | 0 | 9 | 3 | 27 |
| 46 | PZ18_RNAComposer_2 | 3 | 28 | 23 | 11 | 0 | 0 | 10 | 4 | 25 |
| 47 | PZ18_RNAComposer_3 | 3 | 29 | 26 | 13 | 0 | 0 | 10 | 3 | 26 |
| 48 | PZ18_RNAComposer_4 | 3 | 29 | 26 | 13 | 0 | 0 | 11 | 2 | 26 |
| 49 | PZ18_RNAComposer_5 | 4 | 28 | 25 | 11 | 0 | 0 | 12 | 1 | 24 |
| 50 | PZ18_SimRNA_1 | 1 | 29 | 24 | 9 | 0 | 0 | 11 | 8 | 28 |
| 51 | PZ18_SimRNA_2 | 2 | 28 | 26 | 9 | 0 | 0 | 13 | 4 | 26 |

| | | | | | | | | | | |
|---|---|---|---|---|---|---|---|---|---|---|
| 52 | PZ18_SimRNA_3 | 2 | 29 | 26 | 8 | 0 | 0 | 15 | 4 | 27 |
| 53 | PZ18_YagoubAli_1 | 3 | 29 | 26 | 15 | 0 | 0 | 9 | 2 | 26 |
| | Average | 2.38 | 28.36 | 25.28 | 11.06 | 0.04 | 0.00 | 11.40 | 3.49 | 25.98 |
| | Standard deviation | 0.69 | 1.53 | 1.17 | 5.33 | 0.19 | 0.00 | 4.87 | 1.32 | 1.82 |
| | Median | 2.00 | 29.00 | 26.00 | 11.00 | 0.00 | 0.00 | 11.00 | 3.00 | 26.00 |

**Table S66**. X3DNA report on structures in Puzzle 19.

| | | | Base pairs | | Dinucleotide steps | | | | | |
|---|---|---|---|---|---|---|---|---|---|---|
| | | | | | **Right-handed** | | **Left-handed** | **Other** | | |
| **No.** | **3D RNA model** | **#Helices** | **#all bps** | **#canonical / wobble bps** | **#A-RNA** | **#B-RNA** | **#Z-RNA** | **#Unclassified steps** | **#Steps without a continuous backbone** | **Total** |
| 1 | PZ19_refstructure | 2 | 27 | 23 | 12 | 0 | 0 | 6 | 7 | 25 |
| 2 | PZ19_3dRNA_1 | 3 | 24 | 22 | 1 | 0 | 0 | 17 | 3 | 21 |
| 3 | PZ19_3dRNA_2 | 2 | 24 | 22 | 1 | 0 | 0 | 16 | 5 | 22 |
| 4 | PZ19_3dRNA_3 | 3 | 25 | 21 | 2 | 2 | 0 | 13 | 5 | 22 |
| 5 | PZ19_3dRNA_4 | 2 | 24 | 21 | 1 | 0 | 0 | 17 | 4 | 22 |
| 6 | PZ19_3dRNA_5 | 3 | 23 | 22 | 1 | 0 | 0 | 15 | 4 | 20 |
| 7 | PZ19_Adamiak_1 | 2 | 28 | 24 | 14 | 0 | 0 | 9 | 3 | 26 |
| 8 | PZ19_Adamiak_2 | 2 | 25 | 21 | 12 | 0 | 0 | 9 | 2 | 23 |
| 9 | PZ19_Adamiak_3 | 3 | 24 | 19 | 11 | 0 | 0 | 6 | 4 | 21 |
| 10 | PZ19_Adamiak_4 | 2 | 27 | 21 | 13 | 0 | 0 | 8 | 4 | 25 |
| 11 | PZ19_Adamiak_5 | 3 | 24 | 19 | 10 | 0 | 0 | 7 | 4 | 21 |
| 12 | PZ19_Bujnicki_1 | 2 | 27 | 25 | 15 | 0 | 0 | 5 | 5 | 25 |
| 13 | PZ19_Bujnicki_2 | 2 | 27 | 25 | 16 | 0 | 0 | 4 | 5 | 25 |
| 14 | PZ19_Bujnicki_3 | 2 | 25 | 24 | 11 | 0 | 0 | 6 | 6 | 23 |
| 15 | PZ19_Bujnicki_4 | 2 | 27 | 25 | 10 | 0 | 0 | 10 | 5 | 25 |
| 16 | PZ19_Bujnicki_5 | 2 | 27 | 26 | 16 | 0 | 0 | 4 | 5 | 25 |
| 17 | PZ19_Chen_1 | 2 | 24 | 22 | 11 | 1 | 0 | 5 | 5 | 22 |
| 18 | PZ19_Chen_2 | 2 | 24 | 21 | 5 | 0 | 0 | 12 | 5 | 22 |
| 19 | PZ19_Chen_3 | 2 | 23 | 20 | 6 | 0 | 0 | 11 | 4 | 21 |
| 20 | PZ19_Chen_4 | 3 | 24 | 23 | 8 | 0 | 0 | 10 | 3 | 21 |

| | | | | | | | | | | |
|---|---|---|---|---|---|---|---|---|---|---|
| 21 | PZ19_Chen_5 | 1 | 21 | 17 | 0 | 2 | 0 | 13 | 5 | 20 |
| 22 | PZ19_Das_1 | 4 | 25 | 24 | 16 | 0 | 0 | 1 | 4 | 21 |
| 23 | PZ19_Das_2 | 2 | 26 | 21 | 15 | 0 | 0 | 4 | 5 | 24 |
| 24 | PZ19_Das_3 | 3 | 24 | 20 | 16 | 0 | 0 | 1 | 4 | 21 |
| 25 | PZ19_Das_4 | 3 | 22 | 20 | 16 | 0 | 0 | 1 | 2 | 19 |
| 26 | PZ19_Das_5 | 2 | 27 | 24 | 14 | 0 | 0 | 7 | 4 | 25 |
| 27 | PZ19_Ding_1 | 2 | 26 | 23 | 10 | 0 | 0 | 9 | 5 | 24 |
| 28 | PZ19_Ding_2 | 3 | 23 | 22 | 11 | 0 | 0 | 7 | 2 | 20 |
| 29 | PZ19_Ding_3 | 2 | 24 | 23 | 11 | 0 | 0 | 8 | 3 | 22 |
| 30 | PZ19_Ding_4 | 3 | 20 | 16 | 0 | 0 | 0 | 15 | 2 | 17 |
| 31 | PZ19_Ding_5 | 4 | 18 | 14 | 0 | 0 | 0 | 10 | 4 | 14 |
| 32 | PZ19_Dokholyan_1 | 3 | 22 | 22 | 10 | 0 | 0 | 7 | 2 | 19 |
| 33 | PZ19_FARFAR_1 | 2 | 26 | 20 | 12 | 0 | 0 | 9 | 3 | 24 |
| 34 | PZ19_FARFAR_10 | 3 | 26 | 20 | 5 | 0 | 0 | 14 | 4 | 23 |
| 35 | PZ19_FARFAR_2 | 2 | 26 | 20 | 11 | 0 | 0 | 11 | 2 | 24 |
| 36 | PZ19_FARFAR_3 | 2 | 27 | 20 | 12 | 0 | 0 | 10 | 3 | 25 |
| 37 | PZ19_FARFAR_4 | 2 | 25 | 21 | 11 | 0 | 0 | 7 | 5 | 23 |
| 38 | PZ19_FARFAR_5 | 2 | 26 | 20 | 11 | 0 | 0 | 9 | 4 | 24 |
| 39 | PZ19_FARFAR_6 | 1 | 25 | 21 | 11 | 0 | 0 | 9 | 4 | 24 |
| 40 | PZ19_FARFAR_7 | 1 | 25 | 20 | 12 | 0 | 0 | 8 | 4 | 24 |
| 41 | PZ19_FARFAR_8 | 2 | 26 | 20 | 11 | 0 | 0 | 10 | 3 | 24 |
| 42 | PZ19_FARFAR_9 | 2 | 27 | 20 | 12 | 0 | 0 | 10 | 3 | 25 |
| 43 | PZ19_LeeServer_1 | 3 | 24 | 22 | 0 | 0 | 0 | 19 | 2 | 21 |
| 44 | PZ19_LeeServer_2 | 2 | 24 | 22 | 0 | 0 | 0 | 17 | 5 | 22 |
| 45 | PZ19_LeeServer_3 | 2 | 28 | 24 | 0 | 0 | 0 | 22 | 4 | 26 |
| 46 | PZ19_LeeServer_4 | 3 | 20 | 17 | 7 | 0 | 0 | 9 | 1 | 17 |
| 47 | PZ19_LeeServer_5 | 3 | 18 | 17 | 9 | 0 | 0 | 5 | 1 | 15 |
| 48 | PZ19_RNAComposer_1 | 2 | 25 | 21 | 10 | 0 | 0 | 9 | 4 | 23 |
| 49 | PZ19_RNAComposer_2 | 2 | 26 | 23 | 5 | 0 | 0 | 15 | 4 | 24 |
| 50 | PZ19_RNAComposer_3 | 2 | 27 | 24 | 7 | 0 | 0 | 14 | 4 | 25 |
| 51 | PZ19_RNAComposer_4 | 3 | 26 | 24 | 10 | 0 | 0 | 10 | 3 | 23 |
| 52 | PZ19_RNAComposer_5 | 3 | 25 | 23 | 9 | 0 | 0 | 10 | 3 | 22 |
| 53 | PZ19_SimRNA_1 | 2 | 26 | 25 | 11 | 0 | 0 | 7 | 6 | 24 |

| | | | | | | | | | | |
|---|---|---|---|---|---|---|---|---|---|---|
| 54 | PZ19_SimRNA_2 | 3 | 25 | 23 | 11 | 0 | 0 | 6 | 5 | 22 |
| 55 | PZ19_SimRNA_3 | 4 | 25 | 25 | 11 | 0 | 0 | 7 | 3 | 21 |
| | Average | 2.38 | 24.71 | 21.53 | 8.96 | 0.09 | 0.00 | 9.45 | 3.82 | 22.33 |
| | Standard deviation | 0.68 | 2.23 | 2.47 | 5.00 | 0.40 | 0.00 | 4.50 | 1.26 | 2.60 |
| | Median | 2.00 | 25.00 | 22.00 | 11.00 | 0.00 | 0.00 | 9.00 | 4.00 | 23.00 |

**Table S67**. X3DNA report on structures in Puzzle 20.

| | | | Base pairs | | Dinucleotide steps | | | | | |
|---|---|---|---|---|---|---|---|---|---|---|
| | | | | | **Right-handed** | | **Left-handed** | **Other** | | |
| **No.** | **3D RNA model** | **#Helices** | **#all bps** | **#canonical / wobble bps** | **#A-RNA** | **#B-RNA** | **#Z-RNA** | **#Unclassified steps** | **#Steps without a continuous backbone** | **Total** |
| 1 | PZ20_Adamiak_1 | 2 | 30 | 25 | 14 | 0 | 0 | 10 | 4 | 28 |
| 2 | PZ20_Adamiak_2 | 3 | 29 | 25 | 11 | 0 | 0 | 11 | 4 | 26 |
| 3 | PZ20_Adamiak_3 | 2 | 30 | 25 | 14 | 0 | 0 | 10 | 4 | 28 |
| 4 | PZ20_Adamiak_4 | 2 | 30 | 25 | 14 | 0 | 0 | 10 | 4 | 28 |
| 5 | PZ20_Adamiak_5 | 2 | 30 | 25 | 9 | 0 | 0 | 15 | 4 | 28 |
| 6 | PZ20_Bujnicki_1 | 2 | 29 | 23 | 16 | 0 | 0 | 4 | 7 | 27 |
| 7 | PZ20_Bujnicki_2 | 2 | 29 | 23 | 16 | 0 | 0 | 4 | 7 | 27 |
| 8 | PZ20_Bujnicki_3 | 4 | 28 | 23 | 15 | 0 | 0 | 5 | 4 | 24 |
| 9 | PZ20_Bujnicki_4 | 2 | 27 | 23 | 12 | 0 | 0 | 6 | 7 | 25 |
| 10 | PZ20_Bujnicki_5 | 2 | 27 | 23 | 8 | 0 | 0 | 12 | 5 | 25 |
| 11 | PZ20_Das_1 | 4 | 26 | 21 | 16 | 0 | 0 | 2 | 4 | 22 |
| 12 | PZ20_Das_2 | 2 | 28 | 23 | 15 | 0 | 0 | 5 | 6 | 26 |
| 13 | PZ20_Das_3 | 4 | 30 | 24 | 16 | 0 | 0 | 6 | 4 | 26 |
| 14 | PZ20_Das_4 | 4 | 26 | 22 | 16 | 0 | 0 | 1 | 5 | 22 |
| 15 | PZ20_Das_5 | 5 | 24 | 21 | 16 | 0 | 0 | 1 | 2 | 19 |
| 16 | PZ20_FARFAR_1 | 2 | 30 | 26 | 17 | 0 | 0 | 7 | 4 | 28 |
| 17 | PZ20_FARFAR_10 | 2 | 30 | 26 | 17 | 0 | 0 | 7 | 4 | 28 |
| 18 | PZ20_FARFAR_2 | 2 | 30 | 25 | 17 | 0 | 0 | 7 | 4 | 28 |
| 19 | PZ20_FARFAR_3 | 2 | 29 | 26 | 17 | 0 | 0 | 7 | 3 | 27 |
| 20 | PZ20_FARFAR_4 | 2 | 30 | 26 | 18 | 0 | 0 | 6 | 4 | 28 |

| | | | | | | | | | | |
|---|---|---|---|---|---|---|---|---|---|---|
| 21 | PZ20_FARFAR_5 | 2 | 30 | 26 | 17 | 0 | 0 | 7 | 4 | 28 |
| 22 | PZ20_FARFAR_6 | 2 | 31 | 25 | 17 | 0 | 0 | 8 | 4 | 29 |
| 23 | PZ20_FARFAR_7 | 2 | 31 | 26 | 17 | 0 | 0 | 8 | 4 | 29 |
| 24 | PZ20_FARFAR_8 | 2 | 30 | 27 | 16 | 0 | 0 | 6 | 6 | 28 |
| 25 | PZ20_FARFAR_9 | 2 | 31 | 25 | 18 | 0 | 0 | 7 | 4 | 29 |
| 26 | PZ20_RNAComposer_1 | 2 | 30 | 24 | 11 | 0 | 0 | 13 | 4 | 28 |
| 27 | PZ20_RNAComposer_2 | 2 | 29 | 25 | 13 | 0 | 0 | 9 | 5 | 27 |
| 28 | PZ20_RNAComposer_3 | 4 | 27 | 24 | 13 | 0 | 0 | 7 | 3 | 23 |
| 29 | PZ20_RNAComposer_4 | 2 | 28 | 24 | 10 | 0 | 0 | 11 | 5 | 26 |
| 30 | PZ20_RNAComposer_5 | 3 | 28 | 23 | 13 | 0 | 0 | 9 | 3 | 25 |
| 31 | PZ20_SimRNA_1 | 2 | 28 | 24 | 1 | 0 | 0 | 17 | 8 | 26 |
| 32 | PZ20_SimRNA_2 | 2 | 25 | 21 | 9 | 0 | 0 | 7 | 7 | 23 |
| 33 | PZ20_SimRNA_3 | 2 | 27 | 26 | 8 | 0 | 0 | 6 | 11 | 25 |
| 34 | PZ20_SimRNA_4 | 3 | 26 | 25 | 12 | 0 | 0 | 6 | 5 | 23 |
| 35 | PZ20_SimRNA_5 | 1 | 26 | 23 | 10 | 0 | 0 | 6 | 9 | 25 |
| 36 | PZ20_Xiao3_1 | 4 | 20 | 16 | 4 | 0 | 0 | 10 | 2 | 16 |
| 37 | PZ20_Xiao3_2 | 3 | 18 | 13 | 3 | 0 | 0 | 11 | 1 | 15 |
| 38 | PZ20_Xiao3_3 | 3 | 21 | 13 | 3 | 0 | 0 | 11 | 4 | 18 |
| 39 | PZ20_Xiao3_4 | 4 | 20 | 15 | 2 | 0 | 0 | 10 | 4 | 16 |
| 40 | PZ20_Xiao3_5 | 4 | 21 | 17 | 2 | 0 | 0 | 11 | 4 | 17 |
| 41 | PZ20_refstructure | 2 | 30 | 24 | 17 | 0 | 0 | 4 | 7 | 28 |
| | Average | 2.56 | 27.54 | 23.07 | 12.44 | 0.00 | 0.00 | 7.80 | 4.73 | 24.98 |
| | Standard deviation | 0.92 | 3.34 | 3.49 | 5.04 | 0.00 | 0.00 | 3.44 | 1.91 | 3.93 |
| | Median | 2.00 | 29.00 | 24.00 | 14.00 | 0.00 | 0.00 | 7.00 | 4.00 | 26.00 |

**Table S68**. X3DNA report on structures in Puzzle 21.

| | | | Base pairs | | Dinucleotide steps | | | | | |
|---|---|---|---|---|---|---|---|---|---|---|
| | | | | | **Right-handed** | | **Left-handed** | **Other** | | **Total** |
| **No.** | **3D RNA model** | **#Helices** | **#all bps** | **#canonical / wobble bps** | **#A-RNA** | **#B-RNA** | **#Z-RNA** | **#Unclassified steps** | **#Steps without a continuous backbone** | |
| 1 | PZ21_3dRNA_1 | 1 | 4 | 4 | 3 | 0 | 0 | 0 | 0 | 3 |
| 2 | PZ21_3dRNA_2 | 1 | 5 | 4 | 0 | 0 | 0 | 3 | 1 | 4 |
| 3 | PZ21_3dRNA_3 | 1 | 4 | 4 | 0 | 0 | 0 | 3 | 0 | 3 |
| 4 | PZ21_3dRNA_4 | 2 | 12 | 9 | 0 | 0 | 0 | 9 | 1 | 10 |
| 5 | PZ21_3dRNA_5 | 2 | 15 | 11 | 0 | 0 | 0 | 11 | 2 | 13 |
| 6 | PZ21_Adamiak_1 | 2 | 12 | 12 | 4 | 0 | 0 | 5 | 1 | 10 |
| 7 | PZ21_Adamiak_2 | 2 | 12 | 12 | 6 | 0 | 0 | 3 | 1 | 10 |
| 8 | PZ21_Adamiak_3 | 2 | 11 | 11 | 5 | 0 | 0 | 4 | 0 | 9 |
| 9 | PZ21_Adamiak_4 | 2 | 12 | 11 | 6 | 0 | 0 | 3 | 1 | 10 |
| 10 | PZ21_Adamiak_5 | 2 | 11 | 11 | 6 | 0 | 0 | 3 | 0 | 9 |
| 11 | PZ21_Bujnicki_1 | 2 | 13 | 10 | 3 | 0 | 0 | 6 | 2 | 11 |
| 12 | PZ21_Bujnicki_2 | 1 | 13 | 11 | 8 | 0 | 0 | 2 | 2 | 12 |
| 13 | PZ21_Bujnicki_3 | 2 | 17 | 15 | 2 | 0 | 0 | 10 | 3 | 15 |
| 14 | PZ21_Bujnicki_4 | 1 | 12 | 11 | 9 | 0 | 0 | 1 | 1 | 11 |
| 15 | PZ21_Bujnicki_5 | 1 | 10 | 9 | 2 | 0 | 0 | 4 | 3 | 9 |
| 16 | PZ21_ChenHighLig_1 | 1 | 13 | 11 | 3 | 0 | 0 | 8 | 1 | 12 |
| 17 | PZ21_ChenHighLig_2 | 1 | 13 | 11 | 6 | 0 | 0 | 5 | 1 | 12 |
| 18 | PZ21_ChenHighLig_3 | 2 | 13 | 11 | 7 | 0 | 0 | 2 | 2 | 11 |
| 19 | PZ21_ChenHighLig_4 | 1 | 13 | 10 | 5 | 0 | 0 | 4 | 3 | 12 |
| 20 | PZ21_ChenHighLig_5 | 2 | 14 | 11 | 2 | 0 | 0 | 9 | 1 | 12 |
| 21 | PZ21_ChenLowLig_1 | 1 | 13 | 11 | 3 | 0 | 0 | 8 | 1 | 12 |
| 22 | PZ21_ChenLowLig_2 | 1 | 13 | 11 | 6 | 0 | 0 | 5 | 1 | 12 |
| 23 | PZ21_ChenLowLig_3 | 2 | 13 | 11 | 7 | 0 | 0 | 2 | 2 | 11 |
| 24 | PZ21_ChenLowLig_4 | 1 | 13 | 10 | 5 | 0 | 0 | 4 | 3 | 12 |
| 25 | PZ21_ChenLowLig_5 | 2 | 14 | 11 | 2 | 0 | 0 | 9 | 1 | 12 |
| 26 | PZ21_Das_1 | 1 | 13 | 11 | 10 | 0 | 0 | 1 | 1 | 12 |

| | | | | | | | | | | |
|---|---|---|---|---|---|---|---|---|---|---|
| 27 | PZ21_Das_2 | 1 | 12 | 11 | 10 | 0 | 0 | 0 | 1 | 11 |
| 28 | PZ21_Das_3 | 1 | 13 | 11 | 8 | 0 | 0 | 2 | 2 | 12 |
| 29 | PZ21_Das_4 | 2 | 13 | 12 | 9 | 0 | 0 | 0 | 2 | 11 |
| 30 | PZ21_Das_5 | 1 | 13 | 11 | 9 | 0 | 0 | 1 | 2 | 12 |
| 31 | PZ21_DasLORES_1 | 1 | 13 | 11 | 9 | 0 | 0 | 1 | 2 | 12 |
| 32 | PZ21_DasLORES_2 | 1 | 13 | 11 | 10 | 0 | 0 | 1 | 1 | 12 |
| 33 | PZ21_DasLORES_3 | 2 | 15 | 13 | 9 | 0 | 0 | 0 | 4 | 13 |
| 34 | PZ21_DasLORES_4 | 1 | 12 | 12 | 9 | 0 | 0 | 0 | 2 | 11 |
| 35 | PZ21_DasLORES_5 | 2 | 15 | 12 | 9 | 0 | 0 | 1 | 3 | 13 |
| 36 | PZ21_FARFAR_1 | 2 | 16 | 12 | 8 | 0 | 0 | 3 | 3 | 14 |
| 37 | PZ21_FARFAR_10 | 2 | 17 | 12 | 8 | 0 | 0 | 3 | 4 | 15 |
| 38 | PZ21_FARFAR_2 | 3 | 17 | 12 | 8 | 0 | 0 | 4 | 2 | 14 |
| 39 | PZ21_FARFAR_3 | 1 | 17 | 11 | 9 | 0 | 0 | 5 | 2 | 16 |
| 40 | PZ21_FARFAR_4 | 2 | 16 | 12 | 9 | 0 | 0 | 2 | 3 | 14 |
| 41 | PZ21_FARFAR_5 | 1 | 15 | 12 | 8 | 0 | 0 | 3 | 3 | 14 |
| 42 | PZ21_FARFAR_6 | 2 | 16 | 11 | 9 | 0 | 0 | 3 | 2 | 14 |
| 43 | PZ21_FARFAR_7 | 2 | 17 | 12 | 9 | 0 | 0 | 3 | 3 | 15 |
| 44 | PZ21_FARFAR_8 | 1 | 15 | 12 | 8 | 0 | 0 | 2 | 4 | 14 |
| 45 | PZ21_FARFAR_9 | 1 | 17 | 11 | 8 | 0 | 0 | 6 | 2 | 16 |
| 46 | PZ21_RNAComposer_1 | 2 | 13 | 8 | 6 | 0 | 0 | 2 | 3 | 11 |
| 47 | PZ21_RNAComposer_2 | 1 | 12 | 9 | 7 | 0 | 0 | 2 | 2 | 11 |
| 48 | PZ21_RNAComposer_3 | 2 | 11 | 11 | 3 | 0 | 0 | 6 | 0 | 9 |
| 49 | PZ21_RNAComposer_4 | 2 | 11 | 11 | 1 | 0 | 0 | 8 | 0 | 9 |
| 50 | PZ21_RNAComposer_5 | 3 | 14 | 11 | 8 | 0 | 0 | 2 | 1 | 11 |
| 51 | PZ21_Sanbonmatsu_1 | 3 | 11 | 10 | 8 | 0 | 0 | 0 | 0 | 8 |
| 52 | PZ21_Sanbonmatsu_2 | 3 | 11 | 10 | 8 | 0 | 0 | 0 | 0 | 8 |
| 53 | PZ21_Sanbonmatsu_3 | 2 | 13 | 12 | 9 | 0 | 0 | 0 | 2 | 11 |
| 54 | PZ21_Sanbonmatsu_4 | 2 | 13 | 12 | 9 | 0 | 0 | 0 | 2 | 11 |
| 55 | PZ21_SimRNA_1 | 2 | 16 | 15 | 5 | 0 | 0 | 6 | 3 | 14 |
| 56 | PZ21_SimRNA_2 | 1 | 17 | 16 | 4 | 0 | 0 | 8 | 4 | 16 |
| 57 | PZ21_SimRNA_3 | 2 | 17 | 14 | 10 | 0 | 0 | 3 | 2 | 15 |
| 58 | PZ21_SimRNA_4 | 1 | 17 | 16 | 8 | 0 | 0 | 4 | 4 | 16 |
| 59 | PZ21_SimRNA_5 | 2 | 17 | 14 | 10 | 0 | 0 | 3 | 2 | 15 |

| | | | | | | | | | | |
|---|---|---|---|---|---|---|---|---|---|---|
| 60 | PZ21_refstructure | 2 | 13 | 12 | 6 | 0 | 0 | 2 | 3 | 11 |
| | Average | 1.63 | 13.27 | 11.10 | 6.30 | 0.00 | 0.00 | 3.50 | 1.83 | 11.63 |
| | Standard deviation | 0.61 | 2.86 | 2.22 | 3.00 | 0.00 | 0.00 | 2.85 | 1.15 | 2.80 |
| | Median | 2.00 | 13.00 | 11.00 | 7.50 | 0.00 | 0.00 | 3.00 | 2.00 | 12.00 |

**Table S69**. X3DNA report on structures in Puzzle 24.

| | | | Base pairs | | Dinucleotide steps | | | | | |
|---|---|---|---|---|---|---|---|---|---|---|
| | | | | | **Right-handed** | | **Left-handed** | **Other** | | **Total** |
| **No.** | **3D RNA model** | **#Helices** | **#all bps** | **#canonical / wobble bps** | **#A-RNA** | **#B-RNA** | **#Z-RNA** | **#Unclassified steps** | **#Steps without a continuous backbone** | |
| 1 | PZ24_refstructure | 3 | 49 | 46 | 32 | 0 | 0 | 11 | 3 | 46 |
| 2 | PZ24_3dRNA_1 | 4 | 40 | 35 | 19 | 0 | 0 | 15 | 2 | 36 |
| 3 | PZ24_3dRNA_2 | 4 | 40 | 36 | 22 | 0 | 0 | 13 | 1 | 36 |
| 4 | PZ24_3dRNA_3 | 4 | 45 | 39 | 22 | 0 | 0 | 17 | 2 | 41 |
| 5 | PZ24_3dRNA_4 | 3 | 43 | 39 | 26 | 0 | 0 | 12 | 2 | 40 |
| 6 | PZ24_3dRNA_5 | 3 | 44 | 39 | 23 | 0 | 0 | 16 | 2 | 41 |
| 7 | PZ24_Adamiak_1 | 4 | 46 | 42 | 18 | 1 | 0 | 19 | 4 | 42 |
| 8 | PZ24_Adamiak_2 | 4 | 46 | 42 | 14 | 0 | 0 | 24 | 4 | 42 |
| 9 | PZ24_Adamiak_3 | 4 | 46 | 43 | 20 | 0 | 0 | 20 | 2 | 42 |
| 10 | PZ24_Adamiak_4 | 4 | 46 | 44 | 23 | 0 | 0 | 17 | 2 | 42 |
| 11 | PZ24_Adamiak_5 | 4 | 46 | 44 | 28 | 0 | 0 | 12 | 2 | 42 |
| 12 | PZ24_Bujnicki_1 | 3 | 47 | 42 | 15 | 0 | 0 | 24 | 5 | 44 |
| 13 | PZ24_Bujnicki_2 | 3 | 48 | 46 | 23 | 0 | 0 | 16 | 6 | 45 |
| 14 | PZ24_Bujnicki_3 | 3 | 47 | 44 | 23 | 0 | 0 | 16 | 5 | 44 |
| 15 | PZ24_Bujnicki_4 | 4 | 49 | 46 | 19 | 0 | 0 | 21 | 5 | 45 |
| 16 | PZ24_Bujnicki_5 | 3 | 49 | 46 | 29 | 0 | 0 | 11 | 6 | 46 |
| 17 | PZ24_Das_1 | 3 | 50 | 45 | 31 | 0 | 0 | 11 | 5 | 47 |
| 18 | PZ24_Das_10 | 3 | 50 | 45 | 30 | 0 | 0 | 12 | 5 | 47 |
| 19 | PZ24_Das_2 | 3 | 50 | 45 | 31 | 0 | 0 | 11 | 5 | 47 |
| 20 | PZ24_Das_3 | 3 | 50 | 45 | 32 | 0 | 0 | 10 | 5 | 47 |
| 21 | PZ24_Das_4 | 3 | 50 | 45 | 33 | 0 | 0 | 9 | 5 | 47 |

| | | | | | | | | | | |
|---|---|---|---|---|---|---|---|---|---|---|
| 22 | PZ24_Das_5 | 3 | 50 | 45 | 31 | 0 | 0 | 11 | 5 | 47 |
| 23 | PZ24_Das_6 | 3 | 50 | 45 | 28 | 0 | 0 | 14 | 5 | 47 |
| 24 | PZ24_Das_7 | 3 | 50 | 45 | 30 | 0 | 0 | 12 | 5 | 47 |
| 25 | PZ24_Das_8 | 3 | 50 | 45 | 31 | 0 | 0 | 11 | 5 | 47 |
| 26 | PZ24_Das_9 | 3 | 50 | 45 | 29 | 0 | 0 | 13 | 5 | 47 |
| 27 | PZ24_DasTFN_1 | 4 | 50 | 44 | 27 | 0 | 0 | 17 | 2 | 46 |
| 28 | PZ24_DasTFN_10 | 3 | 50 | 43 | 29 | 0 | 0 | 13 | 5 | 47 |
| 29 | PZ24_DasTFN_2 | 3 | 50 | 45 | 25 | 0 | 0 | 19 | 3 | 47 |
| 30 | PZ24_DasTFN_3 | 3 | 50 | 44 | 29 | 0 | 0 | 15 | 3 | 47 |
| 31 | PZ24_DasTFN_4 | 2 | 51 | 44 | 23 | 0 | 0 | 22 | 4 | 49 |
| 32 | PZ24_DasTFN_5 | 2 | 49 | 44 | 30 | 0 | 0 | 13 | 4 | 47 |
| 33 | PZ24_DasTFN_6 | 4 | 50 | 44 | 25 | 0 | 0 | 19 | 2 | 46 |
| 34 | PZ24_DasTFN_7 | 3 | 50 | 44 | 24 | 0 | 0 | 19 | 4 | 47 |
| 35 | PZ24_DasTFN_8 | 3 | 51 | 46 | 25 | 0 | 0 | 20 | 3 | 48 |
| 36 | PZ24_DasTFN_9 | 3 | 51 | 45 | 25 | 0 | 0 | 19 | 4 | 48 |
| 37 | PZ24_Ding_1 | 3 | 26 | 23 | 11 | 0 | 0 | 7 | 5 | 23 |
| 38 | PZ24_Ding_10 | 3 | 31 | 24 | 11 | 0 | 0 | 10 | 7 | 28 |
| 39 | PZ24_Ding_2 | 4 | 30 | 23 | 12 | 0 | 0 | 9 | 5 | 26 |
| 40 | PZ24_Ding_3 | 3 | 31 | 24 | 9 | 0 | 0 | 12 | 7 | 28 |
| 41 | PZ24_Ding_4 | 3 | 30 | 25 | 15 | 0 | 0 | 8 | 4 | 27 |
| 42 | PZ24_Ding_5 | 5 | 30 | 24 | 12 | 0 | 0 | 8 | 5 | 25 |
| 43 | PZ24_Ding_6 | 2 | 28 | 25 | 8 | 0 | 0 | 11 | 7 | 26 |
| 44 | PZ24_Ding_7 | 3 | 29 | 25 | 11 | 0 | 0 | 10 | 5 | 26 |
| 45 | PZ24_Ding_8 | 4 | 26 | 22 | 10 | 0 | 0 | 10 | 2 | 22 |
| 46 | PZ24_Ding_9 | 3 | 26 | 23 | 10 | 0 | 0 | 8 | 5 | 23 |
| 47 | PZ24_FARFAR2_1 | 3 | 50 | 44 | 29 | 0 | 0 | 15 | 3 | 47 |
| 48 | PZ24_FARFAR2_10 | 3 | 49 | 45 | 27 | 0 | 0 | 15 | 4 | 46 |
| 49 | PZ24_FARFAR2_2 | 2 | 52 | 45 | 31 | 0 | 0 | 13 | 6 | 50 |
| 50 | PZ24_FARFAR2_3 | 3 | 50 | 44 | 28 | 0 | 0 | 15 | 4 | 47 |
| 51 | PZ24_FARFAR2_4 | 3 | 50 | 43 | 28 | 0 | 0 | 15 | 4 | 47 |
| 52 | PZ24_FARFAR2_5 | 4 | 49 | 44 | 31 | 0 | 0 | 12 | 2 | 45 |
| 53 | PZ24_FARFAR2_6 | 3 | 51 | 45 | 25 | 0 | 0 | 19 | 4 | 48 |
| 54 | PZ24_FARFAR2_7 | 3 | 49 | 44 | 32 | 0 | 0 | 9 | 5 | 46 |

| | | | | | | | | | | |
|---|---|---|---|---|---|---|---|---|---|---|
| 55 | PZ24_FARFAR2_8 | 3 | 48 | 44 | 28 | 0 | 0 | 13 | 4 | 45 |
| 56 | PZ24_FARFAR2_9 | 3 | 49 | 44 | 28 | 0 | 0 | 15 | 3 | 46 |
| 57 | PZ24_iFoldRNA_1 | 3 | 45 | 43 | 11 | 0 | 0 | 26 | 5 | 42 |
| 58 | PZ24_iFoldRNA_2 | 3 | 45 | 42 | 9 | 0 | 0 | 29 | 4 | 42 |
| 59 | PZ24_iFoldRNA_3 | 4 | 42 | 39 | 11 | 1 | 0 | 21 | 5 | 38 |
| 60 | PZ24_iFoldRNA_4 | 3 | 45 | 37 | 11 | 0 | 0 | 25 | 6 | 42 |
| 61 | PZ24_iFoldRNA_5 | 7 | 40 | 36 | 3 | 2 | 0 | 26 | 2 | 33 |
| 62 | PZ24_Kollmann_1 | 8 | 40 | 36 | 21 | 0 | 0 | 10 | 1 | 32 |
| 63 | PZ24_Kollmann_10 | 7 | 43 | 36 | 17 | 0 | 0 | 13 | 6 | 36 |
| 64 | PZ24_Kollmann_2 | 6 | 42 | 34 | 19 | 0 | 0 | 9 | 8 | 36 |
| 65 | PZ24_Kollmann_3 | 6 | 37 | 30 | 19 | 0 | 0 | 6 | 6 | 31 |
| 66 | PZ24_Kollmann_4 | 7 | 37 | 32 | 22 | 0 | 0 | 4 | 4 | 30 |
| 67 | PZ24_Kollmann_5 | 5 | 43 | 41 | 22 | 0 | 0 | 10 | 6 | 38 |
| 68 | PZ24_Kollmann_6 | 6 | 46 | 39 | 23 | 0 | 0 | 9 | 8 | 40 |
| 69 | PZ24_Kollmann_7 | 8 | 43 | 36 | 18 | 0 | 0 | 12 | 5 | 35 |
| 70 | PZ24_Kollmann_8 | 6 | 39 | 32 | 21 | 0 | 0 | 7 | 5 | 33 |
| 71 | PZ24_Kollmann_9 | 5 | 40 | 35 | 17 | 0 | 0 | 12 | 6 | 35 |
| 72 | PZ24_RNAComposer_1 | 4 | 47 | 44 | 19 | 0 | 0 | 21 | 3 | 43 |
| 73 | PZ24_RNAComposer_2 | 4 | 48 | 43 | 17 | 0 | 0 | 24 | 3 | 44 |
| 74 | PZ24_RNAComposer_3 | 3 | 43 | 41 | 23 | 0 | 0 | 14 | 3 | 40 |
| 75 | PZ24_RNAComposer_4 | 3 | 44 | 42 | 26 | 0 | 0 | 12 | 3 | 41 |
| 76 | PZ24_RNAComposer_5 | 4 | 44 | 42 | 20 | 0 | 0 | 17 | 3 | 40 |
| 77 | PZ24_SimRNA_1 | 3 | 46 | 43 | 17 | 0 | 0 | 17 | 9 | 43 |
| 78 | PZ24_SimRNA_2 | 2 | 43 | 41 | 17 | 0 | 0 | 13 | 11 | 41 |
| 79 | PZ24_SimRNA_3 | 4 | 42 | 41 | 23 | 0 | 0 | 10 | 5 | 38 |
| 80 | PZ24_SimRNA_4 | 6 | 44 | 44 | 11 | 0 | 0 | 22 | 5 | 38 |
| 81 | PZ24_SimRNA_5 | 4 | 42 | 40 | 20 | 0 | 0 | 14 | 4 | 38 |
| 82 | PZ24_Vfold3D_1 | 4 | 47 | 44 | 15 | 1 | 0 | 25 | 2 | 43 |
| 83 | PZ24_Vfold3D_2 | 4 | 46 | 45 | 15 | 0 | 0 | 25 | 2 | 42 |
| 84 | PZ24_Vfold3D_3 | 5 | 47 | 45 | 10 | 0 | 0 | 30 | 2 | 42 |
| 85 | PZ24_Vfold3D_4 | 4 | 48 | 47 | 9 | 3 | 0 | 30 | 2 | 44 |
| 86 | PZ24_Vfold3D_5 | 3 | 46 | 45 | 11 | 1 | 0 | 28 | 3 | 43 |
| 87 | PZ24_VfoldLA_1 | 4 | 43 | 39 | 10 | 0 | 0 | 23 | 6 | 39 |

| | | | | | | | | | | |
|---|---|---|---|---|---|---|---|---|---|---|
| 88 | PZ24_VfoldLA_2 | 4 | 44 | 41 | 12 | 0 | 0 | 15 | 13 | 40 |
| 89 | PZ24_VfoldLA_3 | 5 | 44 | 40 | 10 | 0 | 0 | 17 | 12 | 39 |
| 90 | PZ24_VfoldLA_4 | 4 | 44 | 42 | 14 | 0 | 0 | 18 | 8 | 40 |
| 91 | PZ24_VfoldLA_5 | 6 | 42 | 40 | 10 | 0 | 0 | 14 | 12 | 36 |
| | Average | 3.78 | 44.37 | 40.03 | 20.58 | 0.10 | 0.00 | 15.34 | 4.57 | 40.59 |
| | Standard deviation | 1.28 | 6.57 | 6.75 | 7.64 | 0.42 | 0.00 | 5.87 | 2.32 | 6.98 |
| | Median | 3.00 | 46.00 | 43.00 | 22.00 | 0.00 | 0.00 | 14.00 | 5.00 | 42.00 |

MolProbity reports:

(a) Clash score is the number of serious steric overlaps (> 0.4 Å) per 1000 atoms.
(b) 100th percentile is the best among structures of comparable resolution; 0th percentile is the worst. For clashscore the comparative set of structures was selected in 2004, for MolProbity score in 2006.
(c) RNA backbone was recently shown to be rotameric. Outliers are RNA suites that don't fall into recognized rotamers.

**Table S70**. MolProbity report on the reference structures.

| No. | 3D RNA model | (a) Clash score | (b) Rank-ing | #Bad bonds | #Bad angles | #Chiral handedness swaps | #Tetrahedral geometry outli-ers | #Probably wrong sugar puckers | (c) #Bad back-bone confor-mations | To-tal |
|---|---|---|---|---|---|---|---|---|---|---|
| 1 | PZ01_refstructure | 2.03 | 99 | 5 | 34 | 0 | 0 | 1 | 2 | 42 |
| 2 | PZ02_refstructure | 29.34 | 16 | 7 | 2 | 0 | 0 | 1 | 25 | 35 |
| 3 | PZ03_refstructure | 0.73 | 99 | 0 | 4 | 0 | 0 | 2 | 14 | 20 |
| 4 | PZ04_refstructure | 26.15 | 19 | 0 | 31 | 0 | 2 | 10 | 21 | 64 |
| 5 | PZ05_refstructure | 7.95 | 82 | 0 | 6 | 0 | 2 | 1 | 28 | 37 |
| 6 | PZ06_refstructure | 9.74 | 73 | 0 | 0 | 0 | 0 | 3 | 33 | 36 |
| 7 | PZ07_refstructure | 3.02 | 98 | 0 | 3 | 0 | 0 | 0 | 21 | 24 |
| 8 | PZ08_refstructure | 7.04 | 86 | 1 | 10 | 0 | 0 | 3 | 13 | 27 |
| 9 | PZ09_refstructure | 4.39 | 95 | 0 | 2 | 0 | 0 | 0 | 8 | 10 |
| 10 | PZ10_refstructure | 4.17 | 96 | 0 | 0 | 0 | 0 | 2 | 29 | 31 |
| 11 | PZ11_refstructure | 0.00 | 100 | 1 | 1 | 0 | 0 | 5 | 12 | 19 |
| 12 | PZ12_refstructure | 11.35 | 65 | 0 | 0 | 0 | 0 | 2 | 23 | 25 |
| 13 | PZ13_refstructure | 0.00 | 100 | 0 | 0 | 0 | 0 | 0 | 4 | 4 |
| 14 | PZ14_refstructure_bound | 4.56 | 95 | 1 | 0 | 0 | 0 | 0 | 10 | 11 |
| 15 | PZ14_refstructure_free | 13.38 | 56 | 0 | 0 | 0 | 0 | 1 | 13 | 14 |
| 16 | PZ15_refstructure | 0.91 | 99 | 1 | 0 | 0 | 0 | 0 | 5 | 6 |
| 17 | PZ17_refstructure | 15.59 | 47 | 0 | 3 | 0 | 0 | 1 | 11 | 15 |
| 18 | PZ18_refstructure | 3.48 | 97 | 0 | 0 | 0 | 0 | 0 | 7 | 7 |
| 19 | PZ19_refstructure | 2.50 | 99 | 1 | 6 | 0 | 0 | 1 | 6 | 14 |
| 20 | PZ20_refstructure | 0.46 | 99 | 0 | 0 | 0 | 0 | 0 | 5 | 5 |
| 21 | PZ21_refstructure | 20.29 | 31 | 10 | 2 | 0 | 0 | 0 | 16 | 28 |
| 22 | PZ24_refstructure | 0.28 | 99 | 0 | 0 | 0 | 0 | 1 | 10 | 11 |

| | | | | | | | | | | |
|---|---|---|---|---|---|---|---|---|---|---|
| | Average | 7.61 | 79.55 | 1.23 | 4.73 | 0.00 | 0.18 | 1.55 | 14.36 | 22.05 |
| | Standard deviation | 8.54 | 28.01 | 2.64 | 9.37 | 0.00 | 0.59 | 2.28 | 8.96 | 14.77 |
| | Median | 4.28 | 95.50 | 0.00 | 1.50 | 0.00 | 0.00 | 1.00 | 12.50 | 19.50 |

**Table S71**. MolProbity report on structures in Puzzle 01.

| **No.** | **3D RNA model** | **(a) Clash score** | **(b) Ranking** | **#Bad bonds** | **#Bad angles** | **#Chiral hand-edness swaps** | **#Tetrahedral geometry outli-ers** | **#Probably wrong sugar puckers** | **(c) #Bad backbone conformations** | **To-tal** |
|---|---|---|---|---|---|---|---|---|---|---|
| 1 | PZ01_Das_1 | 0.00 | 100 | 0 | 0 | 0 | 0 | 0 | 3 | 3 |
| 2 | PZ01_Das_4 | 0.00 | 100 | 0 | 0 | 0 | 0 | 0 | 3 | 3 |
| 3 | PZ01_Das_2 | 0.68 | 99 | 0 | 2 | 0 | 0 | 0 | 3 | 5 |
| 4 | PZ01_Das_3 | 0.00 | 100 | 0 | 1 | 0 | 0 | 0 | 4 | 5 |
| 5 | PZ01_Das_5 | 0.00 | 100 | 0 | 4 | 0 | 0 | 0 | 5 | 9 |
| 6 | PZ01_Chen_1 | 0.00 | 100 | 2 | 17 | 0 | 0 | 0 | 2 | 21 |
| 7 | PZ01_Bujnicki_1 | 0.00 | 100 | 0 | 18 | 1 | 0 | 2 | 3 | 24 |
| 8 | PZ01_refstructure | 2.03 | 99 | 5 | 34 | 0 | 0 | 1 | 2 | 42 |
| 9 | PZ01_Dokholyan_1 | 26.33 | 19 | 3 | 51 | 0 | 0 | 0 | 6 | 60 |
| 10 | PZ01_Bujnicki_3 | 47.30 | 5 | 33 | 50 | 0 | 0 | 0 | 0 | 83 |
| 11 | PZ01_Bujnicki_5 | 65.72 | 1 | 29 | 52 | 0 | 0 | 1 | 2 | 84 |
| 12 | PZ01_Bujnicki_4 | 69.11 | 1 | 30 | 61 | 0 | 0 | 0 | 1 | 92 |
| 13 | PZ01_Santalucia_1 | 25.00 | 22 | 65 | 26 | 0 | 0 | 1 | 8 | 100 |
| 14 | PZ01_Bujnicki_2 | 60.09 | 2 | 42 | 74 | 0 | 2 | 1 | 4 | 123 |
| 15 | PZ01_Major_1 | 55.48 | 3 | 84 | 276 | 0 | 0 | 7 | 25 | 392 |
| | Average | 23.45 | 56.73 | 19.53 | 44.40 | 0.07 | 0.13 | 0.87 | 4.73 | 69.73 |
| | Standard deviation | 28.15 | 47.96 | 26.98 | 68.70 | 0.26 | 0.52 | 1.81 | 5.95 | 98.12 |
| | Median | 2.03 | 99.00 | 3.00 | 26.00 | 0.00 | 0.00 | 0.00 | 3.00 | 42.00 |

**Table S72**. MolProbity report on structures in Puzzle 02.

| No. | 3D RNA model | (a) Clash score | (b) Ranking | #Bad bonds | #Bad angles | #Chiral handedness swaps | #Tetrahedral geometry outliers | #Probably wrong sugar puckers | (c) #Bad backbone conformations | Total |
|---|---|---|---|---|---|---|---|---|---|---|
| 1 | PZ02_Das_1 | 14.23 | 53 | 0 | 0 | 0 | 0 | 1 | 23 | 24 |
| 2 | PZ02_Das_2 | 17.33 | 40 | 0 | 1 | 0 | 0 | 1 | 26 | 28 |
| 3 | PZ02_Das_4 | 16.71 | 43 | 1 | 1 | 0 | 0 | 1 | 26 | 29 |
| 4 | PZ02_Das_3 | 17.33 | 40 | 0 | 0 | 0 | 0 | 2 | 31 | 33 |
| 5 | PZ02_refstructure | 29.34 | 16 | 7 | 2 | 0 | 0 | 1 | 25 | 35 |
| 6 | PZ02_Das_5 | 17.95 | 38 | 2 | 3 | 0 | 0 | 1 | 29 | 35 |
| 7 | PZ02_Santalucia_1 | 18.29 | 37 | 36 | 22 | 0 | 0 | 0 | 2 | 60 |
| 8 | PZ02_Bujnicki_3 | 0.31 | 99 | 0 | 71 | 0 | 0 | 3 | 17 | 91 |
| 9 | PZ02_Bujnicki_2 | 12.69 | 60 | 0 | 71 | 0 | 0 | 3 | 25 | 99 |
| 10 | PZ02_Dokholyan_1 | 12.07 | 62 | 6 | 98 | 0 | 0 | 4 | 34 | 142 |
| 11 | PZ02_Bujnicki_1 | 10.83 | 67 | 4 | 103 | 1 | 4 | 6 | 43 | 161 |
| 12 | PZ02_Wildauer_1 | 153.39 | 0 | 23 | 63 | 0 | 0 | 15 | 68 | 169 |
| 13 | PZ02_Chen_1 | 21.05 | 30 | 22 | 135 | 2 | 3 | 5 | 28 | 195 |
| 14 | PZ02_Major_1 | 125.58 | 0 | 140 | 182 | 0 | 12 | 0 | 64 | 398 |
| | Average | 33.36 | 41.79 | 17.21 | 53.71 | 0.21 | 1.36 | 3.07 | 31.50 | 107.07 |
| | Standard deviation | 45.73 | 26.48 | 37.07 | 59.30 | 0.58 | 3.32 | 3.89 | 17.19 | 103.03 |
| | Median | 17.33 | 40.00 | 3.00 | 42.50 | 0.00 | 0.00 | 1.50 | 27.00 | 75.50 |

**Table S73**. MolProbity report on structures in Puzzle 03.

| No. | 3D RNA model | (a) Clash score | (b) Rank-ing | #Bad bonds | #Bad angles | #Chiral handedness swaps | #Tetrahedral geometry outli-ers | #Probably wrong sugar puck-ers | (c) #Bad back-bone confor-mations | Total |
|---|---|---|---|---|---|---|---|---|---|---|
| 1 | PZ03_Das_1 | 0.00 | 100 | 1 | 3 | 0 | 0 | 1 | 5 | 10 |
| 2 | PZ03_Das_2 | 0.74 | 99 | 1 | 2 | 0 | 0 | 0 | 9 | 12 |
| 3 | PZ03_Das_3 | 1.10 | 99 | 0 | 4 | 0 | 0 | 0 | 12 | 16 |
| 4 | PZ03_Das_5 | 1.10 | 99 | 2 | 2 | 0 | 0 | 2 | 10 | 16 |
| 5 | PZ03_Das_4 | 2.21 | 99 | 1 | 6 | 0 | 0 | 2 | 11 | 20 |
| 6 | PZ03_refstructure | 0.73 | 99 | 0 | 4 | 0 | 0 | 2 | 14 | 20 |
| 7 | PZ03_Bujnicki_2 | 3.68 | 97 | 0 | 38 | 0 | 0 | 1 | 7 | 46 |
| 8 | PZ03_Chen_1 | 0.00 | 100 | 1 | 74 | 0 | 0 | 2 | 22 | 99 |
| 9 | PZ03_Bujnicki_1 | 9.20 | 75 | 1 | 82 | 0 | 0 | 2 | 19 | 104 |
| 10 | PZ03_Dokholyan_2 | 32.01 | 13 | 1 | 108 | 0 | 0 | 7 | 30 | 146 |
| 11 | PZ03_Dokholyan_1 | 32.74 | 13 | 1 | 113 | 0 | 0 | 8 | 28 | 150 |
| 12 | PZ03_Major_2 | 76.16 | 0 | 167 | 546 | 1 | 1 | 35 | 62 | 812 |
| 13 | PZ03_Major_1 | 61.83 | 2 | 176 | 617 | 0 | 3 | 24 | 65 | 885 |
|  | Average | 17.04 | 68.85 | 27.08 | 123.00 | 0.08 | 0.31 | 6.62 | 22.62 | 179.69 |
|  | Standard deviation | 25.90 | 43.55 | 64.13 | 208.22 | 0.28 | 0.85 | 10.67 | 19.72 | 301.51 |
|  | Median | 2.21 | 99.00 | 1.00 | 38.00 | 0.00 | 0.00 | 2.00 | 14.00 | 46.00 |

**Table S74**. MolProbity report on structures in Puzzle 04.

| No. | 3D RNA model | (a) Clash score | (b) Rank-ing | #Bad bonds | #Bad angles | #Chiral handedness swaps | #Tetrahedral geometry outli-ers | #Probably wrong sugar puck-ers | (c) #Bad back-bone confor-mations | Total |
|---|---|---|---|---|---|---|---|---|---|---|
| 1 | PZ04_Das_1 | 8.60 | 79 | 1 | 7 | 0 | 0 | 0 | 3 | 11 |
| 2 | PZ04_Das_2 | 8.60 | 79 | 1 | 7 | 0 | 0 | 0 | 3 | 11 |
| 3 | PZ04_Das_3 | 8.60 | 79 | 1 | 7 | 0 | 0 | 0 | 3 | 11 |
| 4 | PZ04_Das_4 | 8.60 | 79 | 1 | 7 | 0 | 0 | 0 | 3 | 11 |
| 5 | PZ04_Das_5 | 8.60 | 79 | 1 | 7 | 0 | 0 | 0 | 3 | 11 |

| | | | | | | | | | | |
|---|---|---|---|---|---|---|---|---|---|---|
| 6 | PZ04_Adamiak_3 | 18.44 | 36 | 0 | 0 | 0 | 0 | 8 | 45 | 53 |
| 7 | PZ04_Adamiak_2 | 19.91 | 33 | 0 | 0 | 0 | 0 | 8 | 46 | 54 |
| 8 | PZ04_refstructure | 26.15 | 19 | 0 | 31 | 0 | 2 | 10 | 21 | 64 |
| 9 | PZ04_Adamiak_1 | 17.70 | 39 | 0 | 2 | 0 | 0 | 8 | 59 | 69 |
| 10 | PZ04_Adamiak_5 | 69.08 | 1 | 0 | 0 | 0 | 0 | 7 | 68 | 75 |
| 11 | PZ04_Santalucia_1 | 33.19 | 13 | 30 | 23 | 0 | 3 | 5 | 22 | 83 |
| 12 | PZ04_Adamiak_4 | 69.08 | 1 | 0 | 0 | 0 | 0 | 8 | 79 | 87 |
| 13 | PZ04_Bujnicki_3 | 22.13 | 27 | 29 | 49 | 0 | 1 | 8 | 27 | 114 |
| 14 | PZ04_Bujnicki_4 | 25.82 | 20 | 29 | 45 | 0 | 1 | 10 | 29 | 114 |
| 15 | PZ04_Bujnicki_5 | 21.15 | 30 | 29 | 49 | 0 | 1 | 9 | 27 | 115 |
| 16 | PZ04_Bujnicki_2 | 25.82 | 20 | 30 | 46 | 0 | 1 | 10 | 29 | 116 |
| 17 | PZ04_Dokholyan_1 | 12.54 | 60 | 0 | 75 | 1 | 3 | 10 | 27 | 116 |
| 18 | PZ04_Dokholyan_2 | 12.54 | 60 | 0 | 75 | 1 | 3 | 10 | 27 | 116 |
| 19 | PZ04_Bujnicki_1 | 20.90 | 30 | 31 | 52 | 0 | 1 | 8 | 26 | 118 |
| 20 | PZ04_Mikolajczak_1 | 29.53 | 16 | 31 | 59 | 0 | 4 | 14 | 30 | 138 |
| 21 | PZ04_Chen_1 | 1.97 | 99 | 1 | 106 | 0 | 0 | 9 | 23 | 139 |
| 22 | PZ04_Chen_4 | 3.45 | 97 | 1 | 119 | 0 | 1 | 9 | 20 | 150 |
| 23 | PZ04_Chen_6 | 2.46 | 99 | 1 | 122 | 0 | 1 | 9 | 19 | 152 |
| 24 | PZ04_Major_1 | 0.49 | 99 | 0 | 108 | 1 | 0 | 8 | 35 | 152 |
| 25 | PZ04_Chen_10 | 1.97 | 99 | 1 | 124 | 0 | 1 | 9 | 19 | 154 |
| 26 | PZ04_Chen_8 | 3.46 | 97 | 2 | 126 | 0 | 0 | 9 | 21 | 158 |
| 27 | PZ04_Chen_2 | 2.22 | 99 | 1 | 132 | 0 | 1 | 9 | 20 | 163 |
| 28 | PZ04_Chen_7 | 2.96 | 98 | 1 | 130 | 0 | 1 | 10 | 22 | 164 |
| 29 | PZ04_Chen_3 | 2.46 | 99 | 2 | 136 | 0 | 1 | 9 | 20 | 168 |
| 30 | PZ04_Chen_9 | 1.98 | 99 | 4 | 141 | 0 | 1 | 9 | 19 | 174 |
| 31 | PZ04_Chen_5 | 2.71 | 98 | 8 | 150 | 0 | 1 | 9 | 19 | 187 |
| | Average | 15.91 | 60.74 | 7.61 | 62.42 | 0.10 | 0.90 | 7.48 | 26.26 | 104.77 |
| | Standard deviation | 17.19 | 35.91 | 12.31 | 53.46 | 0.30 | 1.08 | 3.61 | 17.87 | 55.12 |
| | Median | 8.60 | 79.00 | 1.00 | 49.00 | 0.00 | 1.00 | 9.00 | 22.00 | 116.00 |

**Table S75**. MolProbity report on structures in Puzzle 05.

| No. | 3D RNA model | (a) Clash score | (b) Ranking | #Bad bonds | #Bad angles | #Chiral handedness swaps | #Tetrahedral geometry outliers | #Probably wrong sugar puckers | (c) #Bad backbone conformations | Total |
|---|---|---|---|---|---|---|---|---|---|---|
| 1 | PZ05_refstructure | 7.95 | 82 | 0 | 6 | 0 | 2 | 1 | 28 | 37 |
| 2 | PZ05_Adamiak_1 | 16.56 | 43 | 0 | 2 | 0 | 0 | 6 | 45 | 53 |
| 3 | PZ05_Bujnicki_1 | 1.49 | 99 | 0 | 45 | 0 | 10 | 0 | 35 | 90 |
| 4 | PZ05_Chen_6 | 0.33 | 99 | 0 | 23 | 5 | 0 | 16 | 46 | 90 |
| 5 | PZ05_Chen_7 | 0.00 | 100 | 0 | 29 | 5 | 0 | 14 | 47 | 95 |
| 6 | PZ05_Das_2 | 9.93 | 72 | 10 | 50 | 0 | 0 | 2 | 33 | 95 |
| 7 | PZ05_Bujnicki_4 | 1.32 | 99 | 0 | 42 | 0 | 13 | 3 | 40 | 98 |
| 8 | PZ05_Bujnicki_5 | 0.99 | 99 | 0 | 52 | 0 | 9 | 2 | 41 | 104 |
| 9 | PZ05_Das_1 | 11.75 | 64 | 13 | 46 | 0 | 0 | 6 | 44 | 109 |
| 10 | PZ05_Bujnicki_3 | 6.13 | 90 | 0 | 53 | 0 | 10 | 2 | 47 | 112 |
| 11 | PZ05_Chen_3 | 11.42 | 65 | 24 | 40 | 4 | 2 | 10 | 43 | 123 |
| 12 | PZ05_Bujnicki_2 | 2.65 | 98 | 0 | 67 | 0 | 19 | 1 | 44 | 131 |
| 13 | PZ05_Chen_1 | 5.63 | 92 | 10 | 53 | 5 | 5 | 14 | 47 | 134 |
| 14 | PZ05_Chen_4 | 9.11 | 76 | 20 | 53 | 5 | 6 | 15 | 49 | 148 |
| 15 | PZ05_Chen_5 | 14.24 | 53 | 28 | 54 | 6 | 10 | 14 | 41 | 153 |
| 16 | PZ05_Chen_2 | 8.94 | 77 | 21 | 58 | 5 | 6 | 18 | 56 | 164 |
| 17 | PZ05_Dokholyan_3 | 10.43 | 69 | 0 | 242 | 0 | 0 | 7 | 57 | 306 |
| 18 | PZ05_Dokholyan_5 | 10.93 | 67 | 0 | 242 | 0 | 0 | 13 | 64 | 319 |
| 19 | PZ05_Xiao_1 | 0.00 | 100 | 23 | 206 | 2 | 2 | 16 | 72 | 321 |
| 20 | PZ05_Dokholyan_2 | 11.26 | 66 | 0 | 251 | 0 | 0 | 13 | 59 | 323 |
| 21 | PZ05_Dokholyan_4 | 11.26 | 66 | 0 | 251 | 0 | 0 | 13 | 59 | 323 |
| 22 | PZ05_Dokholyan_1 | 9.77 | 73 | 0 | 270 | 0 | 0 | 13 | 58 | 341 |
| 23 | PZ05_Dokholyan_6 | 14.24 | 53 | 4 | 284 | 0 | 1 | 14 | 50 | 353 |
| 24 | PZ05_Dokholyan_7 | 10.76 | 68 | 11 | 284 | 0 | 0 | 14 | 55 | 364 |
| 25 | PZ05_Dokholyan_8 | 11.59 | 64 | 7 | 315 | 0 | 0 | 18 | 67 | 407 |
| 26 | PZ05_Xiao_2 | 0.17 | 99 | 57 | 270 | 2 | 3 | 29 | 82 | 443 |
|  | Average | 7.65 | 78.19 | 8.77 | 126.46 | 1.50 | 3.77 | 10.54 | 50.35 | 201.38 |
|  | Standard deviation | 5.14 | 17.46 | 13.47 | 111.25 | 2.25 | 5.14 | 7.04 | 12.29 | 125.48 |
|  | Median | 9.44 | 74.50 | 0.00 | 53.50 | 0.00 | 1.50 | 13.00 | 47.00 | 141.00 |

**Table S76**. MolProbity report on structures in Puzzle 06.

| No. | 3D RNA model | (a) Clash score | (b) Ranking | #Bad bonds | #Bad angles | #Chiral handedness swaps | #Tetrahedral geometry outliers | #Probably wrong sugar puckers: | (c) #Bad backbone conformations | Total |
|---|---|---|---|---|---|---|---|---|---|---|
| 1 | PZ06_refstructure | 9.74 | 73 | 0 | 0 | 0 | 0 | 3 | 33 | 36 |
| 2 | PZ06_Bujnicki_1 | 0.55 | 99 | 0 | 33 | 0 | 9 | 1 | 19 | 62 |
| 3 | PZ06_Bujnicki_3 | 1.10 | 99 | 0 | 32 | 0 | 7 | 1 | 26 | 66 |
| 4 | PZ06_Bujnicki_4 | 1.83 | 99 | 0 | 31 | 0 | 8 | 0 | 30 | 69 |
| 5 | PZ06_Das_5 | 14.67 | 51 | 7 | 24 | 0 | 0 | 2 | 41 | 74 |
| 6 | PZ06_Bujnicki_2 | 1.28 | 99 | 0 | 40 | 0 | 9 | 1 | 25 | 75 |
| 7 | PZ06_Das_8 | 15.22 | 48 | 7 | 24 | 0 | 0 | 3 | 41 | 75 |
| 8 | PZ06_Das_9 | 20.54 | 31 | 11 | 29 | 0 | 0 | 4 | 33 | 77 |
| 9 | PZ06_Das_10 | 23.67 | 23 | 13 | 26 | 0 | 0 | 4 | 38 | 81 |
| 10 | PZ06_Das_2 | 20.55 | 31 | 12 | 31 | 0 | 0 | 3 | 35 | 81 |
| 11 | PZ06_Das_4 | 27.88 | 17 | 12 | 37 | 0 | 0 | 4 | 37 | 90 |
| 12 | PZ06_Das_1 | 28.99 | 16 | 12 | 37 | 0 | 0 | 5 | 37 | 91 |
| 13 | PZ06_Das_6 | 29.72 | 16 | 13 | 40 | 0 | 0 | 5 | 37 | 95 |
| 14 | PZ06_Das_7 | 15.96 | 46 | 10 | 29 | 0 | 0 | 5 | 56 | 100 |
| 15 | PZ06_Das_3 | 18.71 | 36 | 10 | 27 | 0 | 0 | 4 | 60 | 101 |
| 16 | PZ06_Chen_6 | 2.20 | 99 | 9 | 73 | 0 | 1 | 6 | 52 | 141 |
| 17 | PZ06_Chen_1 | 0.37 | 99 | 1 | 63 | 0 | 0 | 18 | 69 | 151 |
| 18 | PZ06_Chen_7 | 9.90 | 72 | 18 | 57 | 0 | 5 | 8 | 70 | 158 |
| 19 | PZ06_Major_4 | 0.92 | 99 | 0 | 90 | 4 | 0 | 13 | 57 | 164 |
| 20 | PZ06_Major_6 | 0.73 | 99 | 1 | 94 | 3 | 0 | 16 | 50 | 164 |
| 21 | PZ06_Chen_4 | 1.83 | 99 | 5 | 97 | 0 | 0 | 8 | 57 | 167 |
| 22 | PZ06_Major_1 | 0.73 | 99 | 0 | 89 | 4 | 0 | 16 | 61 | 170 |
| 23 | PZ06_Chen_2 | 0.18 | 99 | 2 | 70 | 0 | 0 | 12 | 93 | 177 |
| 24 | PZ06_Major_3 | 0.92 | 99 | 0 | 104 | 5 | 0 | 16 | 53 | 178 |
| 25 | PZ06_Major_7 | 1.10 | 99 | 0 | 114 | 4 | 0 | 13 | 54 | 185 |
| 26 | PZ06_Major_2 | 0.55 | 99 | 0 | 102 | 7 | 0 | 17 | 62 | 188 |
| 27 | PZ06_Chen_3 | 0.55 | 99 | 8 | 83 | 0 | 0 | 12 | 96 | 199 |

| | | | | | | | | | | |
|---|---|---|---|---|---|---|---|---|---|---|
| 28 | PZ06_Major_5 | 0.73 | 99 | 0 | 129 | 5 | 0 | 18 | 59 | 211 |
| 29 | PZ06_Chen_5 | 17.43 | 40 | 34 | 150 | 0 | 6 | 15 | 57 | 262 |
| 30 | PZ06_Dokholyan_5 | 10.46 | 69 | 5 | 240 | 0 | 0 | 9 | 47 | 301 |
| 31 | PZ06_Dokholyan_2 | 9.36 | 75 | 2 | 248 | 0 | 0 | 12 | 51 | 313 |
| 32 | PZ06_Dokholyan_1 | 11.01 | 67 | 5 | 246 | 0 | 0 | 10 | 54 | 315 |
| 33 | PZ06_Dokholyan_3 | 9.17 | 76 | 3 | 259 | 0 | 0 | 9 | 46 | 317 |
| 34 | PZ06_Dokholyan_6 | 14.68 | 51 | 0 | 255 | 0 | 0 | 17 | 59 | 331 |
| 35 | PZ06_Dokholyan_4 | 9.54 | 74 | 7 | 278 | 0 | 0 | 9 | 58 | 352 |
| | Average | 9.51 | 71.31 | 5.91 | 93.74 | 0.91 | 1.29 | 8.54 | 50.09 | 160.49 |
| | Standard deviation | 9.42 | 30.31 | 7.14 | 81.82 | 1.93 | 2.87 | 5.73 | 16.93 | 90.67 |
| | Median | 9.36 | 75.00 | 5.00 | 70.00 | 0.00 | 0.00 | 8.00 | 52.00 | 158.00 |

**Table S77**. MolProbity report on structures in Puzzle 07.

| No. | 3D RNA model | (a) Clash score | (b) Ranking | #Bad bonds | #Bad angles | #Chiral handedness swaps | #Tetrahedral geometry outliers | #Probably wrong sugar puckers: | (c) #Bad backbone conformations | Total |
|---|---|---|---|---|---|---|---|---|---|---|
| 1 | PZ07_refstructure | 3.02 | 98 | 0 | 3 | 0 | 0 | 0 | 21 | 24 |
| 2 | PZ07_Adamiak_5 | 9.91 | 72 | 0 | 0 | 0 | 0 | 1 | 46 | 47 |
| 3 | PZ07_Adamiak_4 | 14.61 | 51 | 0 | 1 | 0 | 0 | 3 | 45 | 49 |
| 4 | PZ07_Adamiak_1 | 14.44 | 52 | 0 | 0 | 0 | 0 | 2 | 59 | 61 |
| 5 | PZ07_Adamiak_2 | 14.95 | 50 | 0 | 0 | 0 | 0 | 4 | 57 | 61 |
| 6 | PZ07_Adamiak_3 | 17.64 | 39 | 0 | 0 | 0 | 0 | 5 | 56 | 61 |
| 7 | PZ07_Das_5 | 7.39 | 85 | 1 | 8 | 0 | 0 | 4 | 48 | 61 |
| 8 | PZ07_Das_8 | 7.73 | 83 | 4 | 14 | 0 | 0 | 6 | 41 | 65 |
| 9 | PZ07_Das_7 | 13.29 | 57 | 4 | 13 | 0 | 0 | 6 | 54 | 77 |
| 10 | PZ07_Das_2 | 7.89 | 82 | 6 | 21 | 0 | 1 | 6 | 45 | 79 |
| 11 | PZ07_Das_3 | 8.40 | 80 | 3 | 21 | 0 | 0 | 6 | 49 | 79 |
| 12 | PZ07_Das_1 | 11.25 | 66 | 4 | 21 | 0 | 0 | 8 | 47 | 80 |
| 13 | PZ07_Das_6 | 10.58 | 68 | 4 | 16 | 0 | 1 | 6 | 57 | 84 |
| 14 | PZ07_Bujnicki_4 | 1.01 | 99 | 0 | 45 | 7 | 1 | 9 | 24 | 86 |
| 15 | PZ07_Bujnicki_5 | 1.01 | 99 | 0 | 48 | 7 | 1 | 9 | 24 | 89 |
| 16 | PZ07_Das_4 | 9.57 | 74 | 7 | 23 | 0 | 2 | 8 | 57 | 97 |

| | | | | | | | | | | |
|---|---|---|---|---|---|---|---|---|---|---|
| 17 | PZ07_Bujnicki_7 | 1.01 | 99 | 0 | 61 | 5 | 0 | 9 | 31 | 106 |
| 18 | PZ07_Bujnicki_3 | 0.50 | 99 | 1 | 56 | 9 | 0 | 14 | 30 | 110 |
| 19 | PZ07_Bujnicki_2 | 0.34 | 99 | 0 | 69 | 6 | 4 | 10 | 25 | 114 |
| 20 | PZ07_Bujnicki_1 | 1.01 | 99 | 0 | 71 | 7 | 3 | 9 | 25 | 115 |
| 21 | PZ07_Bujnicki_6 | 0.50 | 99 | 1 | 63 | 10 | 0 | 15 | 34 | 123 |
| 22 | PZ07_Chen_8 | 55.32 | 3 | 89 | 167 | 0 | 0 | 6 | 26 | 288 |
| 23 | PZ07_Chen_9 | 64.15 | 1 | 90 | 167 | 0 | 0 | 6 | 26 | 289 |
| 24 | PZ07_Chen_10 | 47.47 | 4 | 91 | 169 | 0 | 0 | 7 | 24 | 291 |
| 25 | PZ07_Dokholyan_2 | 8.73 | 78 | 0 | 231 | 0 | 0 | 6 | 56 | 293 |
| 26 | PZ07_Chen_5 | 61.90 | 2 | 96 | 177 | 0 | 0 | 4 | 25 | 302 |
| 27 | PZ07_Chen_3 | 53.52 | 3 | 97 | 181 | 0 | 0 | 6 | 23 | 307 |
| 28 | PZ07_Chen_6 | 60.76 | 2 | 96 | 179 | 0 | 0 | 5 | 27 | 307 |
| 29 | PZ07_Chen_2 | 59.79 | 2 | 88 | 178 | 0 | 0 | 10 | 32 | 308 |
| 30 | PZ07_Chen_7 | 71.32 | 1 | 86 | 185 | 0 | 0 | 10 | 31 | 312 |
| 31 | PZ07_Chen_4 | 65.65 | 1 | 96 | 191 | 0 | 0 | 7 | 23 | 317 |
| 32 | PZ07_Dokholyan_1 | 7.05 | 86 | 0 | 253 | 1 | 0 | 9 | 61 | 324 |
| 33 | PZ07_Ding_10 | 11.25 | 66 | 9 | 280 | 0 | 0 | 7 | 46 | 342 |
| 34 | PZ07_Ding_8 | 13.77 | 55 | 5 | 278 | 0 | 0 | 8 | 57 | 348 |
| 35 | PZ07_Ding_4 | 14.28 | 52 | 7 | 274 | 0 | 0 | 12 | 59 | 352 |
| 36 | PZ07_Ding_3 | 15.45 | 48 | 4 | 287 | 0 | 0 | 14 | 54 | 359 |
| 37 | PZ07_Ding_9 | 11.08 | 66 | 10 | 275 | 0 | 0 | 12 | 64 | 361 |
| 38 | PZ07_Ding_1 | 14.61 | 51 | 8 | 286 | 0 | 1 | 17 | 51 | 363 |
| 39 | PZ07_Ding_5 | 11.76 | 64 | 5 | 284 | 0 | 0 | 13 | 63 | 365 |
| 40 | PZ07_Ding_6 | 10.92 | 67 | 8 | 282 | 0 | 0 | 12 | 68 | 370 |
| 41 | PZ07_Ding_7 | 12.26 | 62 | 6 | 291 | 0 | 0 | 12 | 61 | 370 |
| 42 | PZ07_Ding_2 | 10.92 | 67 | 6 | 304 | 0 | 0 | 8 | 60 | 378 |
| 43 | PZ07_Chen_1 | 0.00 | 100 | 281 | 349 | 3 | 0 | 22 | 91 | 746 |
| 44 | PZ07_Major_4 | 0.00 | 100 | 61 | 577 | 4 | 5 | 25 | 91 | 763 |
| 45 | PZ07_Major_7 | 0.00 | 100 | 61 | 595 | 2 | 7 | 26 | 89 | 780 |
| 46 | PZ07_Major_5 | 0.00 | 100 | 60 | 603 | 2 | 6 | 24 | 89 | 784 |
| 47 | PZ07_Major_8 | 0.00 | 100 | 55 | 643 | 0 | 7 | 23 | 116 | 844 |
| 48 | PZ07_Major_6 | 0.00 | 100 | 91 | 641 | 1 | 6 | 33 | 100 | 872 |
| 49 | PZ07_Major_1 | 0.00 | 100 | 62 | 669 | 2 | 8 | 29 | 116 | 886 |

| | | | | | | | | | | |
|---|---|---|---|---|---|---|---|---|---|---|
| 50 | PZ07_Major_9 | 0.00 | 100 | 84 | 683 | 2 | 12 | 33 | 110 | 924 |
| 51 | PZ07_Major_3 | 0.00 | 100 | 90 | 680 | 3 | 12 | 36 | 105 | 926 |
| 52 | PZ07_Major_10 | 0.00 | 100 | 95 | 700 | 2 | 10 | 33 | 111 | 951 |
| 53 | PZ07_Major_2 | 0.00 | 100 | 91 | 715 | 3 | 15 | 29 | 115 | 968 |
| | Average | 15.81 | 66.62 | 37.04 | 232.60 | 1.43 | 1.92 | 12.15 | 55.57 | 340.72 |
| | Standard deviation | 21.08 | 34.59 | 52.27 | 229.65 | 2.58 | 3.69 | 9.31 | 28.35 | 293.59 |
| | Median | 9.91 | 72.00 | 6.00 | 179.00 | 0.00 | 0.00 | 9.00 | 54.00 | 307.00 |

**Table S78**. MolProbity report on structures in Puzzle 08.

| No. | 3D RNA model | (a) Clash score | (b) Ranking | #Bad bonds | #Bad angles | #Chiral handedness swaps | #Tetrahedral geometry outliers | #Probably wrong sugar puckers: | (c) #Bad backbone conformations | Total |
|---|---|---|---|---|---|---|---|---|---|---|
| 1 | PZ08_refstructure | 7.04 | 86 | 1 | 10 | 0 | 0 | 3 | 13 | 27 |
| 2 | PZ08_Adamiak_1 | 10.25 | 70 | 0 | 0 | 0 | 0 | 1 | 19 | 20 |
| 3 | PZ08_Adamiak_2 | 11.21 | 66 | 0 | 0 | 0 | 0 | 1 | 24 | 25 |
| 4 | PZ08_Bujnicki_1 | 8.47 | 79 | 14 | 146 | 2 | 3 | 8 | 23 | 196 |
| 5 | PZ08_Bujnicki_10 | 9.38 | 74 | 14 | 133 | 8 | 1 | 16 | 25 | 197 |
| 6 | PZ08_Bujnicki_2 | 12.81 | 59 | 16 | 225 | 0 | 6 | 8 | 26 | 281 |
| 7 | PZ08_Bujnicki_3 | 29.32 | 16 | 20 | 180 | 0 | 5 | 6 | 22 | 233 |
| 8 | PZ08_Bujnicki_4 | 26.06 | 20 | 13 | 183 | 1 | 2 | 6 | 20 | 225 |
| 9 | PZ08_Bujnicki_5 | 18.89 | 35 | 17 | 180 | 1 | 3 | 7 | 25 | 233 |
| 10 | PZ08_Bujnicki_6 | 11.85 | 63 | 12 | 150 | 3 | 2 | 9 | 24 | 200 |
| 11 | PZ08_Bujnicki_7 | 0.96 | 99 | 0 | 36 | 8 | 0 | 9 | 20 | 73 |
| 12 | PZ08_Bujnicki_8 | 5.76 | 91 | 7 | 92 | 7 | 2 | 10 | 19 | 137 |
| 13 | PZ08_Bujnicki_9 | 4.80 | 94 | 8 | 72 | 2 | 0 | 10 | 22 | 114 |
| 14 | PZ08_Chen_1 | 38.45 | 9 | 25 | 87 | 1 | 2 | 0 | 41 | 156 |
| 15 | PZ08_Chen_10 | 24.90 | 22 | 11 | 50 | 2 | 0 | 0 | 40 | 103 |
| 16 | PZ08_Chen_2 | 24.90 | 22 | 3 | 37 | 0 | 0 | 0 | 34 | 74 |
| 17 | PZ08_Chen_3 | 30.55 | 15 | 8 | 59 | 0 | 1 | 0 | 37 | 105 |
| 18 | PZ08_Chen_4 | 27.54 | 18 | 5 | 55 | 0 | 0 | 0 | 37 | 97 |
| 19 | PZ08_Chen_5 | 30.55 | 15 | 5 | 53 | 0 | 0 | 0 | 29 | 87 |
| 20 | PZ08_Chen_6 | 37.34 | 9 | 133 | 205 | 0 | 1 | 0 | 52 | 391 |

| | | | | | | | | | | |
|---|---|---|---|---|---|---|---|---|---|---|
| 21 | PZ08_Chen_7 | 63.37 | 1 | 45 | 117 | 0 | 4 | 0 | 39 | 205 |
| 22 | PZ08_Chen_8 | 34.70 | 11 | 164 | 181 | 1 | 0 | 0 | 46 | 392 |
| 23 | PZ08_Chen_9 | 29.80 | 16 | 167 | 236 | 1 | 0 | 0 | 50 | 454 |
| 24 | PZ08_Das_1 | 14.41 | 52 | 1 | 4 | 0 | 0 | 1 | 13 | 19 |
| 25 | PZ08_Das_2 | 12.81 | 59 | 1 | 4 | 0 | 0 | 1 | 15 | 21 |
| 26 | PZ08_Das_3 | 13.77 | 55 | 2 | 5 | 0 | 0 | 0 | 16 | 23 |
| 27 | PZ08_Das_4 | 7.36 | 85 | 1 | 5 | 0 | 0 | 1 | 23 | 30 |
| 28 | PZ08_Das_5 | 7.68 | 83 | 1 | 5 | 0 | 0 | 3 | 28 | 37 |
| 29 | PZ08_Das_6 | 5.44 | 92 | 0 | 5 | 0 | 0 | 0 | 27 | 32 |
| 30 | PZ08_Ding_1 | 14.09 | 53 | 4 | 148 | 0 | 0 | 14 | 33 | 199 |
| 31 | PZ08_Ding_10 | 12.49 | 61 | 2 | 152 | 0 | 0 | 12 | 35 | 201 |
| 32 | PZ08_Ding_2 | 17.61 | 40 | 2 | 160 | 0 | 0 | 6 | 35 | 203 |
| 33 | PZ08_Ding_3 | 14.41 | 52 | 4 | 164 | 0 | 0 | 11 | 31 | 210 |
| 34 | PZ08_Ding_4 | 10.89 | 67 | 1 | 154 | 0 | 0 | 5 | 29 | 189 |
| 35 | PZ08_Ding_5 | 12.17 | 62 | 2 | 141 | 0 | 0 | 8 | 36 | 187 |
| 36 | PZ08_Ding_6 | 7.36 | 85 | 3 | 147 | 0 | 0 | 9 | 29 | 188 |
| 37 | PZ08_Ding_7 | 11.21 | 66 | 4 | 147 | 0 | 0 | 7 | 30 | 188 |
| 38 | PZ08_Ding_8 | 13.77 | 55 | 3 | 137 | 0 | 0 | 11 | 32 | 183 |
| 39 | PZ08_Ding_9 | 13.45 | 56 | 4 | 162 | 0 | 0 | 6 | 33 | 205 |
| 40 | PZ08_Dokholyan_1 | 7.36 | 85 | 0 | 130 | 0 | 0 | 5 | 22 | 157 |
| 41 | PZ08_Dokholyan_2 | 11.53 | 65 | 0 | 136 | 0 | 0 | 4 | 30 | 170 |
| 42 | PZ08_Dokholyan_3 | 6.40 | 89 | 0 | 119 | 0 | 0 | 6 | 31 | 156 |
| 43 | PZ08_Dokholyan_4 | 9.61 | 73 | 0 | 127 | 0 | 0 | 3 | 31 | 161 |
| | Average | 16.81 | 54.07 | 16.81 | 105.56 | 0.86 | 0.74 | 4.81 | 28.98 | 157.77 |
| | Standard deviation | 12.06 | 28.70 | 39.36 | 70.10 | 2.02 | 1.47 | 4.51 | 9.14 | 102.08 |
| | Median | 12.81 | 59.00 | 4.00 | 130.00 | 0.00 | 0.00 | 5.00 | 29.00 | 170.00 |

**Table S79**. MolProbity report on structures in Puzzle 09.

| No. | 3D RNA model | (a) Clash score | (b) Ranking | #Bad bonds | #Bad angles | #Chiral handedness swaps | #Tetrahedral geometry outliers | #Probably wrong sugar puckers: | (c) #Bad backbone conformations | Total |
|---|---|---|---|---|---|---|---|---|---|---|
| 1 | PZ09_refstructure | 4.39 | 95 | 0 | 2 | 0 | 0 | 0 | 8 | 10 |
| 2 | PZ09_Das_8 | 2.20 | 99 | 0 | 3 | 0 | 0 | 0 | 11 | 14 |
| 3 | PZ09_Das_2 | 2.20 | 99 | 2 | 6 | 1 | 0 | 0 | 6 | 15 |
| 4 | PZ09_Chen_2 | 26.42 | 19 | 6 | 6 | 0 | 0 | 0 | 5 | 17 |
| 5 | PZ09_Das_6 | 9.22 | 75 | 1 | 1 | 0 | 0 | 1 | 14 | 17 |
| 6 | PZ09_Das_9 | 3.07 | 98 | 1 | 5 | 0 | 0 | 1 | 12 | 19 |
| 7 | PZ09_Das_4 | 3.51 | 97 | 2 | 5 | 1 | 0 | 0 | 12 | 20 |
| 8 | PZ09_Das_7 | 9.22 | 75 | 1 | 1 | 0 | 0 | 4 | 14 | 20 |
| 9 | PZ09_Das_5 | 4.83 | 94 | 2 | 6 | 1 | 0 | 1 | 11 | 21 |
| 10 | PZ09_Das_1 | 4.83 | 94 | 2 | 7 | 1 | 0 | 0 | 12 | 22 |
| 11 | PZ09_Das_3 | 15.81 | 46 | 2 | 8 | 1 | 0 | 2 | 10 | 23 |
| 12 | PZ09_Chen_4 | 23.76 | 23 | 6 | 8 | 0 | 1 | 2 | 8 | 25 |
| 13 | PZ09_Chen_5 | 33.01 | 13 | 7 | 10 | 0 | 0 | 8 | 17 | 42 |
| 14 | PZ09_Chen_3 | 29.49 | 16 | 8 | 27 | 0 | 1 | 2 | 8 | 46 |
| 15 | PZ09_Chen_1 | 34.73 | 11 | 10 | 38 | 0 | 0 | 0 | 4 | 52 |
| 16 | PZ09_Chen_7 | 18.08 | 37 | 11 | 36 | 0 | 0 | 0 | 5 | 52 |
| 17 | PZ09_Chen_8 | 65.82 | 1 | 15 | 52 | 0 | 0 | 0 | 4 | 71 |
| 18 | PZ09_Bujnicki_3 | 1.32 | 99 | 0 | 26 | 12 | 0 | 18 | 31 | 87 |
| 19 | PZ09_Bujnicki_4 | 1.32 | 99 | 0 | 26 | 12 | 0 | 18 | 31 | 87 |
| 20 | PZ09_Bujnicki_5 | 1.32 | 99 | 0 | 26 | 12 | 0 | 18 | 31 | 87 |
| 21 | PZ09_Chen_6 | 64.62 | 1 | 20 | 66 | 0 | 0 | 1 | 9 | 96 |
| 22 | PZ09_Dokholyan_4 | 4.83 | 94 | 0 | 88 | 0 | 0 | 3 | 20 | 111 |
| 23 | PZ09_Ding_5 | 14.05 | 53 | 2 | 99 | 0 | 0 | 1 | 17 | 119 |
| 24 | PZ09_Dokholyan_1 | 7.91 | 82 | 0 | 96 | 0 | 0 | 7 | 28 | 131 |
| 25 | PZ09_Dokholyan_3 | 6.59 | 89 | 0 | 103 | 0 | 0 | 8 | 24 | 135 |
| 26 | PZ09_Ding_10 | 8.78 | 78 | 2 | 109 | 0 | 0 | 4 | 22 | 137 |
| 27 | PZ09_Ding_9 | 10.54 | 68 | 6 | 109 | 0 | 1 | 2 | 19 | 137 |
| 28 | PZ09_Ding_6 | 10.54 | 68 | 1 | 116 | 0 | 0 | 4 | 21 | 142 |

| | | | | | | | | | | |
|---|---|---|---|---|---|---|---|---|---|---|
| 29 | PZ09_Ding_7 | 9.66 | 73 | 2 | 111 | 0 | 0 | 6 | 24 | 143 |
| 30 | PZ09_Dokholyan_2 | 7.03 | 86 | 0 | 120 | 0 | 0 | 4 | 20 | 144 |
| 31 | PZ09_Ding_8 | 14.05 | 53 | 5 | 123 | 0 | 0 | 3 | 16 | 147 |
| 32 | PZ09_Ding_4 | 12.30 | 61 | 5 | 114 | 0 | 0 | 7 | 22 | 148 |
| 33 | PZ09_Ding_1 | 14.93 | 50 | 2 | 120 | 0 | 0 | 6 | 22 | 150 |
| 34 | PZ09_Ding_3 | 10.54 | 68 | 6 | 123 | 0 | 0 | 4 | 17 | 150 |
| 35 | PZ09_Ding_2 | 14.05 | 53 | 3 | 126 | 0 | 0 | 8 | 26 | 163 |
| | Average | 14.43 | 64.74 | 3.71 | 54.91 | 1.17 | 0.09 | 4.09 | 16.03 | 80.00 |
| | Standard deviation | 15.48 | 32.08 | 4.60 | 49.20 | 3.38 | 0.28 | 5.05 | 8.11 | 54.89 |
| | Median | 9.66 | 73.00 | 2.00 | 36.00 | 0.00 | 0.00 | 2.00 | 16.00 | 87.00 |

**Table S80**. MolProbity report on structures in Puzzle 10.

| No. | 3D RNA model | (a) Clash score | (b) Ranking | #Bad bonds | #Bad angles | #Chiral handedness swaps | #Tetrahedral geometry outliers | #Probably wrong sugar puckers: | (c) #Bad backbone conformations | Total |
|---|---|---|---|---|---|---|---|---|---|---|
| 1 | PZ10_refstructure | 4.17 | 96 | 0 | 0 | 0 | 0 | 2 | 29 | 31 |
| 2 | PZ10_Das_5 | 13.24 | 57 | 5 | 27 | 0 | 0 | 6 | 40 | 78 |
| 3 | PZ10_Das_2 | 13.61 | 55 | 7 | 32 | 0 | 0 | 6 | 39 | 84 |
| 4 | PZ10_Das_3 | 13.79 | 55 | 8 | 31 | 0 | 0 | 6 | 41 | 86 |
| 5 | PZ10_Das_4 | 12.34 | 61 | 8 | 30 | 0 | 0 | 8 | 40 | 86 |
| 6 | PZ10_Das_1 | 14.15 | 53 | 6 | 34 | 0 | 1 | 9 | 46 | 96 |
| 7 | PZ10_Bujnicki_4 | 0.29 | 99 | 0 | 94 | 23 | 1 | 24 | 54 | 196 |
| 8 | PZ10_Bujnicki_1 | 1.17 | 99 | 0 | 98 | 26 | 0 | 27 | 56 | 207 |
| 9 | PZ10_Bujnicki_7 | 1.58 | 99 | 0 | 116 | 22 | 0 | 21 | 51 | 210 |
| 10 | PZ10_Bujnicki_2 | 1.75 | 99 | 0 | 99 | 24 | 1 | 28 | 59 | 211 |
| 11 | PZ10_Bujnicki_5 | 1.46 | 99 | 2 | 109 | 23 | 1 | 26 | 51 | 212 |
| 12 | PZ10_Bujnicki_6 | 1.46 | 99 | 1 | 108 | 24 | 1 | 25 | 57 | 216 |
| 13 | PZ10_Bujnicki_3 | 0.29 | 99 | 2 | 88 | 32 | 0 | 33 | 64 | 219 |
| 14 | PZ10_Bujnicki_8 | 2.34 | 99 | 0 | 128 | 18 | 2 | 21 | 53 | 222 |
| 15 | PZ10_Bujnicki_9 | 0.32 | 99 | 0 | 117 | 24 | 1 | 25 | 62 | 229 |
| 16 | PZ10_Bujnicki_10 | 0.00 | 100 | 1 | 123 | 30 | 1 | 30 | 68 | 253 |
| 17 | PZ10_Chen_1 | 9.08 | 77 | 36 | 146 | 2 | 5 | 21 | 79 | 289 |

| | | | | | | | | | | |
|---|---|---|---|---|---|---|---|---|---|---|
| 18 | PZ10_Dokholyan_2 | 11.61 | 64 | 8 | 249 | 0 | 0 | 16 | 61 | 334 |
| 19 | PZ10_Dokholyan_3 | 9.98 | 71 | 12 | 265 | 0 | 0 | 14 | 56 | 347 |
| 20 | PZ10_Dokholyan_5 | 11.79 | 64 | 3 | 272 | 0 | 0 | 11 | 62 | 348 |
| 21 | PZ10_Dokholyan_9 | 13.43 | 56 | 7 | 270 | 0 | 0 | 14 | 69 | 360 |
| 22 | PZ10_Dokholyan_7 | 8.71 | 78 | 10 | 279 | 0 | 0 | 19 | 53 | 361 |
| 23 | PZ10_Dokholyan_1 | 14.70 | 51 | 9 | 293 | 0 | 0 | 12 | 48 | 362 |
| 24 | PZ10_Dokholyan_8 | 12.70 | 60 | 7 | 303 | 0 | 0 | 8 | 46 | 364 |
| 25 | PZ10_Dokholyan_4 | 12.34 | 61 | 8 | 292 | 0 | 1 | 13 | 54 | 368 |
| 26 | PZ10_Dokholyan_6 | 9.25 | 75 | 6 | 301 | 1 | 0 | 16 | 63 | 387 |
| 27 | PZ10_Dokholyan_10 | 15.06 | 49 | 5 | 300 | 1 | 0 | 18 | 65 | 389 |
| | Average | 7.80 | 76.81 | 5.59 | 155.70 | 9.26 | 0.56 | 17.00 | 54.30 | 242.41 |
| | Standard deviation | 5.71 | 19.87 | 7.13 | 105.64 | 12.22 | 1.05 | 8.48 | 11.02 | 111.51 |
| | Median | 9.25 | 75.00 | 5.00 | 117.00 | 0.00 | 0.00 | 16.00 | 54.00 | 222.00 |

**Table S81**. MolProbity report on structures in Puzzle 11.

| **No.** | **3D RNA model** | **(a) Clash score** | **(b) Rank-ing** | **#Bad bonds** | **#Bad angles** | **#Chiral handedness swaps** | **#Tetrahedral geometry outli-ers** | **#Probably wrong sugar puck-ers:** | **(c) #Bad back-bone confor-mations** | **Total** |
|---|---|---|---|---|---|---|---|---|---|---|
| 1 | PZ11_Adamiak_3 | 6.59 | 89 | 0 | 0 | 0 | 0 | 0 | 9 | 9 |
| 2 | PZ11_Adamiak_9 | 10.97 | 67 | 0 | 0 | 0 | 0 | 1 | 9 | 10 |
| 3 | PZ11_Adamiak_5 | 6.58 | 89 | 0 | 0 | 0 | 0 | 1 | 11 | 12 |
| 4 | PZ11_Adamiak_4 | 10.42 | 69 | 0 | 0 | 0 | 0 | 1 | 12 | 13 |
| 5 | PZ11_Chen_10 | 8.23 | 81 | 0 | 1 | 0 | 0 | 0 | 12 | 13 |
| 6 | PZ11_Chen_6 | 17.55 | 40 | 1 | 5 | 0 | 0 | 1 | 6 | 13 |
| 7 | PZ11_Chen_7 | 12.61 | 60 | 1 | 4 | 0 | 0 | 2 | 7 | 14 |
| 8 | PZ11_Chen_8 | 9.87 | 72 | 0 | 1 | 0 | 0 | 1 | 12 | 14 |
| 9 | PZ11_Adamiak_1 | 8.23 | 81 | 0 | 0 | 0 | 0 | 0 | 15 | 15 |
| 10 | PZ11_Adamiak_6 | 10.97 | 67 | 0 | 0 | 0 | 0 | 3 | 12 | 15 |
| 11 | PZ11_Adamiak_2 | 9.87 | 72 | 0 | 0 | 0 | 0 | 3 | 13 | 16 |
| 12 | PZ11_Das_10 | 10.42 | 69 | 0 | 0 | 0 | 0 | 2 | 14 | 16 |
| 13 | PZ11_Adamiak_10 | 9.33 | 75 | 0 | 0 | 0 | 0 | 0 | 18 | 18 |
| 14 | PZ11_Adamiak_8 | 9.33 | 75 | 0 | 0 | 0 | 0 | 2 | 16 | 18 |

| | | | | | | | | | | |
|---|---|---|---|---|---|---|---|---|---|---|
| 15 | PZ11_Adamiak_7 | 7.13 | 86 | 0 | 0 | 0 | 0 | 1 | 18 | 19 |
| 16 | PZ11_refstructure | 0.00 | 100 | 1 | 1 | 0 | 0 | 5 | 12 | 19 |
| 17 | PZ11_Chen_9 | 14.80 | 50 | 1 | 5 | 0 | 0 | 2 | 12 | 20 |
| 18 | PZ11_Das_2 | 12.07 | 62 | 3 | 7 | 0 | 0 | 2 | 8 | 20 |
| 19 | PZ11_Das_3 | 8.78 | 78 | 2 | 5 | 0 | 0 | 2 | 12 | 21 |
| 20 | PZ11_Das_5 | 10.97 | 67 | 2 | 3 | 0 | 0 | 3 | 13 | 21 |
| 21 | PZ11_Das_7 | 9.33 | 75 | 4 | 6 | 0 | 0 | 2 | 9 | 21 |
| 22 | PZ11_Das_8 | 7.68 | 83 | 2 | 5 | 0 | 0 | 2 | 12 | 21 |
| 23 | PZ11_Das_1 | 8.23 | 81 | 3 | 5 | 0 | 0 | 3 | 11 | 22 |
| 24 | PZ11_Das_9 | 12.07 | 62 | 0 | 2 | 0 | 0 | 3 | 17 | 22 |
| 25 | PZ11_Bujnicki_5 | 0.00 | 100 | 0 | 18 | 2 | 0 | 1 | 3 | 24 |
| 26 | PZ11_Bujnicki_8 | 0.00 | 100 | 0 | 13 | 3 | 0 | 1 | 7 | 24 |
| 27 | PZ11_Bujnicki_1 | 0.55 | 99 | 0 | 13 | 2 | 0 | 4 | 8 | 27 |
| 28 | PZ11_Bujnicki_7 | 1.10 | 99 | 0 | 21 | 2 | 0 | 1 | 5 | 29 |
| 29 | PZ11_Das_6 | 8.78 | 78 | 2 | 7 | 0 | 0 | 3 | 17 | 29 |
| 30 | PZ11_Das_4 | 10.42 | 69 | 2 | 7 | 0 | 1 | 5 | 17 | 32 |
| 31 | PZ11_Bujnicki_2 | 0.00 | 100 | 0 | 16 | 3 | 0 | 4 | 10 | 33 |
| 32 | PZ11_Bujnicki_6 | 0.55 | 99 | 0 | 23 | 3 | 0 | 1 | 7 | 34 |
| 33 | PZ11_Bujnicki_9 | 1.10 | 99 | 0 | 20 | 4 | 0 | 2 | 10 | 36 |
| 34 | PZ11_Bujnicki_3 | 0.00 | 100 | 0 | 17 | 5 | 0 | 7 | 13 | 42 |
| 35 | PZ11_Chen_4 | 0.55 | 99 | 0 | 20 | 0 | 0 | 5 | 20 | 45 |
| 36 | PZ11_Chen_1 | 1.10 | 99 | 1 | 24 | 0 | 0 | 2 | 19 | 46 |
| 37 | PZ11_Chen_3 | 1.10 | 99 | 2 | 21 | 0 | 0 | 3 | 21 | 47 |
| 38 | PZ11_Bujnicki_10 | 0.00 | 100 | 0 | 20 | 7 | 0 | 7 | 14 | 48 |
| 39 | PZ11_Xiao_3 | 1.10 | 99 | 1 | 40 | 0 | 0 | 1 | 10 | 52 |
| 40 | PZ11_Chen_2 | 2.74 | 98 | 1 | 27 | 0 | 0 | 5 | 20 | 53 |
| 41 | PZ11_Chen_5 | 0.55 | 99 | 3 | 30 | 0 | 0 | 3 | 18 | 54 |
| 42 | PZ11_Xiao_2 | 1.10 | 99 | 1 | 40 | 0 | 0 | 2 | 14 | 57 |
| 43 | PZ11_Xiao_1 | 4.39 | 95 | 1 | 47 | 0 | 0 | 5 | 19 | 72 |
| 44 | PZ11_Ding_3 | 13.17 | 57 | 3 | 78 | 0 | 0 | 3 | 15 | 99 |
| 45 | PZ11_Ding_1 | 11.52 | 65 | 2 | 82 | 0 | 0 | 6 | 18 | 108 |
| 46 | PZ11_Ding_10 | 13.71 | 55 | 3 | 85 | 0 | 0 | 5 | 17 | 110 |
| 47 | PZ11_Ding_9 | 9.87 | 72 | 0 | 93 | 0 | 0 | 3 | 17 | 113 |

| | | | | | | | | | | |
|---|---|---|---|---|---|---|---|---|---|---|
| 48 | PZ11_Ding_6 | 10.97 | 67 | 0 | 94 | 0 | 0 | 5 | 17 | 116 |
| 49 | PZ11_Ding_8 | 9.87 | 72 | 1 | 88 | 0 | 0 | 5 | 23 | 117 |
| 50 | PZ11_Ding_7 | 14.26 | 53 | 2 | 100 | 0 | 0 | 6 | 15 | 123 |
| 51 | PZ11_Ding_2 | 14.26 | 53 | 2 | 94 | 0 | 0 | 6 | 24 | 126 |
| 52 | PZ11_Ding_4 | 9.87 | 72 | 2 | 98 | 0 | 0 | 7 | 25 | 132 |
| 53 | PZ11_Ding_5 | 12.62 | 60 | 1 | 100 | 0 | 0 | 7 | 26 | 134 |
| 54 | PZ11_Bujnicki_4 | 2.78 | 98 | 57 | 205 | 5 | 0 | 13 | 27 | 307 |
| | Average | 7.22 | 79.70 | 1.98 | 29.46 | 0.67 | 0.02 | 3.15 | 14.19 | 49.46 |
| | Standard deviation | 5.09 | 17.00 | 7.71 | 41.35 | 1.57 | 0.14 | 2.44 | 5.44 | 52.21 |
| | Median | 8.78 | 78.00 | 1.00 | 13.00 | 0.00 | 0.00 | 3.00 | 13.50 | 28.00 |

**Table S82**. MolProbity report on structures in Puzzle 12.

| No. | 3D RNA model | (a) Clash score | (b) Ranking | #Bad bonds | #Bad angles | #Chiral handedness swaps | #Tetrahedral geometry outliers | #Probably wrong sugar puckers: | (c) #Bad backbone conformations | Total |
|---|---|---|---|---|---|---|---|---|---|---|
| 1 | PZ12_Adamiak_3 | 9.15 | 76 | 0 | 0 | 0 | 0 | 0 | 18 | 18 |
| 2 | PZ12_Bujnicki_7 | 18.55 | 36 | 0 | 3 | 0 | 1 | 0 | 18 | 22 |
| 3 | PZ12_Adamiak_2 | 7.91 | 82 | 0 | 0 | 0 | 0 | 0 | 24 | 24 |
| 4 | PZ12_refstructure | 11.35 | 65 | 0 | 0 | 0 | 0 | 2 | 23 | 25 |
| 5 | PZ12_Bujnicki_10 | 27.94 | 17 | 0 | 0 | 0 | 0 | 2 | 23 | 25 |
| 6 | PZ12_Bujnicki_8 | 33.14 | 13 | 0 | 2 | 0 | 1 | 2 | 22 | 27 |
| 7 | PZ12_Bujnicki_4 | 22.01 | 27 | 0 | 3 | 0 | 1 | 0 | 24 | 28 |
| 8 | PZ12_Bujnicki_6 | 21.51 | 29 | 0 | 0 | 0 | 0 | 1 | 27 | 28 |
| 9 | PZ12_Bujnicki_9 | 25.72 | 20 | 0 | 6 | 0 | 0 | 2 | 20 | 28 |
| 10 | PZ12_Bujnicki_1 | 25.22 | 21 | 0 | 0 | 0 | 0 | 2 | 32 | 34 |
| 11 | PZ12_Adamiak_1 | 10.14 | 71 | 0 | 0 | 0 | 0 | 3 | 32 | 35 |
| 12 | PZ12_Bujnicki_3 | 27.94 | 17 | 0 | 2 | 0 | 1 | 3 | 29 | 35 |
| 13 | PZ12_Bujnicki_5 | 28.19 | 17 | 0 | 6 | 0 | 0 | 3 | 27 | 36 |
| 14 | PZ12_Das_8 | 2.97 | 98 | 0 | 2 | 0 | 0 | 1 | 37 | 40 |
| 15 | PZ12_Das_10 | 3.22 | 97 | 1 | 8 | 0 | 0 | 2 | 31 | 42 |
| 16 | PZ12_Das_9 | 3.46 | 97 | 0 | 10 | 0 | 0 | 1 | 32 | 43 |
| 17 | PZ12_Das_5 | 8.65 | 79 | 0 | 2 | 0 | 0 | 3 | 45 | 50 |

| | | | | | | | | | | |
|---|---|---|---|---|---|---|---|---|---|---|
| 18 | PZ12_Das_1 | 8.16 | 81 | 1 | 7 | 0 | 1 | 3 | 43 | 55 |
| 19 | PZ12_Das_2 | 10.14 | 71 | 1 | 11 | 0 | 0 | 2 | 43 | 57 |
| 20 | PZ12_Das_3 | 11.62 | 64 | 0 | 7 | 0 | 1 | 1 | 52 | 61 |
| 21 | PZ12_Das_6 | 8.65 | 79 | 2 | 11 | 0 | 0 | 3 | 47 | 63 |
| 22 | PZ12_Das_4 | 7.91 | 82 | 4 | 8 | 0 | 0 | 5 | 51 | 68 |
| 23 | PZ12_Das_7 | 12.61 | 60 | 2 | 15 | 0 | 0 | 4 | 54 | 75 |
| 24 | PZ12_Bujnicki_2 | 11.13 | 66 | 0 | 4 | 0 | 0 | 5 | 75 | 84 |
| 25 | PZ12_Chen_1 | 0.00 | 100 | 3 | 44 | 0 | 1 | 8 | 45 | 101 |
| 26 | PZ12_Chen_3 | 0.00 | 100 | 12 | 61 | 0 | 1 | 10 | 26 | 110 |
| 27 | PZ12_Chen_2 | 0.00 | 100 | 2 | 47 | 2 | 1 | 10 | 52 | 114 |
| 28 | PZ12_Chen_4 | 0.00 | 100 | 17 | 64 | 0 | 1 | 11 | 36 | 129 |
| 29 | PZ12_Chen_7 | 0.00 | 100 | 12 | 72 | 1 | 3 | 12 | 31 | 131 |
| 30 | PZ12_Weeks_1 | 6.68 | 88 | 0 | 105 | 0 | 0 | 4 | 23 | 132 |
| 31 | PZ12_Chen_6 | 0.00 | 100 | 23 | 72 | 0 | 1 | 12 | 32 | 140 |
| 32 | PZ12_Weeks_2 | 9.64 | 73 | 0 | 107 | 0 | 0 | 4 | 31 | 142 |
| 33 | PZ12_Weeks_3 | 8.16 | 81 | 0 | 113 | 0 | 0 | 6 | 36 | 155 |
| 34 | PZ12_Chen_5 | 0.00 | 100 | 20 | 91 | 0 | 2 | 12 | 31 | 156 |
| 35 | PZ12_Chen_9 | 0.00 | 100 | 33 | 90 | 2 | 2 | 13 | 40 | 180 |
| 36 | PZ12_Chen_8 | 0.00 | 100 | 39 | 126 | 0 | 4 | 15 | 33 | 217 |
| 37 | PZ12_Chen_10 | 0.00 | 100 | 48 | 117 | 3 | 6 | 12 | 35 | 221 |
| 38 | PZ12_Ding_3 | 16.33 | 44 | 5 | 181 | 0 | 0 | 7 | 48 | 241 |
| 39 | PZ12_Ding_6 | 15.33 | 48 | 3 | 190 | 0 | 0 | 11 | 40 | 244 |
| 40 | PZ12_Ding_10 | 12.12 | 62 | 6 | 202 | 0 | 1 | 9 | 32 | 250 |
| 41 | PZ12_Ding_5 | 9.39 | 74 | 6 | 196 | 0 | 0 | 9 | 40 | 251 |
| 42 | PZ12_Ding_7 | 10.14 | 71 | 5 | 200 | 1 | 0 | 7 | 40 | 253 |
| 43 | PZ12_Ding_1 | 14.84 | 50 | 6 | 205 | 0 | 0 | 8 | 38 | 257 |
| 44 | PZ12_Ding_11 | 11.38 | 65 | 5 | 202 | 0 | 0 | 11 | 43 | 261 |
| 45 | PZ12_Ding_4 | 13.11 | 57 | 4 | 207 | 0 | 0 | 14 | 40 | 265 |
| 46 | PZ12_Ding_12 | 12.61 | 60 | 8 | 216 | 0 | 0 | 10 | 42 | 276 |
| 47 | PZ12_Ding_2 | 11.87 | 63 | 6 | 204 | 0 | 0 | 12 | 56 | 278 |
| 48 | PZ12_Ding_9 | 12.86 | 59 | 5 | 214 | 0 | 0 | 15 | 45 | 279 |
| 49 | PZ12_Ding_8 | 13.85 | 54 | 9 | 219 | 0 | 0 | 10 | 42 | 280 |
| 50 | PZ12_Xiao_3 | 20.05 | 32 | 29 | 224 | 6 | 5 | 29 | 63 | 356 |

| | | | | | | | | | | |
|---|---|---|---|---|---|---|---|---|---|---|
| 51 | PZ12_Xiao_1 | 45.04 | 5 | 72 | 312 | 3 | 8 | 28 | 66 | 489 |
| 52 | PZ12_Xiao_2 | 50.45 | 4 | 72 | 301 | 6 | 6 | 38 | 70 | 493 |
| | Average | 12.52 | 64.52 | 8.87 | 86.33 | 0.46 | 0.92 | 7.63 | 38.17 | 142.38 |
| | Standard deviation | 11.11 | 29.56 | 16.57 | 94.02 | 1.32 | 1.78 | 7.60 | 13.08 | 119.52 |
| | Median | 10.64 | 68.50 | 2.00 | 54.00 | 0.00 | 0.00 | 5.50 | 36.50 | 112.00 |

**Table S83**. MolProbity report on structures in Puzzle 13.

| No. | 3D RNA model | (a) Clash score | (b) Rank-ing | #Bad bonds | #Bad angles | #Chiral handedness swaps | #Tetrahedral geometry out-liers | #Probably wrong sugar puck-ers: | (c) #Bad back-bone confor-mations | Total |
|---|---|---|---|---|---|---|---|---|---|---|
| 1 | PZ13_refstructure | 0.00 | 100 | 0 | 0 | 0 | 0 | 0 | 4 | 4 |
| 2 | PZ13_Adamiak_1 | 6.07 | 90 | 0 | 0 | 0 | 0 | 2 | 15 | 17 |
| 3 | PZ13_Das_2 | 4.34 | 96 | 0 | 2 | 0 | 0 | 1 | 16 | 19 |
| 4 | PZ13_Das_4 | 8.68 | 78 | 0 | 5 | 0 | 0 | 2 | 18 | 25 |
| 5 | PZ13_Das_9 | 11.71 | 64 | 0 | 0 | 0 | 0 | 3 | 22 | 25 |
| 6 | PZ13_Das_1 | 5.21 | 93 | 0 | 5 | 0 | 0 | 2 | 19 | 26 |
| 7 | PZ13_Das_10 | 5.64 | 92 | 0 | 3 | 0 | 0 | 2 | 21 | 26 |
| 8 | PZ13_Das_6 | 8.68 | 78 | 0 | 1 | 0 | 0 | 1 | 25 | 27 |
| 9 | PZ13_Das_8 | 4.77 | 94 | 1 | 2 | 0 | 0 | 3 | 23 | 29 |
| 10 | PZ13_Das_7 | 10.85 | 67 | 0 | 3 | 0 | 1 | 4 | 25 | 33 |
| 11 | PZ13_Das_5 | 15.18 | 49 | 0 | 7 | 0 | 0 | 6 | 28 | 41 |
| 12 | PZ13_Das_3 | 6.07 | 90 | 2 | 7 | 6 | 1 | 4 | 25 | 45 |
| 13 | PZ13_Chen_6 | 0.00 | 100 | 6 | 27 | 1 | 0 | 4 | 26 | 64 |
| 14 | PZ13_Bujnicki_8 | 1.74 | 99 | 0 | 49 | 1 | 0 | 5 | 14 | 69 |
| 15 | PZ13_Chen_1 | 0.00 | 100 | 1 | 28 | 2 | 0 | 12 | 38 | 81 |
| 16 | PZ13_Chen_3 | 1.74 | 99 | 4 | 34 | 2 | 0 | 10 | 33 | 83 |
| 17 | PZ13_Bujnicki_10 | 3.04 | 98 | 0 | 64 | 5 | 1 | 7 | 21 | 98 |
| 18 | PZ13_Chen_4 | 0.00 | 100 | 4 | 47 | 2 | 0 | 11 | 37 | 101 |
| 19 | PZ13_Dokholyan_3 | 11.28 | 66 | 0 | 76 | 0 | 0 | 4 | 21 | 101 |
| 20 | PZ13_Dokholyan_2 | 10.41 | 69 | 0 | 90 | 0 | 0 | 2 | 18 | 110 |
| 21 | PZ13_Bujnicki_9 | 1.74 | 99 | 1 | 79 | 5 | 0 | 10 | 18 | 113 |
| 22 | PZ13_Ding_10 | 11.28 | 66 | 3 | 89 | 0 | 0 | 5 | 20 | 117 |

| | | | | | | | | | | |
|---|---|---|---|---|---|---|---|---|---|---|
| 23 | PZ13_Chen_8 | 2.17 | 99 | 8 | 64 | 1 | 0 | 8 | 38 | 119 |
| 24 | PZ13_Chen_7 | 0.43 | 99 | 3 | 62 | 1 | 0 | 11 | 45 | 122 |
| 25 | PZ13_Ding_7 | 18.22 | 37 | 2 | 97 | 0 | 0 | 4 | 22 | 125 |
| 26 | PZ13_Ding_4 | 10.41 | 69 | 1 | 103 | 0 | 0 | 2 | 20 | 126 |
| 27 | PZ13_Chen_5 | 0.43 | 99 | 8 | 62 | 4 | 0 | 9 | 45 | 128 |
| 28 | PZ13_Ding_5 | 13.02 | 58 | 4 | 99 | 0 | 0 | 5 | 20 | 128 |
| 29 | PZ13_Ding_1 | 13.45 | 56 | 2 | 107 | 0 | 0 | 2 | 18 | 129 |
| 30 | PZ13_Chen_9 | 0.43 | 99 | 25 | 57 | 4 | 0 | 8 | 36 | 130 |
| 31 | PZ13_Bujnicki_6 | 6.51 | 89 | 0 | 118 | 0 | 0 | 2 | 11 | 131 |
| 32 | PZ13_Chen_2 | 0.43 | 99 | 10 | 55 | 5 | 0 | 15 | 49 | 134 |
| 33 | PZ13_Ding_9 | 10.85 | 67 | 2 | 109 | 0 | 0 | 9 | 21 | 141 |
| 34 | PZ13_Ding_6 | 12.15 | 62 | 5 | 118 | 0 | 0 | 1 | 18 | 142 |
| 35 | PZ13_Ding_3 | 13.02 | 58 | 3 | 116 | 0 | 0 | 4 | 21 | 144 |
| 36 | PZ13_Ding_8 | 6.94 | 87 | 3 | 115 | 0 | 0 | 4 | 23 | 145 |
| 37 | PZ13_Dokholyan_1 | 13.88 | 54 | 0 | 109 | 0 | 0 | 4 | 32 | 145 |
| 38 | PZ13_Dokholyan_4 | 13.45 | 56 | 0 | 109 | 2 | 0 | 11 | 31 | 153 |
| 39 | PZ13_Xiao_8 | 1.30 | 99 | 4 | 103 | 3 | 0 | 12 | 31 | 153 |
| 40 | PZ13_Dokholyan_5 | 14.75 | 50 | 0 | 108 | 0 | 0 | 11 | 35 | 154 |
| 41 | PZ13_Xiao_9 | 2.17 | 99 | 4 | 100 | 3 | 0 | 14 | 33 | 154 |
| 42 | PZ13_Xiao_10 | 1.30 | 99 | 0 | 105 | 2 | 0 | 12 | 36 | 155 |
| 43 | PZ13_Xiao_6 | 1.74 | 99 | 1 | 101 | 3 | 0 | 12 | 39 | 156 |
| 44 | PZ13_Ding_2 | 19.09 | 35 | 6 | 125 | 0 | 0 | 6 | 22 | 159 |
| 45 | PZ13_Bujnicki_7 | 12.15 | 62 | 5 | 178 | 0 | 3 | 5 | 16 | 207 |
| 46 | PZ13_Bujnicki_2 | 20.39 | 31 | 6 | 179 | 0 | 4 | 7 | 22 | 218 |
| 47 | PZ13_Bujnicki_5 | 9.11 | 76 | 11 | 176 | 0 | 5 | 7 | 20 | 219 |
| 48 | PZ13_Bujnicki_3 | 22.13 | 27 | 10 | 191 | 0 | 3 | 2 | 22 | 228 |
| 49 | PZ13_Bujnicki_4 | 41.25 | 7 | 20 | 230 | 0 | 5 | 4 | 13 | 272 |
| 50 | PZ13_Xiao_4 | 0.00 | 100 | 77 | 175 | 3 | 0 | 6 | 14 | 275 |
| 51 | PZ13_Xiao_1 | 4.42 | 95 | 68 | 191 | 3 | 0 | 6 | 24 | 292 |
| 52 | PZ13_Xiao_7 | 32.13 | 13 | 37 | 196 | 3 | 3 | 19 | 38 | 296 |
| 53 | PZ13_Bujnicki_1 | 37.38 | 9 | 23 | 251 | 0 | 9 | 10 | 25 | 318 |
| 54 | PZ13_Xiao_5 | 26.46 | 19 | 96 | 190 | 4 | 1 | 10 | 21 | 322 |
| 55 | PZ13_Xiao_2 | 27.74 | 17 | 82 | 217 | 6 | 1 | 8 | 28 | 342 |

| | | | | | | | | | | |
|---|---|---|---|---|---|---|---|---|---|---|
| 56 | PZ13_Xiao_3 | 16.29 | 44 | 93 | 217 | 6 | 3 | 10 | 33 | 362 |
| | Average | 9.89 | 72.43 | 11.45 | 91.98 | 1.38 | 0.71 | 6.43 | 25.16 | 137.11 |
| | Standard deviation | 9.55 | 28.37 | 23.92 | 69.71 | 1.92 | 1.70 | 4.21 | 9.26 | 90.06 |
| | Median | 8.68 | 78.00 | 2.50 | 98.00 | 0.00 | 0.00 | 5.50 | 22.00 | 128.50 |

**Table S84**. MolProbity report on structures in Puzzle 14a.

| No. | 3D RNA model | (a) Clash score | (b) Rank-ing | #Bad bonds | #Bad angles | #Chiral handedness swaps | #Tetrahedral geometry out-liers | #Probably wrong sugar puck-ers: | (c) #Bad back-bone confor-mations | Total |
|---|---|---|---|---|---|---|---|---|---|---|
| 1 | PZ14_AdamiakPostExp_1 | 6.08 | 90 | 0 | 0 | 0 | 0 | 0 | 8 | 8 |
| 2 | PZ14_AdamiakPreExp_1 | 5.57 | 92 | 0 | 0 | 0 | 0 | 0 | 8 | 8 |
| 3 | PZ14_refstructure_bound | 4.56 | 95 | 1 | 0 | 0 | 0 | 0 | 10 | 11 |
| 4 | PZ14_AdamiakPreExp_2 | 8.61 | 79 | 0 | 0 | 0 | 0 | 1 | 11 | 12 |
| 5 | PZ14_AdamiakPostExp_2 | 14.69 | 51 | 0 | 0 | 0 | 0 | 0 | 14 | 14 |
| 6 | PZ14_DasPostExp_10 | 5.57 | 92 | 0 | 0 | 0 | 0 | 2 | 12 | 14 |
| 7 | PZ14_DasPreExp_5 | 5.57 | 92 | 0 | 0 | 0 | 0 | 2 | 12 | 14 |
| 8 | PZ14_DasPostExp_9 | 6.08 | 90 | 0 | 1 | 0 | 0 | 0 | 14 | 15 |
| 9 | PZ14_DasPreExp_10 | 6.08 | 90 | 0 | 1 | 0 | 0 | 0 | 14 | 15 |
| 10 | PZ14_DasPostExp_3 | 14.18 | 53 | 1 | 3 | 0 | 0 | 2 | 12 | 18 |
| 11 | PZ14_DasPostExp_4 | 5.07 | 93 | 0 | 2 | 0 | 0 | 1 | 16 | 19 |
| 12 | PZ14_DasPreExp_8 | 6.59 | 89 | 0 | 7 | 0 | 0 | 1 | 11 | 19 |
| 13 | PZ14_DasPreExp_9 | 6.08 | 90 | 0 | 5 | 0 | 0 | 1 | 13 | 19 |
| 14 | PZ14_DasPostExp_5 | 6.08 | 90 | 0 | 4 | 0 | 0 | 0 | 16 | 20 |
| 15 | PZ14_DasPostExp_7 | 2.53 | 98 | 0 | 5 | 0 | 0 | 1 | 14 | 20 |
| 16 | PZ14_DasPreExp_2 | 2.53 | 98 | 0 | 5 | 0 | 0 | 1 | 14 | 20 |
| 17 | PZ14_DasPreExp_7 | 12.16 | 62 | 1 | 2 | 0 | 0 | 1 | 16 | 20 |
| 18 | PZ14_DasPostExp_2 | 8.61 | 79 | 0 | 0 | 0 | 0 | 1 | 21 | 22 |
| 19 | PZ14_DasPreExp_6 | 11.65 | 64 | 0 | 1 | 0 | 0 | 3 | 18 | 22 |
| 20 | PZ14_DasPostExp_6 | 7.09 | 86 | 0 | 3 | 0 | 0 | 2 | 19 | 24 |
| 21 | PZ14_DasPreExp_1 | 7.09 | 86 | 0 | 3 | 0 | 0 | 2 | 19 | 24 |
| 22 | PZ14_DasPostExp_1 | 12.16 | 62 | 0 | 4 | 0 | 0 | 3 | 19 | 26 |
| 23 | PZ14_BujnickiPreExp_4 | 11.65 | 64 | 1 | 6 | 2 | 0 | 3 | 16 | 28 |

| | | | | | | | | | | |
|---|---|---|---|---|---|---|---|---|---|---|
| 24 | PZ14_ChenPostExp_1 | 0.00 | 100 | 0 | 8 | 0 | 0 | 4 | 16 | 28 |
| 25 | PZ14_DasPostExp_8 | 11.66 | 64 | 0 | 6 | 0 | 0 | 2 | 21 | 29 |
| 26 | PZ14_DasPreExp_3 | 11.66 | 64 | 0 | 6 | 0 | 0 | 2 | 21 | 29 |
| 27 | PZ14_DasPreExp_4 | 13.68 | 55 | 0 | 6 | 0 | 0 | 1 | 26 | 33 |
| 28 | PZ14_BujnickiPreExp_5 | 0.00 | 100 | 0 | 13 | 2 | 0 | 3 | 17 | 35 |
| 29 | PZ14_ChenPostExp_2 | 0.00 | 100 | 0 | 16 | 0 | 0 | 2 | 23 | 41 |
| 30 | PZ14_ChenPostExp_4 | 0.00 | 100 | 0 | 18 | 0 | 0 | 2 | 24 | 44 |
| 31 | PZ14_ChenPostExp_5 | 0.00 | 100 | 1 | 18 | 1 | 0 | 2 | 24 | 46 |
| 32 | PZ14_ChenPostExp_8 | 0.00 | 100 | 2 | 27 | 0 | 0 | 5 | 16 | 50 |
| 33 | PZ14_BujnickiPreExp_6 | 11.35 | 65 | 0 | 23 | 5 | 0 | 8 | 19 | 55 |
| 34 | PZ14_ChenPostExp_9 | 0.00 | 100 | 6 | 27 | 0 | 1 | 3 | 20 | 57 |
| 35 | PZ14_ChenPostExp_10 | 0.00 | 100 | 12 | 25 | 0 | 0 | 4 | 17 | 58 |
| 36 | PZ14_ChenPostExp_7 | 0.00 | 100 | 12 | 28 | 0 | 0 | 3 | 20 | 63 |
| 37 | PZ14_ChenPostExp_6 | 0.00 | 100 | 7 | 35 | 0 | 0 | 5 | 17 | 64 |
| 38 | PZ14_BujnickiPostExp_3 | 12.66 | 60 | 2 | 46 | 9 | 0 | 10 | 21 | 88 |
| 39 | PZ14_ChenPostExp_3 | 0.00 | 100 | 22 | 45 | 0 | 0 | 5 | 18 | 90 |
| 40 | PZ14_BujnickiPreExp_7 | 8.61 | 79 | 0 | 52 | 10 | 0 | 12 | 25 | 99 |
| 41 | PZ14_DingPreExp_3 | 16.21 | 45 | 5 | 85 | 0 | 0 | 3 | 18 | 111 |
| 42 | PZ14_DingPostExp_3 | 10.64 | 68 | 6 | 88 | 0 | 0 | 2 | 20 | 116 |
| 43 | PZ14_DingPreExp_5 | 12.66 | 60 | 1 | 96 | 0 | 0 | 4 | 15 | 116 |
| 44 | PZ14_DingPostExp_4 | 15.70 | 47 | 7 | 93 | 0 | 0 | 2 | 17 | 119 |
| 45 | PZ14_DingPreExp_2 | 16.21 | 45 | 7 | 90 | 0 | 0 | 3 | 19 | 119 |
| 46 | PZ14_DingPostExp_7 | 20.77 | 30 | 6 | 94 | 0 | 0 | 3 | 18 | 121 |
| 47 | PZ14_DingPostExp_5 | 21.28 | 29 | 5 | 93 | 0 | 0 | 4 | 20 | 122 |
| 48 | PZ14_DingPreExp_4 | 17.73 | 39 | 5 | 98 | 0 | 0 | 3 | 16 | 122 |
| 49 | PZ14_DingPostExp_6 | 17.22 | 41 | 6 | 100 | 0 | 0 | 3 | 14 | 123 |
| 50 | PZ14_DingPostExp_10 | 13.68 | 55 | 4 | 94 | 0 | 0 | 4 | 24 | 126 |
| 51 | PZ14_DingPreExp_7 | 16.21 | 45 | 3 | 106 | 0 | 0 | 2 | 20 | 131 |
| 52 | PZ14_DingPreExp_8 | 13.68 | 55 | 4 | 95 | 0 | 0 | 8 | 25 | 132 |
| 53 | PZ14_DingPostExp_9 | 11.14 | 66 | 0 | 107 | 0 | 0 | 3 | 23 | 133 |
| 54 | PZ14_DingPostExp_2 | 13.68 | 55 | 4 | 109 | 0 | 1 | 5 | 17 | 136 |
| 55 | PZ14_DingPreExp_6 | 8.61 | 79 | 2 | 108 | 0 | 0 | 5 | 21 | 136 |
| 56 | PZ14_DingPostExp_1 | 11.14 | 66 | 1 | 107 | 0 | 0 | 5 | 25 | 138 |

| | | | | | | | | | | |
|---|---|---|---|---|---|---|---|---|---|---|
| 57 | PZ14_DingPreExp_1 | 13.68 | 55 | 4 | 110 | 0 | 0 | 4 | 24 | 142 |
| 58 | PZ14_DingPostExp_8 | 16.20 | 45 | 5 | 125 | 0 | 0 | 1 | 15 | 146 |
| 59 | PZ14_BujnickiPostExp_1 | 22.88 | 25 | 27 | 103 | 5 | 1 | 9 | 21 | 166 |
| 60 | PZ14_BujnickiPostExp_5 | 23.30 | 24 | 20 | 92 | 12 | 3 | 16 | 31 | 174 |
| 61 | PZ14_BujnickiPostExp_4 | 28.40 | 17 | 36 | 148 | 2 | 4 | 6 | 16 | 212 |
| 62 | PZ14_BujnickiPostExp_2 | 29.38 | 16 | 31 | 149 | 5 | 0 | 10 | 23 | 218 |
| | Average | 9.94 | 70.63 | 4.15 | 44.21 | 0.85 | 0.16 | 3.31 | 17.97 | 70.65 |
| | Standard deviation | 7.22 | 24.80 | 7.66 | 47.26 | 2.46 | 0.66 | 3.09 | 4.66 | 56.97 |
| | Median | 10.89 | 67.00 | 1.00 | 20.50 | 0.00 | 0.00 | 3.00 | 18.00 | 48.00 |

**Table S85**. MolProbity report on structures in Puzzle 14b.

| No. | 3D RNA model | (a) Clash score | (b) Ranking | #Bad bonds | #Bad angles | #Chiral handedness swaps | #Tetrahedral geometry outliers | #Probably wrong sugar puckers: | (c) #Bad backbone conformations | Total |
|---|---|---|---|---|---|---|---|---|---|---|
| 1 | PZ14_DasPostExp_5 | 5.58 | 92 | 0 | 2 | 0 | 0 | 2 | 9 | 13 |
| 2 | PZ14_refstructure_free | 13.38 | 56 | 0 | 0 | 0 | 0 | 1 | 13 | 14 |
| 3 | PZ14_DasPostExp_7 | 5.58 | 92 | 0 | 5 | 0 | 0 | 1 | 11 | 17 |
| 4 | PZ14_DasPreExp_2 | 5.58 | 92 | 0 | 5 | 0 | 0 | 1 | 11 | 17 |
| 5 | PZ14_DasPreExp_7 | 5.58 | 92 | 0 | 2 | 0 | 0 | 2 | 15 | 19 |
| 6 | PZ14_DasPostExp_3 | 5.58 | 92 | 1 | 2 | 0 | 0 | 0 | 17 | 20 |
| 7 | PZ14_DasPostExp_4 | 6.60 | 89 | 0 | 2 | 0 | 0 | 2 | 16 | 20 |
| 8 | PZ14_DasPostExp_6 | 16.24 | 45 | 1 | 1 | 0 | 0 | 4 | 14 | 20 |
| 9 | PZ14_DasPreExp_1 | 16.24 | 45 | 1 | 1 | 0 | 0 | 4 | 14 | 20 |
| 10 | PZ14_DasPreExp_9 | 4.57 | 95 | 0 | 8 | 0 | 0 | 1 | 12 | 21 |
| 11 | PZ14_DasPreExp_8 | 4.57 | 95 | 0 | 7 | 0 | 0 | 1 | 14 | 22 |
| 12 | PZ14_DasPostExp_10 | 6.09 | 90 | 0 | 1 | 0 | 0 | 2 | 20 | 23 |
| 13 | PZ14_DasPreExp_5 | 6.09 | 90 | 0 | 1 | 0 | 0 | 2 | 20 | 23 |
| 14 | PZ14_DasPostExp_9 | 6.60 | 89 | 1 | 8 | 0 | 1 | 2 | 12 | 24 |
| 15 | PZ14_DasPreExp_10 | 6.60 | 89 | 1 | 8 | 0 | 1 | 2 | 12 | 24 |
| 16 | PZ14_DasPostExp_8 | 12.20 | 62 | 1 | 5 | 0 | 0 | 1 | 18 | 25 |
| 17 | PZ14_DasPreExp_4 | 12.20 | 62 | 1 | 5 | 0 | 0 | 1 | 18 | 25 |
| 18 | PZ14_DasPreExp_3 | 8.63 | 79 | 0 | 0 | 0 | 0 | 2 | 24 | 26 |

| | | | | | | | | | | |
|---|---|---|---|---|---|---|---|---|---|---|
| 19 | PZ14_DasPostExp_2 | 10.66 | 68 | 0 | 2 | 0 | 1 | 2 | 22 | 27 |
| 20 | PZ14_DasPostExp_1 | 10.15 | 71 | 1 | 4 | 0 | 1 | 4 | 24 | 34 |
| 21 | PZ14_DasPreExp_6 | 10.15 | 71 | 1 | 4 | 0 | 1 | 4 | 24 | 34 |
| 22 | PZ14_ChenPostExp_1 | 0.00 | 100 | 7 | 29 | 0 | 5 | 5 | 16 | 62 |
| 23 | PZ14_BujnickiPreExp_4 | 7.61 | 83 | 0 | 31 | 6 | 1 | 7 | 20 | 65 |
| 24 | PZ14_ChenPostExp_2 | 0.00 | 100 | 5 | 34 | 0 | 2 | 2 | 23 | 66 |
| 25 | PZ14_BujnickiPreExp_2 | 12.12 | 62 | 0 | 27 | 10 | 0 | 10 | 21 | 68 |
| 26 | PZ14_BujnickiPostExp_1 | 14.74 | 50 | 4 | 38 | 4 | 0 | 7 | 16 | 69 |
| 27 | PZ14_BujnickiPreExp_3 | 11.11 | 66 | 1 | 29 | 9 | 0 | 11 | 24 | 74 |
| 28 | PZ14_ChenPostExp_4 | 0.00 | 100 | 8 | 37 | 0 | 6 | 2 | 23 | 76 |
| 29 | PZ14_ChenPostExp_8 | 0.00 | 100 | 9 | 44 | 0 | 4 | 5 | 16 | 78 |
| 30 | PZ14_ChenPostExp_9 | 0.00 | 100 | 12 | 44 | 0 | 7 | 2 | 19 | 84 |
| 31 | PZ14_ChenPostExp_5 | 0.00 | 100 | 11 | 42 | 1 | 5 | 2 | 24 | 85 |
| 32 | PZ14_ChenPostExp_6 | 0.00 | 100 | 14 | 53 | 0 | 2 | 5 | 18 | 92 |
| 33 | PZ14_ChenPostExp_10 | 0.00 | 100 | 22 | 46 | 0 | 5 | 4 | 18 | 95 |
| 34 | PZ14_ChenPostExp_7 | 0.00 | 100 | 18 | 49 | 0 | 7 | 4 | 20 | 98 |
| 35 | PZ14_BujnickiPostExp_2 | 22.85 | 25 | 10 | 70 | 5 | 0 | 6 | 12 | 103 |
| 36 | PZ14_DingPostExp_1 | 14.21 | 53 | 3 | 83 | 0 | 0 | 3 | 16 | 105 |
| 37 | PZ14_DingPostExp_2 | 15.23 | 48 | 5 | 87 | 0 | 0 | 0 | 15 | 107 |
| 38 | PZ14_ChenPostExp_3 | 0.00 | 100 | 29 | 54 | 0 | 6 | 5 | 18 | 112 |
| 39 | PZ14_DingPreExp_4 | 16.75 | 43 | 5 | 91 | 0 | 0 | 1 | 17 | 114 |
| 40 | PZ14_BujnickiPostExp_3 | 12.70 | 60 | 21 | 71 | 3 | 0 | 5 | 16 | 116 |
| 41 | PZ14_DingPostExp_5 | 7.61 | 83 | 5 | 100 | 0 | 0 | 1 | 11 | 117 |
| 42 | PZ14_DingPostExp_6 | 13.20 | 57 | 3 | 94 | 0 | 0 | 3 | 18 | 118 |
| 43 | PZ14_DingPostExp_3 | 15.23 | 48 | 4 | 97 | 0 | 0 | 3 | 19 | 123 |
| 44 | PZ14_DingPreExp_1 | 11.68 | 64 | 5 | 99 | 0 | 0 | 5 | 18 | 127 |
| 45 | PZ14_DingPostExp_9 | 11.17 | 66 | 4 | 103 | 0 | 0 | 5 | 22 | 134 |
| 46 | PZ14_DingPostExp_4 | 13.71 | 55 | 6 | 108 | 1 | 0 | 5 | 23 | 143 |
| 47 | PZ14_DingPostExp_7 | 9.64 | 73 | 4 | 120 | 0 | 0 | 3 | 16 | 143 |
| 48 | PZ14_DingPreExp_2 | 11.17 | 66 | 4 | 109 | 0 | 0 | 8 | 24 | 145 |
| 49 | PZ14_DingPreExp_3 | 13.71 | 55 | 7 | 106 | 0 | 0 | 8 | 24 | 145 |
| 50 | PZ14_BujnickiPreExp_1 | 4.04 | 96 | 5 | 95 | 9 | 2 | 11 | 25 | 147 |
| 51 | PZ14_DingPostExp_10 | 11.68 | 64 | 1 | 122 | 0 | 0 | 5 | 22 | 150 |

| | | | | | | | | | | |
|---|---|---|---|---|---|---|---|---|---|---|
| 52 | PZ14_DingPostExp_8 | 14.21 | 53 | 6 | 109 | 0 | 0 | 8 | 29 | 152 |
| 53 | PZ14_BujnickiPostExp_4 | 23.86 | 23 | 20 | 101 | 13 | 3 | 16 | 31 | 184 |
| | Average | 8.81 | 74.51 | 5.04 | 45.19 | 1.15 | 1.13 | 3.92 | 18.38 | 74.81 |
| | Standard deviation | 6.00 | 21.47 | 6.62 | 42.04 | 2.94 | 2.06 | 3.18 | 4.86 | 49.58 |
| | Median | 9.64 | 73.00 | 3.00 | 37.00 | 0.00 | 0.00 | 3.00 | 18.00 | 74.00 |

**Table S86**. MolProbity report on structures in Puzzle 15.

| **No.** | **3D RNA model** | **(a) Clash score** | **(b) Rank-ing** | **#Bad bonds** | **#Bad angles** | **#Chiral handedness swaps** | **#Tetrahedral geometry out-liers** | **#Probably wrong sugar puck-ers:** | **(c) #Bad back-bone confor-mations** | **Total** |
|---|---|---|---|---|---|---|---|---|---|---|
| 1 | PZ15_Adamiak_10 | 8.64 | 79 | 0 | 0 | 0 | 0 | 0 | 6 | 6 |
| 2 | PZ15_Adamiak_3 | 5.46 | 92 | 0 | 0 | 0 | 0 | 0 | 6 | 6 |
| 3 | PZ15_RNAComposer1_2 | 5.46 | 92 | 0 | 0 | 0 | 0 | 0 | 6 | 6 |
| 4 | PZ15_refstructure | 0.91 | 99 | 1 | 0 | 0 | 0 | 0 | 5 | 6 |
| 5 | PZ15_Adamiak_2 | 6.37 | 89 | 0 | 0 | 0 | 0 | 0 | 7 | 7 |
| 6 | PZ15_RNAComposer1_1 | 5.91 | 91 | 0 | 0 | 0 | 0 | 0 | 7 | 7 |
| 7 | PZ15_Adamiak_1 | 26.39 | 19 | 0 | 0 | 0 | 0 | 0 | 11 | 11 |
| 8 | PZ15_Adamiak_4 | 10.92 | 67 | 0 | 0 | 0 | 0 | 0 | 11 | 11 |
| 9 | PZ15_Adamiak_5 | 10.46 | 69 | 0 | 0 | 0 | 0 | 0 | 11 | 11 |
| 10 | PZ15_Adamiak_8 | 12.74 | 60 | 0 | 1 | 0 | 0 | 1 | 9 | 11 |
| 11 | PZ15_Adamiak_9 | 11.83 | 64 | 0 | 0 | 0 | 0 | 0 | 11 | 11 |
| 12 | PZ15_RNAComposer2_1 | 10.46 | 69 | 0 | 0 | 0 | 0 | 0 | 11 | 11 |
| 13 | PZ15_RNAComposer2_2 | 9.55 | 74 | 0 | 0 | 0 | 0 | 0 | 11 | 11 |
| 14 | PZ15_Adamiak_7 | 9.55 | 74 | 0 | 0 | 0 | 0 | 0 | 13 | 13 |
| 15 | PZ15_Adamiak_6 | 10.46 | 69 | 0 | 0 | 0 | 0 | 1 | 13 | 14 |
| 16 | PZ15_FARFAR2_10 | 7.73 | 83 | 0 | 4 | 0 | 0 | 0 | 11 | 15 |
| 17 | PZ15_FARFAR2_6 | 7.28 | 85 | 0 | 2 | 0 | 0 | 1 | 13 | 16 |
| 18 | PZ15_FARFAR2_2 | 7.28 | 85 | 0 | 2 | 0 | 0 | 0 | 15 | 17 |
| 19 | PZ15_FARFAR2_3 | 9.10 | 77 | 0 | 6 | 0 | 0 | 2 | 9 | 17 |
| 20 | PZ15_FARFAR2_1 | 10.46 | 69 | 1 | 5 | 0 | 0 | 1 | 11 | 18 |
| 21 | PZ15_FARFAR2_7 | 10.46 | 69 | 1 | 2 | 0 | 0 | 1 | 15 | 19 |
| 22 | PZ15_FARFAR2_8 | 5.00 | 94 | 0 | 2 | 0 | 0 | 0 | 17 | 19 |

| | | | | | | | | | | |
|---|---|---|---|---|---|---|---|---|---|---|
| 23 | PZ15_FARFAR2_9 | 9.10 | 77 | 1 | 4 | 0 | 1 | 4 | 10 | 20 |
| 24 | PZ15_FARFAR2_5 | 9.55 | 74 | 0 | 2 | 0 | 0 | 2 | 17 | 21 |
| 25 | PZ15_FARFAR2_4 | 12.74 | 60 | 0 | 1 | 0 | 0 | 0 | 24 | 25 |
| 26 | PZ15_Chen_3 | 0.45 | 99 | 0 | 17 | 0 | 0 | 2 | 13 | 32 |
| 27 | PZ15_Chen_2 | 0.00 | 100 | 2 | 17 | 0 | 0 | 5 | 19 | 43 |
| 28 | PZ15_Chen_1 | 0.45 | 99 | 4 | 24 | 0 | 0 | 4 | 22 | 54 |
| 29 | PZ15_Chen_6 | 0.46 | 99 | 9 | 42 | 0 | 0 | 5 | 24 | 80 |
| 30 | PZ15_Chen_4 | 0.00 | 100 | 4 | 39 | 0 | 0 | 7 | 36 | 86 |
| 31 | PZ15_Chen_5 | 0.00 | 100 | 11 | 56 | 0 | 1 | 5 | 23 | 96 |
| 32 | PZ15_Chen_9 | 0.91 | 99 | 13 | 60 | 3 | 0 | 8 | 42 | 126 |
| 33 | PZ15_Chen_10 | 0.91 | 99 | 30 | 66 | 2 | 3 | 8 | 22 | 131 |
| 34 | PZ15_Chen_7 | 3.64 | 97 | 25 | 70 | 0 | 0 | 10 | 26 | 131 |
| 35 | PZ15_Chen_8 | 1.36 | 99 | 27 | 63 | 1 | 1 | 11 | 32 | 135 |
| 36 | PZ15_SimRNA2_10 | 72.86 | 1 | 53 | 158 | 0 | 2 | 4 | 17 | 234 |
| 37 | PZ15_SimRNA2_2 | 73.18 | 0 | 60 | 153 | 0 | 1 | 4 | 18 | 236 |
| 38 | PZ15_SimRNA1_7 | 70.55 | 1 | 68 | 149 | 0 | 4 | 3 | 19 | 243 |
| 39 | PZ15_SimRNA1_1 | 94.72 | 0 | 67 | 160 | 1 | 2 | 3 | 18 | 251 |
| 40 | PZ15_SimRNA2_7 | 81.09 | 0 | 65 | 160 | 2 | 3 | 4 | 21 | 255 |
| 41 | PZ15_3dRNA2_9 | 34.56 | 11 | 18 | 163 | 20 | 9 | 15 | 32 | 257 |
| 42 | PZ15_3dRNA2_10 | 34.11 | 11 | 21 | 163 | 20 | 9 | 14 | 32 | 259 |
| 43 | PZ15_SimRNA1_3 | 78.53 | 0 | 70 | 164 | 1 | 2 | 2 | 21 | 260 |
| 44 | PZ15_3dRNA2_1 | 35.02 | 11 | 22 | 164 | 20 | 9 | 14 | 32 | 261 |
| 45 | PZ15_SimRNA2_1 | 38.25 | 9 | 67 | 175 | 2 | 0 | 2 | 15 | 261 |
| 46 | PZ15_SimRNA2_9 | 84.58 | 0 | 66 | 164 | 1 | 5 | 5 | 20 | 261 |
| 47 | PZ15_3dRNA2_2 | 33.65 | 12 | 22 | 165 | 20 | 9 | 15 | 31 | 262 |
| 48 | PZ15_3dRNA2_8 | 34.56 | 11 | 20 | 168 | 20 | 9 | 14 | 32 | 263 |
| 49 | PZ15_SimRNA1_8 | 68.27 | 1 | 78 | 158 | 1 | 2 | 3 | 22 | 264 |
| 50 | PZ15_3dRNA2_7 | 32.74 | 13 | 23 | 168 | 21 | 8 | 14 | 31 | 265 |
| 51 | PZ15_SimRNA1_5 | 85.23 | 0 | 69 | 168 | 2 | 1 | 1 | 26 | 267 |
| 52 | PZ15_3dRNA2_4 | 32.74 | 13 | 24 | 171 | 20 | 9 | 14 | 31 | 269 |
| 53 | PZ15_3dRNA2_3 | 32.74 | 13 | 24 | 176 | 21 | 8 | 15 | 31 | 275 |
| 54 | PZ15_3dRNA2_5 | 32.29 | 13 | 25 | 176 | 21 | 7 | 16 | 31 | 276 |
| 55 | PZ15_SimRNA2_6 | 78.82 | 0 | 80 | 169 | 1 | 3 | 4 | 20 | 277 |

| | | | | | | | | | | |
|---|---|---|---|---|---|---|---|---|---|---|
| 56 | PZ15_3dRNA2_6 | 31.83 | 14 | 25 | 177 | 21 | 8 | 15 | 32 | 278 |
| 57 | PZ15_SimRNA2_5 | 80.15 | 0 | 74 | 182 | 0 | 2 | 5 | 15 | 278 |
| 58 | PZ15_SimRNA1_6 | 97.77 | 0 | 74 | 177 | 1 | 4 | 6 | 17 | 279 |
| 59 | PZ15_SimRNA2_3 | 85.12 | 0 | 78 | 174 | 1 | 3 | 2 | 22 | 280 |
| 60 | PZ15_SimRNA1_2 | 96.00 | 0 | 66 | 182 | 0 | 7 | 4 | 23 | 282 |
| 61 | PZ15_SimRNA2_4 | 63.21 | 1 | 67 | 190 | 0 | 5 | 5 | 16 | 283 |
| 62 | PZ15_SimRNA1_10 | 107.22 | 0 | 67 | 176 | 1 | 4 | 11 | 26 | 285 |
| 63 | PZ15_SimRNA2_8 | 59.49 | 2 | 73 | 178 | 2 | 4 | 6 | 25 | 288 |
| 64 | PZ15_SimRNA1_4 | 88.76 | 0 | 78 | 181 | 1 | 2 | 5 | 23 | 290 |
| 65 | PZ15_SimRNA1_9 | 64.78 | 1 | 71 | 198 | 1 | 1 | 2 | 22 | 295 |
| 66 | PZ15_FARFAR1_1 | Rejected as unacceptable - Bond distance > max_reasonable_bond_distance: 128.024 > 50: distance: 1553 - 1554: 126.362 | | | | | | | | |
| 67 | PZ15_FARFAR1_10 | Rejected as unacceptable - Bond distance > max_reasonable_bond_distance: 110.638 > 50: distance: 1553 - 1554: 110.120 | | | | | | | | |
| 68 | PZ15_FARFAR1_2 | Rejected as unacceptable - Bond distance > max_reasonable_bond_distance: 110.873 > 50: distance: 1553 - 1554: 110.360 | | | | | | | | |
| 69 | PZ15_FARFAR1_3 | Rejected as unacceptable - Bond distance > max_reasonable_bond_distance: 111.621 > 50: distance: 1553 - 1554: 111.175 | | | | | | | | |
| 70 | PZ15_FARFAR1_4 | Rejected as unacceptable - Bond distance > max_reasonable_bond_distance: 123.221 > 50: distance: 1553 - 1554: 121.354 | | | | | | | | |
| 71 | PZ15_FARFAR1_5 | Rejected as unacceptable - Bond distance > max_reasonable_bond_distance: 123.078 > 50: distance: 1553 - 1554: 122.427 | | | | | | | | |
| 72 | PZ15_FARFAR1_6 | Rejected as unacceptable - Bond distance > max_reasonable_bond_distance: 119.902 > 50: distance: 1553 - 1554: 118.260 | | | | | | | | |
| 73 | PZ15_FARFAR1_7 | Rejected as unacceptable - Bond distance > max_reasonable_bond_distance: 126.282 > 50: distance: 1553 - 1554: 124.545 | | | | | | | | |
| 74 | PZ15_FARFAR1_8 | Rejected as unacceptable - Bond distance > max_reasonable_bond_distance: 134.966 > 50: distance: 1553 - 1554: 133.430 | | | | | | | | |
| 75 | PZ15_FARFAR1_9 | Rejected as unacceptable - Bond distance > max_reasonable_bond_distance: 120.129 > 50: distance: 1553 - 1554: 118.685 | | | | | | | | |
| | Average | 33.00 | 46.29 | 26.83 | 86.03 | 3.51 | 2.28 | 4.69 | 19.38 | 142.72 |
| | Standard deviation | 33.25 | 41.10 | 30.23 | 80.64 | 7.29 | 3.11 | 5.07 | 8.75 | 120.96 |
| | Median | 12.74 | 60.00 | 18.00 | 63.00 | 0.00 | 1.00 | 3.00 | 19.00 | 131.00 |

**Table S87**. MolProbity report on structures in Puzzle 17.

| No. | 3D RNA model | (a) Clash score | (b) Rank-ing | #Bad bonds | #Bad angles | #Chiral handedness swaps | #Tetrahedral geometry out-liers | #Probably wrong sugar puckers: | (c) #Bad back-bone confor-mations | Total |
|---|---|---|---|---|---|---|---|---|---|---|
| 1 | PZ17_RNAComposer1_3 | 9.55 | 74 | 0 | 0 | 0 | 0 | 1 | 7 | 8 |
| 2 | PZ17_RNAComposer1_5 | 15.08 | 49 | 0 | 0 | 0 | 0 | 1 | 9 | 10 |
| 3 | PZ17_RNAComposer2_2 | 9.55 | 74 | 0 | 0 | 0 | 0 | 0 | 12 | 12 |
| 4 | PZ17_RNAComposer2_5 | 12.56 | 60 | 0 | 0 | 0 | 0 | 0 | 12 | 12 |

| | | | | | | | | | | |
|---|---|---|---|---|---|---|---|---|---|---|
| 5 | PZ17_RNAComposer2_4 | 9.55 | 74 | 0 | 0 | 0 | 0 | 0 | 14 | 14 |
| 6 | PZ17_refstructure | 15.59 | 47 | 0 | 3 | 0 | 0 | 1 | 11 | 15 |
| 7 | PZ17_RNAComposer2_1 | 6.53 | 89 | 0 | 0 | 0 | 0 | 0 | 15 | 15 |
| 8 | PZ17_RNAComposer1_10 | 11.06 | 66 | 0 | 0 | 0 | 0 | 2 | 14 | 16 |
| 9 | PZ17_RNAComposer2_6 | 16.59 | 43 | 0 | 0 | 0 | 0 | 0 | 16 | 16 |
| 10 | PZ17_Das_1 | 3.02 | 98 | 0 | 6 | 0 | 0 | 0 | 11 | 17 |
| 11 | PZ17_Das_4 | 6.53 | 89 | 0 | 1 | 0 | 0 | 2 | 14 | 17 |
| 12 | PZ17_Das_9 | 6.53 | 89 | 0 | 1 | 0 | 0 | 3 | 13 | 17 |
| 13 | PZ17_RNAComposer1_1 | 10.55 | 68 | 0 | 0 | 0 | 0 | 1 | 16 | 17 |
| 14 | PZ17_RNAComposer1_4 | 10.55 | 68 | 0 | 0 | 0 | 0 | 1 | 16 | 17 |
| 15 | PZ17_RNAComposer2_8 | 11.06 | 66 | 0 | 0 | 0 | 0 | 0 | 17 | 17 |
| 16 | PZ17_RNAComposer2_9 | 11.06 | 66 | 0 | 0 | 0 | 0 | 1 | 16 | 17 |
| 17 | PZ17_Das_6 | 4.02 | 96 | 0 | 4 | 0 | 0 | 1 | 13 | 18 |
| 18 | PZ17_RNAComposer1_7 | 12.56 | 60 | 0 | 0 | 0 | 0 | 1 | 17 | 18 |
| 19 | PZ17_RNAComposer1_8 | 12.56 | 60 | 0 | 0 | 0 | 0 | 2 | 16 | 18 |
| 20 | PZ17_RNAComposer2_7 | 13.07 | 58 | 0 | 0 | 0 | 0 | 0 | 18 | 18 |
| 21 | PZ17_Das_5 | 8.54 | 79 | 0 | 1 | 0 | 0 | 3 | 15 | 19 |
| 22 | PZ17_DasExtraInfo_1 | 10.05 | 71 | 0 | 2 | 0 | 0 | 0 | 17 | 19 |
| 23 | PZ17_Das_8 | 7.04 | 86 | 0 | 4 | 0 | 0 | 1 | 15 | 20 |
| 24 | PZ17_RNAComposer2_10 | 10.05 | 71 | 0 | 0 | 0 | 0 | 1 | 19 | 20 |
| 25 | PZ17_Das_7 | 13.07 | 58 | 0 | 4 | 0 | 0 | 4 | 14 | 22 |
| 26 | PZ17_RNAComposer1_2 | 8.04 | 82 | 0 | 0 | 0 | 0 | 2 | 20 | 22 |
| 27 | PZ17_RNAComposer1_9 | 8.54 | 79 | 0 | 0 | 0 | 0 | 2 | 20 | 22 |
| 28 | PZ17_Adamiak_2 | 15.58 | 47 | 0 | 1 | 0 | 0 | 2 | 20 | 23 |
| 29 | PZ17_DasExtraInfo_3 | 12.06 | 62 | 1 | 3 | 0 | 0 | 2 | 18 | 24 |
| 30 | PZ17_Das_2 | 7.04 | 86 | 0 | 1 | 0 | 0 | 5 | 19 | 25 |
| 31 | PZ17_RNAComposer1_6 | 16.58 | 43 | 0 | 0 | 0 | 0 | 2 | 23 | 25 |
| 32 | PZ17_Das_3 | 10.05 | 71 | 1 | 1 | 0 | 0 | 2 | 22 | 26 |
| 33 | PZ17_Adamiak_3 | 11.06 | 66 | 0 | 0 | 0 | 0 | 1 | 26 | 27 |
| 34 | PZ17_DasExtraInfo_2 | 9.05 | 77 | 1 | 5 | 1 | 0 | 1 | 19 | 27 |
| 35 | PZ17_Adamiak_1 | 10.55 | 68 | 0 | 0 | 0 | 0 | 1 | 27 | 28 |
| 36 | PZ17_Bujnicki_4 | 0.00 | 100 | 0 | 17 | 1 | 0 | 2 | 10 | 30 |
| 37 | PZ17_Bujnicki_8 | 0.00 | 100 | 0 | 18 | 0 | 0 | 2 | 10 | 30 |

| | | | | | | | | | | |
|---|---|---|---|---|---|---|---|---|---|---|
| 38 | PZ17_Bujnicki_3 | 0.00 | 100 | 0 | 15 | 1 | 0 | 2 | 13 | 31 |
| 39 | PZ17_Das_10 | 13.57 | 56 | 0 | 3 | 0 | 0 | 1 | 27 | 31 |
| 40 | PZ17_Bujnicki_10 | 0.00 | 100 | 0 | 15 | 1 | 0 | 4 | 13 | 33 |
| 41 | PZ17_Bujnicki_2 | 0.00 | 100 | 0 | 15 | 1 | 0 | 4 | 13 | 33 |
| 42 | PZ17_Bujnicki_7 | 1.01 | 99 | 0 | 23 | 0 | 0 | 3 | 7 | 33 |
| 43 | PZ17_Bujnicki_6 | 0.00 | 100 | 0 | 21 | 0 | 0 | 1 | 13 | 35 |
| 44 | PZ17_Chen_10 | 0.50 | 99 | 0 | 12 | 1 | 0 | 4 | 19 | 36 |
| 45 | PZ17_Major_2 | 0.00 | 100 | 1 | 30 | 0 | 0 | 3 | 15 | 49 |
| 46 | PZ17_Major_5 | 0.00 | 100 | 0 | 28 | 0 | 0 | 2 | 22 | 52 |
| 47 | PZ17_Major_1 | 0.50 | 99 | 1 | 33 | 0 | 0 | 5 | 16 | 55 |
| 48 | PZ17_Major_4 | 0.00 | 100 | 0 | 34 | 0 | 0 | 9 | 20 | 63 |
| 49 | PZ17_Major_3 | 0.00 | 100 | 0 | 34 | 3 | 0 | 10 | 23 | 70 |
| 50 | PZ17_Chen_3 | 0.00 | 100 | 2 | 33 | 2 | 0 | 11 | 35 | 83 |
| 51 | PZ17_Ding_1 | 8.54 | 79 | 0 | 93 | 0 | 0 | 4 | 13 | 110 |
| 52 | PZ17_Ding_5 | 8.54 | 79 | 3 | 88 | 0 | 0 | 4 | 20 | 115 |
| 53 | PZ17_Ding_8 | 13.57 | 56 | 5 | 90 | 0 | 0 | 4 | 17 | 116 |
| 54 | PZ17_Major_6 | 1.01 | 99 | 0 | 67 | 5 | 0 | 13 | 32 | 117 |
| 55 | PZ17_Major_8 | 0.00 | 100 | 1 | 60 | 4 | 0 | 15 | 38 | 118 |
| 56 | PZ17_Dohkolyan_1 | 17.59 | 40 | 0 | 93 | 0 | 0 | 7 | 19 | 119 |
| 57 | PZ17_RNAComposer2_3 | 68.45 | 1 | 48 | 64 | 0 | 0 | 0 | 8 | 120 |
| 58 | PZ17_Ding_6 | 10.05 | 71 | 3 | 95 | 0 | 0 | 5 | 19 | 122 |
| 59 | PZ17_Ding_9 | 7.04 | 86 | 3 | 97 | 0 | 0 | 4 | 18 | 122 |
| 60 | PZ17_Dohkolyan_2 | 22.61 | 26 | 0 | 100 | 0 | 0 | 5 | 19 | 124 |
| 61 | PZ17_Ding_4 | 10.05 | 71 | 3 | 100 | 0 | 0 | 3 | 21 | 127 |
| 62 | PZ17_Ding_3 | 9.55 | 74 | 3 | 103 | 0 | 0 | 4 | 18 | 128 |
| 63 | PZ17_Ding_10 | 8.04 | 82 | 5 | 104 | 0 | 0 | 2 | 18 | 129 |
| 64 | PZ17_Dohkolyan_3 | 19.60 | 34 | 0 | 102 | 0 | 0 | 4 | 25 | 131 |
| 65 | PZ17_Ding_2 | 11.06 | 66 | 6 | 96 | 0 | 0 | 7 | 28 | 137 |
| 66 | PZ17_Major_7 | 2.01 | 99 | 1 | 81 | 4 | 0 | 15 | 38 | 139 |
| 67 | PZ17_Bujnicki_1 | 3.02 | 98 | 1 | 109 | 3 | 4 | 6 | 17 | 140 |
| 68 | PZ17_Bujnicki_9 | 3.02 | 98 | 1 | 109 | 3 | 4 | 6 | 17 | 140 |
| 69 | PZ17_Ding_7 | 8.54 | 79 | 3 | 110 | 0 | 0 | 9 | 28 | 150 |
| 70 | PZ17_Chen_7 | 4.52 | 95 | 35 | 78 | 2 | 1 | 10 | 29 | 155 |

| | | | | | | | | | | |
|---|---|---|---|---|---|---|---|---|---|---|
| 71 | PZ17_Xiao_7 | 0.50 | 99 | 1 | 73 | 31 | 1 | 24 | 44 | 174 |
| 72 | PZ17_Xiao_2 | 0.00 | 100 | 1 | 74 | 36 | 0 | 21 | 43 | 175 |
| 73 | PZ17_Xiao_1 | 12.56 | 60 | 11 | 94 | 20 | 0 | 16 | 35 | 176 |
| 74 | PZ17_Xiao_3 | 3.02 | 98 | 1 | 77 | 40 | 0 | 23 | 41 | 182 |
| 75 | PZ17_Major_9 | 13.57 | 56 | 8 | 97 | 10 | 3 | 24 | 49 | 191 |
| 76 | PZ17_SimRNA1_2 | 109.15 | 0 | 53 | 121 | 0 | 2 | 3 | 12 | 191 |
| 77 | PZ17_Xiao_4 | 4.02 | 96 | 1 | 88 | 32 | 0 | 30 | 42 | 193 |
| 78 | PZ17_Chen_8 | 1.51 | 99 | 41 | 108 | 2 | 2 | 14 | 34 | 201 |
| 79 | PZ17_Major_10 | 15.08 | 49 | 8 | 116 | 5 | 4 | 22 | 46 | 201 |
| 80 | PZ17_Bujnicki_5 | 5.03 | 94 | 6 | 164 | 2 | 5 | 10 | 26 | 213 |
| 81 | PZ17_SimRNA1_7 | 140.42 | 0 | 61 | 134 | 0 | 1 | 3 | 17 | 216 |
| 82 | PZ17_SimRNA1_9 | 141.93 | 0 | 61 | 138 | 0 | 2 | 1 | 14 | 216 |
| 83 | PZ17_SimRNA1_8 | 121.91 | 0 | 54 | 143 | 0 | 3 | 2 | 15 | 217 |
| 84 | PZ17_Xiao_10 | 4.52 | 95 | 1 | 94 | 47 | 0 | 28 | 47 | 217 |
| 85 | PZ17_SimRNA1_1 | 140.48 | 0 | 57 | 140 | 0 | 4 | 4 | 13 | 218 |
| 86 | PZ17_Xiao_5 | 27.14 | 18 | 21 | 118 | 23 | 0 | 18 | 38 | 218 |
| 87 | PZ17_Chen_2 | 101.35 | 0 | 47 | 146 | 0 | 1 | 9 | 17 | 220 |
| 88 | PZ17_Chen_6 | 177.18 | 0 | 51 | 140 | 0 | 2 | 9 | 18 | 220 |
| 89 | PZ17_SimRNA2_4 | 102.72 | 0 | 65 | 132 | 1 | 2 | 3 | 17 | 220 |
| 90 | PZ17_SimRNA2_2 | 134.24 | 0 | 63 | 135 | 0 | 5 | 2 | 16 | 221 |
| 91 | PZ17_SimRNA1_5 | 161.45 | 0 | 62 | 140 | 0 | 5 | 3 | 15 | 225 |
| 92 | PZ17_SimRNA1_3 | 176.65 | 0 | 61 | 148 | 0 | 3 | 3 | 13 | 228 |
| 93 | PZ17_SimRNA2_7 | 158.61 | 0 | 66 | 142 | 1 | 1 | 2 | 17 | 229 |
| 94 | PZ17_Chen_9 | 107.65 | 0 | 49 | 156 | 0 | 2 | 5 | 18 | 230 |
| 95 | PZ17_Chen_1 | 102.46 | 0 | 39 | 165 | 0 | 0 | 8 | 22 | 234 |
| 96 | PZ17_SimRNA2_9 | 160.97 | 0 | 62 | 147 | 1 | 3 | 6 | 16 | 235 |
| 97 | PZ17_Xiao_6 | 30.15 | 15 | 22 | 125 | 25 | 1 | 23 | 40 | 236 |
| 98 | PZ17_SimRNA1_6 | 139.63 | 0 | 67 | 152 | 1 | 2 | 2 | 14 | 238 |
| 99 | PZ17_Chen_5 | 156.64 | 0 | 49 | 169 | 1 | 0 | 7 | 19 | 245 |
| 100 | PZ17_Xiao_8 | 16.08 | 45 | 15 | 119 | 40 | 1 | 27 | 43 | 245 |
| 101 | PZ17_SimRNA2_1 | 151.48 | 0 | 59 | 170 | 2 | 1 | 2 | 15 | 249 |
| 102 | PZ17_Chen_4 | 169.44 | 0 | 52 | 169 | 0 | 0 | 11 | 23 | 255 |
| 103 | PZ17_SimRNA2_8 | 154.46 | 0 | 69 | 160 | 0 | 4 | 3 | 22 | 258 |

| | | | | | | | | | | |
|---|---|---|---|---|---|---|---|---|---|---|
| 104 | PZ17_SimRNA2_6 | 122.29 | 0 | 64 | 166 | 1 | 5 | 4 | 20 | 260 |
| 105 | PZ17_SimRNA2_3 | 126.88 | 0 | 75 | 152 | 0 | 9 | 2 | 23 | 261 |
| 106 | PZ17_SimRNA2_5 | 128.01 | 0 | 67 | 162 | 1 | 3 | 7 | 23 | 263 |
| 107 | PZ17_Xiao_9 | 14.07 | 53 | 15 | 137 | 39 | 1 | 30 | 45 | 267 |
| 108 | PZ17_SimRNA1_4 | 162.23 | 0 | 65 | 175 | 0 | 6 | 3 | 21 | 270 |
| | Average | 38.08 | 58.36 | 15.72 | 67.53 | 3.64 | 0.86 | 6.02 | 20.94 | 114.71 |
| | Standard deviation | 56.18 | 37.43 | 24.78 | 61.21 | 9.94 | 1.67 | 7.29 | 9.77 | 91.03 |
| | Median | 10.55 | 68.00 | 1.00 | 73.50 | 0.00 | 0.00 | 3.00 | 18.00 | 117.50 |

**Table S88**. MolProbity report on structures in Puzzle 18.

| No. | 3D RNA model | (a) Clash score | (b) Rank-ing | #Bad bonds | #Bad angles | #Chiral handedness swaps | #Tetrahedral geometry outli-ers | #Probably wrong sugar puckers: | (c) #Bad back-bone confor-mations | Total |
|---|---|---|---|---|---|---|---|---|---|---|
| 1 | PZ18_refstructure | 3.48 | 97 | 0 | 0 | 0 | 0 | 0 | 7 | 7 |
| 2 | PZ18_FARFAR_2 | 5.22 | 93 | 0 | 0 | 0 | 0 | 0 | 8 | 8 |
| 3 | PZ18_FARFAR_10 | 6.52 | 89 | 0 | 0 | 0 | 0 | 1 | 8 | 9 |
| 4 | PZ18_Lee_4 | 3.91 | 96 | 1 | 4 | 0 | 0 | 1 | 3 | 9 |
| 5 | PZ18_FARFAR_6 | 3.91 | 96 | 0 | 0 | 0 | 0 | 0 | 11 | 11 |
| 6 | PZ18_Lee_1 | 1.74 | 99 | 1 | 3 | 0 | 0 | 2 | 6 | 12 |
| 7 | PZ18_Lee_5 | 2.17 | 99 | 1 | 5 | 0 | 0 | 1 | 5 | 12 |
| 8 | PZ18_FARFAR_3 | 10.44 | 69 | 1 | 2 | 0 | 0 | 1 | 9 | 13 |
| 9 | PZ18_FARFAR_4 | 3.48 | 97 | 0 | 0 | 0 | 0 | 0 | 13 | 13 |
| 10 | PZ18_Lee_2 | 3.91 | 96 | 1 | 4 | 0 | 0 | 2 | 6 | 13 |
| 11 | PZ18_FARFAR_7 | 7.39 | 85 | 0 | 2 | 0 | 0 | 1 | 11 | 14 |
| 12 | PZ18_FARFAR_8 | 3.48 | 97 | 0 | 1 | 0 | 0 | 1 | 12 | 14 |
| 13 | PZ18_LeeServer_1 | 0.87 | 99 | 1 | 5 | 0 | 0 | 2 | 6 | 14 |
| 14 | PZ18_RNAComposer_1 | 12.61 | 60 | 0 | 0 | 0 | 0 | 0 | 14 | 14 |
| 15 | PZ18_FARFAR_1 | 6.52 | 89 | 0 | 1 | 0 | 0 | 1 | 13 | 15 |
| 16 | PZ18_FARFAR_9 | 1.74 | 99 | 1 | 1 | 0 | 0 | 1 | 15 | 18 |
| 17 | PZ18_RNAComposer_2 | 10.87 | 67 | 0 | 0 | 0 | 0 | 1 | 17 | 18 |
| 18 | PZ18_RNAComposer_3 | 10.87 | 67 | 0 | 0 | 0 | 0 | 2 | 16 | 18 |
| 19 | PZ18_RNAComposer_4 | 10.87 | 67 | 0 | 0 | 0 | 0 | 2 | 16 | 18 |

| | | | | | | | | | | |
|---|---|---|---|---|---|---|---|---|---|---|
| 20 | PZ18_Das_2 | 16.09 | 45 | 1 | 5 | 0 | 0 | 0 | 13 | 19 |
| 21 | PZ18_LeeServer_5 | 2.17 | 99 | 1 | 6 | 0 | 0 | 2 | 10 | 19 |
| 22 | PZ18_Lee_3 | 1.74 | 99 | 1 | 7 | 0 | 0 | 1 | 11 | 20 |
| 23 | PZ18_LeeServer_4 | 4.78 | 94 | 1 | 7 | 0 | 0 | 4 | 8 | 20 |
| 24 | PZ18_RNAComposer_5 | 13.92 | 54 | 0 | 0 | 0 | 0 | 4 | 16 | 20 |
| 25 | PZ18_Das_3 | 9.57 | 74 | 0 | 6 | 0 | 0 | 0 | 15 | 21 |
| 26 | PZ18_Das_4 | 13.05 | 58 | 0 | 6 | 0 | 0 | 1 | 14 | 21 |
| 27 | PZ18_FARFAR_5 | 8.26 | 81 | 0 | 5 | 0 | 0 | 2 | 14 | 21 |
| 28 | PZ18_Das_1 | 13.05 | 58 | 1 | 9 | 0 | 0 | 0 | 15 | 25 |
| 29 | PZ18_LeeServer_2 | 4.35 | 95 | 1 | 11 | 0 | 0 | 5 | 15 | 32 |
| 30 | PZ18_LeeServer_3 | 3.48 | 97 | 1 | 8 | 0 | 1 | 5 | 17 | 32 |
| 31 | PZ18_Das_5 | 19.57 | 34 | 1 | 10 | 0 | 0 | 2 | 21 | 34 |
| 32 | PZ18_YagoubAli_1 | 8.26 | 81 | 2 | 4 | 0 | 0 | 4 | 26 | 36 |
| 33 | PZ18_3dRNA_2 | 0.43 | 99 | 1 | 42 | 3 | 0 | 5 | 12 | 63 |
| 34 | PZ18_Chen_5 | 1.31 | 99 | 7 | 34 | 0 | 2 | 3 | 17 | 63 |
| 35 | PZ18_Chen_1 | 0.43 | 99 | 5 | 40 | 0 | 0 | 7 | 32 | 84 |
| 36 | PZ18_Chen_4 | 4.78 | 94 | 9 | 45 | 0 | 1 | 3 | 28 | 86 |
| 37 | PZ18_Chen_3 | 4.78 | 94 | 9 | 56 | 0 | 1 | 5 | 31 | 102 |
| 38 | PZ18_Chen_2 | 2.17 | 99 | 9 | 51 | 0 | 0 | 11 | 35 | 106 |
| 39 | PZ18_3dRNA_4 | 0.87 | 99 | 1 | 57 | 19 | 1 | 10 | 22 | 110 |
| 40 | PZ18_Ding_2 | 10.87 | 67 | 3 | 94 | 0 | 0 | 1 | 15 | 113 |
| 41 | PZ18_Ding_4 | 8.26 | 81 | 4 | 97 | 0 | 1 | 2 | 11 | 115 |
| 42 | PZ18_Ding_1 | 10.44 | 69 | 6 | 97 | 0 | 0 | 1 | 14 | 118 |
| 43 | PZ18_Ding_5 | 9.13 | 76 | 2 | 98 | 0 | 0 | 5 | 17 | 122 |
| 44 | PZ18_Dokholyan_2 | 14.79 | 50 | 1 | 93 | 0 | 0 | 5 | 24 | 123 |
| 45 | PZ18_3dRNA_5 | 0.43 | 99 | 1 | 66 | 16 | 0 | 11 | 31 | 125 |
| 46 | PZ18_3dRNA_3 | 12.61 | 60 | 10 | 83 | 9 | 0 | 9 | 22 | 133 |
| 47 | PZ18_Dokholyan_3 | 13.48 | 56 | 0 | 94 | 0 | 0 | 8 | 31 | 133 |
| 48 | PZ18_Ding_3 | 7.83 | 82 | 5 | 108 | 0 | 0 | 5 | 20 | 138 |
| 49 | PZ18_Dokholyan_1 | 9.57 | 74 | 0 | 108 | 0 | 0 | 5 | 29 | 142 |
| 50 | PZ18_3dRNA_1 | 11.74 | 64 | 9 | 84 | 13 | 0 | 10 | 31 | 147 |
| 51 | PZ18_SimRNA_1 | 100.48 | 0 | 62 | 131 | 1 | 1 | 3 | 17 | 215 |
| 52 | PZ18_SimRNA_3 | 100.78 | 0 | 74 | 133 | 0 | 2 | 4 | 16 | 229 |

| | | | | | | | | | | |
|---|---|---|---|---|---|---|---|---|---|---|
| 53 | PZ18_SimRNA_2 | 87.75 | 0 | 67 | 162 | 1 | 3 | 1 | 13 | 247 |
| | Average | 12.10 | 77.19 | 5.70 | 35.57 | 1.17 | 0.25 | 3.08 | 16.21 | 61.96 |
| | Standard deviation | 21.41 | 25.85 | 15.63 | 45.30 | 3.94 | 0.62 | 3.02 | 7.90 | 62.93 |
| | Median | 7.39 | 85.00 | 1.00 | 7.00 | 0.00 | 0.00 | 2.00 | 15.00 | 21.00 |

**Table S89**. MolProbity report on structures in Puzzle 19.

| No. | 3D RNA model | (a) Clash score | (b) Rank-ing | #Bad bonds | #Bad angles | #Chiral handedness swaps | #Tetrahedral geometry out-liers | #Probably wrong sugar puckers: | (c) #Bad back-bone confor-mations | Total |
|---|---|---|---|---|---|---|---|---|---|---|
| 1 | PZ19_Adamiak_4 | 6.99 | 87 | 0 | 0 | 0 | 0 | 0 | 4 | 4 |
| 2 | PZ19_Adamiak_2 | 6.99 | 87 | 0 | 0 | 0 | 0 | 0 | 6 | 6 |
| 3 | PZ19_Adamiak_3 | 7.49 | 84 | 0 | 0 | 0 | 0 | 2 | 4 | 6 |
| 4 | PZ19_RNAComposer_1 | 6.49 | 89 | 0 | 0 | 0 | 0 | 0 | 8 | 8 |
| 5 | PZ19_Adamiak_1 | 7.99 | 82 | 0 | 0 | 0 | 0 | 2 | 8 | 10 |
| 6 | PZ19_FARFAR_9 | 5.99 | 90 | 0 | 1 | 0 | 0 | 2 | 8 | 11 |
| 7 | PZ19_FARFAR_7 | 6.99 | 87 | 0 | 3 | 0 | 0 | 1 | 8 | 12 |
| 8 | PZ19_RNAComposer_3 | 11.48 | 65 | 0 | 0 | 0 | 0 | 1 | 11 | 12 |
| 9 | PZ19_LeeServer_5 | 4.49 | 95 | 2 | 4 | 0 | 0 | 0 | 7 | 13 |
| 10 | PZ19_refstructure | 2.50 | 99 | 1 | 6 | 0 | 0 | 1 | 6 | 14 |
| 11 | PZ19_Adamiak_5 | 5.49 | 92 | 0 | 0 | 0 | 0 | 1 | 13 | 14 |
| 12 | PZ19_RNAComposer_2 | 9.99 | 71 | 0 | 0 | 0 | 0 | 2 | 12 | 14 |
| 13 | PZ19_FARFAR_1 | 4.00 | 96 | 0 | 2 | 0 | 0 | 1 | 12 | 15 |
| 14 | PZ19_FARFAR_8 | 5.00 | 94 | 1 | 2 | 0 | 0 | 2 | 10 | 15 |
| 15 | PZ19_RNAComposer_5 | 11.48 | 65 | 0 | 0 | 0 | 0 | 4 | 11 | 15 |
| 16 | PZ19_FARFAR_2 | 7.49 | 84 | 0 | 2 | 0 | 0 | 3 | 11 | 16 |
| 17 | PZ19_RNAComposer_4 | 12.98 | 58 | 0 | 0 | 0 | 0 | 2 | 14 | 16 |
| 18 | PZ19_FARFAR_3 | 4.50 | 95 | 0 | 4 | 0 | 0 | 1 | 12 | 17 |
| 19 | PZ19_FARFAR_6 | 5.00 | 94 | 1 | 3 | 0 | 0 | 3 | 10 | 17 |
| 20 | PZ19_FARFAR_10 | 7.49 | 84 | 1 | 0 | 0 | 0 | 2 | 15 | 18 |
| 21 | PZ19_FARFAR_5 | 7.99 | 82 | 0 | 3 | 0 | 0 | 2 | 13 | 18 |
| 22 | PZ19_LeeServer_4 | 6.49 | 89 | 2 | 7 | 0 | 1 | 2 | 10 | 22 |
| 23 | PZ19_FARFAR_4 | 10.99 | 67 | 0 | 10 | 0 | 0 | 3 | 11 | 24 |

| | | | | | | | | | | |
|---|---|---|---|---|---|---|---|---|---|---|
| 24 | PZ19_Bujnicki_1 | 1.50 | 99 | 0 | 17 | 2 | 0 | 2 | 11 | 32 |
| 25 | PZ19_Chen_4 | 0.00 | 100 | 5 | 16 | 0 | 0 | 3 | 21 | 45 |
| 26 | PZ19_Chen_3 | 0.00 | 100 | 1 | 30 | 0 | 0 | 4 | 22 | 57 |
| 27 | PZ19_Bujnicki_2 | 4.49 | 95 | 0 | 42 | 5 | 0 | 6 | 13 | 66 |
| 28 | PZ19_Dokholyan_1 | 6.49 | 89 | 0 | 50 | 0 | 0 | 3 | 15 | 68 |
| 29 | PZ19_Bujnicki_5 | 1.00 | 99 | 0 | 40 | 11 | 1 | 10 | 18 | 80 |
| 30 | PZ19_Bujnicki_3 | 3.99 | 96 | 0 | 63 | 4 | 0 | 5 | 11 | 83 |
| 31 | PZ19_Chen_2 | 0.50 | 99 | 8 | 43 | 0 | 0 | 4 | 29 | 84 |
| 32 | PZ19_LeeServer_3 | 1.50 | 99 | 2 | 12 | 49 | 0 | 11 | 17 | 91 |
| 33 | PZ19_LeeServer_2 | 1.00 | 99 | 2 | 7 | 51 | 0 | 12 | 20 | 92 |
| 34 | PZ19_LeeServer_1 | 1.50 | 99 | 2 | 12 | 50 | 0 | 11 | 18 | 93 |
| 35 | PZ19_Ding_2 | 10.98 | 67 | 1 | 77 | 0 | 0 | 3 | 14 | 95 |
| 36 | PZ19_Ding_4 | 12.98 | 58 | 2 | 93 | 0 | 0 | 2 | 16 | 113 |
| 37 | PZ19_Ding_1 | 8.49 | 79 | 5 | 97 | 0 | 0 | 2 | 10 | 114 |
| 38 | PZ19_Ding_5 | 8.49 | 79 | 3 | 95 | 0 | 1 | 2 | 13 | 114 |
| 39 | PZ19_Ding_3 | 7.99 | 82 | 5 | 89 | 0 | 0 | 3 | 18 | 115 |
| 40 | PZ19_Bujnicki_4 | 4.99 | 94 | 1 | 114 | 9 | 2 | 8 | 18 | 152 |
| 41 | PZ19_Chen_1 | 18.97 | 35 | 22 | 92 | 1 | 1 | 10 | 26 | 152 |
| 42 | PZ19_3dRNA_5 | 5.50 | 92 | 15 | 102 | 11 | 1 | 8 | 30 | 167 |
| 43 | PZ19_3dRNA_2 | 4.00 | 96 | 15 | 94 | 11 | 1 | 14 | 34 | 169 |
| 44 | PZ19_3dRNA_4 | 4.00 | 96 | 16 | 101 | 15 | 1 | 11 | 33 | 177 |
| 45 | PZ19_3dRNA_1 | 4.50 | 95 | 15 | 103 | 12 | 1 | 15 | 33 | 179 |
| 46 | PZ19_3dRNA_3 | 6.00 | 90 | 7 | 102 | 15 | 2 | 14 | 40 | 180 |
| 47 | PZ19_SimRNA_1 | 108.39 | 0 | 52 | 119 | 1 | 1 | 1 | 9 | 183 |
| 48 | PZ19_Chen_5 | 19.97 | 32 | 41 | 125 | 3 | 5 | 8 | 35 | 217 |
| 49 | PZ19_SimRNA_2 | 134.40 | 0 | 55 | 158 | 0 | 3 | 3 | 17 | 236 |
| 50 | PZ19_SimRNA_3 | 105.68 | 0 | 71 | 148 | 2 | 2 | 5 | 19 | 247 |
| 51 | PZ19_Das_1 | Rejected as unacceptable - Bond distance > max_reasonable_bond_distance: 146.452 > 50: distance: 1298 - 1299: 144.568 | | | | | | | | |
| 52 | PZ19_Das_2 | Rejected as unacceptable - Bond distance > max_reasonable_bond_distance: 104.524 > 50: distance: 1298 - 1299: 103.861 | | | | | | | | |
| 53 | PZ19_Das_3 | Rejected as unacceptable - Bond distance > max_reasonable_bond_distance: 114.045 > 50: distance: 1298 - 1299: 112.953 | | | | | | | | |
| 54 | PZ19_Das_4 | Rejected as unacceptable - Bond distance > max_reasonable_bond_distance: 145.887 > 50: distance: 1298 - 1299: 144.334 | | | | | | | | |
| 55 | PZ19_Das_5 | Rejected as unacceptable - Bond distance > max_reasonable_bond_distance: 76.1157 > 50: distance: 1298 - 1299: 75.621 | | | | | | | | |
| | Average | 13.07 | 80.10 | 7.08 | 41.76 | 5.04 | 0.46 | 4.34 | 15.68 | 74.36 |

| | | | | | | | | | | |
|---|---|---|---|---|---|---|---|---|---|---|
| | Standard deviation | 26.83 | 25.61 | 15.38 | 48.66 | 12.23 | 0.95 | 4.14 | 8.65 | 70.88 |
| | Median | 6.49 | 89.00 | 1.00 | 12.00 | 0.00 | 0.00 | 3.00 | 13.00 | 51.00 |

**Table S90**. MolProbity report on structures in Puzzle 20.

| No. | 3D RNA model | (a) Clash score | (b) Ranking | #Bad bonds | #Bad angles | #Chiral handedness swaps | #Tetrahedral geometry outliers | #Probably wrong sugar puckers: | (c) #Bad backbone conformations | Total |
|---|---|---|---|---|---|---|---|---|---|---|
| 1 | PZ20_FARFAR_2 | 3.19 | 97 | 0 | 0 | 0 | 0 | 0 | 4 | 4 |
| 2 | PZ20_refstructure | 0.46 | 99 | 0 | 0 | 0 | 0 | 0 | 5 | 5 |
| 3 | PZ20_FARFAR_9 | 2.28 | 99 | 0 | 0 | 0 | 0 | 0 | 6 | 6 |
| 4 | PZ20_RNAComposer_3 | 10.48 | 69 | 0 | 0 | 0 | 0 | 0 | 8 | 8 |
| 5 | PZ20_RNAComposer_5 | 9.12 | 76 | 0 | 0 | 0 | 0 | 1 | 10 | 11 |
| 6 | PZ20_Adamiak_2 | 9.12 | 76 | 0 | 0 | 0 | 0 | 1 | 11 | 12 |
| 7 | PZ20_FARFAR_1 | 6.84 | 87 | 0 | 0 | 0 | 0 | 0 | 12 | 12 |
| 8 | PZ20_FARFAR_5 | 5.47 | 92 | 0 | 1 | 0 | 0 | 2 | 9 | 12 |
| 9 | PZ20_FARFAR_3 | 9.57 | 74 | 0 | 3 | 0 | 0 | 1 | 9 | 13 |
| 10 | PZ20_FARFAR_4 | 3.65 | 97 | 0 | 4 | 0 | 0 | 0 | 9 | 13 |
| 11 | PZ20_RNAComposer_1 | 6.84 | 87 | 0 | 0 | 0 | 0 | 1 | 12 | 13 |
| 12 | PZ20_RNAComposer_2 | 5.47 | 92 | 0 | 0 | 0 | 0 | 0 | 13 | 13 |
| 13 | PZ20_FARFAR_10 | 6.84 | 87 | 0 | 3 | 0 | 0 | 3 | 8 | 14 |
| 14 | PZ20_Adamiak_1 | 8.20 | 81 | 0 | 0 | 0 | 0 | 2 | 13 | 15 |
| 15 | PZ20_Adamiak_3 | 10.48 | 69 | 1 | 0 | 0 | 0 | 2 | 12 | 15 |
| 16 | PZ20_Adamiak_4 | 7.29 | 85 | 0 | 0 | 0 | 0 | 1 | 15 | 16 |
| 17 | PZ20_Das_3 | 9.57 | 74 | 3 | 4 | 0 | 0 | 2 | 7 | 16 |
| 18 | PZ20_FARFAR_6 | 6.84 | 87 | 0 | 3 | 0 | 0 | 2 | 11 | 16 |
| 19 | PZ20_FARFAR_7 | 8.20 | 81 | 0 | 3 | 0 | 1 | 3 | 9 | 16 |
| 20 | PZ20_FARFAR_8 | 6.38 | 89 | 0 | 3 | 0 | 0 | 1 | 13 | 17 |
| 21 | PZ20_RNAComposer_4 | 5.47 | 92 | 0 | 0 | 0 | 0 | 1 | 16 | 17 |
| 22 | PZ20_Adamiak_5 | 6.84 | 87 | 0 | 0 | 0 | 0 | 2 | 16 | 18 |
| 23 | PZ20_Das_2 | 15.50 | 48 | 1 | 5 | 0 | 0 | 0 | 12 | 18 |
| 24 | PZ20_Das_1 | 15.50 | 48 | 1 | 5 | 0 | 0 | 1 | 15 | 22 |
| 25 | PZ20_Das_4 | 18.23 | 37 | 1 | 3 | 0 | 0 | 1 | 19 | 24 |

| | | | | | | | | | | |
|---|---|---|---|---|---|---|---|---|---|---|
| 26 | PZ20_Das_5 | 28.70 | 17 | 1 | 5 | 0 | 0 | 3 | 16 | 25 |
| 27 | PZ20_Bujnicki_5 | 0.46 | 99 | 0 | 13 | 3 | 0 | 5 | 11 | 32 |
| 28 | PZ20_Bujnicki_1 | 21.41 | 29 | 23 | 105 | 1 | 0 | 4 | 10 | 143 |
| 29 | PZ20_Xiao3_5 | 1.82 | 99 | 2 | 88 | 34 | 1 | 16 | 34 | 175 |
| 30 | PZ20_SimRNA_2 | 144.49 | 0 | 63 | 139 | 1 | 1 | 4 | 13 | 221 |
| 31 | PZ20_Xiao3_3 | 39.32 | 8 | 27 | 130 | 27 | 0 | 18 | 34 | 236 |
| 32 | PZ20_SimRNA_5 | 98.63 | 0 | 64 | 149 | 0 | 3 | 2 | 20 | 238 |
| 33 | PZ20_Xiao3_2 | 13.70 | 55 | 8 | 133 | 37 | 0 | 19 | 41 | 238 |
| 34 | PZ20_Bujnicki_3 | 27.37 | 18 | 55 | 158 | 1 | 5 | 6 | 14 | 239 |
| 35 | PZ20_SimRNA_4 | 102.91 | 0 | 79 | 145 | 0 | 3 | 3 | 18 | 248 |
| 36 | PZ20_Bujnicki_4 | 37.37 | 9 | 41 | 180 | 1 | 6 | 7 | 19 | 254 |
| 37 | PZ20_SimRNA_1 | 129.15 | 0 | 77 | 154 | 0 | 2 | 7 | 19 | 259 |
| 38 | PZ20_Bujnicki_2 | 32.80 | 13 | 40 | 195 | 1 | 4 | 7 | 14 | 261 |
| 39 | PZ20_SimRNA_3 | 101.32 | 0 | 69 | 169 | 2 | 2 | 4 | 23 | 269 |
| 40 | PZ20_Xiao3_4 | 13.68 | 55 | 9 | 137 | 50 | 0 | 27 | 48 | 271 |
| 41 | PZ20_Xiao3_1 | 25.57 | 20 | 21 | 127 | 53 | 2 | 28 | 51 | 282 |
| | Average | 24.79 | 59.32 | 14.29 | 50.34 | 5.15 | 0.73 | 4.56 | 16.07 | 91.15 |
| | Standard deviation | 36.05 | 35.88 | 24.89 | 69.80 | 13.69 | 1.48 | 6.95 | 10.78 | 109.42 |
| | Median | 9.57 | 74.00 | 0.00 | 3.00 | 0.00 | 0.00 | 2.00 | 13.00 | 17.00 |

**Table S91**. MolProbity report on structures in Puzzle 21.

| **No.** | **3D RNA model** | **(a) Clash score** | **(b) Rank-ing** | **#Bad bonds** | **#Bad angles** | **#Chiral handedness swaps** | **#Tetrahedral geometry outli-ers** | **#Probably wrong sugar puck-ers:** | **(c) #Bad back-bone confor-mations** | **Total** |
|---|---|---|---|---|---|---|---|---|---|---|
| 1 | PZ21_Das_1 | 17.29 | 41 | 0 | 0 | 0 | 0 | 0 | 3 | 3 |
| 2 | PZ21_RNAComposer_2 | 9.02 | 77 | 0 | 0 | 0 | 0 | 0 | 3 | 3 |
| 3 | PZ21_Das_2 | 19.55 | 34 | 0 | 0 | 0 | 0 | 0 | 4 | 4 |
| 4 | PZ21_Das_4 | 11.28 | 66 | 0 | 3 | 0 | 0 | 0 | 1 | 4 |
| 5 | PZ21_Das_3 | 15.04 | 49 | 0 | 3 | 0 | 0 | 0 | 3 | 6 |
| 6 | PZ21_RNAComposer_1 | 15.04 | 49 | 0 | 0 | 0 | 0 | 1 | 5 | 6 |
| 7 | PZ21_RNAComposer_5 | 6.77 | 88 | 0 | 0 | 0 | 0 | 0 | 7 | 7 |
| 8 | PZ21_Adamiak_3 | 8.27 | 81 | 0 | 0 | 0 | 0 | 1 | 7 | 8 |

| | | | | | | | | | | |
|---|---|---|---|---|---|---|---|---|---|---|
| 9 | PZ21_DasLORES_4 | 3.01 | 98 | 1 | 0 | 0 | 0 | 0 | 7 | 8 |
| 10 | PZ21_Das_5 | 14.29 | 52 | 0 | 5 | 0 | 0 | 1 | 3 | 9 |
| 11 | PZ21_Adamiak_4 | 13.53 | 56 | 0 | 0 | 0 | 0 | 1 | 9 | 10 |
| 12 | PZ21_FARFAR_8 | 6.02 | 90 | 0 | 1 | 0 | 0 | 2 | 7 | 10 |
| 13 | PZ21_Adamiak_2 | 10.53 | 68 | 0 | 0 | 0 | 0 | 0 | 11 | 11 |
| 14 | PZ21_Adamiak_5 | 13.53 | 56 | 0 | 0 | 0 | 0 | 0 | 11 | 11 |
| 15 | PZ21_DasLORES_3 | 6.77 | 88 | 1 | 2 | 0 | 0 | 0 | 8 | 11 |
| 16 | PZ21_RNAComposer_4 | 9.02 | 77 | 0 | 0 | 0 | 0 | 0 | 11 | 11 |
| 17 | PZ21_Adamiak_1 | 12.78 | 60 | 0 | 0 | 0 | 0 | 0 | 12 | 12 |
| 18 | PZ21_DasLORES_2 | 6.77 | 88 | 0 | 1 | 0 | 0 | 1 | 10 | 12 |
| 19 | PZ21_FARFAR_3 | 8.27 | 81 | 0 | 1 | 0 | 0 | 2 | 10 | 13 |
| 20 | PZ21_FARFAR_6 | 4.51 | 95 | 0 | 1 | 0 | 0 | 2 | 10 | 13 |
| 21 | PZ21_DasLORES_5 | 5.26 | 93 | 0 | 0 | 0 | 0 | 0 | 14 | 14 |
| 22 | PZ21_FARFAR_7 | 10.53 | 68 | 0 | 2 | 0 | 0 | 2 | 10 | 14 |
| 23 | PZ21_FARFAR_9 | 2.26 | 99 | 0 | 4 | 0 | 0 | 0 | 10 | 14 |
| 24 | PZ21_RNAComposer_3 | 18.80 | 36 | 0 | 0 | 0 | 0 | 0 | 14 | 14 |
| 25 | PZ21_FARFAR_5 | 6.02 | 90 | 0 | 3 | 0 | 0 | 3 | 9 | 15 |
| 26 | PZ21_FARFAR_10 | 9.77 | 73 | 0 | 1 | 0 | 0 | 3 | 12 | 16 |
| 27 | PZ21_FARFAR_4 | 7.52 | 84 | 0 | 4 | 0 | 0 | 2 | 10 | 16 |
| 28 | PZ21_Bujnicki_4 | 0.00 | 100 | 0 | 3 | 4 | 0 | 3 | 8 | 18 |
| 29 | PZ21_FARFAR_2 | 9.77 | 73 | 1 | 4 | 0 | 0 | 3 | 10 | 18 |
| 30 | PZ21_DasLORES_1 | 9.77 | 73 | 0 | 4 | 0 | 0 | 2 | 13 | 19 |
| 31 | PZ21_FARFAR_1 | 9.02 | 77 | 0 | 3 | 0 | 0 | 3 | 13 | 19 |
| 32 | PZ21_Bujnicki_1 | 0.00 | 100 | 0 | 7 | 4 | 0 | 4 | 10 | 25 |
| 33 | PZ21_refstructure | 20.29 | 31 | 10 | 2 | 0 | 0 | 0 | 16 | 28 |
| 34 | PZ21_Bujnicki_3 | 0.00 | 100 | 0 | 6 | 6 | 0 | 6 | 13 | 31 |
| 35 | PZ21_Bujnicki_5 | 0.75 | 99 | 0 | 16 | 4 | 0 | 4 | 11 | 35 |
| 36 | PZ21_Sanbonmatsu_2 | 109.19 | 0 | 14 | 33 | 12 | 14 | 6 | 16 | 95 |
| 37 | PZ21_Sanbonmatsu_4 | 122.93 | 0 | 14 | 39 | 8 | 14 | 5 | 16 | 96 |
| 38 | PZ21_Sanbonmatsu_1 | 109.19 | 0 | 14 | 35 | 12 | 14 | 6 | 16 | 97 |
| 39 | PZ21_Sanbonmatsu_3 | 122.93 | 0 | 14 | 42 | 8 | 14 | 5 | 16 | 99 |
| 40 | PZ21_Bujnicki_2 | 8.00 | 82 | 29 | 61 | 0 | 2 | 2 | 6 | 100 |
| 41 | PZ21_3dRNA_5 | 0.75 | 99 | 0 | 54 | 22 | 0 | 10 | 20 | 106 |

| | | | | | | | | | | |
|---|---|---|---|---|---|---|---|---|---|---|
| 42 | PZ21_3dRNA_4 | 0.75 | 99 | 1 | 54 | 26 | 0 | 15 | 31 | 127 |
| 43 | PZ21_ChenHighLig_3 | 107.92 | 0 | 32 | 88 | 0 | 0 | 1 | 6 | 127 |
| 44 | PZ21_ChenLowLig_3 | 107.92 | 0 | 32 | 88 | 0 | 0 | 1 | 6 | 127 |
| 45 | PZ21_3dRNA_3 | 0.00 | 100 | 0 | 66 | 21 | 1 | 14 | 32 | 134 |
| 46 | PZ21_SimRNA_4 | 117.03 | 0 | 38 | 87 | 0 | 2 | 0 | 10 | 137 |
| 47 | PZ21_SimRNA_2 | 116.54 | 0 | 35 | 90 | 0 | 3 | 3 | 9 | 140 |
| 48 | PZ21_SimRNA_1 | 114.95 | 0 | 40 | 101 | 1 | 0 | 1 | 11 | 154 |
| 49 | PZ21_ChenLowLig_1 | 128.21 | 0 | 33 | 116 | 0 | 0 | 3 | 8 | 160 |
| 50 | PZ21_ChenHighLig_1 | 128.21 | 0 | 33 | 117 | 0 | 0 | 3 | 8 | 161 |
| 51 | PZ21_ChenHighLig_2 | 73.04 | 1 | 29 | 120 | 0 | 1 | 2 | 9 | 161 |
| 52 | PZ21_ChenLowLig_2 | 73.04 | 1 | 29 | 120 | 0 | 1 | 2 | 9 | 161 |
| 53 | PZ21_3dRNA_1 | 37.65 | 9 | 19 | 79 | 16 | 0 | 18 | 30 | 162 |
| 54 | PZ21_ChenHighLig_5 | 124.53 | 0 | 29 | 117 | 0 | 0 | 7 | 15 | 168 |
| 55 | PZ21_ChenLowLig_5 | 123.77 | 0 | 29 | 118 | 0 | 0 | 7 | 15 | 169 |
| 56 | PZ21_SimRNA_3 | 114.54 | 0 | 47 | 104 | 1 | 3 | 2 | 12 | 169 |
| 57 | PZ21_SimRNA_5 | 114.54 | 0 | 47 | 104 | 1 | 3 | 2 | 12 | 169 |
| 58 | PZ21_3dRNA_2 | 46.62 | 5 | 21 | 103 | 24 | 0 | 19 | 32 | 199 |
| 59 | PZ21_ChenLowLig_4 | 223.48 | 0 | 41 | 128 | 0 | 2 | 12 | 20 | 203 |
| 60 | PZ21_ChenHighLig_4 | 222.73 | 0 | 41 | 129 | 0 | 2 | 12 | 20 | 204 |
| | Average | 46.16 | 49.77 | 11.23 | 37.82 | 2.83 | 1.27 | 3.40 | 11.67 | 68.22 |
| | Standard deviation | 57.90 | 39.95 | 15.73 | 47.05 | 6.45 | 3.53 | 4.53 | 6.76 | 69.14 |
| | Median | 13.16 | 58.00 | 0.00 | 4.00 | 0.00 | 0.00 | 2.00 | 10.00 | 19.00 |

**Table S92**. MolProbity report on structures in Puzzle 24.

| No. | 3D RNA model | (a) Clash score | (b) Ranking | #Bad bonds | #Bad angles | #Chiral handedness swaps | #Tetrahedral geometry outliers | #Probably wrong sugar puckers: | (c) #Bad backbone conformations | Total |
|---|---|---|---|---|---|---|---|---|---|---|
| 1 | PZ24_FARFAR2_2 | 0.55 | 99 | 0 | 2 | 0 | 0 | 0 | 3 | 5 |
| 2 | PZ24_FARFAR2_5 | 0.83 | 99 | 0 | 1 | 0 | 0 | 0 | 4 | 5 |
| 3 | PZ24_FARFAR2_4 | 0.83 | 99 | 0 | 1 | 0 | 0 | 0 | 5 | 6 |
| 4 | PZ24_FARFAR2_8 | 0.83 | 99 | 0 | 3 | 0 | 0 | 0 | 4 | 7 |
| 5 | PZ24_FARFAR2_3 | 0.83 | 99 | 0 | 3 | 0 | 0 | 0 | 5 | 8 |

| | | | | | | | | | | |
|---|---|---|---|---|---|---|---|---|---|---|
| 6 | PZ24_Das_2 | 1.94 | 99 | 0 | 0 | 0 | 0 | 0 | 9 | 9 |
| 7 | PZ24_Das_3 | 1.94 | 99 | 0 | 0 | 0 | 0 | 0 | 9 | 9 |
| 8 | PZ24_Das_4 | 2.22 | 99 | 0 | 0 | 0 | 0 | 0 | 9 | 9 |
| 9 | PZ24_Das_8 | 1.94 | 99 | 0 | 0 | 0 | 0 | 0 | 9 | 9 |
| 10 | PZ24_Das_1 | 1.94 | 99 | 0 | 0 | 0 | 0 | 0 | 10 | 10 |
| 11 | PZ24_Das_5 | 1.94 | 99 | 0 | 0 | 0 | 0 | 0 | 10 | 10 |
| 12 | PZ24_refstructure | 0.28 | 99 | 0 | 0 | 0 | 0 | 1 | 10 | 11 |
| 13 | PZ24_Das_6 | 2.22 | 99 | 0 | 0 | 0 | 0 | 0 | 12 | 12 |
| 14 | PZ24_RNAComposer_1 | 8.87 | 78 | 0 | 0 | 0 | 0 | 0 | 12 | 12 |
| 15 | PZ24_Das_9 | 1.94 | 99 | 0 | 0 | 0 | 0 | 0 | 13 | 13 |
| 16 | PZ24_Das_10 | 1.94 | 99 | 0 | 1 | 0 | 0 | 0 | 13 | 14 |
| 17 | PZ24_Das_7 | 1.94 | 99 | 0 | 1 | 0 | 0 | 0 | 13 | 14 |
| 18 | PZ24_DasTFN_3 | 1.66 | 99 | 0 | 2 | 0 | 0 | 0 | 12 | 14 |
| 19 | PZ24_FARFAR2_1 | 1.66 | 99 | 0 | 2 | 0 | 0 | 0 | 12 | 14 |
| 20 | PZ24_FARFAR2_7 | 1.66 | 99 | 0 | 0 | 0 | 0 | 0 | 15 | 15 |
| 21 | PZ24_DasTFN_9 | 2.22 | 99 | 1 | 3 | 0 | 0 | 0 | 14 | 18 |
| 22 | PZ24_FARFAR2_6 | 2.22 | 99 | 1 | 3 | 0 | 0 | 0 | 14 | 18 |
| 23 | PZ24_RNAComposer_3 | 6.10 | 90 | 0 | 0 | 0 | 0 | 0 | 19 | 19 |
| 24 | PZ24_RNAComposer_5 | 5.82 | 91 | 0 | 0 | 0 | 0 | 0 | 19 | 19 |
| 25 | PZ24_Adamiak_5 | 11.92 | 63 | 0 | 0 | 0 | 0 | 1 | 19 | 20 |
| 26 | PZ24_RNAComposer_4 | 8.32 | 80 | 0 | 0 | 0 | 0 | 0 | 21 | 21 |
| 27 | PZ24_FARFAR2_10 | 3.05 | 98 | 0 | 3 | 0 | 0 | 3 | 16 | 22 |
| 28 | PZ24_FARFAR2_9 | 1.66 | 99 | 0 | 1 | 0 | 0 | 0 | 21 | 22 |
| 29 | PZ24_Adamiak_1 | 10.82 | 67 | 0 | 0 | 0 | 0 | 2 | 21 | 23 |
| 30 | PZ24_DasTFN_2 | 4.99 | 94 | 0 | 4 | 0 | 0 | 2 | 17 | 23 |
| 31 | PZ24_DasTFN_10 | 0.55 | 99 | 0 | 4 | 0 | 0 | 2 | 18 | 24 |
| 32 | PZ24_Adamiak_4 | 12.20 | 62 | 0 | 0 | 0 | 0 | 2 | 24 | 26 |
| 33 | PZ24_DasTFN_4 | 1.94 | 99 | 0 | 2 | 0 | 0 | 2 | 24 | 28 |
| 34 | PZ24_DasTFN_7 | 5.82 | 91 | 1 | 5 | 0 | 0 | 4 | 19 | 29 |
| 35 | PZ24_Adamiak_3 | 10.54 | 68 | 0 | 0 | 0 | 0 | 1 | 29 | 30 |
| 36 | PZ24_DasTFN_6 | 4.16 | 96 | 1 | 6 | 0 | 0 | 5 | 18 | 30 |
| 37 | PZ24_DasTFN_1 | 3.33 | 97 | 1 | 6 | 0 | 0 | 5 | 20 | 32 |
| 38 | PZ24_DasTFN_5 | 4.16 | 96 | 1 | 7 | 0 | 0 | 3 | 21 | 32 |

| | | | | | | | | | | |
|---|---|---|---|---|---|---|---|---|---|---|
| 39 | PZ24_Adamiak_2 | 14.42 | 52 | 0 | 0 | 0 | 0 | 5 | 29 | 34 |
| 40 | PZ24_DasTFN_8 | 5.82 | 91 | 0 | 9 | 0 | 1 | 3 | 22 | 35 |
| 41 | PZ24_RNAComposer_2 | 11.92 | 63 | 0 | 0 | 0 | 0 | 4 | 32 | 36 |
| 42 | PZ24_3dRNA_4 | 0.00 | 100 | 1 | 18 | 0 | 0 | 4 | 25 | 48 |
| 43 | PZ24_3dRNA_5 | 1.11 | 99 | 1 | 23 | 0 | 0 | 6 | 18 | 48 |
| 44 | PZ24_3dRNA_2 | 0.00 | 100 | 1 | 26 | 0 | 0 | 8 | 19 | 54 |
| 45 | PZ24_Bujnicki_4 | 0.00 | 100 | 0 | 32 | 6 | 0 | 6 | 16 | 60 |
| 46 | PZ24_Bujnicki_2 | 0.00 | 100 | 0 | 27 | 6 | 0 | 8 | 20 | 61 |
| 47 | PZ24_3dRNA_1 | 0.28 | 99 | 1 | 42 | 0 | 0 | 8 | 16 | 67 |
| 48 | PZ24_3dRNA_3 | 0.00 | 100 | 1 | 39 | 1 | 0 | 12 | 24 | 77 |
| 49 | PZ24_iFoldRNA_1 | 0.83 | 99 | 1 | 61 | 0 | 0 | 3 | 15 | 80 |
| 50 | PZ24_Bujnicki_3 | 0.00 | 100 | 1 | 44 | 9 | 0 | 8 | 25 | 87 |
| 51 | PZ24_Bujnicki_1 | 0.00 | 100 | 0 | 24 | 17 | 0 | 16 | 34 | 91 |
| 52 | PZ24_iFoldRNA_3 | 2.22 | 99 | 1 | 75 | 0 | 0 | 2 | 15 | 93 |
| 53 | PZ24_iFoldRNA_2 | 1.67 | 99 | 1 | 79 | 0 | 1 | 3 | 16 | 100 |
| 54 | PZ24_iFoldRNA_5 | 0.55 | 99 | 1 | 84 | 0 | 0 | 4 | 16 | 105 |
| 55 | PZ24_iFoldRNA_4 | 1.94 | 99 | 1 | 83 | 0 | 1 | 4 | 17 | 106 |
| 56 | PZ24_Vfold3D_4 | 0.00 | 100 | 6 | 68 | 2 | 0 | 4 | 44 | 124 |
| 57 | PZ24_Ding_3 | 13.03 | 58 | 0 | 116 | 0 | 0 | 0 | 11 | 127 |
| 58 | PZ24_Ding_10 | 8.04 | 82 | 0 | 104 | 0 | 0 | 4 | 20 | 128 |
| 59 | PZ24_Ding_8 | 9.71 | 73 | 0 | 109 | 0 | 0 | 4 | 18 | 131 |
| 60 | PZ24_Ding_2 | 10.26 | 70 | 0 | 115 | 0 | 0 | 3 | 15 | 133 |
| 61 | PZ24_Vfold3D_5 | 0.00 | 100 | 9 | 85 | 0 | 1 | 4 | 34 | 133 |
| 62 | PZ24_Ding_7 | 10.82 | 67 | 0 | 119 | 0 | 0 | 2 | 13 | 134 |
| 63 | PZ24_Vfold3D_2 | 0.00 | 100 | 6 | 81 | 0 | 0 | 6 | 44 | 137 |
| 64 | PZ24_Ding_4 | 12.76 | 60 | 0 | 127 | 0 | 0 | 2 | 13 | 142 |
| 65 | PZ24_Ding_6 | 11.37 | 65 | 0 | 124 | 0 | 0 | 3 | 15 | 142 |
| 66 | PZ24_Ding_5 | 9.43 | 74 | 0 | 123 | 0 | 0 | 4 | 16 | 143 |
| 67 | PZ24_Ding_9 | 9.98 | 71 | 0 | 119 | 0 | 0 | 4 | 20 | 143 |
| 68 | PZ24_Ding_1 | 7.49 | 84 | 0 | 131 | 0 | 0 | 4 | 15 | 150 |
| 69 | PZ24_Vfold3D_1 | 0.00 | 100 | 11 | 90 | 0 | 1 | 4 | 44 | 150 |
| 70 | PZ24_Vfold3D_3 | 0.00 | 100 | 27 | 93 | 0 | 1 | 7 | 46 | 174 |
| 71 | PZ24_Bujnicki_5 | 0.00 | 100 | 2 | 192 | 6 | 1 | 10 | 22 | 233 |

| | | | | | | | | | | |
|---|---|---|---|---|---|---|---|---|---|---|
| 72 | PZ24_Kollmann_2 | 128.68 | 0 | 62 | 192 | 1 | 6 | 4 | 21 | 286 |
| 73 | PZ24_Kollmann_4 | 123.19 | 0 | 71 | 216 | 1 | 8 | 5 | 22 | 323 |
| 74 | PZ24_Kollmann_7 | 117.55 | 0 | 67 | 215 | 2 | 6 | 10 | 29 | 329 |
| 75 | PZ24_Kollmann_6 | 92.12 | 0 | 77 | 219 | 0 | 5 | 9 | 25 | 335 |
| 76 | PZ24_Kollmann_10 | 113.64 | 0 | 73 | 233 | 0 | 8 | 8 | 26 | 348 |
| 77 | PZ24_Kollmann_8 | 114.07 | 0 | 75 | 235 | 0 | 7 | 8 | 28 | 353 |
| 78 | PZ24_Kollmann_5 | 121.04 | 0 | 89 | 230 | 0 | 11 | 5 | 26 | 361 |
| 79 | PZ24_Kollmann_3 | 120.66 | 0 | 91 | 236 | 1 | 5 | 7 | 26 | 366 |
| 80 | PZ24_Kollmann_9 | 104.33 | 0 | 85 | 241 | 0 | 8 | 10 | 26 | 370 |
| 81 | PZ24_Kollmann_1 | 115.47 | 0 | 98 | 236 | 0 | 8 | 11 | 29 | 382 |
| 82 | PZ24_SimRNA_2 | 136.29 | 0 | 107 | 243 | 0 | 5 | 3 | 27 | 385 |
| 83 | PZ24_SimRNA_5 | 96.00 | 0 | 120 | 256 | 2 | 4 | 3 | 28 | 413 |
| 84 | PZ24_SimRNA_3 | 124.86 | 0 | 119 | 255 | 0 | 2 | 6 | 32 | 414 |
| 85 | PZ24_SimRNA_1 | 120.70 | 0 | 118 | 273 | 2 | 9 | 5 | 34 | 441 |
| 86 | PZ24_VfoldLA_1 | 144.71 | 0 | 85 | 308 | 0 | 0 | 9 | 43 | 445 |
| 87 | PZ24_VfoldLA_2 | 171.09 | 0 | 94 | 334 | 0 | 1 | 13 | 43 | 485 |
| 88 | PZ24_VfoldLA_3 | 158.98 | 0 | 84 | 350 | 0 | 5 | 14 | 44 | 497 |
| 89 | PZ24_VfoldLA_5 | 147.87 | 0 | 99 | 341 | 0 | 3 | 8 | 49 | 500 |
| 90 | PZ24_VfoldLA_4 | 155.67 | 0 | 99 | 342 | 0 | 3 | 15 | 46 | 505 |
| 91 | PZ24_SimRNA_4 | 99.70 | 0 | 129 | 323 | 0 | 9 | 9 | 41 | 511 |
| | Average | 30.64 | 70.88 | 21.10 | 85.71 | 0.62 | 1.32 | 4.00 | 21.29 | 134.03 |
| | Standard deviation | 51.62 | 39.75 | 39.03 | 106.67 | 2.27 | 2.68 | 3.91 | 10.79 | 154.52 |
| | Median | 3.33 | 97.00 | 0.00 | 27.00 | 0.00 | 0.00 | 3.00 | 19.00 | 61.00 |

**Supplementary Material**

**DICTIONARY OF MAXIT PARAMETERS**

**Table D1**. Conditions for close contact identification.

| | Two adjacent residues in the chain | Two nonadjacent residues in the chain | | | | |
|---|---|---|---|---|---|---|
| | | D – D | D – H | D/H – Any | {*} – Any | Any – Any |
| Distance [Å] | < 1.80 | < 1.415 | < 1.35 | < 1.60 | < 1.70 | < 2.195 |

D – Deuterium
{*} – {Na, Cu, Ca, Mn, Zn, Mg, Ni, Co, Hg, Pt, Ba}
Any – any type of atom except for D, H, and atoms in {*}

**Table D2.** Neutral adenine bond lengths in Angstroms [Å].

| Atom 1 | Atom 2 | Type | Minimum | Maximum | Reference | Standard dev |
|---|---|---|---|---|---|---|
| P | OP1 | s | 1.383 | 1.587 | 1.485 | 0.017 |
| P | OP2 | s | 1.383 | 1.587 | 1.485 | 0.017 |
| P | OP3 | s | 1.535 | 1.679 | 1.607 | 0.012 |
| P | O5' | s | 1.533 | 1.653 | 1.593 | 0.010 |
| O5' | C5' | (P)O5'-C5' | 1.344 | 1.536 | 1.440 | 0.016 |
| O5' | C5' | (H)C2'-endo | 1.328 | 1.520 | 1.424 | 0.016 |
| O5' | C5' | (H)C3'-endo | 1.366 | 1.474 | 1.420 | 0.009 |
| C5' | C4' | C2'-endo | 1.437 | 1.581 | 1.509 | 0.012 |
| C5' | C4' | C3'-endo | 1.466 | 1.550 | 1.508 | 0.007 |
| C4' | C3' | C2'-endo | 1.461 | 1.593 | 1.527 | 0.011 |
| C4' | C3' | C3'-endo | 1.461 | 1.581 | 1.521 | 0.010 |
| C3' | C2' | C2'-endo | 1.459 | 1.591 | 1.525 | 0.011 |
| C3' | C2' | C3'-endo | 1.457 | 1.589 | 1.523 | 0.011 |
| C2' | C1' | C2'-endo | 1.478 | 1.574 | 1.526 | 0.008 |
| C2' | C1' | C3'-endo | 1.463 | 1.595 | 1.529 | 0.011 |
| O4' | C1' | C2'-endo | 1.343 | 1.487 | 1.415 | 0.012 |
| O4' | C1' | C3'-endo | 1.334 | 1.490 | 1.412 | 0.013 |
| O4' | C4' | C2'-endo | 1.394 | 1.514 | 1.454 | 0.010 |
| O4' | C4' | C3'-endo | 1.373 | 1.529 | 1.451 | 0.013 |
| O3' | C3' | C2'-endo | 1.355 | 1.499 | 1.427 | 0.012 |
| O3' | C3' | C3'-endo | 1.333 | 1.501 | 1.417 | 0.014 |
| C1' | N9 | C2'-endo | 1.380 | 1.548 | 1.464 | 0.014 |
| C1' | N9 | C3'-endo | 1.393 | 1.573 | 1.483 | 0.015 |
| C2' | O2' | C2'-endo | 1.334 | 1.490 | 1.412 | 0.013 |
| C2' | O2' | C3'-endo | 1.360 | 1.480 | 1.420 | 0.010 |
| N1 | C2 | s | 1.285 | 1.393 | 1.339 | 0.009 |
| C2 | N3 | s | 1.277 | 1.385 | 1.331 | 0.009 |
| N3 | C4 | s | 1.308 | 1.380 | 1.344 | 0.006 |
| C4 | C5 | s | 1.341 | 1.425 | 1.383 | 0.007 |
| C5 | C6 | s | 1.352 | 1.460 | 1.406 | 0.009 |
| C6 | N1 | s | 1.309 | 1.393 | 1.351 | 0.007 |
| C5 | N7 | s | 1.352 | 1.424 | 1.388 | 0.006 |
| N7 | C8 | s | 1.269 | 1.353 | 1.311 | 0.007 |
| C8 | N9 | s | 1.325 | 1.421 | 1.373 | 0.008 |
| N9 | C4 | s | 1.338 | 1.410 | 1.374 | 0.006 |
| C6 | N6 | s | 1.287 | 1.383 | 1.335 | 0.008 |

**Table D3**. Neutral adenine bond angles in degrees [ ° ].

| Atom 1 | Atom 2 | Atom 3 | Type | Minimum | Maximum | Reference | Standard dev |
|---|---|---|---|---|---|---|---|
| OP1 | P | OP2 | s | 110.6 | 128.6 | 119.6 | 1.5 |
| O5' | P | OP1 | large | 103.5 | 117.9 | 110.7 | 1.2 |
| O5' | P | OP1 | small | 100.3 | 111.1 | 105.7 | 0.9 |
| O5' | P | OP2 | large | 103.5 | 117.9 | 110.7 | 1.2 |
| O5' | P | OP2 | small | 100.3 | 111.1 | 105.7 | 0.9 |
| O5' | C5' | C4' | (P)O5'-C5'-C4' | 104.6 | 114.2 | 109.4 | 0.8 |
| O5' | C5' | C4' | C2'-endo | 100.3 | 123.1 | 111.7 | 1.9 |
| O5' | C5' | C4' | C3'-endo | 101.9 | 121.1 | 111.5 | 1.6 |
| P | O5' | C5' | s | 111.3 | 130.5 | 120.9 | 1.6 |
| O4' | C4' | C3' | C2'-endo | 101.3 | 110.9 | 106.1 | 0.8 |
| O4' | C4' | C3' | C3'-endo | 98.0 | 110.0 | 104.0 | 1.0 |
| C5' | C4' | C3' | C2'-endo | 106.8 | 123.6 | 115.2 | 1.4 |
| C5' | C4' | C3' | C3'-endo | 106.4 | 125.6 | 116.0 | 1.6 |
| C5' | C4' | O4' | C2'-endo | 101.9 | 116.3 | 109.1 | 1.2 |
| C5' | C4' | O4' | C3'-endo | 104.4 | 115.2 | 109.8 | 0.9 |
| C1' | O4' | C4' | C2'-endo | 105.5 | 113.9 | 109.7 | 0.7 |
| C1' | O4' | C4' | C3'-endo | 105.1 | 114.7 | 109.9 | 0.8 |
| C4' | C3' | O3' | C2'-endo | 96.8 | 122.0 | 109.4 | 2.1 |
| C4' | C3' | O3' | C3'-endo | 101.0 | 125.0 | 113.0 | 2.0 |
| C2' | C3' | O3' | C2'-endo | 96.3 | 122.7 | 109.5 | 2.2 |
| C2' | C3' | O3' | C3'-endo | 104.1 | 123.3 | 113.7 | 1.6 |
| C4' | C3' | C2' | C2'-endo | 96.6 | 108.6 | 102.6 | 1.0 |
| C4' | C3' | C2' | C3'-endo | 96.6 | 108.6 | 102.6 | 1.0 |
| C3' | C2' | C1' | C2'-endo | 96.7 | 106.3 | 101.5 | 0.8 |
| C3' | C2' | C1' | C3'-endo | 97.1 | 105.5 | 101.3 | 0.7 |
| O4' | C1' | C2' | C2'-endo | 99.8 | 111.8 | 105.8 | 1.0 |
| O4' | C1' | C2' | C3'-endo | 102.2 | 113.0 | 107.6 | 0.9 |
| N9 | C1' | C2' | C2'-endo | 106.2 | 121.8 | 114.0 | 1.3 |
| N9 | C1' | C2' | C3'-endo | 105.4 | 118.6 | 112.0 | 1.1 |
| O4' | C1' | N9 | C2'-endo | 103.4 | 113.0 | 108.2 | 0.8 |
| O4' | C1' | N9 | C3'-endo | 104.3 | 112.7 | 108.5 | 0.7 |
| C1' | C2' | O2' | s | 92.6 | 128.6 | 110.6 | 3.0 |
| C3' | C2' | O2' | s | 95.9 | 130.7 | 113.3 | 2.9 |
| C6 | N1 | C2 | s | 115.0 | 122.2 | 118.6 | 0.6 |
| N1 | C2 | N3 | s | 126.3 | 132.3 | 129.3 | 0.5 |
| C2 | N3 | C4 | s | 107.6 | 113.6 | 110.6 | 0.5 |
| N3 | C4 | C5 | s | 122.6 | 131.0 | 126.8 | 0.7 |
| C4 | C5 | C6 | s | 114.0 | 120.0 | 117.0 | 0.5 |
| C5 | C6 | N1 | s | 114.7 | 120.7 | 117.7 | 0.5 |
| C4 | C5 | N7 | s | 107.7 | 113.7 | 110.7 | 0.5 |
| C5 | N7 | C8 | s | 100.9 | 106.9 | 103.9 | 0.5 |
| N7 | C8 | N9 | s | 110.8 | 116.8 | 113.8 | 0.5 |
| C8 | N9 | C4 | s | 103.4 | 108.2 | 105.8 | 0.4 |
| N9 | C4 | C5 | s | 103.4 | 108.2 | 105.8 | 0.4 |
| N3 | C4 | N9 | s | 122.6 | 132.2 | 127.4 | 0.8 |
| C6 | C5 | N7 | s | 128.1 | 136.5 | 132.3 | 0.7 |
| N1 | C6 | N6 | s | 115.0 | 122.2 | 118.6 | 0.6 |
| C5 | C6 | N6 | s | 118.9 | 128.5 | 123.7 | 0.8 |
| C8 | N9 | C1' | s | 116.9 | 138.5 | 127.7 | 1.8 |
| C4 | N9 | C1' | s | 115.5 | 137.1 | 126.3 | 1.8 |

**Table D4**. Neutral cytosine bond lengths in Angstroms [Å].

| Atom 1 | Atom 2 | Type | Minimum | Maximum | Reference | Standard dev |
|---|---|---|---|---|---|---|
| P | OP1 | s | 1.383 | 1.587 | 1.485 | 0.017 |
| P | OP2 | s | 1.383 | 1.587 | 1.485 | 0.017 |
| P | OP3 | s | 1.535 | 1.679 | 1.607 | 0.012 |
| P | O5' | s | 1.533 | 1.653 | 1.593 | 0.010 |
| O5' | C5' | (P)O5'-C5' | 1.344 | 1.536 | 1.440 | 0.016 |
| O5' | C5' | (H)C2'-endo | 1.328 | 1.520 | 1.424 | 0.016 |
| O5' | C5' | (H)C3'-endo | 1.366 | 1.474 | 1.420 | 0.009 |
| C5' | C4' | C2'-endo | 1.437 | 1.581 | 1.509 | 0.012 |
| C5' | C4' | C3'-endo | 1.466 | 1.550 | 1.508 | 0.007 |
| C4' | C3' | C2'-endo | 1.461 | 1.593 | 1.527 | 0.011 |
| C4' | C3' | C3'-endo | 1.461 | 1.581 | 1.521 | 0.010 |
| C3' | C2' | C2'-endo | 1.459 | 1.591 | 1.525 | 0.011 |
| C3' | C2' | C3'-endo | 1.457 | 1.589 | 1.523 | 0.011 |
| C2' | C1' | C2'-endo | 1.478 | 1.574 | 1.526 | 0.008 |
| C2' | C1' | C3'-endo | 1.463 | 1.595 | 1.529 | 0.011 |
| O4' | C1' | C2'-endo | 1.343 | 1.487 | 1.415 | 0.012 |
| O4' | C1' | C3'-endo | 1.334 | 1.490 | 1.412 | 0.013 |
| O4' | C4' | C2'-endo | 1.394 | 1.514 | 1.454 | 0.010 |
| O4' | C4' | C3'-endo | 1.373 | 1.529 | 1.451 | 0.013 |
| O3' | C3' | C2'-endo | 1.355 | 1.499 | 1.427 | 0.012 |
| O3' | C3' | C3'-endo | 1.333 | 1.501 | 1.417 | 0.014 |
| C1' | N1 | C2'-endo | 1.380 | 1.548 | 1.464 | 0.014 |
| C1' | N1 | C3'-endo | 1.393 | 1.573 | 1.483 | 0.015 |
| C2' | O2' | C2'-endo | 1.334 | 1.490 | 1.412 | 0.013 |
| C2' | O2' | C3'-endo | 1.360 | 1.480 | 1.420 | 0.010 |
| C2 | O2 | s | 1.186 | 1.294 | 1.240 | 0.009 |
| C4 | N4 | s | 1.281 | 1.389 | 1.335 | 0.009 |
| N1 | C2 | s | 1.337 | 1.457 | 1.397 | 0.010 |
| N1 | C6 | s | 1.331 | 1.403 | 1.367 | 0.006 |
| C2 | N3 | s | 1.305 | 1.401 | 1.353 | 0.008 |
| N3 | C4 | s | 1.293 | 1.377 | 1.335 | 0.007 |
| C4 | C5 | s | 1.377 | 1.473 | 1.425 | 0.008 |
| C5 | C6 | s | 1.291 | 1.387 | 1.339 | 0.008 |

**Table D5**. Neutral cytosine bond angles in degrees [ ° ].

| Atom 1 | Atom 2 | Atom 3 | Type | Minimum | Maximum | Reference | Standard dev |
|---|---|---|---|---|---|---|---|
| OP1 | P | OP2 | s | 110.6 | 128.6 | 119.6 | 1.5 |
| O5' | P | OP1 | large | 103.5 | 117.9 | 110.7 | 1.2 |
| O5' | P | OP1 | small | 100.3 | 111.1 | 105.7 | 0.9 |
| O5' | P | OP2 | large | 103.5 | 117.9 | 110.7 | 1.2 |
| O5' | P | OP2 | small | 100.3 | 111.1 | 105.7 | 0.9 |
| O5' | C5' | C4' | (P)O5'-C5'-C4' | 104.6 | 114.2 | 109.4 | 0.8 |
| O5' | C5' | C4' | C2'-endo | 100.3 | 123.1 | 111.7 | 1.9 |
| O5' | C5' | C4' | C3'-endo | 101.9 | 121.1 | 111.5 | 1.6 |
| P | O5' | C5' | s | 111.3 | 130.5 | 120.9 | 1.6 |
| O4' | C4' | C3' | C2'-endo | 101.3 | 110.9 | 106.1 | 0.8 |
| O4' | C4' | C3' | C3'-endo | 98.0 | 110.0 | 104.0 | 1.0 |
| C5' | C4' | C3' | C2'-endo | 106.8 | 123.6 | 115.2 | 1.4 |
| C5' | C4' | C3' | C3'-endo | 106.4 | 125.6 | 116.0 | 1.6 |
| C5' | C4' | O4' | C2'-endo | 101.9 | 116.3 | 109.1 | 1.2 |
| C5' | C4' | O4' | C3'-endo | 104.4 | 115.2 | 109.8 | 0.9 |
| C1' | O4' | C4' | C2'-endo | 105.5 | 113.9 | 109.7 | 0.7 |
| C1' | O4' | C4' | C3'-endo | 105.1 | 114.7 | 109.9 | 0.8 |
| C4' | C3' | O3' | C2'-endo | 96.8 | 122.0 | 109.4 | 2.1 |
| C4' | C3' | O3' | C3'-endo | 101.0 | 125.0 | 113.0 | 2.0 |
| C2' | C3' | O3' | C2'-endo | 96.3 | 122.7 | 109.5 | 2.2 |
| C2' | C3' | O3' | C3'-endo | 104.1 | 123.3 | 113.7 | 1.6 |
| C4' | C3' | C2' | C2'-endo | 96.6 | 108.6 | 102.6 | 1.0 |
| C4' | C3' | C2' | C3'-endo | 96.6 | 108.6 | 102.6 | 1.0 |
| C3' | C2' | C1' | C2'-endo | 96.7 | 106.3 | 101.5 | 0.8 |
| C3' | C2' | C1' | C3'-endo | 97.1 | 105.5 | 101.3 | 0.7 |
| O4' | C1' | C2' | C2'-endo | 99.8 | 111.8 | 105.8 | 1.0 |
| O4' | C1' | C2' | C3'-endo | 102.2 | 113.0 | 107.6 | 0.9 |
| N1 | C1' | C2' | C2'-endo | 106.2 | 121.8 | 114.0 | 1.3 |
| N1 | C1' | C2' | C3'-endo | 105.4 | 118.6 | 112.0 | 1.1 |
| O4' | C1' | N1 | C2'-endo | 103.4 | 113.0 | 108.2 | 0.8 |
| O4' | C1' | N1 | C3'-endo | 104.3 | 112.7 | 108.5 | 0.7 |
| C1' | C2' | O2' | s | 92.6 | 128.6 | 110.6 | 3.0 |
| C3' | C2' | O2' | s | 95.9 | 130.7 | 113.3 | 2.9 |
| C6 | N1 | C2 | s | 117.9 | 122.7 | 120.3 | 0.4 |
| N1 | C2 | N3 | s | 115.0 | 123.4 | 119.2 | 0.7 |
| C2 | N3 | C4 | s | 116.9 | 122.9 | 119.9 | 0.5 |
| N3 | C4 | C5 | s | 119.5 | 124.3 | 121.9 | 0.4 |
| C4 | C5 | C6 | s | 114.4 | 120.4 | 117.4 | 0.5 |
| C5 | C6 | N1 | s | 118.0 | 124.0 | 121.0 | 0.5 |
| N1 | C2 | O2 | s | 115.3 | 122.5 | 118.9 | 0.6 |
| N3 | C2 | O2 | s | 117.7 | 126.1 | 121.9 | 0.7 |
| N3 | C4 | N4 | s | 113.8 | 122.2 | 118.0 | 0.7 |
| C5 | C4 | N4 | s | 116.0 | 124.4 | 120.2 | 0.7 |
| C6 | N1 | C1' | s | 113.6 | 128.0 | 120.8 | 1.2 |
| C2 | N1 | C1' | s | 112.2 | 125.4 | 118.8 | 1.1 |

**Table D6**. Neutral guanine bond lengths in Angstroms [Å].

| Atom 1 | Atom 2 | Type | Minimum | Maximum | Reference | Standard dev |
|---|---|---|---|---|---|---|
| P | OP1 | s | 1.383 | 1.587 | 1.485 | 0.017 |
| P | OP2 | s | 1.383 | 1.587 | 1.485 | 0.017 |
| P | OP3 | s | 1.535 | 1.679 | 1.607 | 0.012 |
| P | O5' | s | 1.533 | 1.653 | 1.593 | 0.010 |
| O5' | C5' | (P)O5'-C5' | 1.344 | 1.536 | 1.440 | 0.016 |
| O5' | C5' | (H)C2'-endo | 1.328 | 1.520 | 1.424 | 0.016 |
| O5' | C5' | (H)C3'-endo | 1.366 | 1.474 | 1.420 | 0.009 |
| C5' | C4' | C2'-endo | 1.437 | 1.581 | 1.509 | 0.012 |
| C5' | C4' | C3'-endo | 1.466 | 1.550 | 1.508 | 0.007 |
| C4' | C3' | C2'-endo | 1.461 | 1.593 | 1.527 | 0.011 |
| C4' | C3' | C3'-endo | 1.461 | 1.581 | 1.521 | 0.010 |
| C3' | C2' | C2'-endo | 1.459 | 1.591 | 1.525 | 0.011 |
| C3' | C2' | C3'-endo | 1.457 | 1.589 | 1.523 | 0.011 |
| C2' | C1' | C2'-endo | 1.478 | 1.574 | 1.526 | 0.008 |
| C2' | C1' | C3'-endo | 1.463 | 1.595 | 1.529 | 0.011 |
| O4' | C1' | C2'-endo | 1.343 | 1.487 | 1.415 | 0.012 |
| O4' | C1' | C3'-endo | 1.334 | 1.490 | 1.412 | 0.013 |
| O4' | C4' | C2'-endo | 1.394 | 1.514 | 1.454 | 0.010 |
| O4' | C4' | C3'-endo | 1.373 | 1.529 | 1.451 | 0.013 |
| O3' | C3' | C2'-endo | 1.355 | 1.499 | 1.427 | 0.012 |
| O3' | C3' | C3'-endo | 1.333 | 1.501 | 1.417 | 0.014 |
| C1' | N9 | C2'-endo | 1.380 | 1.548 | 1.464 | 0.014 |
| C1' | N9 | C3'-endo | 1.393 | 1.573 | 1.483 | 0.015 |
| C2' | O2' | C2'-endo | 1.334 | 1.490 | 1.412 | 0.013 |
| C2' | O2' | C3'-endo | 1.360 | 1.480 | 1.420 | 0.010 |
| N1 | C2 | s | 1.325 | 1.421 | 1.373 | 0.008 |
| C2 | N3 | s | 1.275 | 1.371 | 1.323 | 0.008 |
| N3 | C4 | s | 1.308 | 1.392 | 1.350 | 0.007 |
| C4 | C5 | s | 1.337 | 1.421 | 1.379 | 0.007 |
| C5 | C6 | s | 1.359 | 1.479 | 1.419 | 0.010 |
| C6 | N1 | s | 1.349 | 1.433 | 1.391 | 0.007 |
| C5 | N7 | s | 1.352 | 1.424 | 1.388 | 0.006 |
| N7 | C8 | s | 1.269 | 1.341 | 1.305 | 0.006 |
| C8 | N9 | s | 1.332 | 1.416 | 1.374 | 0.007 |
| N9 | C4 | s | 1.327 | 1.423 | 1.375 | 0.008 |
| C2 | N2 | s | 1.281 | 1.401 | 1.341 | 0.010 |
| C6 | O6 | s | 1.183 | 1.291 | 1.237 | 0.009 |

**Table D7**. Neutral guanine bond angles in degrees [ ° ].

| Atom 1 | Atom 2 | Atom 3 | Type | Minimum | Maximum | Reference | Standard dev |
|---|---|---|---|---|---|---|---|
| OP1 | P | OP2 | s | 110.6 | 128.6 | 119.6 | 1.5 |
| O5' | P | OP1 | large | 103.5 | 117.9 | 110.7 | 1.2 |
| O5' | P | OP1 | small | 100.3 | 111.1 | 105.7 | 0.9 |
| O5' | P | OP2 | large | 103.5 | 117.9 | 110.7 | 1.2 |
| O5' | P | OP2 | small | 100.3 | 111.1 | 105.7 | 0.9 |
| O5' | C5' | C4' | (P)O5'-C5'-C4' | 104.6 | 114.2 | 109.4 | 0.8 |
| O5' | C5' | C4' | C2'-endo | 100.3 | 123.1 | 111.7 | 1.9 |
| O5' | C5' | C4' | C3'-endo | 101.9 | 121.1 | 111.5 | 1.6 |
| P | O5' | C5' | s | 111.3 | 130.5 | 120.9 | 1.6 |
| O4' | C4' | C3' | C2'-endo | 101.3 | 110.9 | 106.1 | 0.8 |
| O4' | C4' | C3' | C3'-endo | 98.0 | 110.0 | 104.0 | 1.0 |
| C5' | C4' | C3' | C2'-endo | 106.8 | 123.6 | 115.2 | 1.4 |
| C5' | C4' | C3' | C3'-endo | 106.4 | 125.6 | 116.0 | 1.6 |
| C5' | C4' | O4' | C2'-endo | 101.9 | 116.3 | 109.1 | 1.2 |
| C5' | C4' | O4' | C3'-endo | 104.4 | 115.2 | 109.8 | 0.9 |
| C1' | O4' | C4' | C2'-endo | 105.5 | 113.9 | 109.7 | 0.7 |
| C1' | O4' | C4' | C3'-endo | 105.1 | 114.7 | 109.9 | 0.8 |
| C4' | C3' | O3' | C2'-endo | 96.8 | 122.0 | 109.4 | 2.1 |
| C4' | C3' | O3' | C3'-endo | 101.0 | 125.0 | 113.0 | 2.0 |
| C2' | C3' | O3' | C2'-endo | 96.3 | 122.7 | 109.5 | 2.2 |
| C2' | C3' | O3' | C3'-endo | 104.1 | 123.3 | 113.7 | 1.6 |
| C4' | C3' | C2' | C2'-endo | 96.6 | 108.6 | 102.6 | 1.0 |
| C4' | C3' | C2' | C3'-endo | 96.6 | 108.6 | 102.6 | 1.0 |
| C3' | C2' | C1' | C2'-endo | 96.7 | 106.3 | 101.5 | 0.8 |
| C3' | C2' | C1' | C3'-endo | 97.1 | 105.5 | 101.3 | 0.7 |
| O4' | C1' | C2' | C2'-endo | 99.8 | 111.8 | 105.8 | 1.0 |
| O4' | C1' | C2' | C3'-endo | 102.2 | 113.0 | 107.6 | 0.9 |
| N9 | C1' | C2' | C2'-endo | 106.2 | 121.8 | 114.0 | 1.3 |
| N9 | C1' | C2' | C3'-endo | 105.4 | 118.6 | 112.0 | 1.1 |
| O4' | C1' | N9 | C2'-endo | 103.4 | 113.0 | 108.2 | 0.8 |
| O4' | C1' | N9 | C3'-endo | 104.3 | 112.7 | 108.5 | 0.7 |
| C1' | C2' | O2' | s | 92.6 | 128.6 | 110.6 | 3.0 |
| C3' | C2' | O2' | s | 95.9 | 130.7 | 113.3 | 2.9 |
| C6 | N1 | C2 | s | 121.5 | 128.7 | 125.1 | 0.6 |
| N1 | C2 | N3 | s | 120.3 | 127.5 | 123.9 | 0.6 |
| C2 | N3 | C4 | s | 108.9 | 114.9 | 111.9 | 0.5 |
| N3 | C4 | C5 | s | 125.6 | 131.6 | 128.6 | 0.5 |
| C4 | C5 | C6 | s | 115.2 | 122.4 | 118.8 | 0.6 |
| C5 | C6 | N1 | s | 108.5 | 114.5 | 111.5 | 0.5 |
| C4 | C5 | N7 | s | 108.4 | 113.2 | 110.8 | 0.4 |
| C5 | N7 | C8 | s | 101.3 | 107.3 | 104.3 | 0.5 |
| N7 | C8 | N9 | s | 110.1 | 116.1 | 113.1 | 0.5 |
| C8 | N9 | C4 | s | 104.0 | 108.8 | 106.4 | 0.4 |
| N9 | C4 | C5 | s | 103.0 | 107.8 | 105.4 | 0.4 |
| N3 | C4 | N9 | s | 122.4 | 129.6 | 126.0 | 0.6 |
| C6 | C5 | N7 | s | 126.8 | 134.0 | 130.4 | 0.6 |
| N1 | C2 | N2 | s | 110.8 | 121.6 | 116.2 | 0.9 |
| N3 | C2 | N2 | s | 115.7 | 124.1 | 119.9 | 0.7 |
| N1 | C6 | O6 | s | 116.3 | 123.5 | 119.9 | 0.6 |
| C5 | C6 | O6 | s | 125.0 | 132.2 | 128.6 | 0.6 |
| C8 | N9 | C1' | s | 119.2 | 134.8 | 127.0 | 1.3 |
| C4 | N9 | C1' | s | 118.7 | 134.3 | 126.5 | 1.3 |

**Table D8**. Neutral uracil bond lengths in Angstroms [Å].

| Atom 1 | Atom 2 | Type | Minimum | Maximum | Reference | Standard dev |
|---|---|---|---|---|---|---|
| P | OP1 | s | 1.383 | 1.587 | 1.485 | 0.017 |
| P | OP2 | s | 1.383 | 1.587 | 1.485 | 0.017 |
| P | OP3 | s | 1.535 | 1.679 | 1.607 | 0.012 |
| P | O5' | s | 1.533 | 1.653 | 1.593 | 0.010 |
| O5' | C5' | (P)O5'-C5' | 1.344 | 1.536 | 1.440 | 0.016 |
| O5' | C5' | (H)C2'-endo | 1.328 | 1.520 | 1.424 | 0.016 |
| O5' | C5' | (H)C3'-endo | 1.366 | 1.474 | 1.420 | 0.009 |
| C5' | C4' | C2'-endo | 1.437 | 1.581 | 1.509 | 0.012 |
| C5' | C4' | C3'-endo | 1.466 | 1.550 | 1.508 | 0.007 |
| C4' | C3' | C2'-endo | 1.461 | 1.593 | 1.527 | 0.011 |
| C4' | C3' | C3'-endo | 1.461 | 1.581 | 1.521 | 0.010 |
| C3' | C2' | C2'-endo | 1.459 | 1.591 | 1.525 | 0.011 |
| C3' | C2' | C3'-endo | 1.457 | 1.589 | 1.523 | 0.011 |
| C2' | C1' | C2'-endo | 1.478 | 1.574 | 1.526 | 0.008 |
| C2' | C1' | C3'-endo | 1.463 | 1.595 | 1.529 | 0.011 |
| O4' | C1' | C2'-endo | 1.343 | 1.487 | 1.415 | 0.012 |
| O4' | C1' | C3'-endo | 1.334 | 1.490 | 1.412 | 0.013 |
| O4' | C4' | C2'-endo | 1.394 | 1.514 | 1.454 | 0.010 |
| O4' | C4' | C3'-endo | 1.373 | 1.529 | 1.451 | 0.013 |
| O3' | C3' | C2'-endo | 1.355 | 1.499 | 1.427 | 0.012 |
| O3' | C3' | C3'-endo | 1.333 | 1.501 | 1.417 | 0.014 |
| C1' | N1 | C2'-endo | 1.380 | 1.548 | 1.464 | 0.014 |
| C1' | N1 | C3'-endo | 1.393 | 1.573 | 1.483 | 0.015 |
| C2' | O2' | C2'-endo | 1.334 | 1.490 | 1.412 | 0.013 |
| C2' | O2' | C3'-endo | 1.360 | 1.480 | 1.420 | 0.010 |
| C2 | O2 | s | 1.165 | 1.273 | 1.219 | 0.009 |
| C4 | O4 | s | 1.184 | 1.280 | 1.232 | 0.008 |
| N1 | C2 | s | 1.327 | 1.435 | 1.381 | 0.009 |
| N1 | C6 | s | 1.321 | 1.429 | 1.375 | 0.009 |
| C2 | N3 | s | 1.331 | 1.415 | 1.373 | 0.007 |
| N3 | C4 | s | 1.326 | 1.434 | 1.380 | 0.009 |
| C4 | C5 | s | 1.377 | 1.485 | 1.431 | 0.009 |
| C5 | C6 | s | 1.283 | 1.391 | 1.337 | 0.009 |

**Table D9**. Neutral uracil bond angles in degrees [ ° ].

| Atom 1 | Atom 2 | Atom 3 | Type | Minimum | Maximum | Reference | Standard dev |
|---|---|---|---|---|---|---|---|
| OP1 | P | OP2 | s | 110.6 | 128.6 | 119.6 | 1.5 |
| O5' | P | OP1 | large | 103.5 | 117.9 | 110.7 | 1.2 |
| O5' | P | OP1 | small | 100.3 | 111.1 | 105.7 | 0.9 |
| O5' | P | OP2 | large | 103.5 | 117.9 | 110.7 | 1.2 |
| O5' | P | OP2 | small | 100.3 | 111.1 | 105.7 | 0.9 |
| O5' | C5' | C4' | (P)O5'-C5'-C4' | 104.6 | 114.2 | 109.4 | 0.8 |
| O5' | C5' | C4' | C2'-endo | 100.3 | 123.1 | 111.7 | 1.9 |
| O5' | C5' | C4' | C3'-endo | 101.9 | 121.1 | 111.5 | 1.6 |
| P | O5' | C5' | s | 111.3 | 130.5 | 120.9 | 1.6 |
| O4' | C4' | C3' | C2'-endo | 101.3 | 110.9 | 106.1 | 0.8 |
| O4' | C4' | C3' | C3'-endo | 98.0 | 110.0 | 104.0 | 1.0 |
| C5' | C4' | C3' | C2'-endo | 106.8 | 123.6 | 115.2 | 1.4 |
| C5' | C4' | C3' | C3'-endo | 106.4 | 125.6 | 116.0 | 1.6 |
| C5' | C4' | O4' | C2'-endo | 101.9 | 116.3 | 109.1 | 1.2 |
| C5' | C4' | O4' | C3'-endo | 104.4 | 115.2 | 109.8 | 0.9 |
| C1' | O4' | C4' | C2'-endo | 105.5 | 113.9 | 109.7 | 0.7 |
| C1' | O4' | C4' | C3'-endo | 105.1 | 114.7 | 109.9 | 0.8 |
| C4' | C3' | O3' | C2'-endo | 96.8 | 122.0 | 109.4 | 2.1 |
| C4' | C3' | O3' | C3'-endo | 101.0 | 125.0 | 113.0 | 2.0 |
| C2' | C3' | O3' | C2'-endo | 96.3 | 122.7 | 109.5 | 2.2 |
| C2' | C3' | O3' | C3'-endo | 104.1 | 123.3 | 113.7 | 1.6 |
| C4' | C3' | C2' | C2'-endo | 96.6 | 108.6 | 102.6 | 1.0 |
| C4' | C3' | C2' | C3'-endo | 96.6 | 108.6 | 102.6 | 1.0 |
| C3' | C2' | C1' | C2'-endo | 96.7 | 106.3 | 101.5 | 0.8 |
| C3' | C2' | C1' | C3'-endo | 97.1 | 105.5 | 101.3 | 0.7 |
| O4' | C1' | C2' | C2'-endo | 99.8 | 111.8 | 105.8 | 1.0 |
| O4' | C1' | C2' | C3'-endo | 102.2 | 113.0 | 107.6 | 0.9 |
| N1 | C1' | C2' | C2'-endo | 106.2 | 121.8 | 114.0 | 1.3 |
| N1 | C1' | C2' | C3'-endo | 105.4 | 118.6 | 112.0 | 1.1 |
| O4' | C1' | N1 | C2'-endo | 103.4 | 113.0 | 108.2 | 0.8 |
| O4' | C1' | N1 | C3'-endo | 104.3 | 112.7 | 108.5 | 0.7 |
| C1' | C2' | O2' | s | 92.6 | 128.6 | 110.6 | 3.0 |
| C3' | C2' | O2' | s | 95.9 | 130.7 | 113.3 | 2.9 |
| C6 | N1 | C2 | s | 117.4 | 124.6 | 121.0 | 0.6 |
| N1 | C2 | N3 | s | 111.3 | 118.5 | 114.9 | 0.6 |
| C2 | N3 | C4 | s | 123.4 | 130.6 | 127.0 | 0.6 |
| N3 | C4 | C5 | s | 111.0 | 118.2 | 114.6 | 0.6 |
| C4 | C5 | C6 | s | 116.1 | 123.3 | 119.7 | 0.6 |
| C5 | C6 | N1 | s | 119.7 | 125.7 | 122.7 | 0.5 |
| N1 | C2 | O2 | s | 118.6 | 127.0 | 122.8 | 0.7 |
| N3 | C2 | O2 | s | 118.0 | 126.4 | 122.2 | 0.7 |
| N3 | C4 | O4 | s | 115.2 | 123.6 | 119.4 | 0.7 |
| C5 | C4 | O4 | s | 122.3 | 129.5 | 125.9 | 0.6 |
| C6 | N1 | C1' | s | 112.8 | 129.6 | 121.2 | 1.4 |
| C2 | N1 | C1' | s | 110.5 | 124.9 | 117.7 | 1.2 |

**Table D10**. Adenine bond lengths in Angstroms [Å].

| Atom 1 | Atom 2 | Type | Minimum | Maximum | Reference | Standard dev |
|---|---|---|---|---|---|---|
| P | OP1 | s | 1.383 | 1.587 | 1.485 | 0.017 |
| P | OP2 | s | 1.383 | 1.587 | 1.485 | 0.017 |
| P | OP3 | s | 1.535 | 1.679 | 1.607 | 0.012 |
| P | O5' | s | 1.533 | 1.653 | 1.593 | 0.010 |
| O5' | C5' | (P)O5'-C5' | 1.344 | 1.536 | 1.440 | 0.016 |
| O5' | C5' | (H)C2'-endo | 1.328 | 1.520 | 1.424 | 0.016 |
| O5' | C5' | (H)C3'-endo | 1.366 | 1.474 | 1.420 | 0.009 |
| C5' | C4' | C2'-endo | 1.437 | 1.581 | 1.509 | 0.012 |
| C5' | C4' | C3'-endo | 1.466 | 1.550 | 1.508 | 0.007 |
| C4' | C3' | C2'-endo | 1.461 | 1.593 | 1.527 | 0.011 |
| C4' | C3' | C3'-endo | 1.461 | 1.581 | 1.521 | 0.010 |
| C3' | C2' | C2'-endo | 1.459 | 1.591 | 1.525 | 0.011 |
| C3' | C2' | C3'-endo | 1.457 | 1.589 | 1.523 | 0.011 |
| C2' | C1' | C2'-endo | 1.478 | 1.574 | 1.526 | 0.008 |
| C2' | C1' | C3'-endo | 1.463 | 1.595 | 1.529 | 0.011 |
| O4' | C1' | C2'-endo | 1.343 | 1.487 | 1.415 | 0.012 |
| O4' | C1' | C3'-endo | 1.334 | 1.490 | 1.412 | 0.013 |
| O4' | C4' | C2'-endo | 1.394 | 1.514 | 1.454 | 0.010 |
| O4' | C4' | C3'-endo | 1.373 | 1.529 | 1.451 | 0.013 |
| O3' | C3' | C2'-endo | 1.355 | 1.499 | 1.427 | 0.012 |
| O3' | C3' | C3'-endo | 1.333 | 1.501 | 1.417 | 0.014 |
| C1' | N9 | C2'-endo | 1.380 | 1.548 | 1.464 | 0.014 |
| C1' | N9 | C3'-endo | 1.393 | 1.573 | 1.483 | 0.015 |
| C2' | O2' | C2'-endo | 1.334 | 1.490 | 1.412 | 0.013 |
| C2' | O2' | C3'-endo | 1.360 | 1.480 | 1.420 | 0.010 |
| N1 | C2 | s | 1.285 | 1.393 | 1.339 | 0.009 |
| C2 | N3 | s | 1.277 | 1.385 | 1.331 | 0.009 |
| N3 | C4 | s | 1.308 | 1.380 | 1.344 | 0.006 |
| C4 | C5 | s | 1.341 | 1.425 | 1.383 | 0.007 |
| C5 | C6 | s | 1.352 | 1.460 | 1.406 | 0.009 |
| C6 | N1 | s | 1.309 | 1.393 | 1.351 | 0.007 |
| C5 | N7 | s | 1.352 | 1.424 | 1.388 | 0.006 |
| N7 | C8 | s | 1.269 | 1.353 | 1.311 | 0.007 |
| C8 | N9 | s | 1.325 | 1.421 | 1.373 | 0.008 |
| N9 | C4 | s | 1.338 | 1.410 | 1.374 | 0.006 |
| C6 | N6 | s | 1.287 | 1.383 | 1.335 | 0.008 |

**Table D11**. Adenine bond angles in degrees [ ° ].

| Atom 1 | Atom 2 | Atom 3 | Type | Minimum | Maximum | Reference | Standard dev |
|---|---|---|---|---|---|---|---|
| OP1 | P | OP2 | s | 110.6 | 128.6 | 119.6 | 1.5 |
| O5' | P | OP1 | large | 103.5 | 117.9 | 110.7 | 1.2 |
| O5' | P | OP1 | small | 100.3 | 111.1 | 105.7 | 0.9 |
| O5' | P | OP2 | large | 103.5 | 117.9 | 110.7 | 1.2 |
| O5' | P | OP2 | small | 100.3 | 111.1 | 105.7 | 0.9 |
| O5' | C5' | C4' | (P)O5'-C5'-C4' | 104.6 | 114.2 | 109.4 | 0.8 |
| O5' | C5' | C4' | C2'-endo | 100.3 | 123.1 | 111.7 | 1.9 |
| O5' | C5' | C4' | C3'-endo | 101.9 | 121.1 | 111.5 | 1.6 |
| P | O5' | C5' | s | 111.3 | 130.5 | 120.9 | 1.6 |
| O4' | C4' | C3' | C2'-endo | 101.3 | 110.9 | 106.1 | 0.8 |
| O4' | C4' | C3' | C3'-endo | 98.0 | 110.0 | 104.0 | 1.0 |
| C5' | C4' | C3' | C2'-endo | 106.8 | 123.6 | 115.2 | 1.4 |
| C5' | C4' | C3' | C3'-endo | 106.4 | 125.6 | 116.0 | 1.6 |
| C5' | C4' | O4' | C2'-endo | 101.9 | 116.3 | 109.1 | 1.2 |
| C5' | C4' | O4' | C3'-endo | 104.4 | 115.2 | 109.8 | 0.9 |
| C1' | O4' | C4' | C2'-endo | 105.5 | 113.9 | 109.7 | 0.7 |
| C1' | O4' | C4' | C3'-endo | 105.1 | 114.7 | 109.9 | 0.8 |
| C4' | C3' | O3' | C2'-endo | 96.8 | 122.0 | 109.4 | 2.1 |
| C4' | C3' | O3' | C3'-endo | 101.0 | 125.0 | 113.0 | 2.0 |
| C2' | C3' | O3' | C2'-endo | 96.3 | 122.7 | 109.5 | 2.2 |
| C2' | C3' | O3' | C3'-endo | 104.1 | 123.3 | 113.7 | 1.6 |
| C4' | C3' | C2' | C2'-endo | 96.6 | 108.6 | 102.6 | 1.0 |
| C4' | C3' | C2' | C3'-endo | 96.6 | 108.6 | 102.6 | 1.0 |
| C3' | C2' | C1' | C2'-endo | 96.7 | 106.3 | 101.5 | 0.8 |
| C3' | C2' | C1' | C3'-endo | 97.1 | 105.5 | 101.3 | 0.7 |
| O4' | C1' | C2' | C2'-endo | 99.8 | 111.8 | 105.8 | 1.0 |
| O4' | C1' | C2' | C3'-endo | 102.2 | 113.0 | 107.6 | 0.9 |
| N9 | C1' | C2' | C2'-endo | 106.2 | 121.8 | 114.0 | 1.3 |
| N9 | C1' | C2' | C3'-endo | 105.4 | 118.6 | 112.0 | 1.1 |
| O4' | C1' | N9 | C2'-endo | 103.4 | 113.0 | 108.2 | 0.8 |
| O4' | C1' | N9 | C3'-endo | 104.3 | 112.7 | 108.5 | 0.7 |
| C1' | C2' | O2' | s | 92.6 | 128.6 | 110.6 | 3.0 |
| C3' | C2' | O2' | s | 95.9 | 130.7 | 113.3 | 2.9 |
| C6 | N1 | C2 | s | 115.0 | 122.2 | 118.6 | 0.6 |
| N1 | C2 | N3 | s | 126.3 | 132.3 | 129.3 | 0.5 |
| C2 | N3 | C4 | s | 107.6 | 113.6 | 110.6 | 0.5 |
| N3 | C4 | C5 | s | 122.6 | 131.0 | 126.8 | 0.7 |
| C4 | C5 | C6 | s | 114.0 | 120.0 | 117.0 | 0.5 |
| C5 | C6 | N1 | s | 114.7 | 120.7 | 117.7 | 0.5 |
| C4 | C5 | N7 | s | 107.7 | 113.7 | 110.7 | 0.5 |
| C5 | N7 | C8 | s | 100.9 | 106.9 | 103.9 | 0.5 |
| N7 | C8 | N9 | s | 110.8 | 116.8 | 113.8 | 0.5 |
| C8 | N9 | C4 | s | 103.4 | 108.2 | 105.8 | 0.4 |
| N9 | C4 | C5 | s | 103.4 | 108.2 | 105.8 | 0.4 |
| N3 | C4 | N9 | s | 122.6 | 132.2 | 127.4 | 0.8 |
| C6 | C5 | N7 | s | 128.1 | 136.5 | 132.3 | 0.7 |
| N1 | C6 | N6 | s | 115.0 | 122.2 | 118.6 | 0.6 |
| C5 | C6 | N6 | s | 118.9 | 128.5 | 123.7 | 0.8 |
| C8 | N9 | C1' | s | 116.9 | 138.5 | 127.7 | 1.8 |
| C4 | N9 | C1' | s | 115.5 | 137.1 | 126.3 | 1.8 |

**Table D12**. Cytosine bond lengths in Angstroms [Å].

| Atom 1 | Atom 2 | Type | Minimum | Maximum | Reference | Standard dev |
|---|---|---|---|---|---|---|
| P | OP1 | s | 1.383 | 1.587 | 1.485 | 0.017 |
| P | OP2 | s | 1.383 | 1.587 | 1.485 | 0.017 |
| P | OP3 | s | 1.535 | 1.679 | 1.607 | 0.012 |
| P | O5' | s | 1.533 | 1.653 | 1.593 | 0.010 |
| O5' | C5' | (P)O5'-C5' | 1.344 | 1.536 | 1.440 | 0.016 |
| O5' | C5' | (H)C2'-endo | 1.328 | 1.520 | 1.424 | 0.016 |
| O5' | C5' | (H)C3'-endo | 1.366 | 1.474 | 1.420 | 0.009 |
| C5' | C4' | C2'-endo | 1.437 | 1.581 | 1.509 | 0.012 |
| C5' | C4' | C3'-endo | 1.466 | 1.550 | 1.508 | 0.007 |
| C4' | C3' | C2'-endo | 1.461 | 1.593 | 1.527 | 0.011 |
| C4' | C3' | C3'-endo | 1.461 | 1.581 | 1.521 | 0.010 |
| C3' | C2' | C2'-endo | 1.459 | 1.591 | 1.525 | 0.011 |
| C3' | C2' | C3'-endo | 1.457 | 1.589 | 1.523 | 0.011 |
| C2' | C1' | C2'-endo | 1.478 | 1.574 | 1.526 | 0.008 |
| C2' | C1' | C3'-endo | 1.463 | 1.595 | 1.529 | 0.011 |
| O4' | C1' | C2'-endo | 1.343 | 1.487 | 1.415 | 0.012 |
| O4' | C1' | C3'-endo | 1.334 | 1.490 | 1.412 | 0.013 |
| O4' | C4' | C2'-endo | 1.394 | 1.514 | 1.454 | 0.010 |
| O4' | C4' | C3'-endo | 1.373 | 1.529 | 1.451 | 0.013 |
| O3' | C3' | C2'-endo | 1.355 | 1.499 | 1.427 | 0.012 |
| O3' | C3' | C3'-endo | 1.333 | 1.501 | 1.417 | 0.014 |
| C1' | N1 | C2'-endo | 1.380 | 1.548 | 1.464 | 0.014 |
| C1' | N1 | C3'-endo | 1.393 | 1.573 | 1.483 | 0.015 |
| C2' | O2' | C2'-endo | 1.334 | 1.490 | 1.412 | 0.013 |
| C2' | O2' | C3'-endo | 1.360 | 1.480 | 1.420 | 0.010 |
| C2 | O2 | s | 1.186 | 1.294 | 1.240 | 0.009 |
| C4 | N4 | s | 1.281 | 1.389 | 1.335 | 0.009 |
| N1 | C2 | s | 1.337 | 1.457 | 1.397 | 0.010 |
| N1 | C6 | s | 1.331 | 1.403 | 1.367 | 0.006 |
| C2 | N3 | s | 1.305 | 1.401 | 1.353 | 0.008 |
| N3 | C4 | s | 1.293 | 1.377 | 1.335 | 0.007 |
| C4 | C5 | s | 1.377 | 1.473 | 1.425 | 0.008 |
| C5 | C6 | s | 1.291 | 1.387 | 1.339 | 0.008 |

**Table D13**. Cytosine bond angles in degrees [ ° ].

| Atom 1 | Atom 2 | Atom 3 | Type | Minimum | Maximum | Reference | Standard dev |
|---|---|---|---|---|---|---|---|
| OP1 | P | OP2 | s | 110.6 | 128.6 | 119.6 | 1.5 |
| O5' | P | OP1 | large | 103.5 | 117.9 | 110.7 | 1.2 |
| O5' | P | OP1 | small | 100.3 | 111.1 | 105.7 | 0.9 |
| O5' | P | OP2 | large | 103.5 | 117.9 | 110.7 | 1.2 |
| O5' | P | OP2 | small | 100.3 | 111.1 | 105.7 | 0.9 |
| O5' | C5' | C4' | (P)O5'-C5'-C4' | 104.6 | 114.2 | 109.4 | 0.8 |
| O5' | C5' | C4' | C2'-endo | 100.3 | 123.1 | 111.7 | 1.9 |
| O5' | C5' | C4' | C3'-endo | 101.9 | 121.1 | 111.5 | 1.6 |
| P | O5' | C5' | s | 111.3 | 130.5 | 120.9 | 1.6 |
| O4' | C4' | C3' | C2'-endo | 101.3 | 110.9 | 106.1 | 0.8 |
| O4' | C4' | C3' | C3'-endo | 98.0 | 110.0 | 104.0 | 1.0 |
| C5' | C4' | C3' | C2'-endo | 106.8 | 123.6 | 115.2 | 1.4 |
| C5' | C4' | C3' | C3'-endo | 106.4 | 125.6 | 116.0 | 1.6 |
| C5' | C4' | O4' | C2'-endo | 101.9 | 116.3 | 109.1 | 1.2 |
| C5' | C4' | O4' | C3'-endo | 104.4 | 115.2 | 109.8 | 0.9 |
| C1' | O4' | C4' | C2'-endo | 105.5 | 113.9 | 109.7 | 0.7 |
| C1' | O4' | C4' | C3'-endo | 105.1 | 114.7 | 109.9 | 0.8 |
| C4' | C3' | O3' | C2'-endo | 96.8 | 122.0 | 109.4 | 2.1 |
| C4' | C3' | O3' | C3'-endo | 101.0 | 125.0 | 113.0 | 2.0 |
| C2' | C3' | O3' | C2'-endo | 96.3 | 122.7 | 109.5 | 2.2 |
| C2' | C3' | O3' | C3'-endo | 104.1 | 123.3 | 113.7 | 1.6 |
| C4' | C3' | C2' | C2'-endo | 96.6 | 108.6 | 102.6 | 1.0 |
| C4' | C3' | C2' | C3'-endo | 96.6 | 108.6 | 102.6 | 1.0 |
| C3' | C2' | C1' | C2'-endo | 96.7 | 106.3 | 101.5 | 0.8 |
| C3' | C2' | C1' | C3'-endo | 97.1 | 105.5 | 101.3 | 0.7 |
| O4' | C1' | C2' | C2'-endo | 99.8 | 111.8 | 105.8 | 1.0 |
| O4' | C1' | C2' | C3'-endo | 102.2 | 113.0 | 107.6 | 0.9 |
| N1 | C1' | C2' | C2'-endo | 106.2 | 121.8 | 114.0 | 1.3 |
| N1 | C1' | C2' | C3'-endo | 105.4 | 118.6 | 112.0 | 1.1 |
| O4' | C1' | N1 | C2'-endo | 103.4 | 113.0 | 108.2 | 0.8 |
| O4' | C1' | N1 | C3'-endo | 104.3 | 112.7 | 108.5 | 0.7 |
| C1' | C2' | O2' | s | 92.6 | 128.6 | 110.6 | 3.0 |
| C3' | C2' | O2' | s | 95.9 | 130.7 | 113.3 | 2.9 |
| C6 | N1 | C2 | s | 117.9 | 122.7 | 120.3 | 0.4 |
| N1 | C2 | N3 | s | 115.0 | 123.4 | 119.2 | 0.7 |
| C2 | N3 | C4 | s | 116.9 | 122.9 | 119.9 | 0.5 |
| N3 | C4 | C5 | s | 119.5 | 124.3 | 121.9 | 0.4 |
| C4 | C5 | C6 | s | 114.4 | 120.4 | 117.4 | 0.5 |
| C5 | C6 | N1 | s | 118.0 | 124.0 | 121.0 | 0.5 |
| N1 | C2 | O2 | s | 115.3 | 122.5 | 118.9 | 0.6 |
| N3 | C2 | O2 | s | 117.7 | 126.1 | 121.9 | 0.7 |
| N3 | C4 | N4 | s | 113.8 | 122.2 | 118.0 | 0.7 |
| C5 | C4 | N4 | s | 116.0 | 124.4 | 120.2 | 0.7 |
| C6 | N1 | C1' | s | 113.6 | 128.0 | 120.8 | 1.2 |
| C2 | N1 | C1' | s | 112.2 | 125.4 | 118.8 | 1.1 |

**Table D14**. Guanine bond lengths in Angstroms [Å].

| Atom 1 | Atom 2 | Type | Minimum | Maximum | Reference | Standard dev |
|---|---|---|---|---|---|---|
| P | OP1 | s | 1.383 | 1.587 | 1.485 | 0.017 |
| P | OP2 | s | 1.383 | 1.587 | 1.485 | 0.017 |
| P | OP3 | s | 1.535 | 1.679 | 1.607 | 0.012 |
| P | O5' | s | 1.533 | 1.653 | 1.593 | 0.010 |
| O5' | C5' | (P)O5'-C5' | 1.344 | 1.536 | 1.440 | 0.016 |
| O5' | C5' | (H)C2'-endo | 1.328 | 1.520 | 1.424 | 0.016 |
| O5' | C5' | (H)C3'-endo | 1.366 | 1.474 | 1.420 | 0.009 |
| C5' | C4' | C2'-endo | 1.437 | 1.581 | 1.509 | 0.012 |
| C5' | C4' | C3'-endo | 1.466 | 1.550 | 1.508 | 0.007 |
| C4' | C3' | C2'-endo | 1.461 | 1.593 | 1.527 | 0.011 |
| C4' | C3' | C3'-endo | 1.461 | 1.581 | 1.521 | 0.010 |
| C3' | C2' | C2'-endo | 1.459 | 1.591 | 1.525 | 0.011 |
| C3' | C2' | C3'-endo | 1.457 | 1.589 | 1.523 | 0.011 |
| C2' | C1' | C2'-endo | 1.478 | 1.574 | 1.526 | 0.008 |
| C2' | C1' | C3'-endo | 1.463 | 1.595 | 1.529 | 0.011 |
| O4' | C1' | C2'-endo | 1.343 | 1.487 | 1.415 | 0.012 |
| O4' | C1' | C3'-endo | 1.334 | 1.490 | 1.412 | 0.013 |
| O4' | C4' | C2'-endo | 1.394 | 1.514 | 1.454 | 0.010 |
| O4' | C4' | C3'-endo | 1.373 | 1.529 | 1.451 | 0.013 |
| O3' | C3' | C2'-endo | 1.355 | 1.499 | 1.427 | 0.012 |
| O3' | C3' | C3'-endo | 1.333 | 1.501 | 1.417 | 0.014 |
| C1' | N9 | C2'-endo | 1.380 | 1.548 | 1.464 | 0.014 |
| C1' | N9 | C3'-endo | 1.393 | 1.573 | 1.483 | 0.015 |
| C2' | O2' | C2'-endo | 1.334 | 1.490 | 1.412 | 0.013 |
| C2' | O2' | C3'-endo | 1.360 | 1.480 | 1.420 | 0.010 |
| N1 | C2 | s | 1.325 | 1.421 | 1.373 | 0.008 |
| C2 | N3 | s | 1.275 | 1.371 | 1.323 | 0.008 |
| N3 | C4 | s | 1.308 | 1.392 | 1.350 | 0.007 |
| C4 | C5 | s | 1.337 | 1.421 | 1.379 | 0.007 |
| C5 | C6 | s | 1.359 | 1.479 | 1.419 | 0.010 |
| C6 | N1 | s | 1.349 | 1.433 | 1.391 | 0.007 |
| C5 | N7 | s | 1.352 | 1.424 | 1.388 | 0.006 |
| N7 | C8 | s | 1.269 | 1.341 | 1.305 | 0.006 |
| C8 | N9 | s | 1.332 | 1.416 | 1.374 | 0.007 |
| N9 | C4 | s | 1.327 | 1.423 | 1.375 | 0.008 |
| C2 | N2 | s | 1.281 | 1.401 | 1.341 | 0.010 |
| C6 | O6 | s | 1.183 | 1.291 | 1.237 | 0.009 |

**Table D15**. Guanine bond angles in degrees [ ° ].

| Atom 1 | Atom 2 | Atom 3 | Type | Minimum | Maximum | Reference | Standard dev |
|---|---|---|---|---|---|---|---|
| OP1 | P | OP2 | s | 110.6 | 128.6 | 119.6 | 1.5 |
| O5' | P | OP1 | large | 103.5 | 117.9 | 110.7 | 1.2 |
| O5' | P | OP1 | small | 100.3 | 111.1 | 105.7 | 0.9 |
| O5' | P | OP2 | large | 103.5 | 117.9 | 110.7 | 1.2 |
| O5' | P | OP2 | small | 100.3 | 111.1 | 105.7 | 0.9 |
| O5' | C5' | C4' | (P)O5'-C5'-C4' | 104.6 | 114.2 | 109.4 | 0.8 |
| O5' | C5' | C4' | C2'-endo | 100.3 | 123.1 | 111.7 | 1.9 |
| O5' | C5' | C4' | C3'-endo | 101.9 | 121.1 | 111.5 | 1.6 |
| P | O5' | C5' | s | 111.3 | 130.5 | 120.9 | 1.6 |
| O4' | C4' | C3' | C2'-endo | 101.3 | 110.9 | 106.1 | 0.8 |
| O4' | C4' | C3' | C3'-endo | 98.0 | 110.0 | 104.0 | 1.0 |
| C5' | C4' | C3' | C2'-endo | 106.8 | 123.6 | 115.2 | 1.4 |
| C5' | C4' | C3' | C3'-endo | 106.4 | 125.6 | 116.0 | 1.6 |
| C5' | C4' | O4' | C2'-endo | 101.9 | 116.3 | 109.1 | 1.2 |
| C5' | C4' | O4' | C3'-endo | 104.4 | 115.2 | 109.8 | 0.9 |
| C1' | O4' | C4' | C2'-endo | 105.5 | 113.9 | 109.7 | 0.7 |
| C1' | O4' | C4' | C3'-endo | 105.1 | 114.7 | 109.9 | 0.8 |
| C4' | C3' | O3' | C2'-endo | 96.8 | 122.0 | 109.4 | 2.1 |
| C4' | C3' | O3' | C3'-endo | 101.0 | 125.0 | 113.0 | 2.0 |
| C2' | C3' | O3' | C2'-endo | 96.3 | 122.7 | 109.5 | 2.2 |
| C2' | C3' | O3' | C3'-endo | 104.1 | 123.3 | 113.7 | 1.6 |
| C4' | C3' | C2' | C2'-endo | 96.6 | 108.6 | 102.6 | 1.0 |
| C4' | C3' | C2' | C3'-endo | 96.6 | 108.6 | 102.6 | 1.0 |
| C3' | C2' | C1' | C2'-endo | 96.7 | 106.3 | 101.5 | 0.8 |
| C3' | C2' | C1' | C3'-endo | 97.1 | 105.5 | 101.3 | 0.7 |
| O4' | C1' | C2' | C2'-endo | 99.8 | 111.8 | 105.8 | 1.0 |
| O4' | C1' | C2' | C3'-endo | 102.2 | 113.0 | 107.6 | 0.9 |
| N9 | C1' | C2' | C2'-endo | 106.2 | 121.8 | 114.0 | 1.3 |
| N9 | C1' | C2' | C3'-endo | 105.4 | 118.6 | 112.0 | 1.1 |
| O4' | C1' | N9 | C2'-endo | 103.4 | 113.0 | 108.2 | 0.8 |
| O4' | C1' | N9 | C3'-endo | 104.3 | 112.7 | 108.5 | 0.7 |
| C1' | C2' | O2' | s | 92.6 | 128.6 | 110.6 | 3.0 |
| C3' | C2' | O2' | s | 95.9 | 130.7 | 113.3 | 2.9 |
| C6 | N1 | C2 | s | 121.5 | 128.7 | 125.1 | 0.6 |
| N1 | C2 | N3 | s | 120.3 | 127.5 | 123.9 | 0.6 |
| C2 | N3 | C4 | s | 108.9 | 114.9 | 111.9 | 0.5 |
| N3 | C4 | C5 | s | 125.6 | 131.6 | 128.6 | 0.5 |
| C4 | C5 | C6 | s | 115.2 | 122.4 | 118.8 | 0.6 |
| C5 | C6 | N1 | s | 108.5 | 114.5 | 111.5 | 0.5 |
| C4 | C5 | N7 | s | 108.4 | 113.2 | 110.8 | 0.4 |
| C5 | N7 | C8 | s | 101.3 | 107.3 | 104.3 | 0.5 |
| N7 | C8 | N9 | s | 110.1 | 116.1 | 113.1 | 0.5 |
| C8 | N9 | C4 | s | 104.0 | 108.8 | 106.4 | 0.4 |
| N9 | C4 | C5 | s | 103.0 | 107.8 | 105.4 | 0.4 |
| N3 | C4 | N9 | s | 122.4 | 129.6 | 126.0 | 0.6 |
| C6 | C5 | N7 | s | 126.8 | 134.0 | 130.4 | 0.6 |
| N1 | C2 | N2 | s | 110.8 | 121.6 | 116.2 | 0.9 |
| N3 | C2 | N2 | s | 115.7 | 124.1 | 119.9 | 0.7 |
| N1 | C6 | O6 | s | 116.3 | 123.5 | 119.9 | 0.6 |
| C5 | C6 | O6 | s | 125.0 | 132.2 | 128.6 | 0.6 |
| C8 | N9 | C1' | s | 119.2 | 134.8 | 127.0 | 1.3 |
| C4 | N9 | C1' | s | 118.7 | 134.3 | 126.5 | 1.3 |

**Table D16**. Uracil bond lengths in Angstroms [Å].

| Atom 1 | Atom 2 | Type | Minimum | Maximum | Reference | Standard dev |
|---|---|---|---|---|---|---|
| P | OP1 | s | 1.383 | 1.587 | 1.485 | 0.017 |
| P | OP2 | s | 1.383 | 1.587 | 1.485 | 0.017 |
| P | OP3 | s | 1.535 | 1.679 | 1.607 | 0.012 |
| P | O5' | s | 1.533 | 1.653 | 1.593 | 0.010 |
| O5' | C5' | (P)O5'-C5' | 1.344 | 1.536 | 1.440 | 0.016 |
| O5' | C5' | (H)C2'-endo | 1.328 | 1.520 | 1.424 | 0.016 |
| O5' | C5' | (H)C3'-endo | 1.366 | 1.474 | 1.420 | 0.009 |
| C5' | C4' | C2'-endo | 1.437 | 1.581 | 1.509 | 0.012 |
| C5' | C4' | C3'-endo | 1.466 | 1.550 | 1.508 | 0.007 |
| C4' | C3' | C2'-endo | 1.461 | 1.593 | 1.527 | 0.011 |
| C4' | C3' | C3'-endo | 1.461 | 1.581 | 1.521 | 0.010 |
| C3' | C2' | C2'-endo | 1.459 | 1.591 | 1.525 | 0.011 |
| C3' | C2' | C3'-endo | 1.457 | 1.589 | 1.523 | 0.011 |
| C2' | C1' | C2'-endo | 1.478 | 1.574 | 1.526 | 0.008 |
| C2' | C1' | C3'-endo | 1.463 | 1.595 | 1.529 | 0.011 |
| O4' | C1' | C2'-endo | 1.343 | 1.487 | 1.415 | 0.012 |
| O4' | C1' | C3'-endo | 1.334 | 1.490 | 1.412 | 0.013 |
| O4' | C4' | C2'-endo | 1.394 | 1.514 | 1.454 | 0.010 |
| O4' | C4' | C3'-endo | 1.373 | 1.529 | 1.451 | 0.013 |
| O3' | C3' | C2'-endo | 1.355 | 1.499 | 1.427 | 0.012 |
| O3' | C3' | C3'-endo | 1.333 | 1.501 | 1.417 | 0.014 |
| C1' | N1 | C2'-endo | 1.380 | 1.548 | 1.464 | 0.014 |
| C1' | N1 | C3'-endo | 1.393 | 1.573 | 1.483 | 0.015 |
| C2' | O2' | C2'-endo | 1.334 | 1.490 | 1.412 | 0.013 |
| C2' | O2' | C3'-endo | 1.360 | 1.480 | 1.420 | 0.010 |
| C2 | O2 | s | 1.165 | 1.273 | 1.219 | 0.009 |
| C4 | O4 | s | 1.184 | 1.280 | 1.232 | 0.008 |
| N1 | C2 | s | 1.327 | 1.435 | 1.381 | 0.009 |
| N1 | C6 | s | 1.321 | 1.429 | 1.375 | 0.009 |
| C2 | N3 | s | 1.331 | 1.415 | 1.373 | 0.007 |
| N3 | C4 | s | 1.326 | 1.434 | 1.380 | 0.009 |
| C4 | C5 | s | 1.377 | 1.485 | 1.431 | 0.009 |
| C5 | C6 | s | 1.283 | 1.391 | 1.337 | 0.009 |

**Table D17**. Uracil bond angles in degrees [ ° ].

| Atom 1 | Atom 2 | Atom 3 | Type | Minimum | Maximum | Reference | Standard dev |
|---|---|---|---|---|---|---|---|
| OP1 | P | OP2 | s | 110.6 | 128.6 | 119.6 | 1.5 |
| O5' | P | OP1 | large | 103.5 | 117.9 | 110.7 | 1.2 |
| O5' | P | OP1 | small | 100.3 | 111.1 | 105.7 | 0.9 |
| O5' | P | OP2 | large | 103.5 | 117.9 | 110.7 | 1.2 |
| O5' | P | OP2 | small | 100.3 | 111.1 | 105.7 | 0.9 |
| O5' | C5' | C4' | (P)O5'-C5'-C4' | 104.6 | 114.2 | 109.4 | 0.8 |
| O5' | C5' | C4' | C2'-endo | 100.3 | 123.1 | 111.7 | 1.9 |
| O5' | C5' | C4' | C3'-endo | 101.9 | 121.1 | 111.5 | 1.6 |
| P | O5' | C5' | s | 111.3 | 130.5 | 120.9 | 1.6 |
| O4' | C4' | C3' | C2'-endo | 101.3 | 110.9 | 106.1 | 0.8 |
| O4' | C4' | C3' | C3'-endo | 98.0 | 110.0 | 104.0 | 1.0 |
| C5' | C4' | C3' | C2'-endo | 106.8 | 123.6 | 115.2 | 1.4 |
| C5' | C4' | C3' | C3'-endo | 106.4 | 125.6 | 116.0 | 1.6 |
| C5' | C4' | O4' | C2'-endo | 101.9 | 116.3 | 109.1 | 1.2 |
| C5' | C4' | O4' | C3'-endo | 104.4 | 115.2 | 109.8 | 0.9 |
| C1' | O4' | C4' | C2'-endo | 105.5 | 113.9 | 109.7 | 0.7 |
| C1' | O4' | C4' | C3'-endo | 105.1 | 114.7 | 109.9 | 0.8 |
| C4' | C3' | O3' | C2'-endo | 96.8 | 122.0 | 109.4 | 2.1 |
| C4' | C3' | O3' | C3'-endo | 101.0 | 125.0 | 113.0 | 2.0 |
| C2' | C3' | O3' | C2'-endo | 96.3 | 122.7 | 109.5 | 2.2 |
| C2' | C3' | O3' | C3'-endo | 104.1 | 123.3 | 113.7 | 1.6 |
| C4' | C3' | C2' | C2'-endo | 96.6 | 108.6 | 102.6 | 1.0 |
| C4' | C3' | C2' | C3'-endo | 96.6 | 108.6 | 102.6 | 1.0 |
| C3' | C2' | C1' | C2'-endo | 96.7 | 106.3 | 101.5 | 0.8 |
| C3' | C2' | C1' | C3'-endo | 97.1 | 105.5 | 101.3 | 0.7 |
| O4' | C1' | C2' | C2'-endo | 99.8 | 111.8 | 105.8 | 1.0 |
| O4' | C1' | C2' | C3'-endo | 102.2 | 113.0 | 107.6 | 0.9 |
| N1 | C1' | C2' | C2'-endo | 106.2 | 121.8 | 114.0 | 1.3 |
| N1 | C1' | C2' | C3'-endo | 105.4 | 118.6 | 112.0 | 1.1 |
| O4' | C1' | N1 | C2'-endo | 103.4 | 113.0 | 108.2 | 0.8 |
| O4' | C1' | N1 | C3'-endo | 104.3 | 112.7 | 108.5 | 0.7 |
| C1' | C2' | O2' | s | 92.6 | 128.6 | 110.6 | 3.0 |
| C3' | C2' | O2' | s | 95.9 | 130.7 | 113.3 | 2.9 |
| C6 | N1 | C2 | s | 117.4 | 124.6 | 121.0 | 0.6 |
| N1 | C2 | N3 | s | 111.3 | 118.5 | 114.9 | 0.6 |
| C2 | N3 | C4 | s | 123.4 | 130.6 | 127.0 | 0.6 |
| N3 | C4 | C5 | s | 111.0 | 118.2 | 114.6 | 0.6 |
| C4 | C5 | C6 | s | 116.1 | 123.3 | 119.7 | 0.6 |
| C5 | C6 | N1 | s | 119.7 | 125.7 | 122.7 | 0.5 |
| N1 | C2 | O2 | s | 118.6 | 127.0 | 122.8 | 0.7 |
| N3 | C2 | O2 | s | 118.0 | 126.4 | 122.2 | 0.7 |
| N3 | C4 | O4 | s | 115.2 | 123.6 | 119.4 | 0.7 |
| C5 | C4 | O4 | s | 122.3 | 129.5 | 125.9 | 0.6 |
| C6 | N1 | C1' | s | 112.8 | 129.6 | 121.2 | 1.4 |
| C2 | N1 | C1' | s | 110.5 | 124.9 | 117.7 | 1.2 |

**Table D18**. Polymer linkage values, bond lengths in Angstroms [Å] and angles in degrees [ ° ].

| Atom 1 | Atom 2 | Atom 3 | Type | Minimum | Maximum | Reference | Standard dev |
|---|---|---|---|---|---|---|---|
| Bond length | | | | | | | |
| O3' | P | | s | 1.535 | 1.679 | 1.607 | 0.012 |
| Angles | | | | | | | |
| C3' | O3' | P | s | 112.5 | 126.9 | 119.7 | 1.2 |
| O3' | P | O5' | s | 92.6 | 115.4 | 104.0 | 1.9 |
| O3' | P | OP2 | large | 103.9 | 117.1 | 110.5 | 1.1 |
| O3' | P | OP2 | small | 92.0 | 118.4 | 105.2 | 2.2 |
| O3' | P | OP1 | large | 103.9 | 117.1 | 110.5 | 1.1 |
| O3' | P | OP1 | small | 92.0 | 118.4 | 105.2 | 2.2 |

**Table D19**. Dihedral angles for nucleic acids.

| Atom 1 | Atom 2 | Atom 3 | Atom 4 | Type | Minimum | Maximum | Reference | Standard dev |
|---|---|---|---|---|---|---|---|---|
| O3' | P | C5' | | | 226.5 | 344.1 | 285.3 | 9.8 |
| | | | | | 8.4 | 153.6 | 81.0 | 12.1 |
| P | O5' | C5' | C4' | | 105.5 | 261.5 | 183.5 | 13.0 |
| O5' | C5' | C4' | C3' | | 18.3 | 86.7 | 52.5 | 5.7 |
| | | | | | 141.0 | 217.8 | 179.4 | 6.4 |
| | | | | | 219.1 | 6.7 | 292.9 | 12.3 |
| C4' | C3' | O3' | P | | 162.4 | 265.6 | 214.0 | 8.6 |
| C3' | O3' | P | O5' | | 260.4 | 318.0 | 289.2 | 4.8 |
| | | | | | 354.9 | 166.5 | 80.7 | 14.3 |
| Sugar | | | | | | | | |
| C5' | C4' | C3' | O3' | C2'-endo | 117.9 | 176.7 | 147.3 | 4.9 |
| O4' | C4' | C3' | O3' | C2'-endo | 236.3 | 299.9 | 268.1 | 5.3 |
| O4' | C1' | C2' | C3' | C2'-endo | 14.8 | 55.6 | 35.2 | 3.4 |
| C1' | C2' | C3' | C4' | C2'-endo | 307.8 | 341.4 | 324.6 | 2.8 |
| C2' | C3' | C4' | O4' | C2'-endo | 357.8 | 50.6 | 24.2 | 4.4 |
| C3' | C4' | O4' | C1' | C2'-endo | 323.5 | 31.9 | 357.7 | 5.7 |
| C4' | O4' | C1' | C2' | C2'-endo | 308.0 | 10.4 | 339.2 | 5.2 |
| C5' | C4' | C3' | C2' | C2'-endo | 238.8 | 288.0 | 263.4 | 4.1 |
| O3' | C3' | C2' | O2' | C2'-endo | 294.5 | 344.9 | 319.7 | 4.2 |
| C4' | O4' | C1' | N1/9 | C2'-endo | 177.5 | 257.9 | 217.7 | 6.7 |
| O4' | C1' | N1 | C2 | C2'-endo | 119.4 | 340.2 | 229.8 | 18.4 |
| O4' | C1' | N9 | C4' | C2'-endo | 91.2 | 22.8 | 237.0 | 24.3 |
| C5' | C4' | C3' | O3' | C3'-endo | 54.6 | 107.4 | 81.0 | 4.4 |
| O4' | C4' | C3' | O3' | C3'-endo | 176.6 | 227.0 | 201.8 | 4.2 |
| O4' | C1' | C2' | C3' | C3'-endo | 306.0 | 4.8 | 335.4 | 4.9 |
| C1' | C2' | C3' | C4' | C3'-endo | 19.1 | 52.7 | 35.9 | 2.8 |
| C2' | C3' | C4' | O4' | C3'-endo | 306.1 | 343.3 | 324.7 | 3.1 |
| C3' | C4' | O4' | C1' | C3'-endo | 349.9 | 51.1 | 20.5 | 5.1 |
| C4' | O4' | C1' | C2' | C3'-endo | 326.2 | 39.4 | 2.8 | 6.1 |
| C5' | C4' | C3' | C2' | C3'-endo | 185.4 | 222.6 | 204.0 | 3.1 |
| O3' | C3' | C2' | O2' | C3'-endo | 17.3 | 71.3 | 44.3 | 4.5 |
| C4' | O4' | C1' | N1/9 | C3'-endo | 202.4 | 280.4 | 241.4 | 6.5 |
| O4' | C1' | N1 | C2 | C3'-endo | 156.1 | 235.3 | 195.7 | 6.6 |
| O4' | C1' | N9 | C4 | C3'-endo | 109.3 | 277.3 | 193.3 | 14.0 |
